# Wiring between close nodes in biological networks evolves more quickly than between distant nodes

**DOI:** 10.1101/2023.05.23.541989

**Authors:** Alejandro Gil-Gomez, Joshua S. Rest

## Abstract

As species diverge, a wide range of evolutionary processes lead to changes in protein-protein interaction networks and metabolic networks. The rate at which biological networks evolve is an important question in evolutionary biology. Previous empirical work has focused on interactomes from model organisms to calculate rewiring rates, but this is limited by the relatively small number of species and sparse nature of network data across species. We present a proxy for variation in network topology: variation in drug-drug interactions (DDIs), obtained by studying drug combinations (DCs) across taxa. Here, we propose the rate at which DDIs change across species as an estimate of the rate at which the underlying biological network changes as species diverge. We computed the evolutionary rates of DDIs using previously published data from a high throughput study in gram-negative bacteria. Using phylogenetic comparative methods, we found that DDIs diverge rapidly over short evolutionary time periods, but that divergence saturates over longer time periods. In parallel, we mapped drugs with known targets in protein-protein interaction and co-functional networks. We found that the targets of synergistic DDIs are closer in these networks than other types of DCs and that synergistic interactions have a higher evolutionary rate, meaning that nodes that are closer evolve at a faster rate. Future studies of network evolution may use DC data to gain larger-scale perspectives on the details of network evolution within and between species.

## Introduction

Biological networks are models of real molecular interactions in the cell. These networks are built by collecting biochemical and genetic interaction data, an approach that has improved in the last decades with the advent of modern high-throughput methods. However, there are still many limitations to the gathering and analysis of biological networks, specifically for studying network variation across and within species. This is because many biological networks have poor quality or are sparse and incomplete and because the number of organisms for which network data is available is still very limited (Cusick et al. 2005; Jin et al. 2013; Ghadie et al. 2018). The major types of biological networks that have been built represent protein interactions, metabolic processes, signaling, and gene regulation.

Biological networks evolve as nodes (e.g. proteins) and edges (e.g. molecular interactions) are added or lost. This can be the result of different processes, including those affecting genes and the interaction of their products, such as gene or motif duplication, loss, horizontal transfer, or whole genome duplication (Wagner 2003; Cork and Purugganan 2004; Bernhardsson et al. 2011; Koch et al. 2017), or a result of processes affecting quantitative properties, including non-synonymous substitutions in the protein nodes affecting their function (Jensen 1976; Ghadie et al. 2018). These mutations may affect the binding of ligands, protein domains, or DNA motifs (Koch et al. 2017), which in turn result in changes in their associated metabolic, signaling, or gene expression networks. The type of network also may affect the rewiring rates of nodes. For example, gene regulatory networks tend to rewire at faster rates than metabolic networks (Shou et al. 2011); suggesting that some network types are less constrained than others.

The genetic and evolutionary events that remove old connections and generate new ones may be random with neutral consequences (Bernhardsson et al. 2011), may be genetically constrained (Wollenberg Valero 2020), or may affect fitness. We know that not all motifs are equally abundant in biological networks (Picard et al. 2008), suggesting the action of natural selection. Natural selection may remove deleterious connections (Jordan et al. 2008) and favor advantageous ones (Laarits et al. 2016; Mehta et al. 2021), including balancing environmental robustness with network functions (Han et al. 2013; Chen and Ho 2014). While higher fitness solutions may include those with lower levels of modularity (Kashtan and Alon 2005), modularity itself may be driven by selection to reduce connection costs (Clune et al. 2013). Higher levels of protein connectivity have been correlated with lower substitution rates (Fraser et al. 2002), potentially due to strong purifying selection acting on the interfacial sites of interacting proteins (Zotenko et al. 2008). Each of the network types, discussed in aggregate above, may be subject to different evolutionary forces and patterns.

Part of the key to understanding the forces acting on networks is a characterization of evolutionary rates of network change. Despite advances in studying the connections of individual nodes and network rewiring, the rate and patterns by which quantitative differences between networks accumulate as a function of species divergence remain relatively unexplored. Beltrao and Serrano (2007) estimated rewiring rates among eukaryotic proteins at ∼10^−5^ interactions/protein pair/million years, found that network rewiring proceeds faster than sequence evolution, and that differences in rewiring rates between functional categories suggest the action of natural selection. One interesting question is whether network rewiring among a given pair of nodes is correlated with metrics of connectivity between those nodes, including minimum path length (the number of steps along the shortest path between nodes), K-edge connectivity (minimum number of edges that can be removed to disconnect two nodes), degree (number of edges connected to the node), and centrality. This is important information for understanding what aspects of network structure that we can quantitate are subject to evolutionary forces acting on variation.

In this study, we use drug-drug interaction (DDI) scores as a quantitative proxy measurement of inter-node connectivity, since it has been shown that DDIs are partially dependent on the underlying network topology between targets (Lehár et al. 2007; Yeh et al. 2009). Among drug combinations (DCs), DDIs occur when the effect of two or more drugs is significantly stronger or weaker than their expected combined effect, respectively named synergies and antagonisms (Greco et al. 1995). DDIs are used in the development of novel pharmacological treatments with higher efficiencies at lower doses and to reduce the evolution of drug resistance (Cowen and Steinbach 2008). The reasons why a DDI occurs are varied, but it has been shown that a DDI can occur as a result of factors such as the topology of the underlying network between drug targets, or the essentiality of the metabolites blocked by the perturbation in combination with the inhibition efficiency of the drug on the drug target (Yeh et al. 2009). For example, synergies can occur if drugs act on parallel pathways, where the individual effect of each drug is small and an alternative pathway can compensate for the effect of a single drug. Alternatively, antagonistic interactions can occur mainly in two different ways: by causing a partial loss of function in two parallel pathways of an essential product, or as a result of drugs acting sequentially along the same pathway. This relation between DDIs and network topology has been further explored using experiments and metabolic flux simulations in yeast, suggesting that specific combinations of perturbations in the network result in different quantifiable interaction scores (Lehár et al. 2007). More recently, it has been shown that most synergistic interactions are the result of drugs targeting the same cellular process, while antagonistic interactions are the result of drugs targeting different processes (Brochado et al. 2018).

Only a few studies have explored interspecific variation of DDI scores (Spitzer et al. 2011; Robbins et al. 2015; Brochado et al. 2018; Davis et al. 2022), but they have demonstrated that DC experiments (used to obtain DDI scores) could be scalable in the number of species and strains. A high throughput study of DDIs in gram-negative bacteria has shown that synergies are more conserved across species than antagonisms and additive combinations (Brochado et al. 2018).

How well does the evolution of DDI scores transmit the evolutionary patterns of their underlying biological networks? This is an important question; assuming there is a reasonable correspondence, then our questions about DDI evolution mirror those about biological networks: Does DDI score divergence accumulate linearly with divergence time, or follow some other function? Do DDIs diverge consistently with neutral processes acting on the underlying network structure? Or is there heterogeneity across the network, for example with differences accumulating slowly in constrained local neighborhoods, and faster between more distant connections? In other words, does the evolutionary rate of DDIs depend on the connectivity of the drug targets?

Another way to parse this latter question is to divide up drug combinations by type. Do synergistic interactions evolve at a slower rate than other types of drug interactions, and do antagonistic interactions evolve at a faster rate? These differences in rates would be a result of synergistic interactions taking place in local neighborhoods of nodes, while antagonistic interactions act across distant network neighborhoods, which may evolve more slowly due to greater network redundancy between distant nodes. Indeed, if we were to map DCs to protein targets (when known), are the drug targets of synergistic drug interactions closer in the network than additive and antagonistic interactions? And, more generally, are more closely connected nodes subject to higher rates of evolution than more distantly related nodes? These questions can be addressed by comparing rates of DDI score evolution with measures of inter-node connectivity.

To address these questions, we used the most complete available dataset of DCs measured across species and strains (Brochado et al. 2018). We modeled these DCs under a phylogenetic comparative framework and applied a multivariate Brownian motion model to estimate the evolutionary rate of interaction scores for different clusters of DCs in six strains (three species) of gram-negative bacteria. We also mapped DCs to their putative protein targets to evaluate them in known biological networks. We show that DDI scores can be used as an effective proxy to evaluate macroevolutionary patterns of network evolution.

## Results

### DDI scores diverge non-linearly

We obtained DDI scores (drug-drug interaction Bliss scores) from a previous study (Brochado et al. 2018) that assessed the effect of 2655 pairwise combinations of 79 different compounds on six strains, two from each of the three gammaproteobacteria species: *Escherichia coli, Salmonella enterica*, and *Pseudomonas aeruginosa*. Hierarchical clustering (UPGMA) of the Euclidean distances among DDI scores from the six strains revealed a ‘DDI score distance’ tree (**Fig. 1B**). This is half the cophenetic distance for each strain pair, the distance from either tip of the pair to the point in the tree where they first come together.

**Figure 1:**
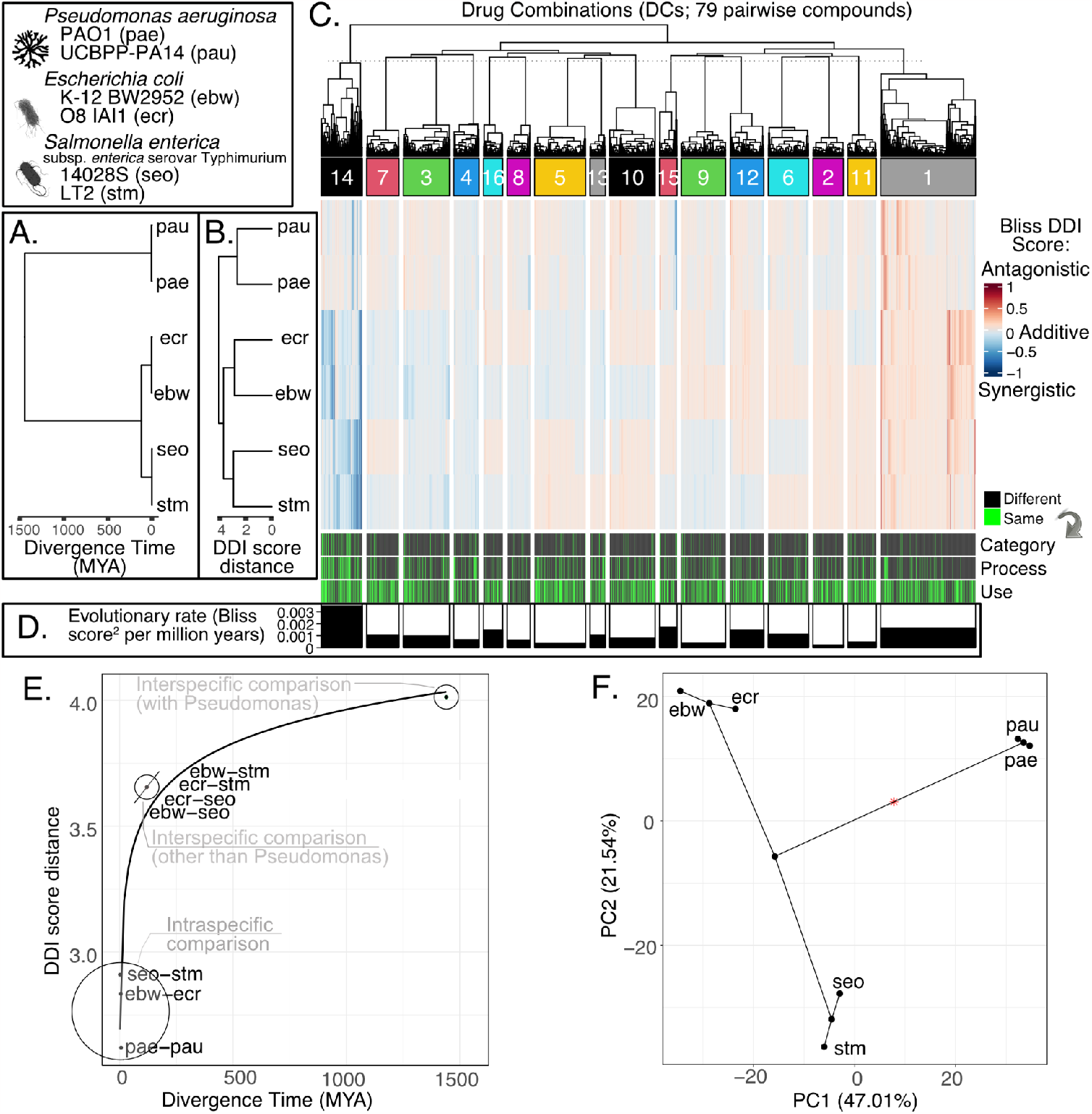
Drug-drug interaction (DDI) score distance diverges non-linearly over time. Species and strain abbreviations are shown in the top left box. **A**. Bayesian phylogeny and divergence time estimates based on an alignment of 27 highly conserved protein sequences. **B**. Hierarchical clustering of strains based on average Euclidean distances across DDIs (i.e. “DDI score distance”); drug combination (DC) data is from Brochado *et al. (2018)*. **C**. Heatmap of DDI scores across strains. Along top, hierarchical clustering of DCs is shown based on Euclidean distances across strains, but these clusters were constrained by t-SNE cluster membership (see text). In the heatmap, synergistic interactions are blue (close to −1), antagonistic interactions are red (close to 1), and additivities are white (close to 0; measured in Bliss score units). The bars below indicate whether the two drugs involved in the interaction are the same (green) or different (black) in terms of: belonging to the same drug category, targeting the same cellular process, or having the same use. **D**. Evolutionary rate of DDI score change, calculated for each cluster. **E**. Pairwise DDI score distances between strains as a function of divergence time between strains. Intraspecific comparisons, comparisons between *Salmonella* and *Escherichia*, and comparisons with *Pseudomonas* are each labeled. **F**. First two axes of a phylomorphospace-PCA for the DDI score data. The percent of variance explained for principal components 1 and 2 are shown.

We also estimated phylogenetic divergence times between the strains and species using calibrated phylogenetic analysis of highly conserved proteins (**Fig. 1A**). The 95% highest probability density of the most recent common ancestor (MRCA) of these three species is 1359-1527 million years ago (MYA), or 100-151 MYA for the two *Enterobacteriaceae* (*E. coli* and *S. enterica*).

To test how DDI scores diverge as a function of species divergence, we regressed the DDI score distances between strains against their divergence times (time to MRCA in MYA). There is a positive correlation between DDI score distance and divergence time (**Fig. 1E**). This is a log-linear relationship (R^2^ =0.93, p-value=6.8×10^−9^; **Sup. Fig. S1**). Specifically, the initial stages of DDI score divergence is characterized by rapid changes that accumulate between closely related strains, the rate of divergence slows down when comparing more distant species, and it saturates among the most distantly related species. This pattern may be consistent with both neutral and selective forces acting on the biological networks underlying DDIs (see Discussion). Indeed, a phylomorphospace plot of the first two axes of a principal components analysis of the DC data (**Fig. 1F**) does not reflect the branch lengths of the DDI distance tree (**Fig. 1B)**, suggesting that DDI score variance is explained by more factors than just time. Here, the largest variance component seems to distinguish all three species from each other (PC1, 47%), while the second component (PC2, 21.5%) seems to distinguish between *Salmonella* and the other species (**Fig. 1F**). This highlights that while the divergence time between *Salmonella* and *Escherichia* is small (**Fig. 1A**), the DDI score distance between these species is comparatively much larger, suggesting that after the divergence from their common ancestor, each lineage rapidly accumulated cellular and biochemical differences that are reflected in their differing DDI scores.

In light of these observations and to further explore these possibilities, we sought to estimate the evolutionary rate of DDI score change for particular DCs. However, there is low power with only six tips to estimate rate shifts for any individual DC. To address this limitation, we used t-SNE to detect clusters of DCs that behave similarly across species, allowing for increased power within each cluster. In order to find DC clusters that are the most biologically relevant from among a large parameter space (after filtering solutions), we first filtered for the top five solutions with the highest degree of phylogenetic modular signal (Adams et al. 2016). From among those five, we chose the solution with the biggest differences in evolutionary rate among clusters, when fitted with a multivariate Brownian motion model. A detailed description of the filters and tests used to select the clusters is in the Methods.

The resulting solution, with a perplexity of 245 and 16 clusters (**Fig. 1C**), was used for the remainder of the analysis. Cluster 14 had the highest evolutionary rate measured using the Brownian motion standard deviation parameter *σ*^2^, with a value of 0.00347 bliss score^2^ per million years, while all the other clusters had rates ranging from 0.00021 to 0.00169 (**Fig. 1D**). Cluster 15 had the second highest rate with a value of 0.00169 bliss score^2^ per million years, and cluster 1 had the third highest rate with a value of 0.00161 bliss score^2^ per million years.

### Synergistic DDIs have the shortest distance between network targets, followed by additive DCs and antagonistic DDIs

We observed that highly synergistic DDIs across all species tend to occur when both drugs belong to the same chemical category and target the same cellular process, as can be seen from the accumulation of green bars for category and process across cluster 14 (**Fig. 1C**; in agreement with (Brochado et al. 2018). This motivated us to formally examine how DCs map onto biological networks. We therefore leave discussion of DDI evolution for now, to first describe our mapping of the DC targets to known biological networks, and how connectivity on these networks relates to combination type (antagonism, additivity, synergy); in the next section we will describe how rates of DDI evolution map onto combination types and network connectivity. We were able to identify protein targets for 39 of the 79 drugs tested by Brochado *et al*. (2018) (see Methods). These drugs had a total of 27 target proteins as identified by their unique protein IDs in *E. coli*. The most common target category was bacterial penicillin-binding protein, a group involved in the biosynthesis of bacterial cell walls. Other target categories mapped include ribosomal RNA, DNA polymerase, topoisomerase, thymidylate synthase, and mitochondrial glycerol-3-phosphate (**Sup. Table S1**).

We then examined the *E. coli* DDI scores as a function of the connectivity of their targets. We used two *E. coli* networks: (1) a positive gold-standard co-functional gene pair network of *E. coli* (**Fig. 2A**), and (2) a small/medium scale protein-protein interaction (PPI) network (**Fig. 2B**), both available at EcoliNet *(Kim et al*. *2015)*. The co-functional network includes data integrated from several sources and contains information about genes and their products that are linked in molecular and metabolic pathways and processes, or linked because they are involved in the same or overlapping biological processes, or co-regulated. The PPI network was derived from curated PPI databases, and contains high-confidence interactions (and their co-functional links) between pairs of *E. coli* proteins.

**Figure 2:**
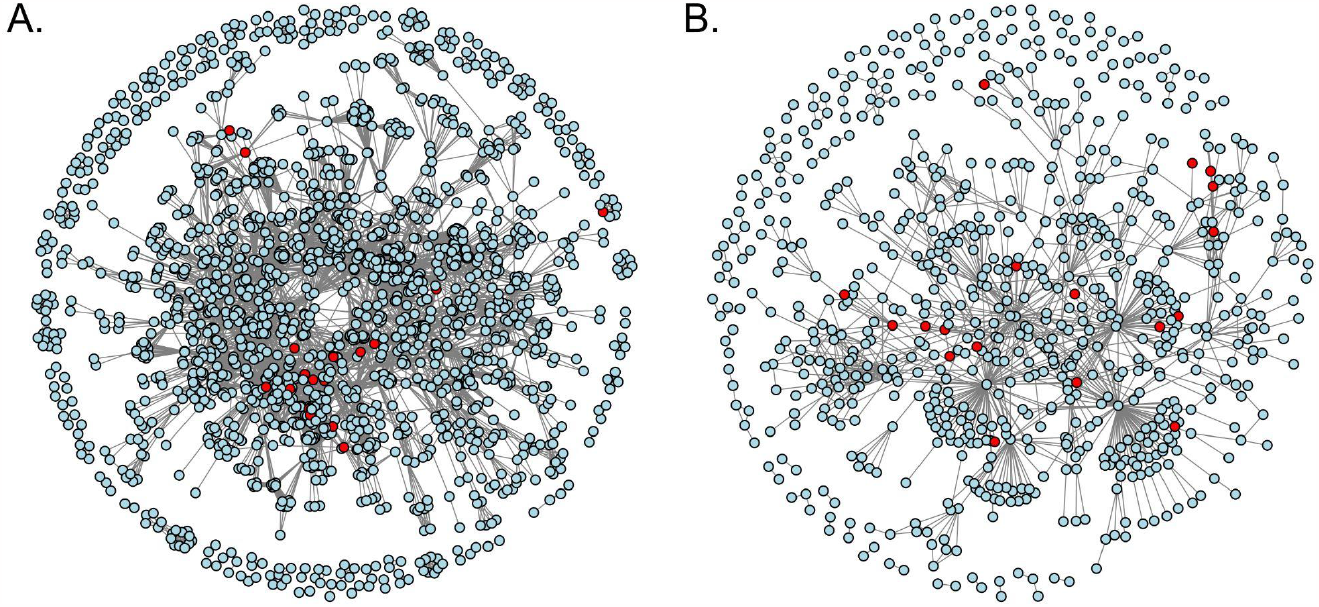
**A**. Graphical representations of the co-functional gene pair network of *E. coli* (EcoliNet: EcoCyc/GO-BP). This network contains 1835 nodes with an average path length of 4.8 and contains 20 proteins targeted by 36 drugs in our analysis. **B**. Graphical representations of the PPI network of *E. coli*, as determined by small and medium-scale experiments (EcoliNet: LC. Small/medium-scale PPI). This network contains 764 nodes with an average path length of 4.9, with 18 proteins targeted by 27 drugs in our analysis. In **A** and **B**, each node represents a unique protein (KEGG ID) in *E. coli*; the red nodes are target proteins identified as participating in DCs in our analysis.

We measured the length of minimum distance paths between all drug targets in the networks. Targets of synergistic DCs have a significantly shorter minimum path length than antagonistic and additive DCs in both networks (**Fig. 3A, B**). This result is consistent with our expectations, given that synergistic DDIs tend to be more common between drugs that target the same cellular process and thus should be closer to each other in biological networks (Brochado et al. 2018). We next examined K-edge connectivity, where two nodes are K-edge-connected if after removing k edges or less the nodes remain connected. For the co-functional network, K-edge connectivity is higher for targets of synergistic DCs than additive DCs (Wilcoxon p-value=0.0032), and higher for targets of additive DCs than antagonistic DCs (Wilcoxon p-value=0.014) (**Fig. 3C**), as expected if targets of synergistic DCs are closer together. There was no significant difference among combination types for the PPI network, although targets of synergistic and additive DCs do have significantly higher connectivity than a background set of non-targets (**Fig. 3D**). Overall, these results suggest that, the closer together and better connected two nodes are to each other, the more likely they are associated with a synergistic DDI.

**Figure 3:**
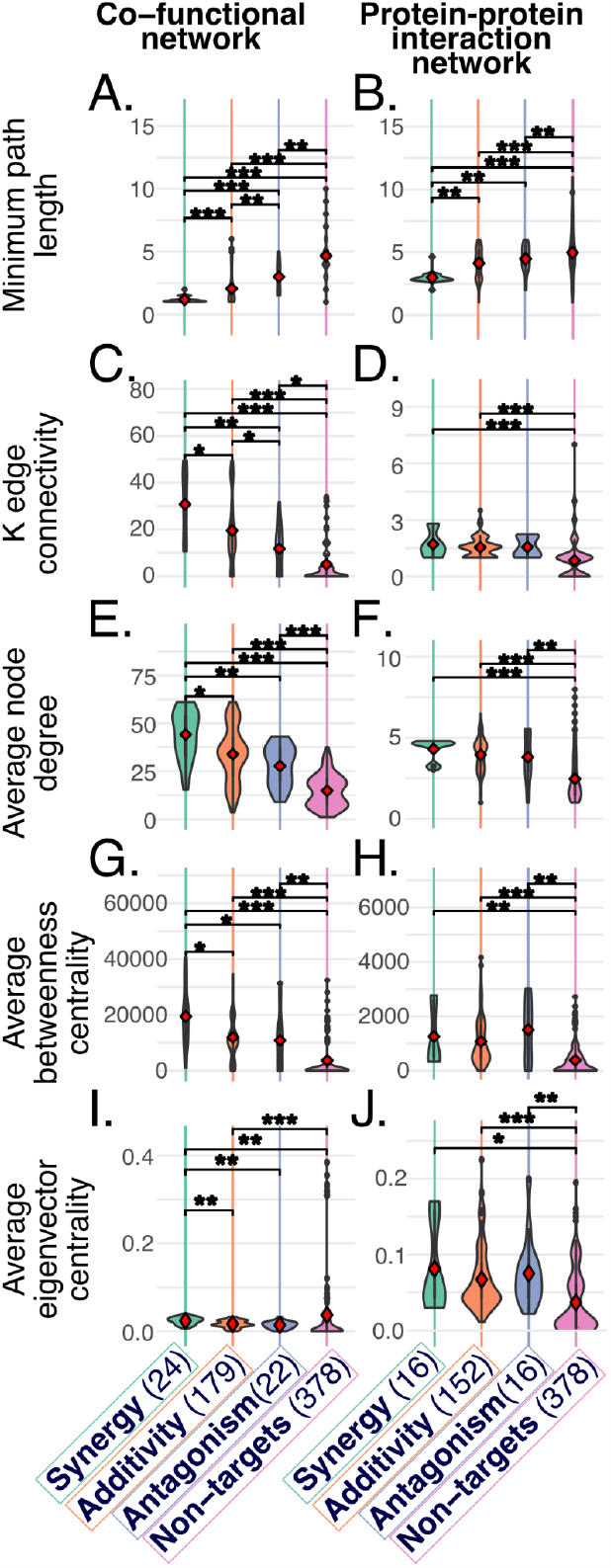
Pairs of protein targets with synergistic drug-drug interactions (DDIs) have lower path length and higher connectivity and centrality measures in *E. coli* co-functional (**A**,**C**,**E**,**G**,**I**) and protein-protein interaction (**B**,**D**,**F**,**H**,**J**) networks. DDI types examined are synergies, additivities, and antagonisms (“non-targets” are a background sample of non-drug target proteins in the network). The number of interactions in each category is in parentheses. Network metrics are: **A-B**: path length between two targets, **C-D**: K-edge connectivity between two targets, **E-F**: mean node degree of the two targets, **G-H**: mean betweenness centrality of the two targets, **I-J**: mean eigenvector centrality of the two targets. In all plots, significance of Wilcoxon test p-values are given for differences in the mean between all pairwise comparisons: p-value < *0.05, **0.0001, ***0.00001. For all plots, Kruskal−Wallis test for difference among groups is p < 1×10^−7^.

We also found that proteins in the co-functional network that are associated with synergistic DDIs are more central and better connected than nodes that are associated with other types of interactions. In the PPI network, nodes that are associated with any type of DC are more central and better connected than nodes that aren’t targeted in our DC set, but don’t differ by combination type. We arrived at these conclusions by examining three metrics (node degree, betweenness centrality, eigenvector centrality) that are characteristics of individual nodes (we averaged between the two target nodes in the DC). Average node degree (the number of connected edges) is significantly higher for targets of synergistic DCs than additive (Wilcoxon p-value=0.0024) or antagonistic DCs (Wilcoxon p-value=0.00018) in the co-functional network (**Fig. 3E**), but not in the PPI network (**Fig. 3F**). Average betweenness centrality (how much each node lies on the shortest paths between pairs of other nodes in the network) is higher for targets associated with synergistic DCs than with additive (Wilcoxon p-value=0.0015) or antagonistic DCs (Wilcoxon p-value=0.013) in the co-functional network (**Fig. 3G**), but not in the PPI network (**Fig. 3H**). Average eigenvector centrality (how much each node is connected to other important nodes in the network) was also higher for targets associated with in synergistic than additive (Wilcoxon p-value=0.0002) or antagonistic DCs (Wilcoxon p-value=0.0005) in the co-functional network (**Fig. 3I**), but these differences were not significant in the PPI network (**Fig. 3J**). Together, these results suggest that network connectivity of a protein affects the likelihood that it will be associated with a DDI, as well as the type of DDI.

### Synergistic DDIs evolve faster than additive and antagonistic DDIs

We return now to describing the evolutionary rates of DDI scores (**Fig. 1D**), as determined on a per-cluster basis (via t-SNE, described above). Cluster 14, which had the highest evolutionary rate (*σ*^2^=0.00347), appears to be rich in synergies in *E. coli* and *Salmonella*, while *Pseudomonas* has more additive DCs in this cluster (**Fig. 1B**). In contrast, cluster 15, which was the second highest rate cluster (*σ*^2^=0.00169), appears to be rich in synergies in *Pseudomonas* while the DCs in *E. coli* and *Salmonella* are primarily additive. The cluster with the third highest rate, cluster 1 (*σ*^2^=0.00161), contains highly antagonistic DDIs in all the species. These observations suggest that the evolutionary rate of DDI scores may vary as a function of the combination (DDI) type and network connectivity of their targets. To more formally investigate this, we examined how the rates vary based on the targets’ connectivity (in *E. coli*) and type of interaction. None of the DCs are exclusively antagonistic across all strains and species, although we found all other combinations (i.e., additivity, synergy, additivity⟷antagonism, additivity⟷synergy, antagonism⟷synergy and additivity⟷antagonism⟷synergy, where the arrow indicates that some strains or species have one DDI type and other strains or species have another DDI type).

Overall, synergistic DDIs and additive⟷synergistic DDIs have faster evolutionary rates than any other class (**Fig. 4A**; all possible group comparisons were significantly different from each other except additivity⟷antagonism⟷synergy vs. additivity⟷antagonism, and excluding antagonism⟷synergy, with only a single observation; see **Sup. Table S2** for all statistical tests in **Fig. 4**; significant differences mentioned have p-value < 0.001 unless indicated). We also aggregated each target pair’s DDI type as a simple sum of its DDI types across strains and species (+1 for synergies, 0 for additivities, −1 for antagonisms), and found again that more synergistic DDIs (sum ≤ −3) have faster evolutionary rates than additivities (3 > sum > −3) and antagonisms (sum ≥ 3) (**Fig. 4B**). This result of higher evolutionary rates for synergistic DDIs is also true for the targets that are in both the PPI and co-functional networks described earlier (**Fig. 4C**; all categories are significantly different, except for additivities vs antagonisms in the PPI network).

**Figure 4:**
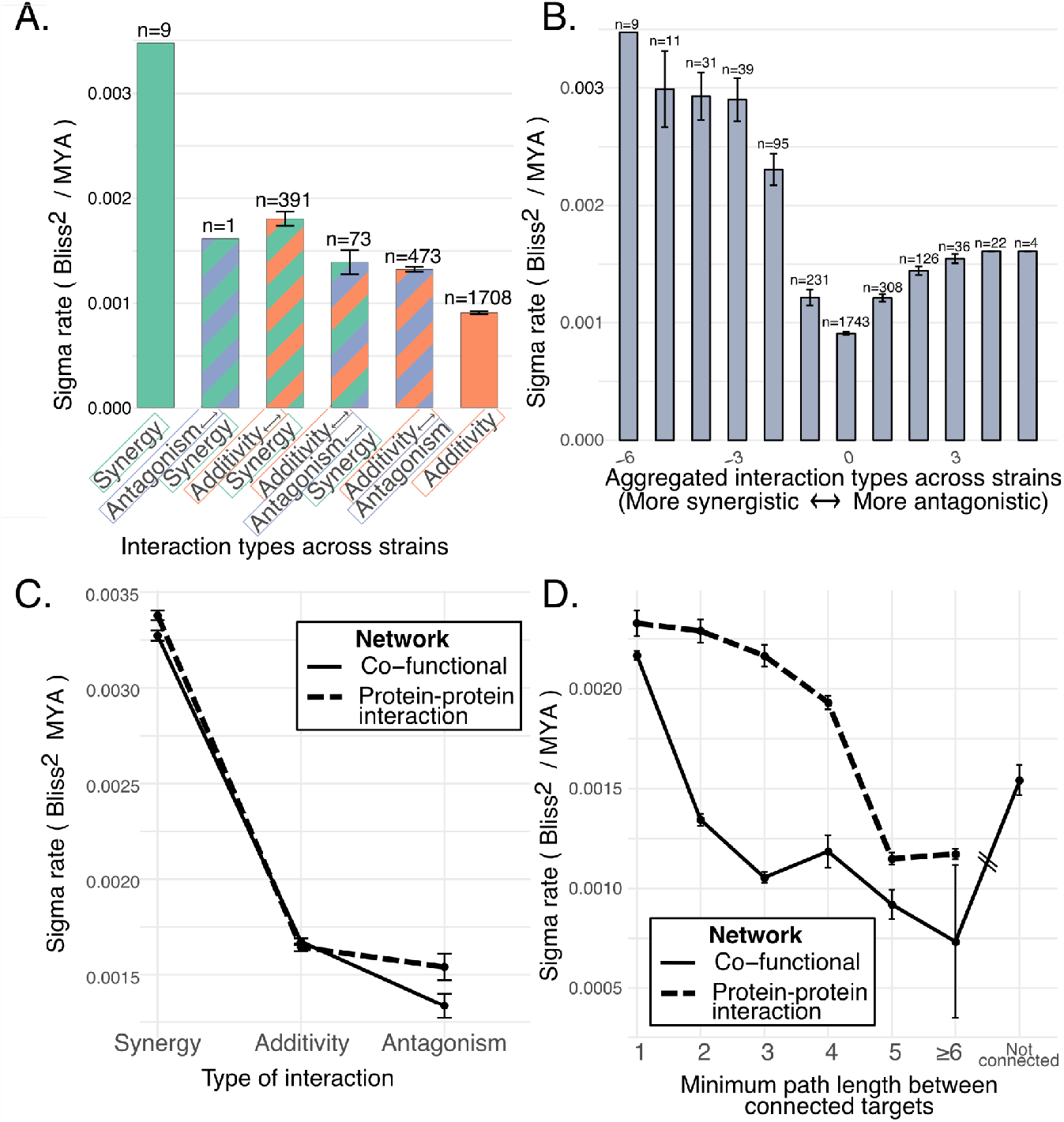
Evolutionary rates of DDI scores as a function of DDI type and network connectivity of targets. **A**. DCs resulting in only synergy or synergy⟷antagonism DDIs across strains have faster evolutionary rates. **B**. Aggregated interaction types reveal that more synergistic DCs have higher evolutionary rates. The X-axis value is the sum, per DC, across strains, where DDIs are scored as −1 for synergies, 0 for additive, and +1 for antagonistic interactions. **C**. Rate of DDI evolution as a function of DDI type, for DCs with protein targets in the co-functional and small/medium scale PPI networks. Synergistic DCs have higher evolutionary rates than additive and antagonistic DCs (combination types and networks from *E. coli*). **D**. Rate of DDI evolution as a function of the minimum distance between DC target proteins reveals that wiring between close nodes in biological networks evolves more quickly than between distant nodes. See **Sup. Table S2** for statistical tests for differences among groups shown here. For all plots, the error bars represent the standard error of the mean.

Furthermore, DCs whose targets are nearby in the network have the highest rate of DDI score evolution (**Fig. 4D**). This is consistent with the fact that cluster 14 has the highest rate (**Fig. 1D**), and contains a majority of synergies (**Fig. 1C**). This result of nearby targets having the highest DDI evolution rates is significant for both the PPI network (p-value=0.007) and the co-functional network (p-value=0.019). The evolutionary rate is also significantly faster when simply comparing DCs that target adjacent vs. non-adjacent targets in the PPI network (p-value=0.0049; **Sup. Fig. S2B**), and for the EcoCyc portion of the co-functional network (**Sup. Fig. S2A**), and when comparing connected vs. disconnected targets for this latter network (**Sup. Fig. S2C**).

Protein pairs with the highest K-edge connectivity (>20) are targets of DCs with higher DDI score evolutionary rates in the co-functional network (but not the PPI network, where all targets have connectivities < 5; **Sup. Fig. S2K**). Interestingly, DCs that have evolved between additive and synergistic DDI types tend to have targets with greater connectivity, degree, and betweenness centrality values in the *E. coli* networks (**Sup. Fig. S2D**,**E**,**F**,**G**). Together, these results indicate that, as the distance and connectivity between two targets increase (as measured, for example, by their minimum connecting path length), the average evolutionary rate of the DDI scores decreases. We interpret this to mean that wiring between close nodes in biological networks evolves faster than between distant nodes.

## Discussion

### Overview

We studied the evolution of drug-drug interaction scores as a proxy for studying the evolution of network topology. We found that the evolutionary rates of DDIs among gram-negative bacteria are initially high, lower at longer evolutionary distances, and plateau at the largest distances. This suggests that chemogenomic variation rapidly accumulates. This helps to explain the observation that therapeutic combination therapies are typically taxon specific; for example, in fungi very few combinations of antifungals act across species (Brown et al. 2014).

We then mapped drugs with known targets to different biological networks in *E. coli* K-12 and found that the targets of synergistic DDIs are closer in biological networks than other types of DDIs, and that synergistic interactions have a higher evolutionary rate, suggesting that connectivity between nodes that are closer in biological networks evolves at a faster rate than between more distant nodes.

### DDI evolution

Examining pairwise distances of DDI scores between species, we observed a rapid accumulation of differences and overall dissimilarity in DDI responses among species. Indeed the DDI score distances curve appears saturated among distantly related species. The high rate of DDI evolution we observed is consistent with the observations from the source paper that most (∼70%) drug interactions are species-specific, and about one fifth are strain-specific (Brochado et al. 2018).

Indeed, it is now generally appreciated that some DDIs show substantial genetic variation (Roemhild et al. 2022). For instance, the interaction between the aminoglycoside tobramycin and different β-lactams can be either synergistic or additive across isolates of multidrug resistant Enterobacteriaceae (Fass 1982). Some interactions between drugs used for treating urinary tract infections are additive across *E. coli* isolates (50%, e.g. mecillinam and ciprofloxacin) or antagonistic across isolates (10%, mecillinam and nitrofurantoin), while two different combination types occur across isolates for other combinations (40%; e.g. mecillinam and trimethoprim) (Fatsis-Kavalopoulos et al. 2020).

We also clustered DDIs by their similarity across strains and species and then evaluated the rate of evolution of these clusters under a Brownian motion model. This approach is analogous to methods used in geometric morphometrics, in which specimen landmarks are grouped to evaluate different features, and modularity tests are used to assess whether there are differences in the evolution rate of different regions of that organ (Smaers and Vanier 2019). Modularity among traits mirrors shared functional, genetic, and developmental pathways (Cheverud 1996). Here, the signal of high modularity among DDI clusters is a result of variance between clusters being greater than the variance within clusters. We observed that the distribution of synergies, additivities and antagonisms among clusters partially explains different rate estimates. Clusters with low rates are composed of mostly additive DDIs; however some of these low-rate estimates are caused by resistance to one or both antibiotics, and a low phylogenetic signal is expected when multiple strains are resistant (Michel et al. 2008). Other clusters with high rates (e.g., clusters 14, 15, and 1) have some highly synergistic or antagonistic interactions in some strains and additivities in other strains.

### Biological network inferences

With these rate estimates and interaction patterns, we asked whether network-based metrics are correlated with DDI divergence, by linking a subset of DCs with their respective protein targets in *E. coli* co-functional (EcoCyc/GO-BP) and PPI networks. We hypothesize that the rate of evolution of a DDI reflects the evolution of underlying connections between the protein targets of the DC.

We first made several observations about the *E. coli* DDI types and network: synergistic DDIs occur when the targets are closer in the network, while additive and antagonistic DDIs target, on average, more distant proteins (**Fig. 3A, B**). This result is not surprising given the fact that synergies are expected to be more prevalent in drug combinations that target the same pathway or that belong to the same chemical class (Cowen and Steinbach 2008; Yeh et al. 2009), however, we think this is the first time this pattern has been shown using network data. We also found that synergistic interactions are more prevalent among targets that are part of the same cellular process or functional category, while antagonisms can take place between targets that are in different parts of the cell (Yeh et al. 2006; Wang et al. 2012; Brochado et al. 2018). We also found that targets associated with synergistic DDIs have a higher node degree (for the co-functional network), and K-edge connectivity than targets of DCs with additive or antagonistic DDIs (**Fig. 3C-F**). This result agrees with a previous study that reports on the positive contribution of node degree in synergy prediction (Liu et al. 2022). Average node betweenness and eigenvector centralities were also found to be more significant among targets of DCs with synergistic DDIs compared to additivities and antagonisms in the co-functional network (**Fig. 3G-J**). This result suggests that targets of DCs with synergistic DDIs tend to be “hubs” in the network (Li et al. 2011; Tilli et al. 2016), or that they are more central to different pathways than targets of DCs with antagonistic or additive DDIs.

At large evolutionary scales, network rewiring rates appear to slow down (Shou et al. 2011); our DDI-based estimates also suggest a pattern of saturation of network divergence at the largest evolutionary distances (**Fig. 1E**). Beltrao and Serrano (2007) estimated a single rewiring rate among proteins that is faster than sequence evolution; here, we demonstrated that there is a distribution of evolutionary rates. We examined the rate of evolution of DDIs whose targets are closer in the PPI and co-functional networks and found that their rate of evolution is higher (**Fig. 4D**). This is correlated with our observation that synergies tend to occur when DC targets are closer in the PPI and co-functional networks (**Fig. 3A,B**) and have a high evolutionary rate (**Fig. 4A,B,C)**. As the path length between targets increases, and for more antagonistic DCs, the evolutionary rate of the DDI score decreases (**Fig. 4**). In some cases, we might speculate that the drug pairs themselves acted through natural selection to cause the observed DDI evolution. Indeed, synergistic DDIs are known to increase the rate of resistance evolution because they increase the selective advantage of resistance mutations, while antagonistic DDIs slow the rate of resistance evolution because they reduce the selective advantage of resistance mutations (Hegreness et al. 2008; Michel et al. 2008).

Our observation of differential evolutionary rates among drug interaction types suggests that different biological pathways have different rewiring rates. Different processes (besides the drugs themselves) may be driving this pattern, for instance, positive selection on mutations that cause co-expression or co-localization of components from different networks may result in novel connections between the previously disconnected pathways. Purifying selection against loss of function may also constrain the rewiring of the network.

Beltrao and Serrano (2007) suggested that differences in rewiring rates between functional categories may be attributed to natural selection. In our DDI analyses, we found that there is variation in functional process annotations among the subset of known target proteins of drugs in clusters 2, 3, 4, 6 and 16 (**Sup. Fig. S6**). These clusters all showed enrichment for anion binding (GO:0043168) and small molecule binding (GO:0036094), while clusters 3, 6 and 16 are also enriched for ion binding (GO:0043167). These results suggest that differences in rates of DDI evolution may be partially explained by differences in natural selection acting on different underlying network components. However, next, we highlight a different type of bias in DDI data that may contribute to our observed signal and cause difficulty in the interpretation of DDI data.

### Network vs. resistance evolution

Another important explanation for our observed signal (high evolutionary rate of synergistic, well-connected targets) lies in the fact that some strains are resistant to particular antibiotics. We suspect that some changes in DDIs that we correlated with changes in network structure are actually caused by changes in resistance to one or both drugs, rather than changes in the drug interaction term itself. Changes in resistance are likely based on structural changes in either target proteins, or other resistance mechanisms. These other resistance mechanisms include transporter proteins or cell membrane permeability characteristics, which affect cross-resistance and collateral sensitivity, and don’t represent the types of biological network changes that we hoped to measure with the DDI. Indeed, there is genetic variation in how drug interactions are affected by the order in which drug combinations are administered, suggesting hysteresis (Roemhild et al. 2022). Note that Brochado *et al*. (2018) reported any strain with fitness values (e.g., ratio of drug/non-drug treated growth) of >0.7 as being resistant to that drug. However, some drugs that are known to be clinically bacteriostatic (inhibit growth) fall in this range, and strains are reported to be resistant; e.g. see the case of sulfamonomethoxine resistance in all species and strains, below. Indeed, Bushby (1973) reported “resistance to one of the drugs as measured by conventional tests may not abolish synergy.” To consider the effect of resistance differences on our results, we repeated the entire analysis filtering for DDIs that are “susceptible” (fitness < 0.7) in at least one of the species for both drugs, assuming that this decreases the proportion of true resistance cases in the dataset. This set included 1127/2655 DCs (42.2% of the total DCs). Our results for this smaller set are consistent with the full dataset (**Sup. Figs. S3-S5**). A minor difference in the subset is that antagonisms have a slightly lower (instead of equivalent) rate than additivities. Although this consistency between the full and reduced dataset is desirable, it doesn’t remove the possibility that differences in resistance among strains are a substantial part of the evolutionary signal.

We used DDIs to learn about the evolution of networks, so discussion of some of the individual DDIs illustrate strengths and limitations of the approach. The evolutionary and network parameters also provide some insight into how and why individual DDIs with clinical relevance may vary across species. Next, we discuss these aspects of the interaction between A22 and novobiocin, and sulfamonomethoxine and trimethoprim or erythromycin.

### A22 and novobiocin

The DDI between A22 and novobiocin is a case example (Sup. Table S3). Briefly, A22 binds the ATP-binding domain of MreB (eco:b3251) (Bean et al. 2009), the actin homolog in prokaryotes, while novobiocin binds gyrase subunit B (gyrB, eco:b3699) and topoisomerase IV (parC, eco:b3019), which are two type II topoisomerases involved in DNA unwinding and DNA duplication (Gellert et al. 1976; Hardy and Cozzarelli 2003). MreB and ParC are known to physically interact (Madabhushi and Marians 2009) and are required for chromosome segregation (Huang et al. 1998). A22 and novobiocin have previously been reported to be synergistic in *E. coli* and *P. aeruginosa* (Taylor 2011; Taylor et al. 2012); however, in the dataset from Brochado *et al. (2018)*, the combination type is variable within species. Both *E. coli* strains are susceptible to both drugs, but they have different interaction terms: one is additive, the other synergistic. *Salmonella* also shows intraspecific differences, where one of the strains is synergistic and the other additive, although in this case, the synergistic strain is resistant to novobiocin, with a fitness of 0.85. Both *Pseudomonas* strains are resistant to novobiocin, with fitnesses greater than 0.86, and they have additive DDIs. This high variability of DDI type within species resulted in an evolutionary rate of 0.008 bliss^2^/MYA for this DC, within the cluster with the highest evolutionary rate. The distance between these targets in the co-functional (EcoCyc/GO-BP) network was 2, and in the DDI network the distance between targets was 4. Why are the rates so high among strains, for this DDI? Strain-specific network predictions at string-DB (Szklarczyk et al. 2023) suggest strain-specific differences in local network content. Of the genes all connected to each other in the local network (mreB, gyrA, parC, gyrB), parC is missing in three strains, while a fourth strain has reduced connectivity for MreB (only connected to gyrB). This type of gene content variation suggests the type of intraspecific network variation that may lead to high observed rates of DDI evolution.

### Sulfamonomethoxine and trimethoprim or erythromycin

Another illustrative example of a known DDI used in the clinic and found in our data is the combination of sulfamonomethoxine and trimethoprim (**Sup. Table S3**) (Bushby 1973). These drugs inhibit successive steps in the synthesis of tetrahydrofolic acid synthesis, necessary for the biosynthesis of amino acids, purines, and thymidine. Sulfamonomethoxine competes against the enzymatic substrate of dihydropteroate synthase (folP, eco:b3177); this blocks the production of dihydrofolic acid, which in turn is a substrate for dihydrofolate reductase. Trimethoprim inhibits dihydrofolate reductase (folA, eco:b0048) by competing against the substrate for the binding site of the enzyme. Even on its own, trimethoprim can contribute to “thymineless death” (Then and Angehrn 1973); but while the effect of each drug on its own is bacteriostatic (prevents growth; Ocampo et al. 2014), when used in combination, the combined effect is bactericidal (kills bacteria). This is the cause of the strong synergistic effect between the two drugs (Bushby 1973; Ocampo et al. 2014).

We measured an evolutionary rate of 0.008 bliss^2^/MYA for this DDI, as it belongs to the cluster with the highest evolutionary rate. This rate can be explained by the difference in DDI type between *Pseudomonas* and non-*Pseudomonas* strains. In the data used in our study, the interaction was synergistic for *E. coli* and *S. enterica*, but additive in *Pseudomonas*. The DDI may be additive in *Pseudomonas* because it is resistant to both drugs, due to differences in its permeability to the drugs and the presence of efflux pumps that remove the drugs (Eliopoulos and Huovinen 2001). In contrast, *E. coli* and *S. enterica* were only resistant to a single drug, sulfamonomethoxine..

Our goal in using the sulfamonomethoxine and trimethoprim DDIs to study the network, in this case, was to capture the rate of evolution of the folate biosynthesis pathway across species. The fact that the two targets act as successive steps in all studied species suggests several possible causes for this observed rate. There are some differences in the presence of pathways components that are peripheral to these steps (Pribat et al. 2010), and it is possible that such differences in the network beyond the direct connection contribute to DDI differences. Indeed, beyond their direct link in the co-functional (EcoCyc/GO-BP) network, the targets are highly connected to each other, with a K-edge connectivity of 21 (68% quantile), meaning that 21 edges have to be removed to disconnect the two targets. It is also possible that there are differences in the direct target enzymes and their connectivity (e.g., concentration, kmax). An alternative explanation is resistance: resistance to sulfamonomethoxine and trimethoprim was detected soon after they were introduced in clinical treatments. Several point mutations have been described in the target genes across different bacteria, and the most common mechanism of resistance is due to the bacteria having an extra copy of the target gene (Estrada et al. 2016). All of this suggests that, for sulfamonomethoxine and trimethoprim, evolution of the DDI is the result of many types of biological differences, not just specific differences in network interactions between the drug targets. This is an important limitation to the DDI approach to studying network evolution.

Trimethoprim also interacts with erythromycin, a macrolide that inhibits protein synthesis and has a bacteriostatic effect. This interaction is additive in *E. coli*, but when sulfate is added to the media the combination type becomes suppressive (a type of antagonistic interaction) (Qi et al. 2021).

Trimethoprim may cause sulfur limitation resulting in changes in expression of sulfur reduction genes. Interestingly, this suppressive effect is dependent on the gene crl, and a crl knockout strain eliminates the suppressive response entirely. This is therefore a clearly demonstrated case of genetic variation affecting a DDI. Because *Pseudomonas* doesn’t have crl (Cavaliere et al. 2015) we might have expected to see a high rate of DDI evolution; however Brochado et al (2018) did not add sulfate to the growth media, and thus the interaction between trimethoprim and erythromycin is additive across all species and in a cluster with one of the lowest evolutionary rates (0.0003 bliss^2^/MYA). We predict that, in the presence of sulfate, the rate of this DDI would be much higher (suppressive in non-*Pseudomonas* strains, and additive in *Pseudomonas*), illustrating a case of DDI evolution as a result of network evolution, as well as a strong dependence on environmental conditions.

## Conclusions

Here, we introduce a novel framework to compare evolutionary rates across entire networks. We examined changes in DC effects among strains and species, and compared these rates of change with the DDI types and, when possible, network topology of the drugs’ targets. This approach has some advantages over direct network analyses, such as increasing the number of species under study, without the need of obtaining the underlying network in each species. DDIs can be quantified using high-throughput experiments across different species and strains, and are phenotypic quantitative traits that can be modeled in a phylogenetic comparative framework, allowing for an independent measurement of evolutionary rates between nodes in the network. An important limitation of our approach is that evolution of resistance to a drug may obscure underlying network-based effects.

Still, our approach suggests a general picture of network evolution, where close nodes in the network (which tend to respond synergistically in response to targeting both at once) evolve at faster rates than more distant nodes.

## Materials and Methods

### DDI scores, strain susceptibility, and combination types

We obtained DDI scores (Bliss scores) from a previous study (Brochado et al. 2018) that assessed 2655 combinations (after removing 228 DDIs with missing data) of 79 different compounds on six strains of three species of gram-negative bacteria: *Escherichia coli* K-12 BW2952 (ebw), *E. coli* O8 IAI1 (ecr), *Salmonella enterica* subsp. *enterica* serovar Typhimurium 14028S (seo), *S. enterica* subsp. *enterica* serovar Typhimurium LT2 (stm), *Pseudomonas aeruginosa* PAO1 (pae), *P. aeruginosa* UCBPP-PA14 (pau). We used the following datasets from Brochado (2018): interaction scores from table ED09C as input for our phylogenetic comparative analysis, strain susceptibility to antibiotics from Sup. Table 1 (used for our **Sup. Figs. S3-S5**), and the categories that DDIs belong to from Sup. Table 2.

### Divergence time estimates

We obtained concatenated alignments of 27 highly conserved protein sequences for the six strains using PhySpeTree (Fang et al. 2019). We simultaneously estimated the phylogeny and divegence times in BEAST2 (Bouckaert et al., 2014) using the following approach: We employed one partition for each protein, with linked trees/clocks. A Yule tree model was used in conjunction with an optimized relaxed molecular clock. The following priors were used for calibrating node time estimates: For the divergence time between *E. coli* and *S. enterica*, 100-160 MYA (Vernikos et al. 2007; Meysman et al. 2013; Knöppel et al. 2018). The divergence time between these species and *P. aeruginosa* occurred within the range of 1350.0 to 1527.7 million years ago (Battistuzzi et al. 2004; Blair Hedges and Kumar 2009; Marin et al. 2017). The Markov chain Monte Carlo analysis was run for 20 million iterations.

### Clustering DDIs using t-SNE

To reduce data dimensionality of the DDI trait space, we classified DDIs into clusters using t-SNE in the R packages bigMap (Garriga and Bartumeus 2018) and bigmemory (Kane et al. 2013) with parameters: 80 threads, 80 layers, and 9 rounds. DDIs that were similar to each other across strains were clustered together. The t-SNE analyses were performed for a range of 251 perplexity values between 5 and 2505 (**Sup. Doc. S1**). The clustering output was evaluated based on the stability and plateauing of cost and effect size, and the variance between threads. We obtained stable solutions between perplexity values 175 and 1455. The pakde algorithm was applied with a perplexity of 1/3 the respective t-SNE perplexity. To find which of the clustering solutions was the most modular, we tested for modularity in the data using the function phylo.modularity within the R package geomorph (Adams et al. 2016) The five most modular clustering patterns were selected with the most negative ZCR coefficients. These clustering patterns had the following perplexity values: 355, 245, 275, 215, and 345; and the following number of clusters: 9, 16, 14, 18, and 12. All five clustering patterns had a strong modular signal, with multivariate effect sizes under −26.6, p-value=0.001 and covariance ratios below 0.91. Lastly, we fitted a multivariate Brownian motion model to each of these 5 clustering patterns and calculated the evolutionary rates per cluster using compare.multi.evol.rates, also using the R package geomorph (Adams et al. 2016). Out of the 5 most modular clustering models, the model with a perplexity of 245 was the one with a higher Z effect in the test, showing the biggest differences between groups. Thus, this clustering pattern was used in the following steps. In addition, the sigma rates calculated for each of the clusters were used as approximations for DDI evolutionary rates that are part of that cluster.

### Identification of drug targets and their biological networks

Each drug was identified with unique Pubchem and CHEMBL IDs using webchem (Szöcs et al. 2020), which were then used to retrieve their mode of action from IUPHAR (Armstrong et al. 2019). We also compared our targets to a previously published dataset on drugs and drug targets (Santos et al. 2017), identified unique Uniprot IDs and KO IDs for each target protein, and converted these IDs into *E. coli* Uniprot IDs using the KEGG Orthology (Kanehisa and Goto 2000). We didn’t include drugs whose mode of action was unknown, or that had non-protein molecules as targets, such as small molecules, RNA, or DNA.

Two biological networks of *E. coli* were downloaded from EcoliNet (https://www.inetbio.org/ecolinet/downloadnetwork.php) (Kim et al. 2015), small/medium-scale protein-protein interactions (LC; 764 genes, 1073 links) and the gold-standard co-functional gene pair network of *E. coli* derived from EcoCyc and GO-BP (1835 genes, 10804 links).

### Inter-node network metrics

For each of the networks, we calculated the average path length and node degree distributions. In addition, the minimum distance between each of the nodes in the network was calculated, as well as the node degree (number of connections per node), the K-edge connectivity between each pair of nodes (i.e. the minimum number of edges that can be removed to disconnect the nodes), betweenness centrality (i.e. a measure of centrality in the network based on shortest paths) and eigenvector centrality (i.e. a measure of the influence of the node in the network). We used the R package igraph (Csardi et al. 2006) to calculate these values in each one of the biological networks and for each node or pair of nodes. We also generated an adjacency matrix, which contains information on whether two nodes are connected directly by an edge or not. In addition, we used K-edge connectivity as a proxy for connectedness between proteins (i.e., a pair with K-edge connectivity equal to zero is disconnected, and proteins with K-edge connectivity different than zero are connected).

### Enrichment analysis per cluster

Differential gene set enrichment analysis was performed across the subset of DDIs with known target proteins for each cluster using the R package clusterProfiler ver 4.10.0 (Yu et al. 2012) and org.EcK12.eg.db. The background set included only the known targets. In many cases, drugs from different clusters target the same protein.

## Data and code availability

The code used for data analysis is available from: https://github.com/Alexggo/ddi-netevo

## References

Adams DC, Collyer M, Kaliontzopoulou A, Sherratt E. 2016. geomorph: Software for geometric morphometric analyses. Available from: https://rune.une.edu.au/web/handle/1959.11/21330

Armstrong JF, Faccenda E, Harding SD, Pawson AJ, Southan C, Sharman JL, Campo B, Cavanagh DR, Alexander SPH, Davenport AP, et al. 2019. The IUPHAR/BPS Guide to PHARMACOLOGY in 2020: extending immunopharmacology content and introducing the IUPHAR/MMV Guide to MALARIA PHARMACOLOGY. Nucleic Acids Res.:gkz951.

Battistuzzi FU, Feijao A, Hedges SB. 2004. A genomic timescale of prokaryote evolution: insights into the origin of methanogenesis, phototrophy, and the colonization of land. BMC Evol. Biol. 4:44.

Bean GJ, Flickinger ST, Westler WM, McCully ME, Sept D, Weibel DB, Amann KJ. 2009. A22 Disrupts the Bacterial Actin Cytoskeleton by Directly Binding and Inducing a Low-Affinity State in MreB. Biochemistry 48:4852–4857.

Beltrao P, Serrano L. 2007. Specificity and evolvability in eukaryotic protein interaction networks. PLoS Comput. Biol. 3:e25.

Bernhardsson S, Gerlee P, Lizana L. 2011. Structural correlations in bacterial metabolic networks. BMC Evol. Biol. 11:20.

Blair Hedges S, Kumar S. 2009. The Timetree of Life. OUP Oxford

Brochado AR, Telzerow A, Bobonis J, Banzhaf M, Mateus A, Selkrig J, Huth E, Bassler S, Zamarreño Beas J, Zietek M, et al. 2018. Species-specific activity of antibacterial drug combinations. Nature 559:259–263.

Brown JCS, Nelson J, VanderSluis B, Deshpande R, Butts A, Kagan S, Polacheck I, Krysan DJ, Myers CL, Madhani HD. 2014. Unraveling the Biology of a Fungal Meningitis Pathogen Using Chemical Genetics. Cell 159:1168–1187.

Bushby SR. 1973. Trimethoprim-sulfamethoxazole: in vitro microbiological aspects. J. Infect. Dis. 128:Suppl:442–462 p.

Cavaliere P, Sizun C, Levi-Acobas F, Nowakowski M, Monteil V, Bontems F, Bellalou J, Mayer C, Norel F. 2015. Binding interface between the Salmonella *σ*(S)/RpoS subunit of RNA polymerase and Crl: hints from bacterial species lacking crl. Sci. Rep. 5:13564.

Chen B-S, Ho S-J. 2014. The stochastic evolutionary game for a population of biological networks under natural selection. Evol. Bioinform. Online 10:17–38.

Cheverud JM. 1996. Developmental integration and the evolution of pleiotropy. Am. Zool. 36:44–50.

Clune J, Mouret J-B, Lipson H. 2013. The evolutionary origins of modularity. Proc. Biol. Sci. 280:20122863.

Cork JM, Purugganan MD. 2004. The evolution of molecular genetic pathways and networks. Bioessays 26:479–484.

Cowen LE, Steinbach WJ. 2008. Stress, drugs, and evolution: the role of cellular signaling in fungal drug resistance. Eukaryot. Cell 7:747–764.

Csardi G, Nepusz T, Others. 2006. The igraph software package for complex network research. InterJournal, complex systems 1695:1–9.

Cusick ME, Klitgord N, Vidal M, Hill DE. 2005. Interactome: gateway into systems biology. Hum. Mol. Genet. 14 Spec No. 2:R171–R181.

Davis KP, Morales Y, McCabe AL, Mecsas J, Aldridge BB. 2022. Critical role of growth medium for detecting drug interactions in Gram-negative bacteria that model in vivo responses. bioRxiv [Internet]:2022.09.20.508761. Available from: https://www.biorxiv.org/content/10.1101/2022.09.20.508761

Eliopoulos GM, Huovinen P. 2001. Resistance to Trimethoprim-Sulfamethoxazole. Clin. Infect. Dis. 32:1608–1614.

Estrada A, Wright DL, Anderson AC. 2016. Antibacterial Antifolates: From Development through Resistance to the Next Generation. Cold Spring Harb. Perspect. Med. [Internet] 6. Available from: 10.1101/cshperspect.a028324

Fang Y, Liu C, Lin J, Li X, Alavian KN, Yang Y, Niu Y. 2019. PhySpeTree: an automated pipeline for reconstructing phylogenetic species trees. BMC Evol. Biol. 19:219.

Fass RJ. 1982. Comparative in vitro activities of beta-lactam-tobramycin combinations against Pseudomonas aeruginosa and multidrug-resistant gram-negative enteric bacilli. Antimicrob. Agents Chemother. 21:1003–1006.

Fatsis-Kavalopoulos N, Roemhild R, Tang P-C, Kreuger J, Andersson DI. 2020. CombiANT: Antibiotic interaction testing made easy. PLoS Biol. 18:e3000856.

Fraser HB, Hirsh AE, Steinmetz LM, Scharfe C, Feldman MW. 2002. Evolutionary rate in the protein interaction network. Science 296:750–752.

Garriga J, Bartumeus F. 2018. bigMap: Big Data Mapping with Parallelized t-SNE. arXiv [cs.LG] [Internet]. Available from: http://arxiv.org/abs/1812.09869

Gellert M, O’Dea MH, Itoh T, Tomizawa J. 1976. Novobiocin and coumermycin inhibit DNA supercoiling catalyzed by DNA gyrase. Proc. Natl. Acad. Sci. U. S. A. 73:4474–4478.

Ghadie MA, Coulombe-Huntington J, Xia Y. 2018. Interactome evolution: insights from genome-wide analyses of protein–protein interactions. Curr. Opin. Struct. Biol. 50:42–48.

Greco WR, Bravo G, Parsons JC. 1995. The search for synergy: a critical review from a response surface perspective. Pharmacol. Rev. 47:331–385.

Han HW, Ohn JH, Moon J, Kim JH. 2013. Yin and Yang of disease genes and death genes between reciprocally scale-free biological networks. Nucleic Acids Res. 41:9209–9217.

Hardy CD, Cozzarelli NR. 2003. Alteration of Escherichia coli topoisomerase IV to novobiocin resistance. Antimicrob. Agents Chemother. 47:941–947.

Hegreness M, Shoresh N, Damian D, Hartl D, Kishony R. 2008. Accelerated evolution of resistance in multidrug environments. Proc. Natl. Acad. Sci. U. S. A. 105:13977–13981.

Huang WM, Libbey JL, van der Hoeven P, Yu SX. 1998. Bipolar localization of Bacillus subtilis topoisomerase IV, an enzyme required for chromosome segregation. Proc. Natl. Acad. Sci. U. S. A. 95:4652–4657.

Jensen RA. 1976. Enzyme Recruitment in Evolution of New Function. Annu. Rev. Microbiol. 30:409–425.

Jin Y, Turaev D, Weinmaier T, Rattei T, Makse HA. 2013. The evolutionary dynamics of protein-protein interaction networks inferred from the reconstruction of ancient networks. PLoS One 8:e58134.

Jordan IK, Katz LS, Denver DR, Streelman JT. 2008. Natural selection governs local, but not global, evolutionary gene coexpression networks in Caenorhabditis elegans. BMC Syst. Biol. 2:96.

Kanehisa M, Goto S. 2000. KEGG: kyoto encyclopedia of genes and genomes. Nucleic Acids Res. 28:27–30.

Kane M, Emerson JW, Weston S. 2013. Scalable Strategies for Computing with Massive Data. J. Stat. Softw. 55:1–19.

Kashtan N, Alon U. 2005. Spontaneous evolution of modularity and network motifs. Proc. Natl. Acad. Sci. U. S. A. 102:13773–13778.

Kim H, Shim JE, Shin J, Lee I. 2015. EcoliNet: a database of cofunctional gene network for Escherichia coli. Database [Internet] 2015. Available from: 10.1093/database/bav001

Knöppel A, Knopp M, Albrecht LM, Lundin E, Lustig U, Näsvall J, Andersson DI. 2018. Genetic Adaptation to Growth Under Laboratory Conditions in Escherichia coli and Salmonella enterica. Front. Microbiol. 9:756.

Koch C, Konieczka J, Delorey T, Lyons A, Socha A, Davis K, Knaack SA, Thompson D, O’Shea EK, Regev A, et al. 2017. Inference and Evolutionary Analysis of Genome-Scale Regulatory Networks in Large Phylogenies. Cell Syst 4:543–558.e8.

Laarits T, Bordalo P, Lemos B. 2016. Genes under weaker stabilizing selection increase network evolvability and rapid regulatory adaptation to an environmental shift. J. Evol. Biol. 29:1602–1616.

Lehár J, Zimmermann GR, Krueger AS, Molnar RA, Ledell JT, Heilbut AM, Short GF 3rd, Giusti LC, Nolan GP, Magid OA, et al. 2007. Chemical combination effects predict connectivity in biological systems. Mol. Syst. Biol. 3:80.

Li S, Zhang B, Zhang N. 2011. Network target for screening synergistic drug combinations with application to traditional Chinese medicine. BMC Syst. Biol. 5 Suppl 1:S10.

Liu X, Song C, Liu S, Li M, Zhou X, Zhang W. 2022. Multi-way relation-enhanced hypergraph representation learning for anti-cancer drug synergy prediction. Bioinformatics 38:4782–4789.

Madabhushi R, Marians KJ. 2009. Actin homolog MreB affects chromosome segregation by regulating topoisomerase IV in Escherichia coli. Mol. Cell 33:171–180.

Marin J, Battistuzzi FU, Brown AC, Hedges SB. 2017. The Timetree of Prokaryotes: New Insights into Their Evolution and Speciation. Mol. Biol. Evol. 34:437–446.

Mehta TK, Koch C, Nash W, Knaack SA, Sudhakar P, Olbei M, Bastkowski S, Penso-Dolfin L, Korcsmaros T, Haerty W, et al. 2021. Evolution of regulatory networks associated with traits under selection in cichlids. Genome Biol. 22:25.

Meysman P, Sánchez-Rodríguez A, Fu Q, Marchal K, Engelen K. 2013. Expression divergence between Escherichia coli and Salmonella enterica serovar Typhimurium reflects their lifestyles. Mol. Biol. Evol. 30:1302–1314.

Michel J-B, Yeh PJ, Chait R, Moellering RC Jr, Kishony R. 2008. Drug interactions modulate the potential for evolution of resistance. Proc. Natl. Acad. Sci. U. S. A. 105:14918–14923.

Ocampo PS, Lázár V, Papp B, Arnoldini M, Abel zur Wiesch P, Busa-Fekete R, Fekete G, Pál C, Ackermann M, Bonhoeffer S. 2014. Antagonism between bacteriostatic and bactericidal antibiotics is prevalent. Antimicrob. Agents Chemother. 58:4573–4582.

Picard F, Daudin J-J, Koskas M, Schbath S, Robin S. 2008. Assessing the exceptionality of network motifs. J. Comput. Biol. 15:1–20.

Pribat A, Blaby IK, Lara-Núñez A, Gregory JF 3rd, de Crécy-Lagard V, Hanson AD. 2010. FolX and FolM are essential for tetrahydromonapterin synthesis in Escherichia coli and Pseudomonas aeruginosa. J. Bacteriol. 192:475–482.

Qi Q, Angermayr SA, Bollenbach T. 2021. Uncovering Key Metabolic Determinants of the Drug Interactions Between Trimethoprim and Erythromycin in Escherichia coli. Front. Microbiol. 12:760017.

Robbins N, Spitzer M, Yu T, Cerone RP, Averette AK, Bahn Y-S, Heitman J, Sheppard DC, Tyers M, Wright GD. 2015. An Antifungal Combination Matrix Identifies a Rich Pool of Adjuvant Molecules that Enhance Drug Activity against Diverse Fungal Pathogens. Cell Rep. 13:1481–1492.

Roemhild R, Bollenbach T, Andersson DI. 2022. The physiology and genetics of bacterial responses to antibiotic combinations. Nat. Rev. Microbiol. 20:478–490.

Shou C, Bhardwaj N, Lam HYK, Yan K-K, Kim PM, Snyder M, Gerstein MB. 2011. Measuring the evolutionary rewiring of biological networks. PLoS Comput. Biol. 7:e1001050.

Smaers JB, Vanier DR. 2019. Brain size expansion in primates and humans is explained by a selective modular expansion of the cortico-cerebellar system. Cortex 118:292–305.

Spitzer M, Griffiths E, Blakely KM, Wildenhain J, Ejim L, Rossi L, De Pascale G, Curak J, Brown E, Tyers M, et al. 2011. Cross-species discovery of syncretic drug combinations that potentiate the antifungal fluconazole. Mol. Syst. Biol. 7:499–499.

Szklarczyk D, Kirsch R, Koutrouli M, Nastou K, Mehryary F, Hachilif R, Gable AL, Fang T, Doncheva NT, Pyysalo S, et al. 2023. The STRING database in 2023: protein-protein association networks and functional enrichment analyses for any sequenced genome of interest. Nucleic Acids Res. 51:D638–D646.

Szöcs E, Stirling T, Scott ER, Scharmüller A, Schäfer RB. 2020. webchem: An R Package to Retrieve Chemical Information from the Web. J. Stat. Softw. 93:1–17.

Taylor P. 2011. Combating intrinsic antibiotic resistance in Gram-negative bacteria. Available from: https://macsphere.mcmaster.ca/handle/11375/11329

Taylor PL, Rossi L, De Pascale G, Wright GD. 2012. A forward chemical screen identifies antibiotic adjuvants in Escherichia coli. ACS Chem. Biol. 7:1547–1555.

Then R, Angehrn P. 1973. Nature of the Bactericidal Action of Sulfonamides and Trimethoprim, Alone and in Combination. J. Infect. Dis. 128:S498–S501.

Tilli TM, Carels N, Tuszynski JA, Pasdar M. 2016. Validation of a network-based strategy for the optimization of combinatorial target selection in breast cancer therapy: siRNA knockdown of network targets in MDA-MB-231 cells as an in vitro model for inhibition of tumor development. Oncotarget 7:63189–63203.

Vernikos GS, Thomson NR, Parkhill J. 2007. Genetic flux over time in the Salmonella lineage. Genome Biol. 8:R100.

Wagner A. 2003. How the global structure of protein interaction networks evolves. Proceedings of the Royal Society of London. Series B: Biological Sciences 270:457–466.

Wang Y-Y, Xu K-J, Song J, Zhao X-M. 2012. Exploring drug combinations in genetic interaction network. BMC Bioinformatics 13 Suppl 7:S7.

Wollenberg Valero KC. 2020. Aligning functional network constraint to evolutionary outcomes. BMC Evol. Biol. 20:58.

Yeh PJ, Hegreness MJ, Aiden AP, Kishony R. 2009. Drug interactions and the evolution of antibiotic resistance. Nat. Rev. Microbiol. 7:460–466.

Yeh P, Tschumi AI, Kishony R. 2006. Functional classification of drugs by properties of their pairwise interactions. Nat. Genet. 38:489–494.

Zotenko E, Mestre J, O’Leary DP, Przytycka TM. 2008. Why do hubs in the yeast protein interaction network tend to be essential: reexamining the connection between the network topology and essentiality. PLoS Comput. Biol. 4:e1000140.

