## Supplementary_Figures_and_Tables for "Wiring between close nodes in biological networks evolves more quickly than between distant nodes"

Sup. Table S1. Drug-target mapping
Sup. Table S2. Statistical summaries for Figure 4
- Statistical summary for Fig. 4A
- Statistical summary for Fig. 4B
- Statistical summary for Fig. 4C
- Statistical summary for Fig. 4D

Sup.Table S3. Details of some of the DDIs analyzed.
Sup. Table S4. Network measurements of DDI
Sup. Document S1. t-SNE results of all DDIs for perplexity range between 5 and 2100.
Sup. Fig. S1: DDI-based Euclidean distances between strains scale as a function of phylogenetic distance plotted on a log scale.
Sup. Fig. S2: Evolutionary rates of DDIs vs network metrics of DDI targets in two E coli networks.
Sup. Fig. S3. Heatmap of DDI scores across strains for the subset of DDI that are susceptible in at least one of the species for both drugs.
Sup. Fig. S4. Differences in minimum path length, k edge connectivity, mean node degree, mean betweenness centrality, and mean eigenvector centrality between targets for different types of drug interactions in E. coli for co-functional and PPI networks for the subset of DDI that are susceptible in at least one of the species for both drugs.
Sup. Fig. S5. Relationships among evolutionary rate, interaction type, and path length for the subset of DDI that are susceptible in at least one of the species for both drugs.
Sup. Fig. S6. Dotplot showing significant GO categories of drug targets across clusters. Clusters 3, 6, and 16 have significant enrichment for ion binding. Clusters 2, 3, 4, 6, and 16 are significantly enriched for anion binding and small molecule binding.

**Sup. Table S1. Drug-target mapping**
See separate CSV File.

**Sup. Table S2. Statistical summaries for Figure 4**

| **Statistical summary for Fig. 4A** | | | | | |
|---|---|---|---|---|---|
| **Group 1** | **Group 2** | **statistical test** | **Statistic** | **p-value** | **df** |
| "Additivity-Synergy" | "Additivity-Antagonism-Synergy" | Welch Two Sample t-test | 3.134 | 0.002 | 128.7 |
| "Additivity-Synergy" | "Additivity" | Welch Two Sample t-test | 13.06 | p < 0.001 | 417.249 |
| "Additivity-Synergy" | "Additivity-Antagonism" | Welch Two Sample t-test | 6.735 | p < 0.001 | 486.877 |
| "Additivity-Synergy" | "Synergy" | Welch Two Sample t-test | -24.847 | p < 0.001 | 390 |
| "Additivity-Synergy" | "Antagonism-Synergy" | Not enough observations | NA | NA | NA |
| "Additivity-Antagonism-Synergy" | "Additivity" | Welch Two Sample t-test | 4.214 | p < .001 | 73.759 |
| "Additivity-Antagonism-Synergy" | "Additivity-Antagonism" | Welch Two Sample t-test | 0.581 | 0.526 | 78.427 |
| "Additivity-Antagonism-Synergy" | "Synergy" | Welch Two Sample t-test | -18.39 | p < .001 | 72 |
| "Additivity-Antagonism-Synergy" | "Antagonism-Synergy" | Not enough observations | NA | NA | NA |
| "Additivity" | "Additivity-Antagonism" | Welch Two Sample t-test | -15.407 | p < .001 | 753.83 |
| "Additivity" | "Synergy" | Welch Two Sample t-test | -205.33 | p < .001 | 1707 |
| "Additivity" | "Antagonism-Synergy" | Not enough observations | NA | NA | NA |
| "Additivity-Antagonism" | "Synergy" | Welch Two Sample t-test | -90.677 | p < .001 | 472 |
| "Additivity-Antagonism" | "Antagonism-Synergy" | Not enough observations | NA | NA | NA |
| "Synergy" | "Antagonism-Synergy" | Not enough observations | NA | NA | NA |

| **Statistical summary for Fig. 4B** | | | | | |
|---|---|---|---|---|---|
| **Group 1** | **Group 2** | **statistical test** | **Statistic** | **p-value** | **df** |
| Synergy | Additivity | Welch Two Sample t-test | 18.902 | p < .001 | 117.717 |
| Synergy | Antagonism | Welch Two Sample t-test | -12.37 | p < .001 | 95.955 |
| Additivity | Antagonism | Welch Two Sample t-test | -16.896 | p < .001 | 91.776 |

**Sup. Table S2 (continued). Statistical summaries for Figure 4**

| Statistical summary for Fig. 4C | | | | | | |
|---|---|---|---|---|---|---|
| **Group 1** | **Group 2** | **statistical test** | **Statistic** | **p-value** | **df** | **Network** |
| Synergy | Additivity | Welch Two Sample t-test | 48.985 | p < .001 | 1542.953 | Co-functional (EcoCyc/GO-BP) |
| Synergy | Antagonism | Welch Two Sample t-test | 27.983 | p < .001 | 226.056 | Co-functional (EcoCyc/GO-BP) |
| Additivity | Antagonism | Welch Two Sample t-test | 5.113 | p < .001 | 188.331 | Co-functional (EcoCyc/GO-BP) |
| Synergy | Additivity | Welch Two Sample t-test | 53.463 | p < .001 | 1233.005 | Small/medium-scale PPI |
| Synergy | Antagonism | Welch Two Sample t-test | 25.244 | p < .001 | 224.621 | Small/medium-scale PPI |
| Additivity | Antagonism | Welch Two Sample t-test | 1.45 | 0.148 | 205.288 | Small/medium-scale PPI |

| Statistical summary for Fig. 4D | | | | | | |
|---|---|---|---|---|---|---|
| **Model** | **Term** | **Estimate** | **std.error** | **statistic** | **p-value** | **Network** |
| OLS(Mean.sigma ~ path.length.corr) | Intercept | 0.00207 | 0.000247 | 8.37 | 0.00111 | Co-functional (EcoCyc/GO-BP) |
| OLS(Mean.sigma ~ path.length.corr) | Slope | -0.00024 | 0.000063 | -3.75 | 0.01992 | Co-functional (EcoCyc/GO-BP) |
| OLS(Mean.sigma ~ path.length.corr) | Intercept | 0.00278 | 0.000212 | 13.15 | 0.00019 | Small/medium-scale PPI |
| OLS(Mean.sigma ~ path.length.corr) | Slope | -0.00027 | 0.000054 | -4.96 | 0.00769 | Small/medium-scale PPI |

**Sup. Table S3. Details of some of the DDIs analyzed.**

| **Drug pair** | **A22_Novobiocin** | **A22_Novobiocin** | **Erythromycin_Trimethoprim** | **Sulfamonomethoxine_Trimethoprim** |
|---|---|---|---|---|
| **Network** | Co-functional EcoCyc/GO-BP | Small/medium scale PPI | NA | Co-functional EcoCyc/GO-BP |
| **Sigma rate** ($bliss^2$/MYA) | 0.0035 | 0.0035 | 0.0003 | 0.0035 |
| **Path length** | 2 | 3.5 | NA | 1 |
| **K-edge** | 12 | 3 | NA | 21 |
| **Mean node degree** | 16 | 5.75 | NA | 26.5 |
| **DDI in ebw (D1-D2)** | Additive (S/S) | | Additive (S/S) | Synergy **(R/S)** |
| **DDI in ecr (D1-D2)** | Synergy (S/S) | | Additive (S/S) | Synergy **(R/S)** |
| **DDI in seo (D1-D2)** | Synergy **(S/R)** | | Additive (S/**R**) | Synergy **(R/S)** |
| **DDI in stm (D1-D2)** | Additive (S/S) | | Additive (S/**R**) | Synergy **(R/S)** |
| **DDI in pae (D1-D2)** | Additive **(R/R)** | | Additive (S/S) | Additive **(R/R)** |
| **DDI in pau (D1-D2)** | Additive **(S/R)** | | Additive (S/S) | Additive **(R/R)** |

**Sup. Table S4. Network measurements of DDIs**
See separate CSV File.

**Sup. Document S1. t-SNE results of all DDIs for perplexity range between 5 and 2100.**
See separate PDF File

**Sup. Fig. S1: DDI-based Euclidean distances between strains scale as a function of phylogenetic distance plotted on a log scale.**

Intraspecific comparisons (in blue) have the lowest amount of sequence and chemical divergence, while comparisons with Pseudomonas are the most distant in terms of sequence and chemical divergence. All drug-drug interaction data is from Brochado et al. (2018).

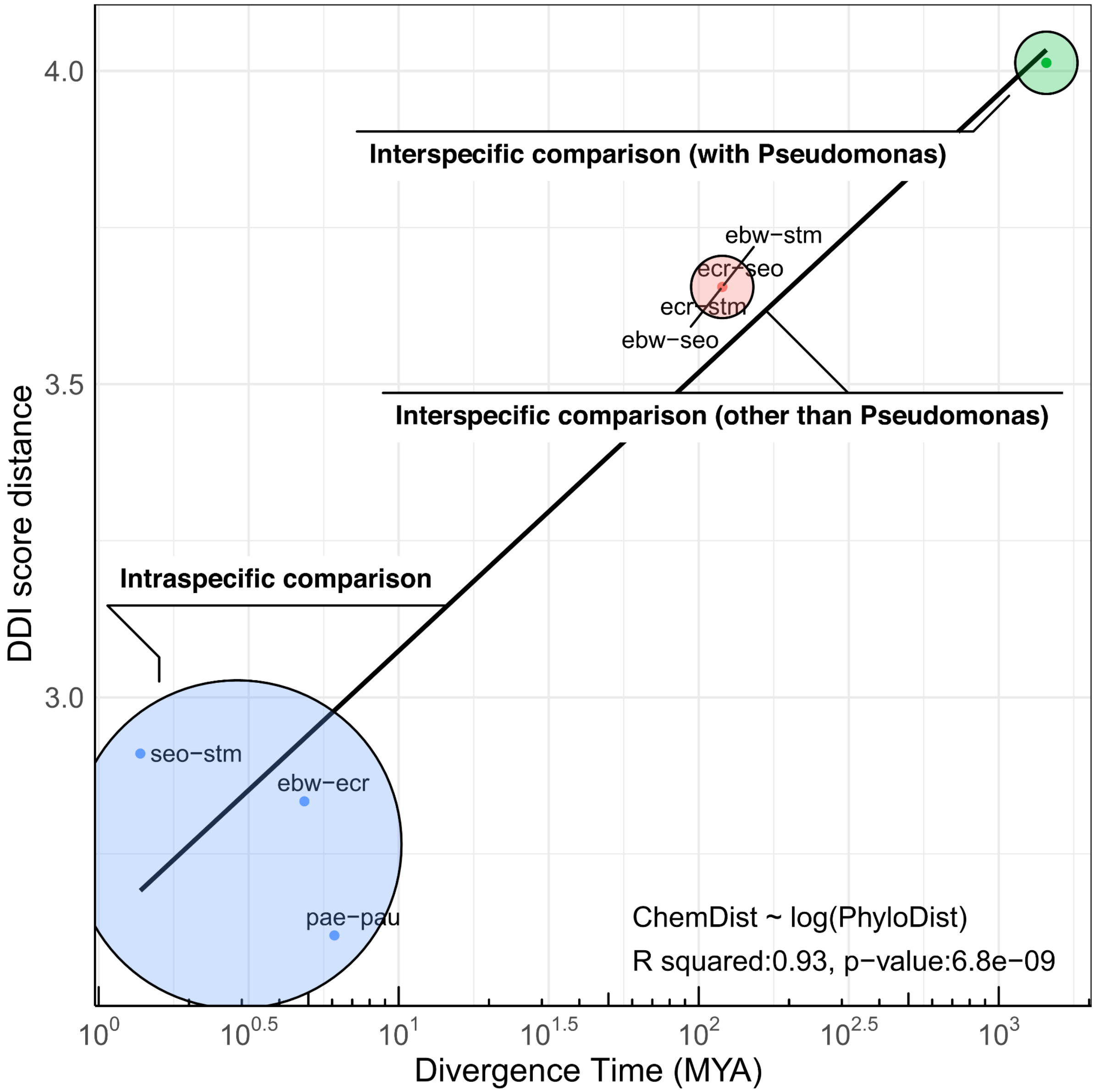

**Sup. Fig. S2: Evolutionary rates of DDIs vs network metrics of DDI targets in two *E coli* networks.**

**A.** Comparison of evolutionary rates between adjacent and non-adjacent targets in the co-functional (EcoCyc) network.
**B.** Comparison of evolutionary rates between adjacent and non-adjacent targets in the PPI network.
**C.** Comparison of evolutionary rates between connected and not-connected targets in the co-functional (EcoCyc) network.
**D.** Comparison of the minimum distance between targets for different combinations of DDI across strains for co-functional (EcoCyc/GO-BP) and PPI networks.
**E.** Comparison of k edge connectivity between targets for different combinations of DDI across strains for (EcoCyc/GO-BP) and PPI networks.
**F.** Comparison of node degree between targets for different combinations of DDI across strains for (EcoCyc/GO-BP) and PPI networks.
**G** and **H.** Comparison of average eigenvector centrality between targets for different combinations of DDI across strains for (EcoCyc/GO-BP) and PPI networks.
**I. and J.** Distribution of connected and disconnected target connections for different types of DDI in the (EcoCyc/GO-BP) and PPI networks.
**K.** Comparison of evolutionary rates for different K-edge connectivity levels in the co-functional (EcoCyc) network.
**L.** Comparison of evolutionary rates for different node degree levels in the co-functional (EcoCyc) network.
**M.** Comparison of evolutionary rates for different betweenness centrality levels in the co-functional (EcoCyc) network.
**N.** Comparison of evolutionary rates for different eigenvector centrality levels in the co-functional (EcoCyc) network.

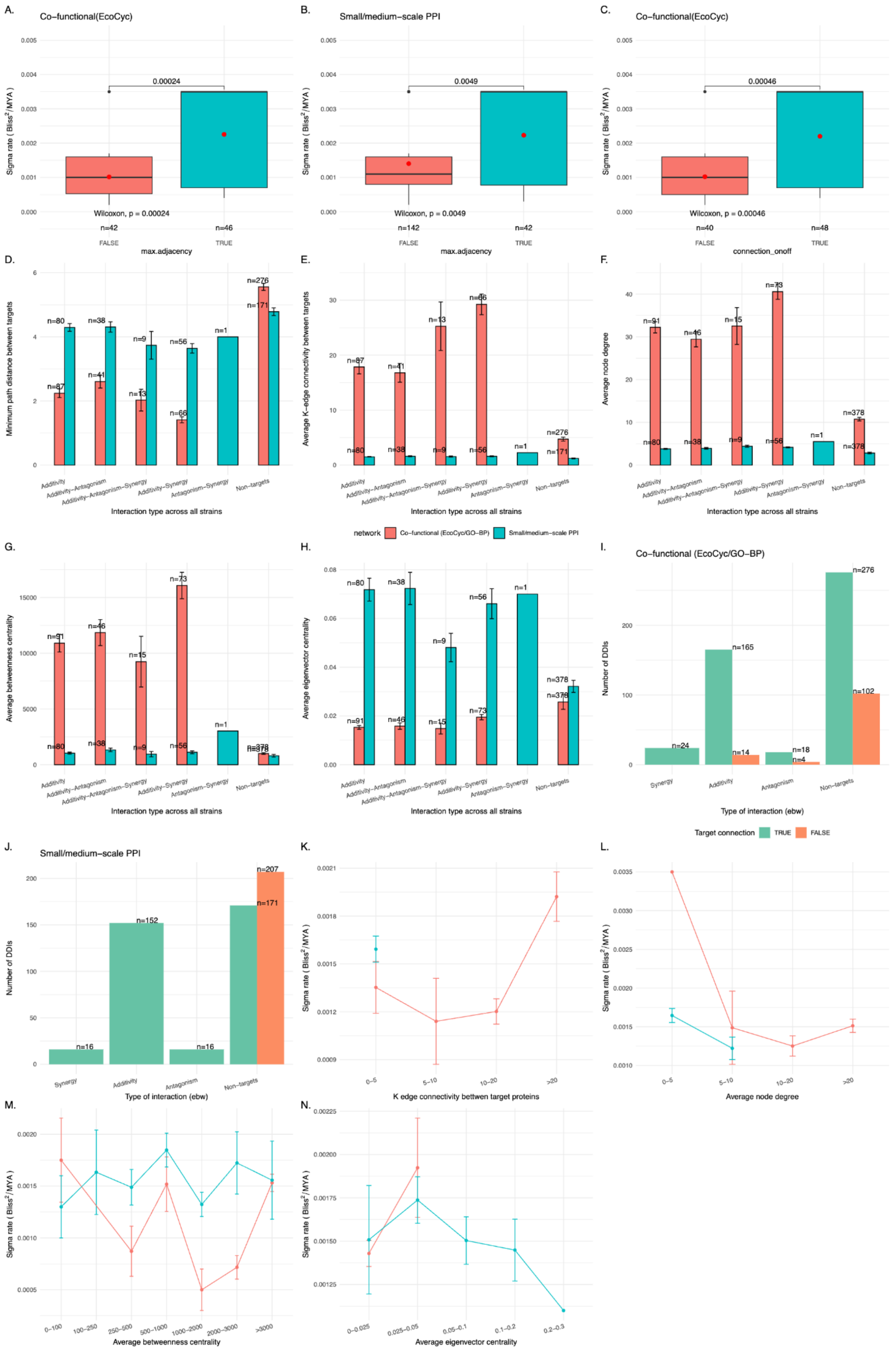

A.
Co−functional(EcoCyc)
Sigma rate ( Bliss$^2$ / MYA )
0.00024
Wilcoxon, p = 0.00024
n=42
n=46
FALSE
TRUE
max.adjacency
B.
Small/medium−scale PPI
0.0049
Wilcoxon, p = 0.0049
n=142
n=42
max.adjacency
C.
Co−functional(EcoCyc)
0.00046
Wilcoxon, p = 0.00046
n=40
n=48
connection_onoff
D.
Minimum path distance between targets
E.
Average K−edge connectivity between targets
F.
Average node degree
Additivity
Additivity−Antagonism
Additivity−Antagonism−Synergy
Additivity−Synergy
Antagonism−Synergy
Non−targets
Interaction type across all strains
network
Co−functional (EcoCyc/GO−BP)
Small/medium−scale PPI
G.
Average betweenness centrality
H.
Average eigenvector centrality
I.
Co−functional (EcoCyc/GO−BP)
Number of DDIs
Synergy
Additivity
Antagonism
Non−targets
Type of interaction (ebw)
Target connection
TRUE
FALSE
J.
Small/medium−scale PPI
Number of DDIs
Type of interaction (ebw)
K.
K edge connectivity bettwen target proteins
L.
Average node degree
M.
Average betweenness centrality
N.
Average eigenvector centrality

**Sup. Fig. S3. Heatmap of DDI scores across strains for the subset of DDI that are susceptible in at least one of the species for both drugs.**

Hierarchical clustering based on Euclidean distances across columns (strains: *E. coli* K-12 BW2952 (ebw), *Escherichia coli* O8 IAI1 (ecr), *S. enterica subsp. enterica* serovar Typhimurium 14028S (seo), *Salmonella enterica* subsp. enterica serovar Typhimurium LT2 (stm), *Pseudomonas aeruginosa* PAO1 (pae), *P. aeruginosa* UCBPP-PA14 (pau)), and rows (DDIs); the latter was constrained by tSNE clusters. Synergistic interactions are shown in blue and antagonistic interactions are in red, all DDIs are measured in bliss score units. The annotations on the right indicate whether the two drugs involved in the interaction belong to the same drug category, target the same cellular process, or have the same use (in green) or not (black). The bottom row indicates the evolutionary rate calculated in each cluster, measured in bliss score units$^2$ per million years.

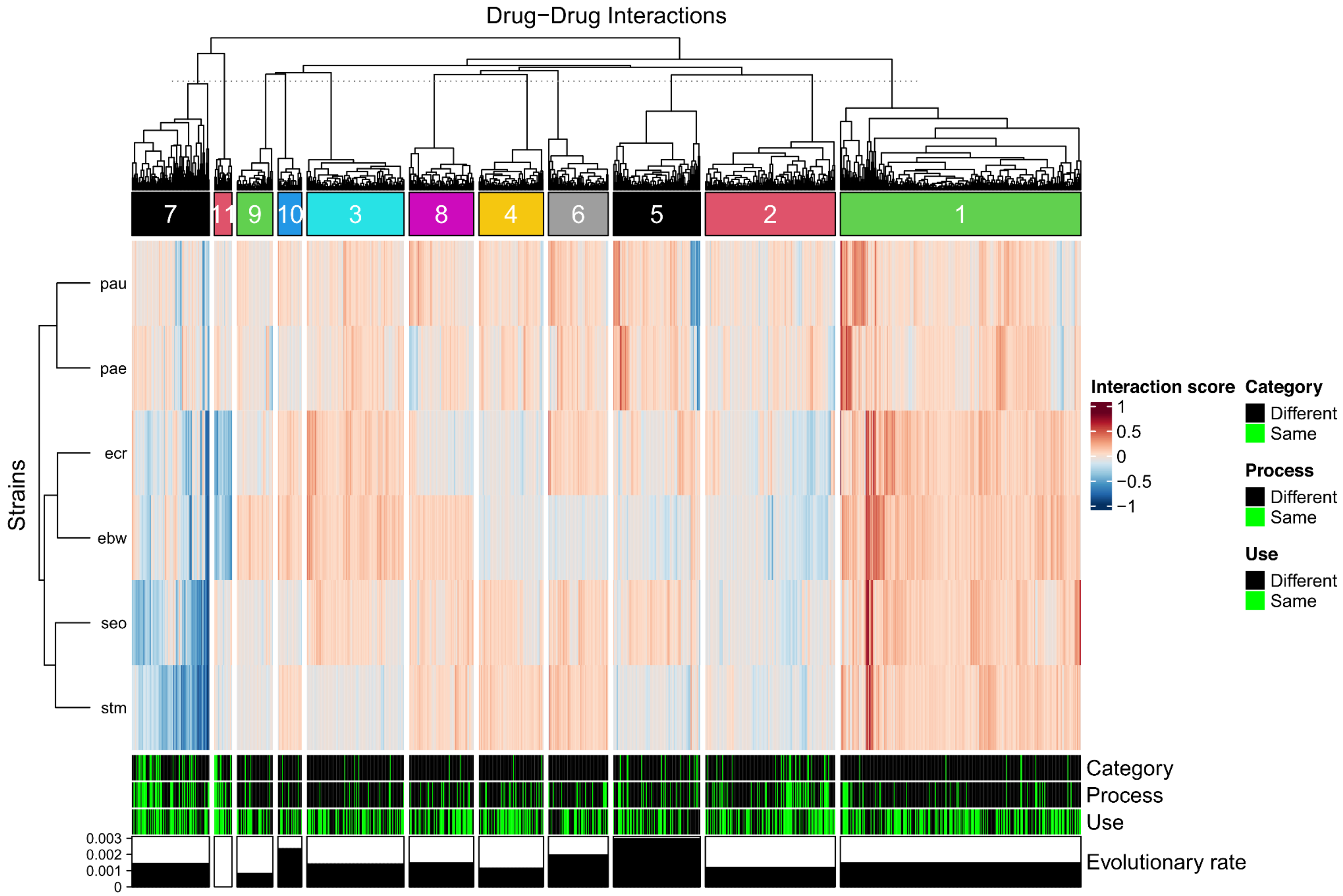

**Sup. Fig. S4. Differences in minimum path length, k edge connectivity, mean node degree, mean betweenness centrality, and mean eigenvector centrality between targets for different types of drug interactions in *E. coli* for co-functional and PPI networks for the subset of DDI that are susceptible in at least one of the species for both drugs.**

**A-B**. Synergistic interactions have a lower average path length than additive interactions and antagonistic interactions. **C-D.** K-edge connectivity is higher in synergies than in additive and antagonistic interactions in the co-functional network, but not in the PPI network. **E-F.** The mean node degree is higher in synergies than in additive or antagonistic interactions in the co-functional network, but not in the PPI network. **G-H**. Mean betweenness centrality is higher in synergies than in additive or antagonistic interactions in the co-functional network, but not in the PPI network. **I-J**. Mean eigenvector centrality is higher in synergies than in additive or antagonistic interactions in the co-functional network, but not in the PPI network.

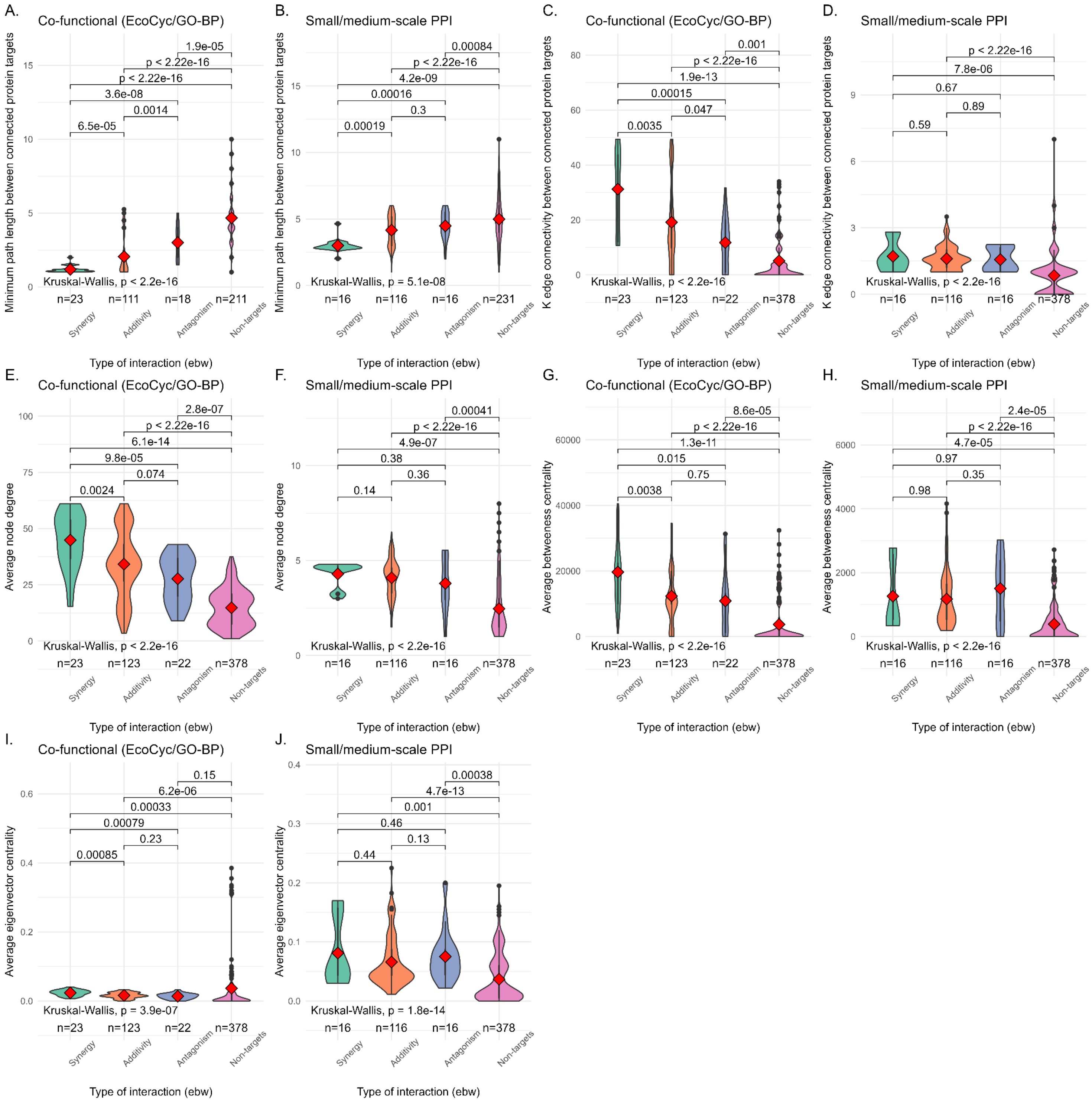

**Sup. Fig. S5. Relationships among evolutionary rate, interaction type, and path length for the subset of DDI that are susceptible in at least one of the species for both drugs.**

**A.** DDIs containing only synergies or synergy-antagonism combinations across all strains have faster evolutionary rates. Error bars represent the standard error of the mean. **B.** DDIs that are more synergistic across all strains have higher evolutionary rates. The score is the aggregated sum across strains, where DDIs were scored as +1 for synergies, 0 for additive, and -1 for antagonistic interactions. Error bars represent the standard error of the mean. **C.** Evolutionary rate as a function of interaction type for the co-functional (pink) and small/medium scale PPI network (blue). Synergistic interactions have higher evolutionary rates than additive and antagonistic interactions in *E. coli*. **D.** Evolutionary rate as a function of the minimum distance between targets. Wiring between close nodes in biological networks evolves more quickly than between distant nodes.

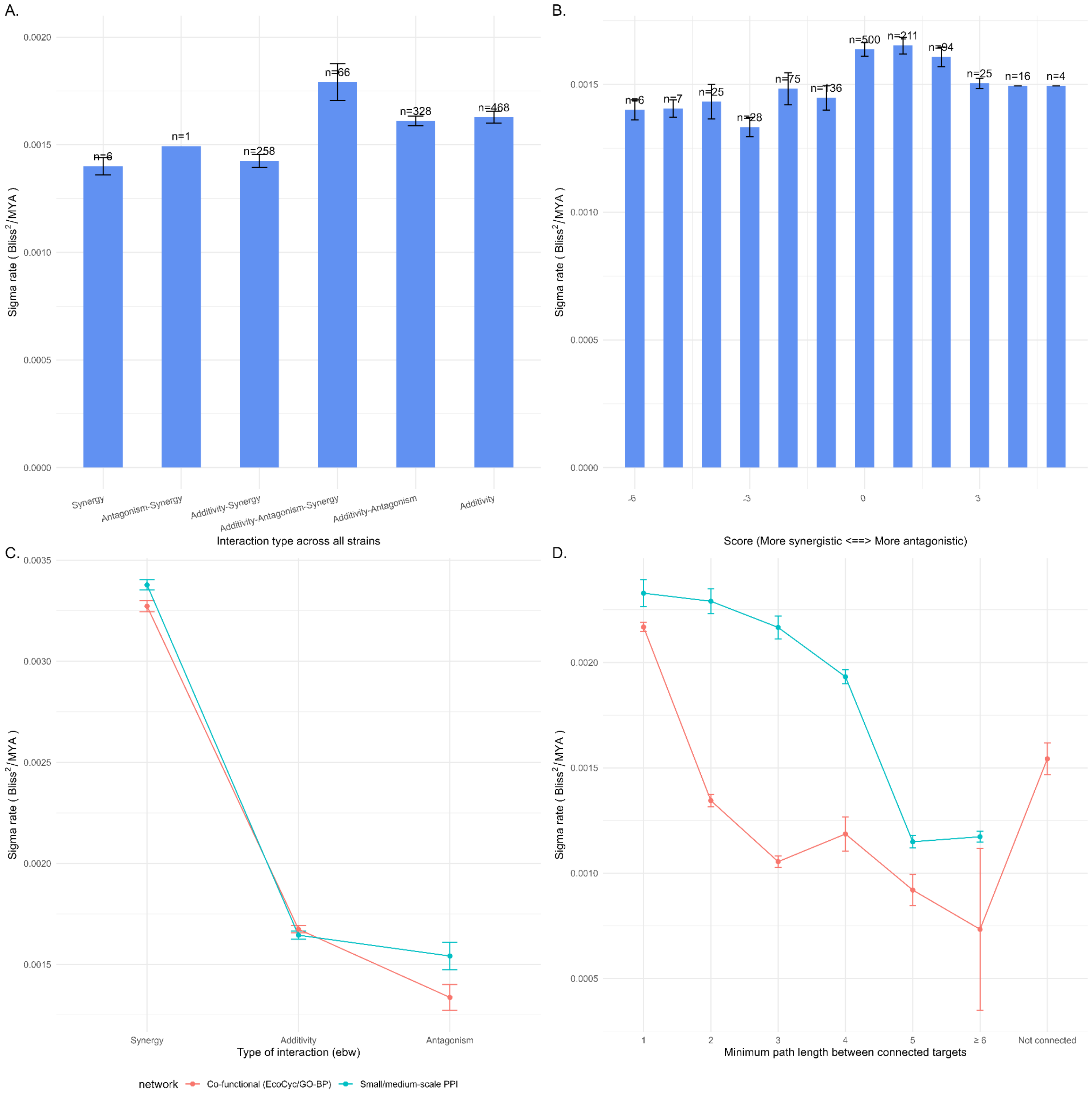

**Sup. Fig. S6. Dotplot showing significant GO categories of drug targets across clusters.** Clusters 3, 6, and 16 have significant enrichment for ion binding. Clusters 2, 3, 4, 6, and 16 are significantly enriched for anion binding and small molecule binding.

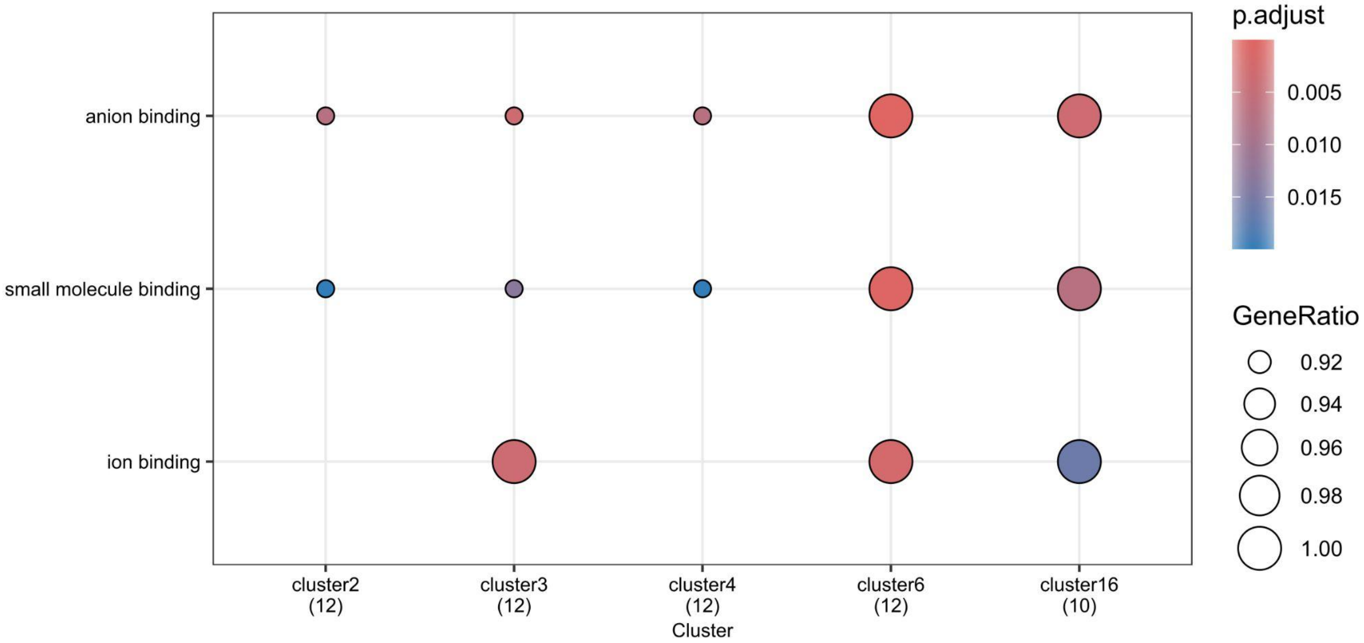
