## Supplementary_Document_S1 for "Wiring between close nodes in biological networks evolves more quickly than between distant nodes"

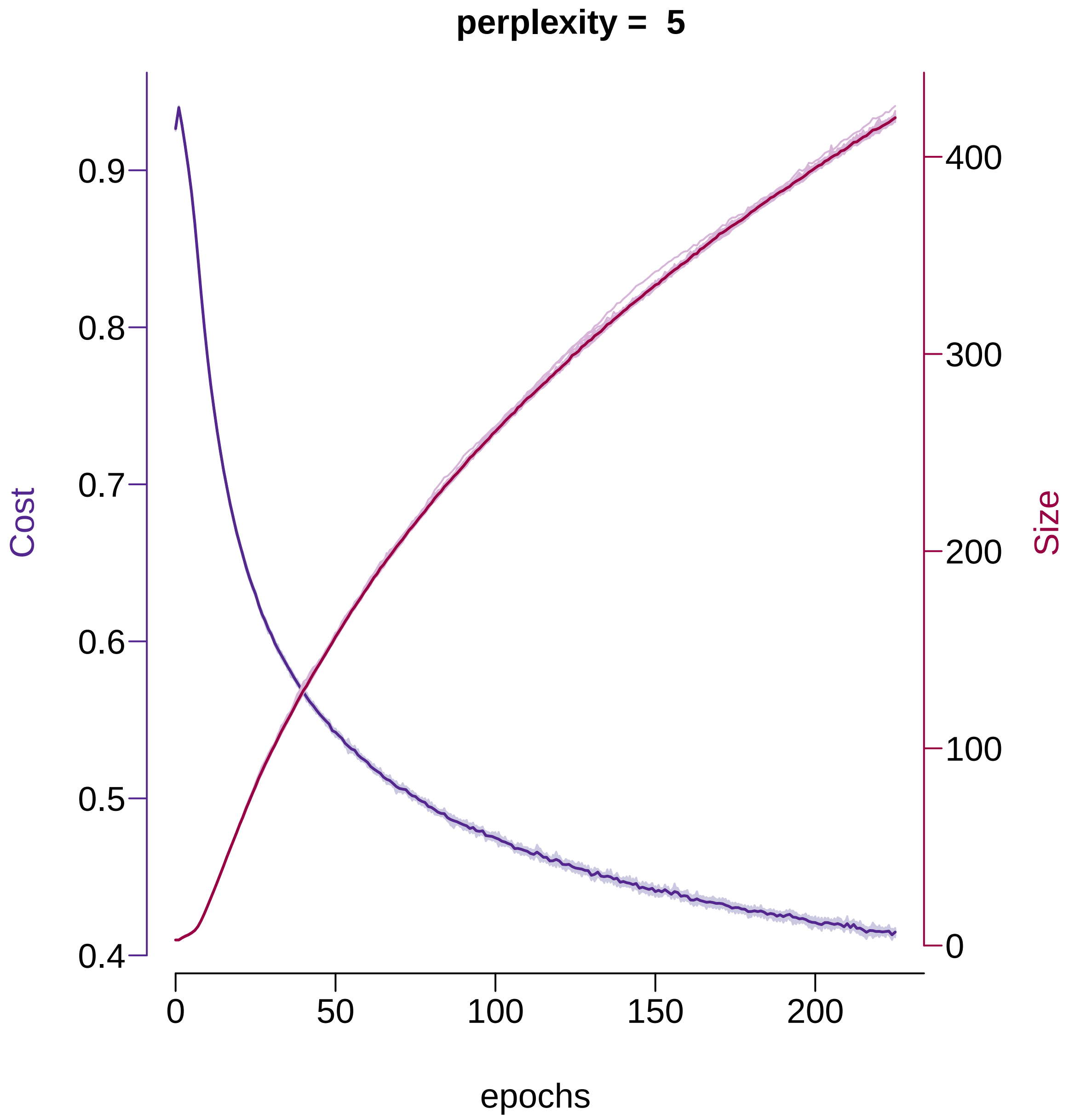
perplexity = 5
Cost
0.4
0.5
0.6
0.7
0.8
0.9
Size
0
100
200
300
400
epochs
0
50
100
150
200

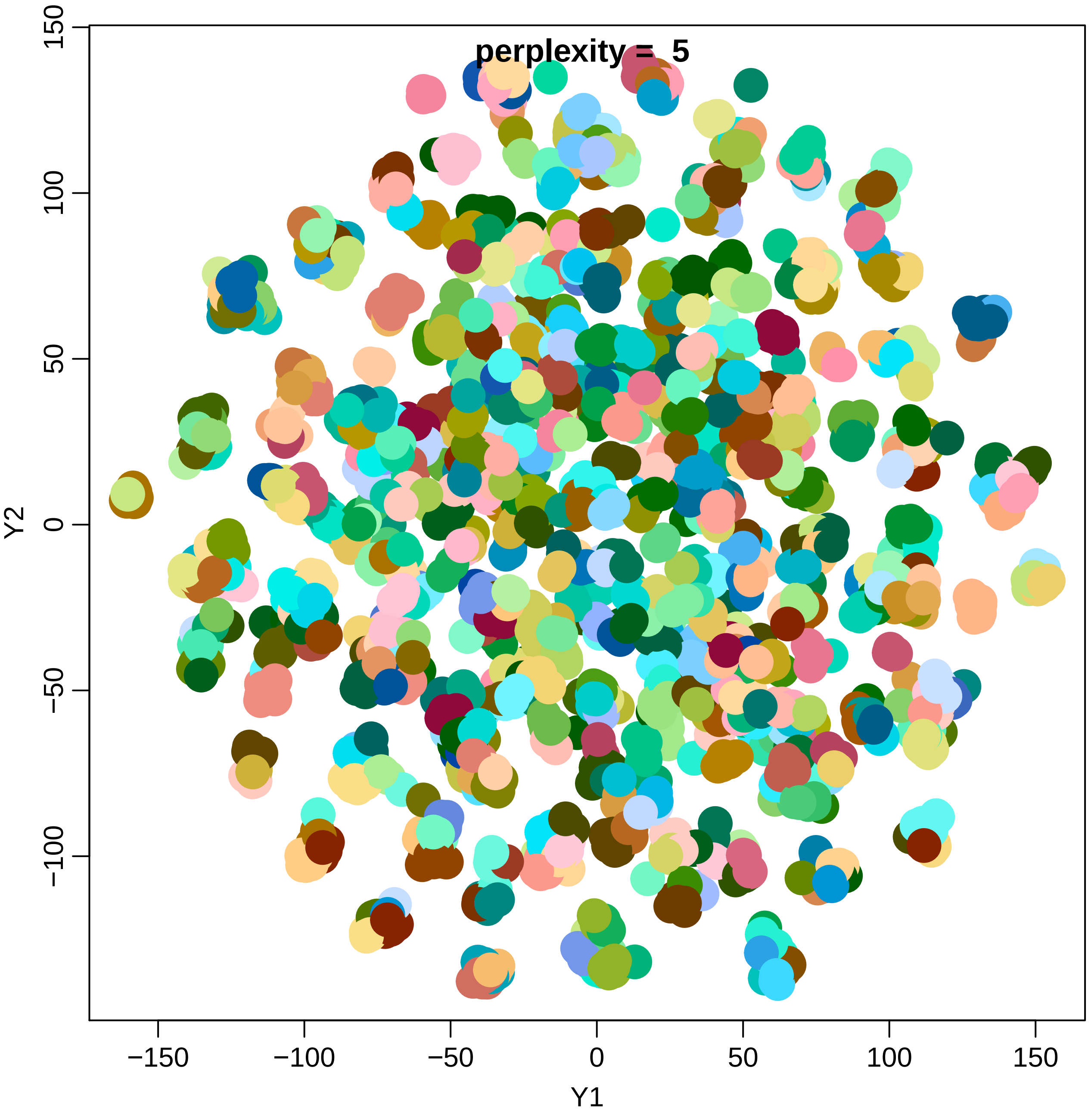

perplexity = 5
Y2
Y1
150
100
50
0
−50
−100
−150
−100
−50
0
50
100
150

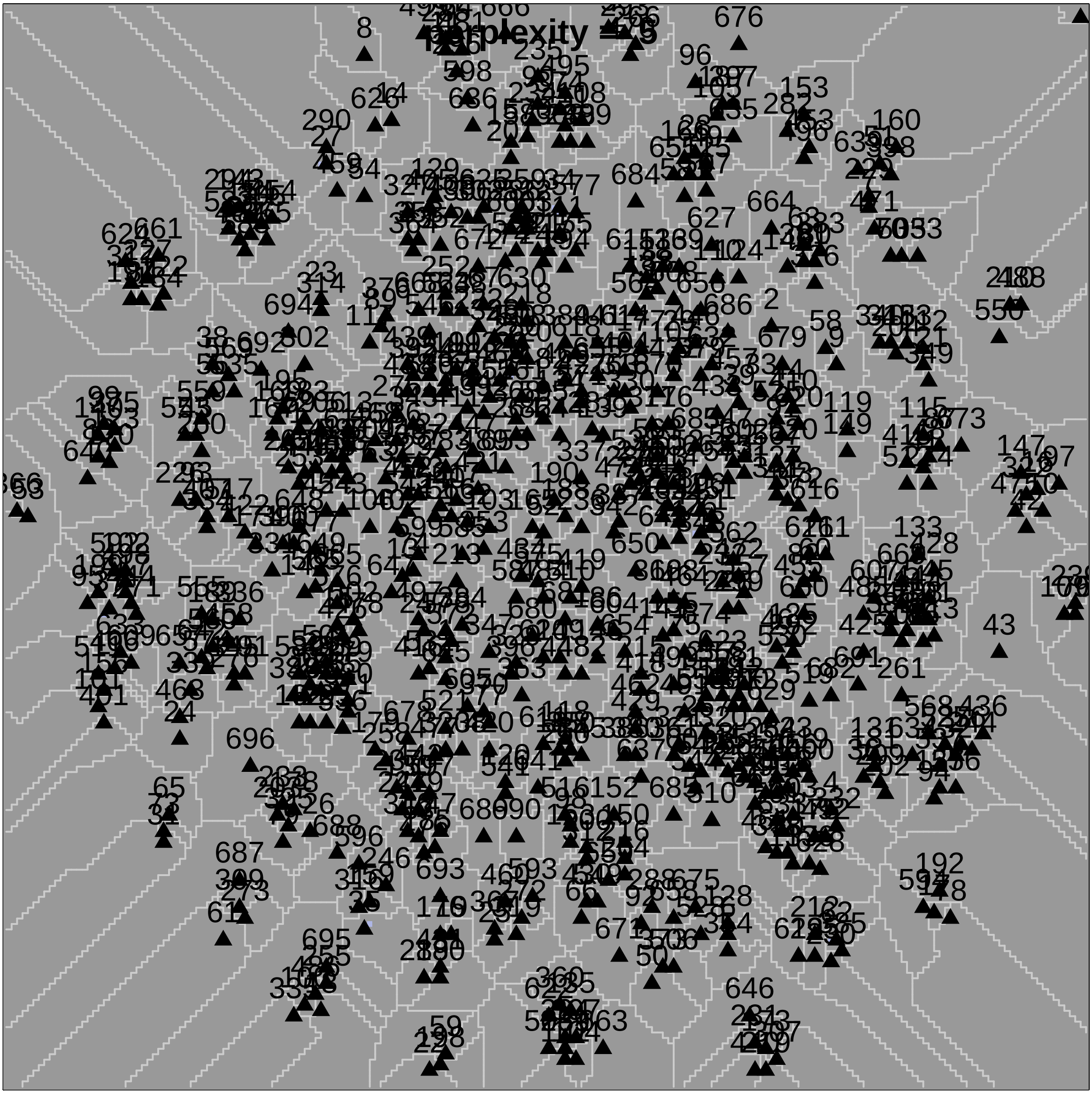

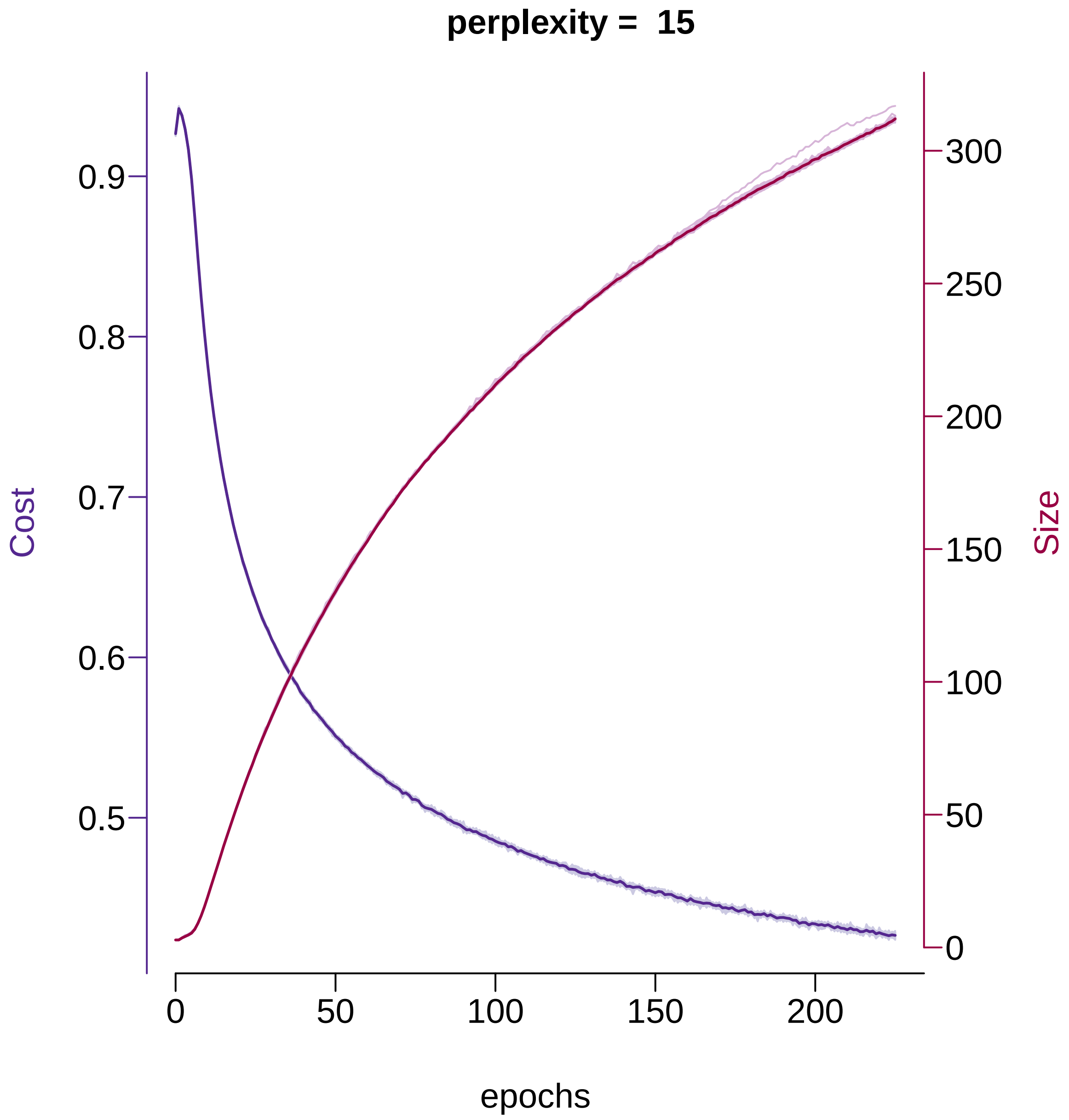

perplexity = 15
Cost
Size
epochs
0.9
0.8
0.7
0.6
0.5
300
250
200
150
100
50
0
0
50
100
150
200

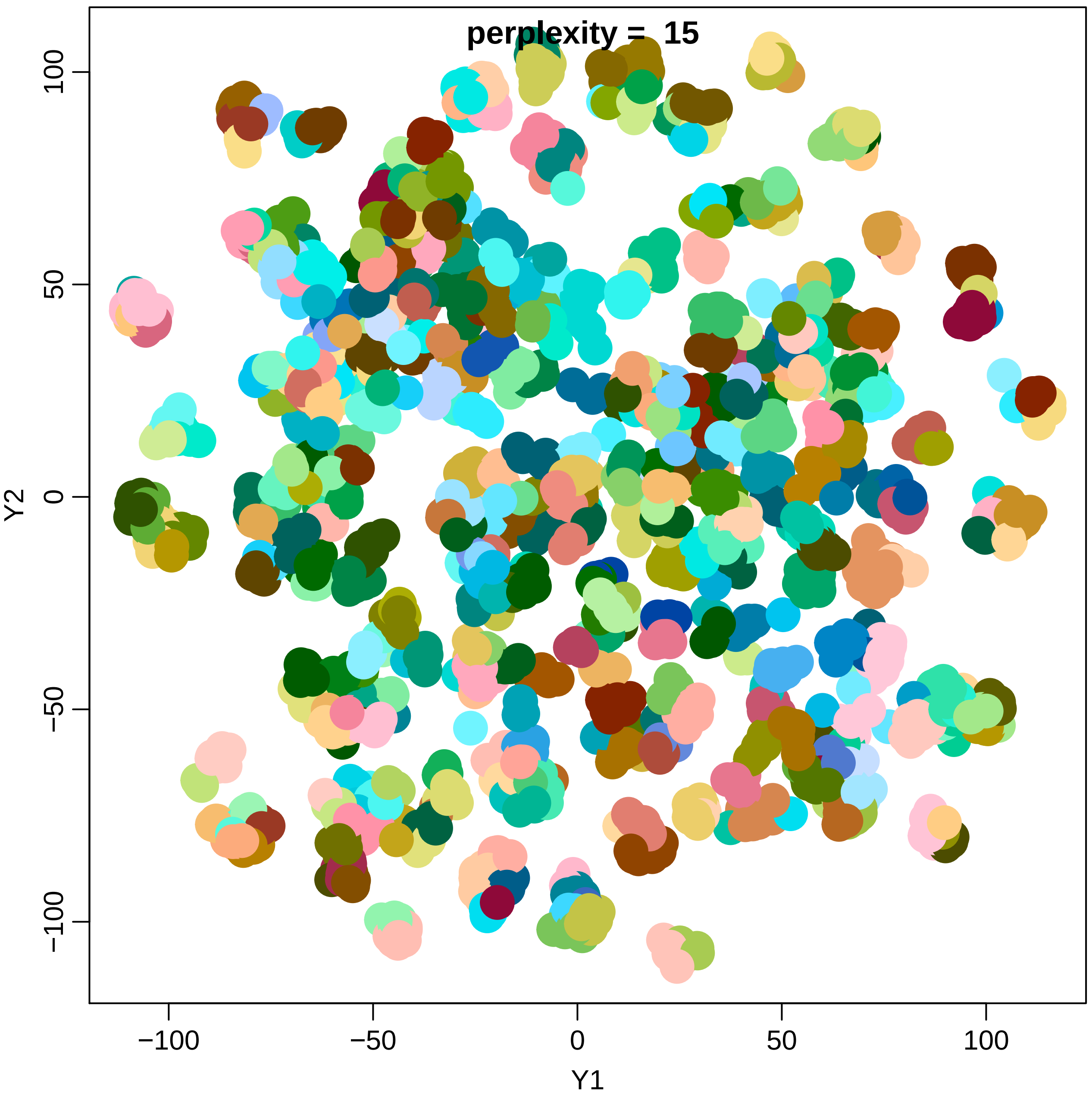
perplexity = 15
Y2
Y1
100
50
0
−50
−100
−100
−50
0
50
100

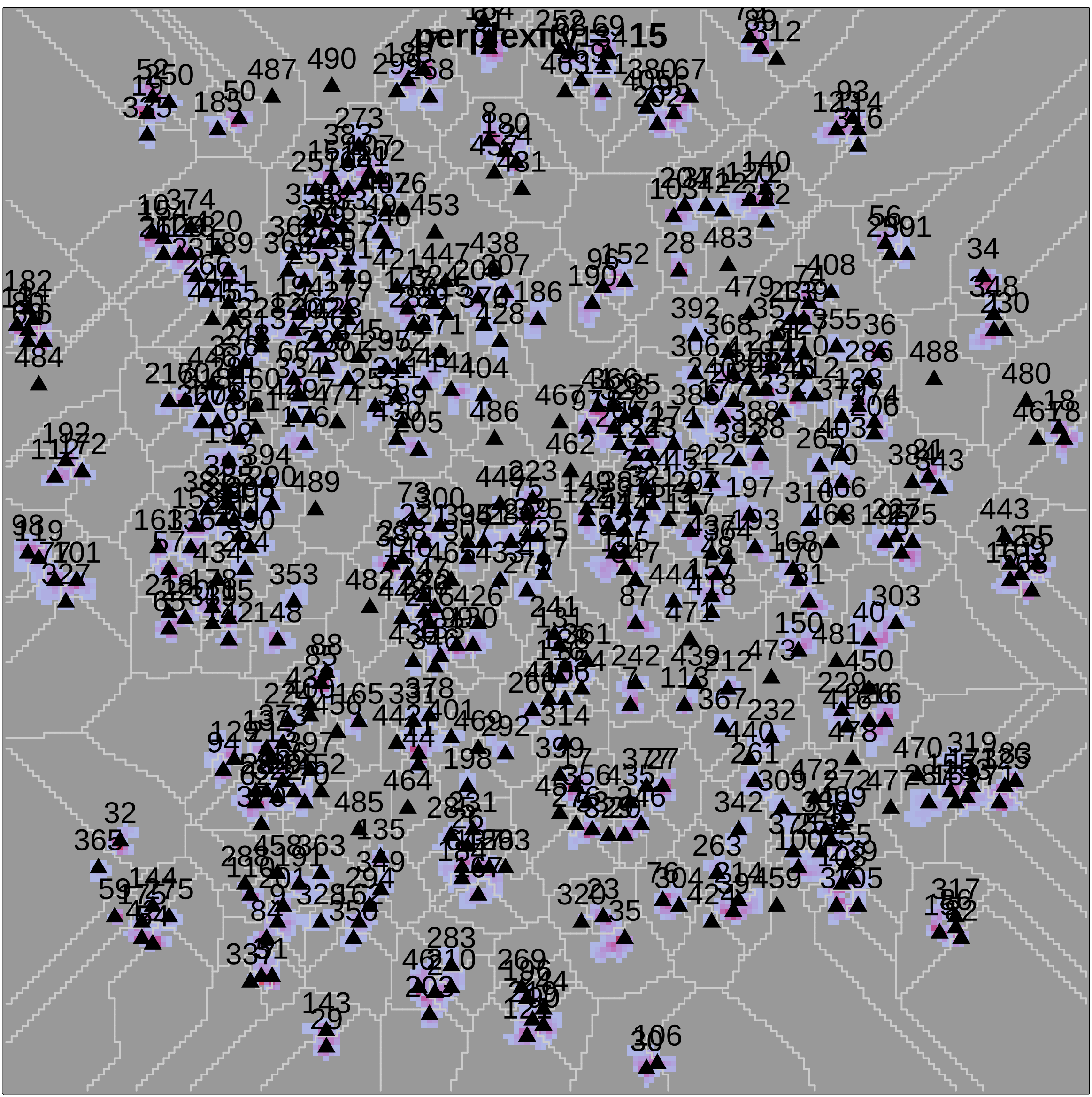

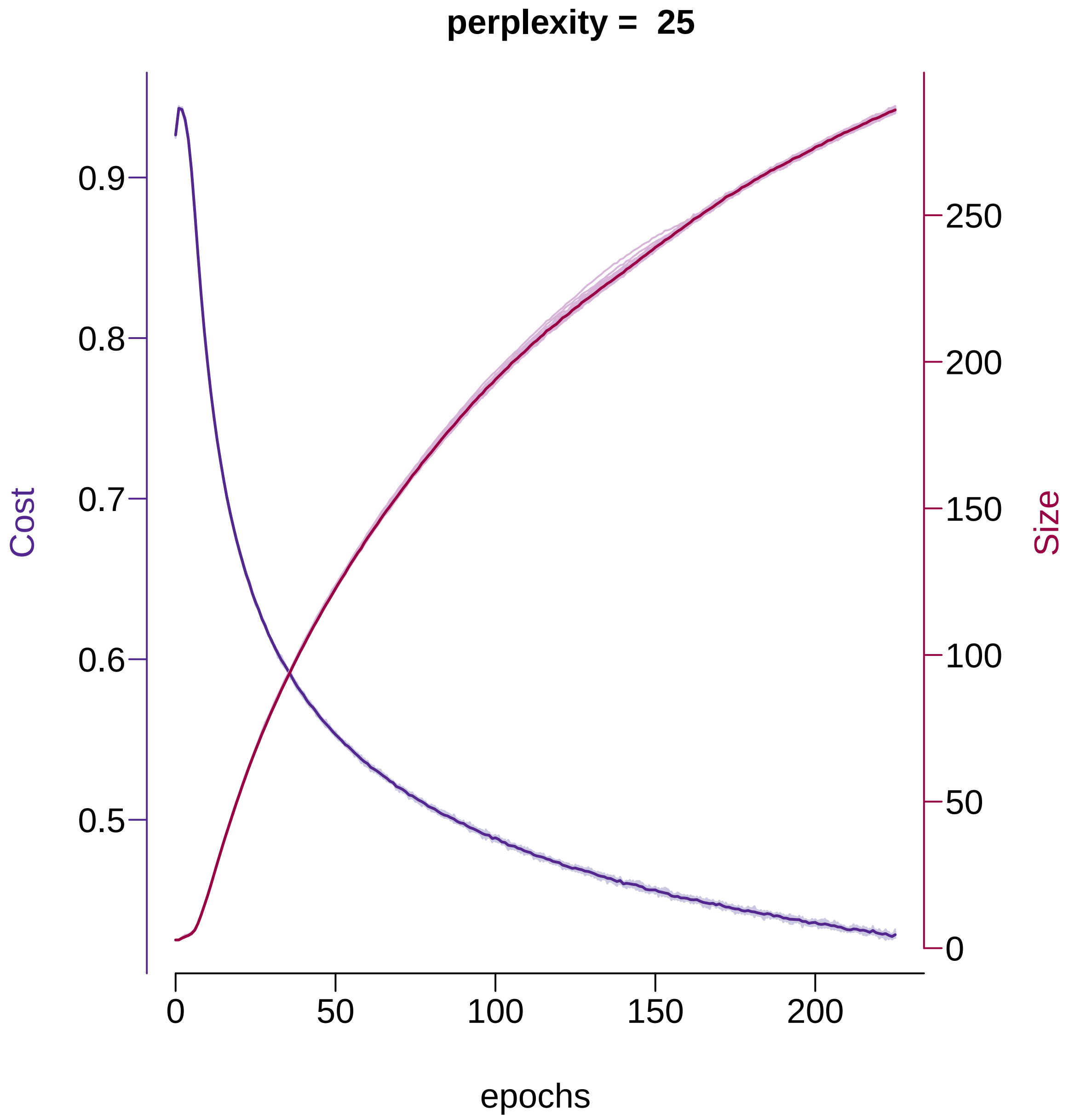

perplexity = 25
Cost
0.9
0.8
0.7
0.6
0.5
Size
250
200
150
100
50
0
0
50
100
150
200
epochs

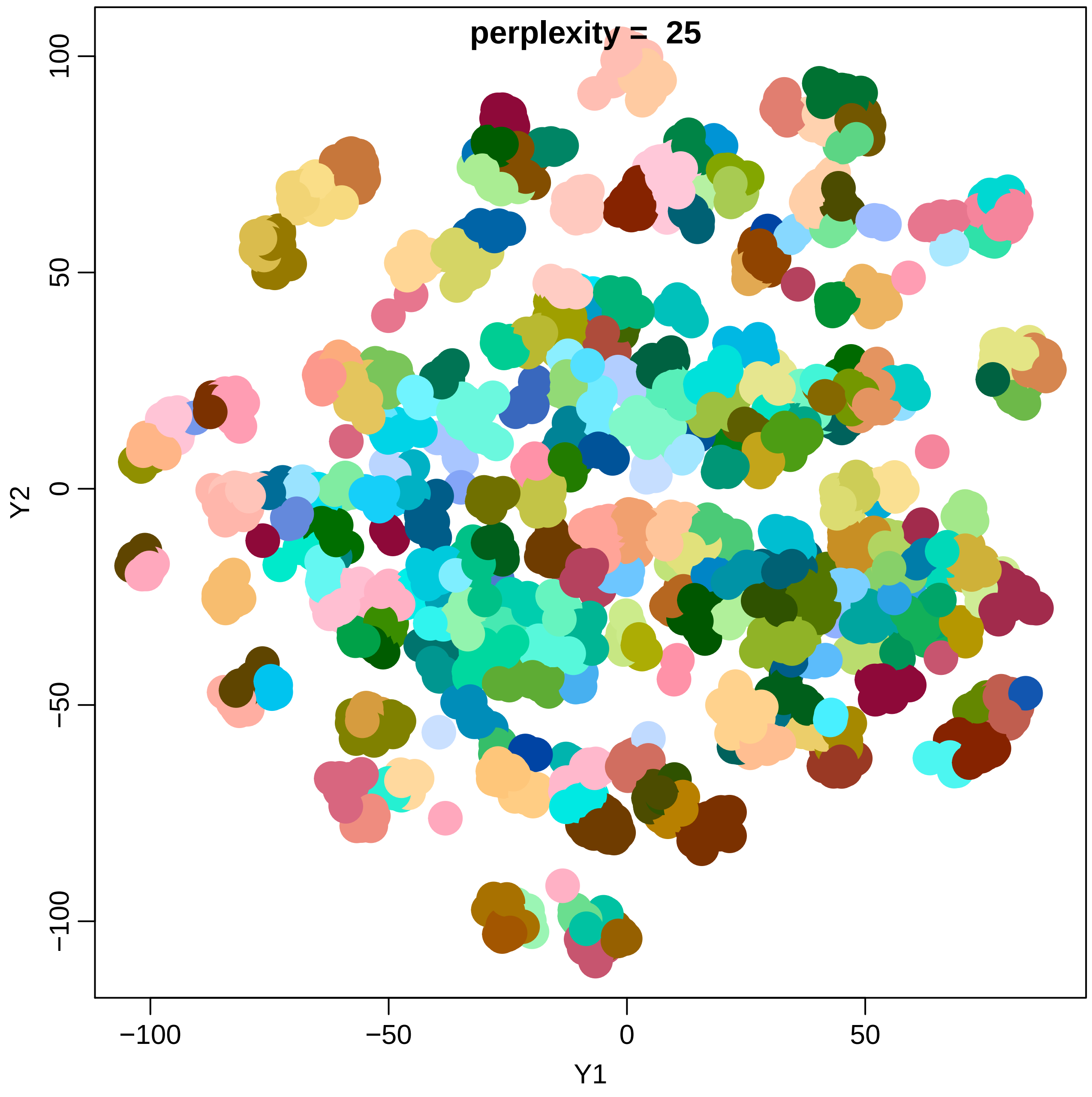

perplexity = 25
Y2
Y1
100
50
0
−50
−100
−100
−50
0
50

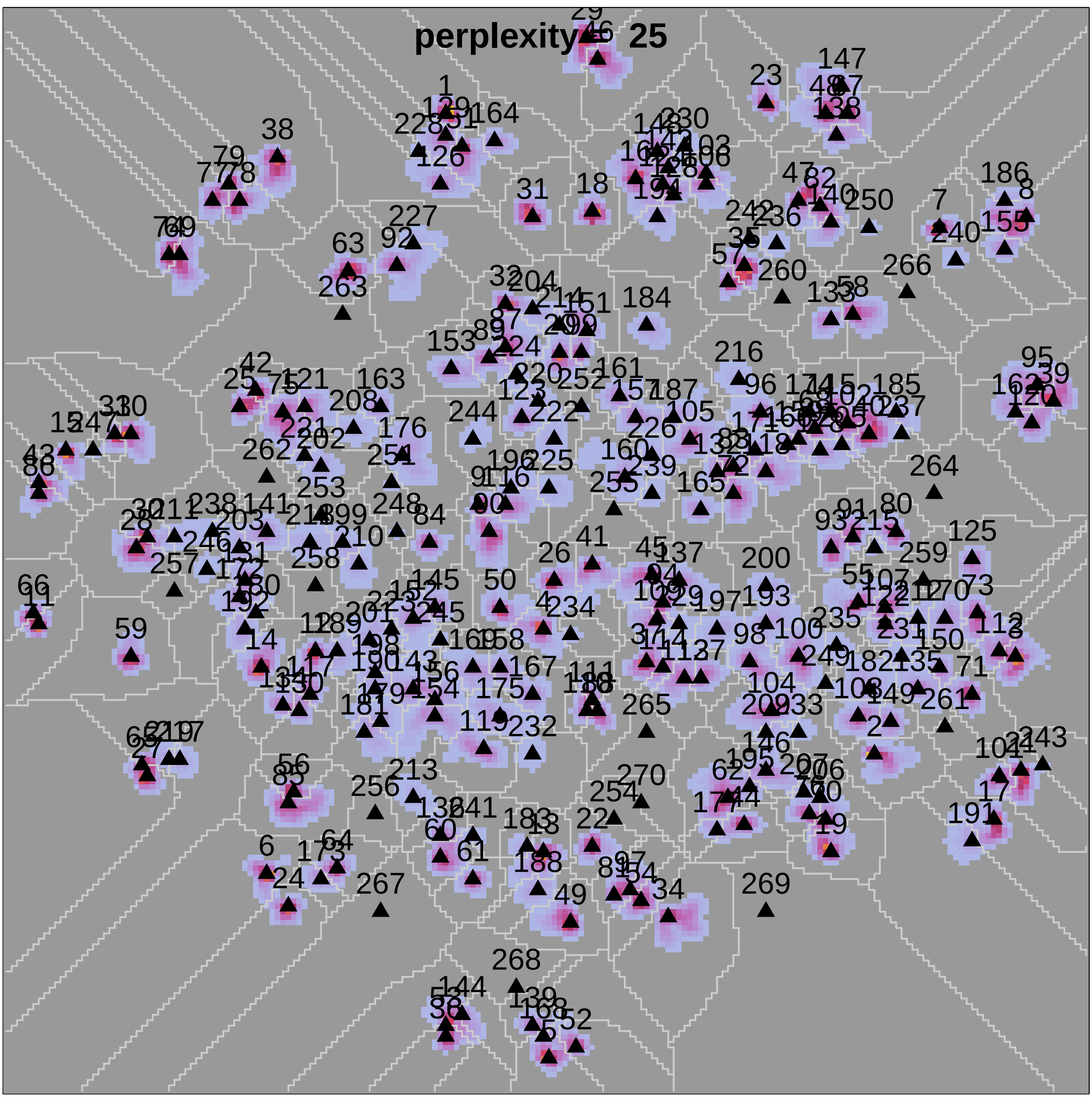
perplexity = 25

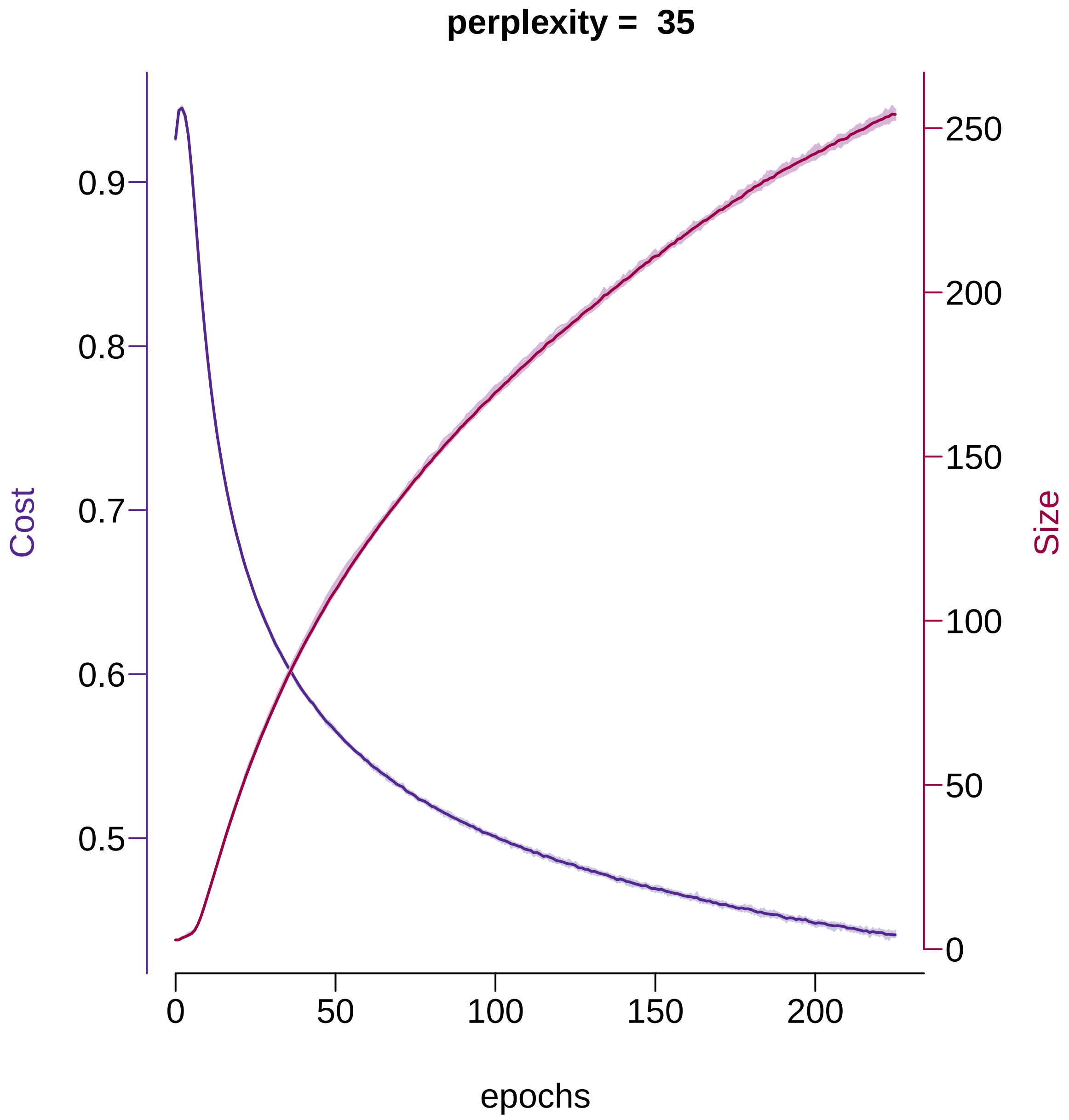

perplexity = 35
Cost
0.9
0.8
0.7
0.6
0.5
Size
250
200
150
100
50
0
0
50
100
150
200
epochs

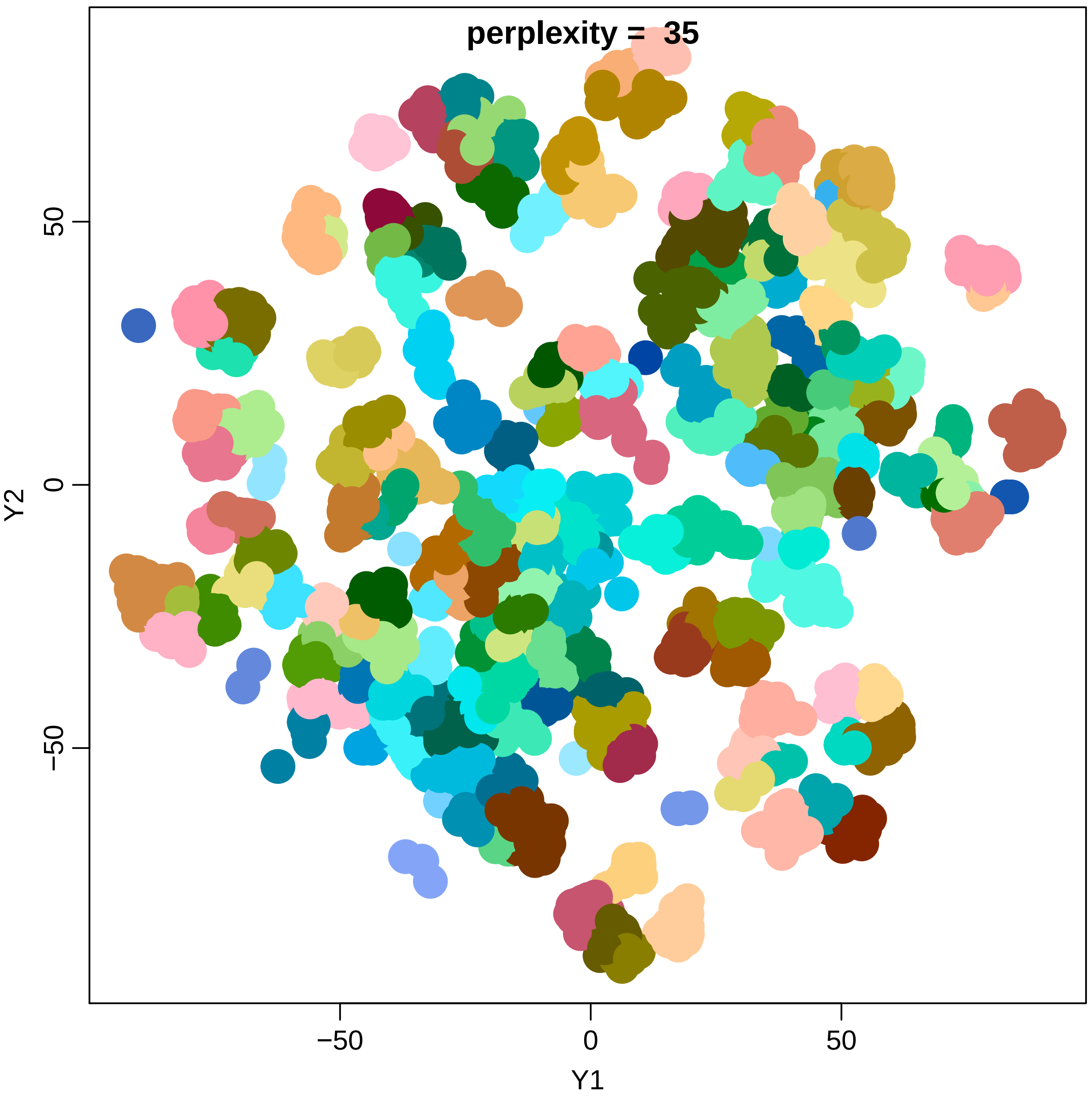

perplexity = 35
Y2
Y1
50
0
−50
−50
0
50

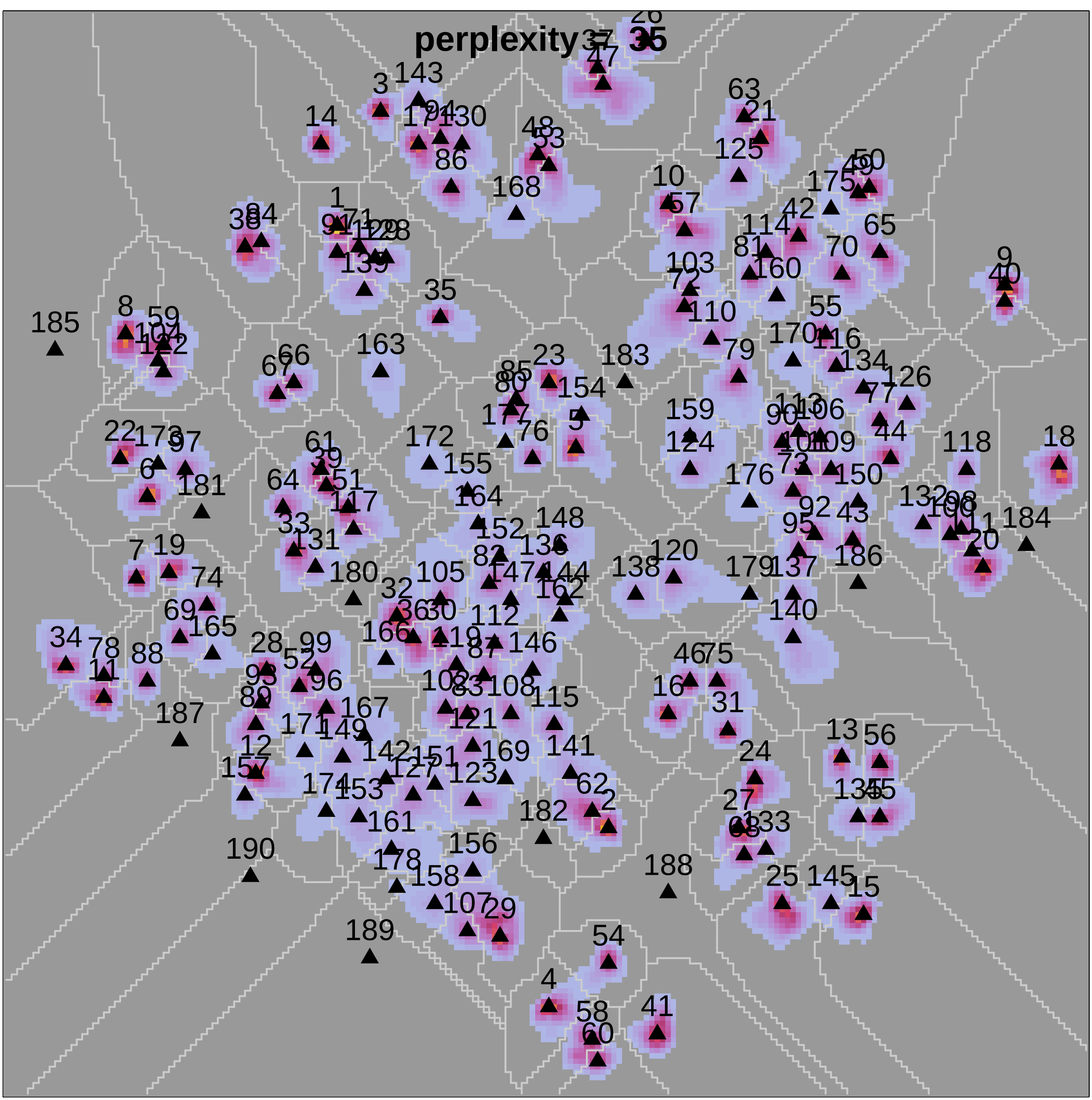
perplexity = 35

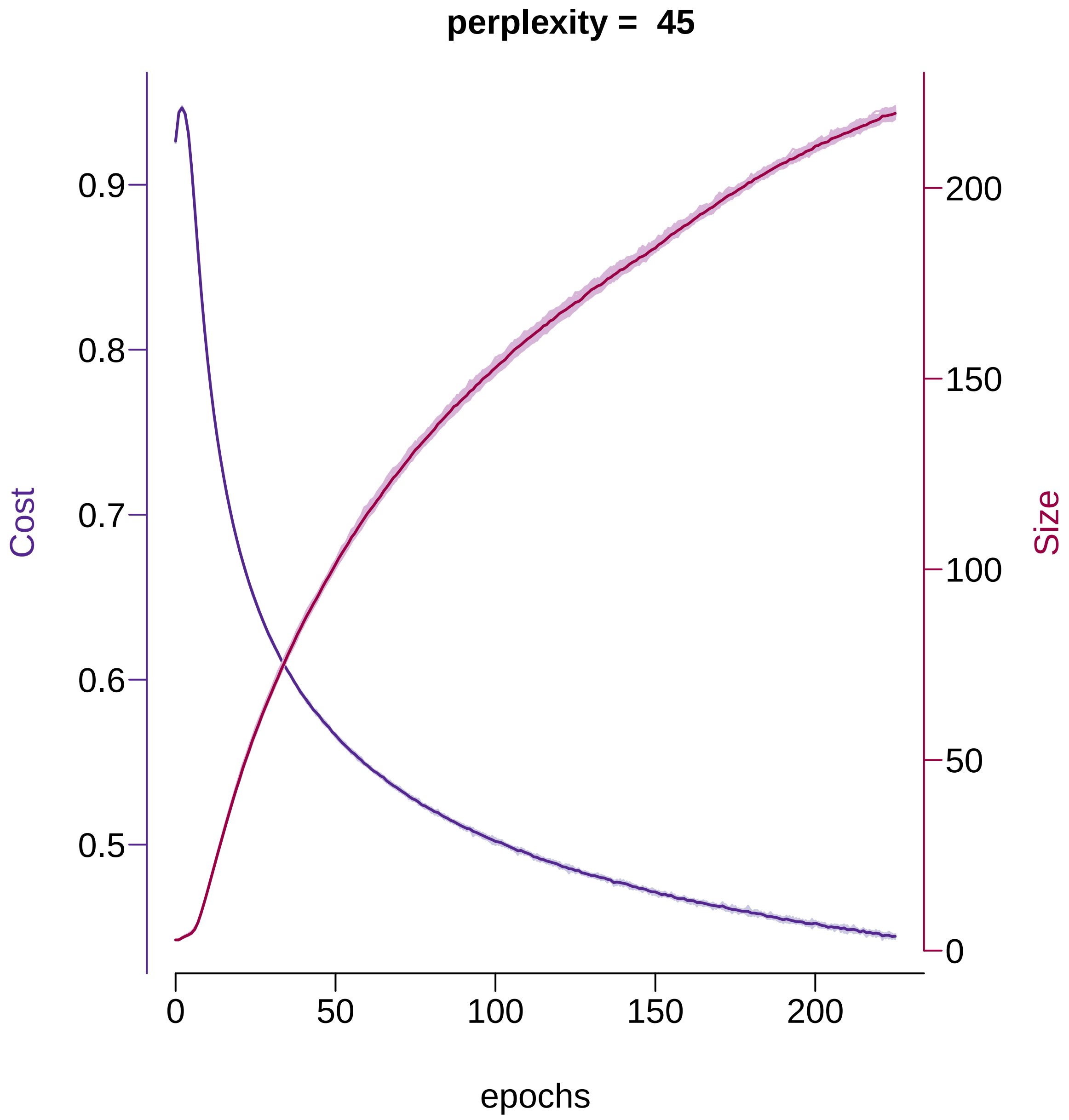

perplexity = 45
Cost
Size
0.9
0.8
0.7
0.6
0.5
200
150
100
50
0
0
50
100
150
200
epochs

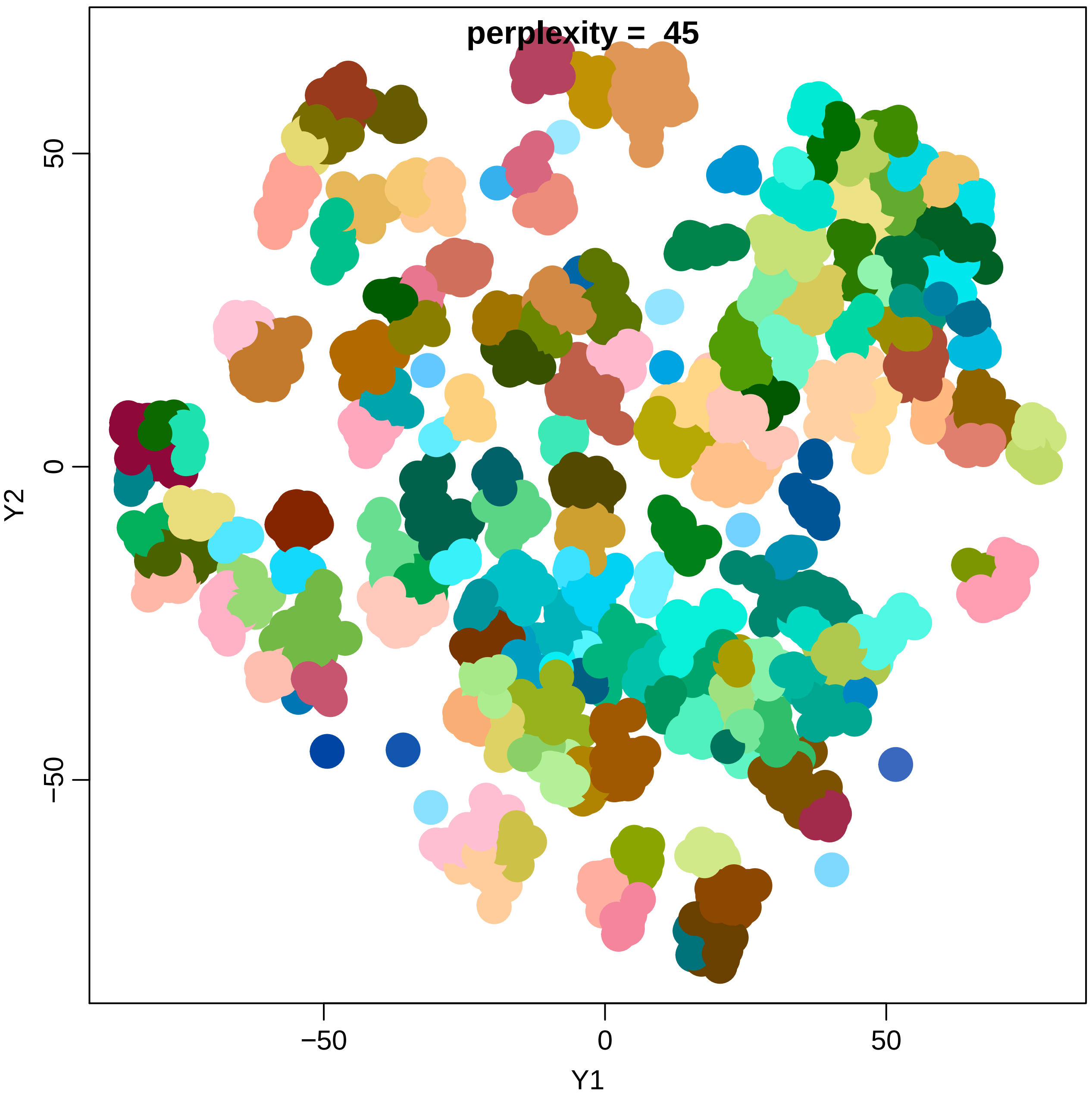
perplexity = 45
Y2
Y1
50
0
−50
−50
0
50

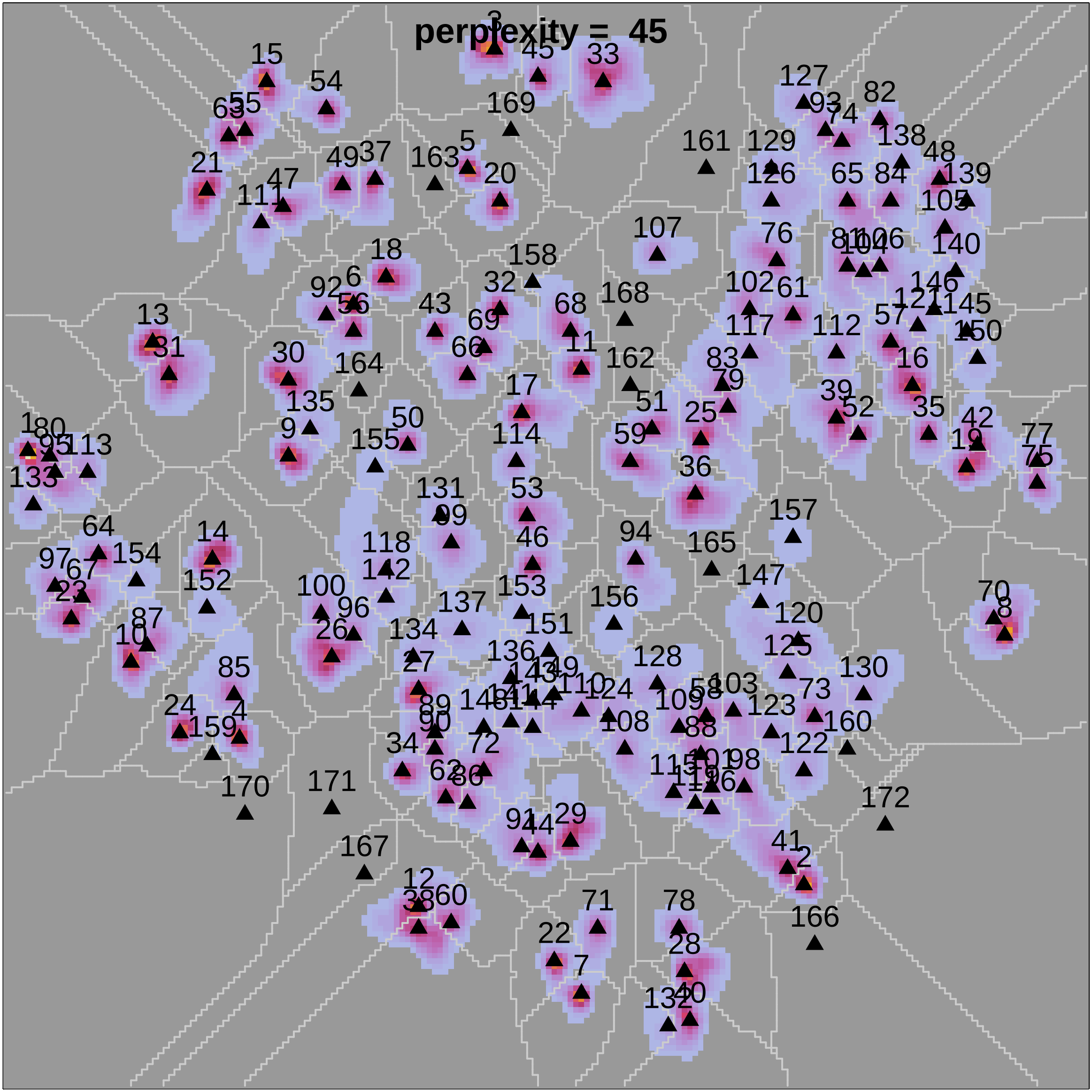
perplexity = 45

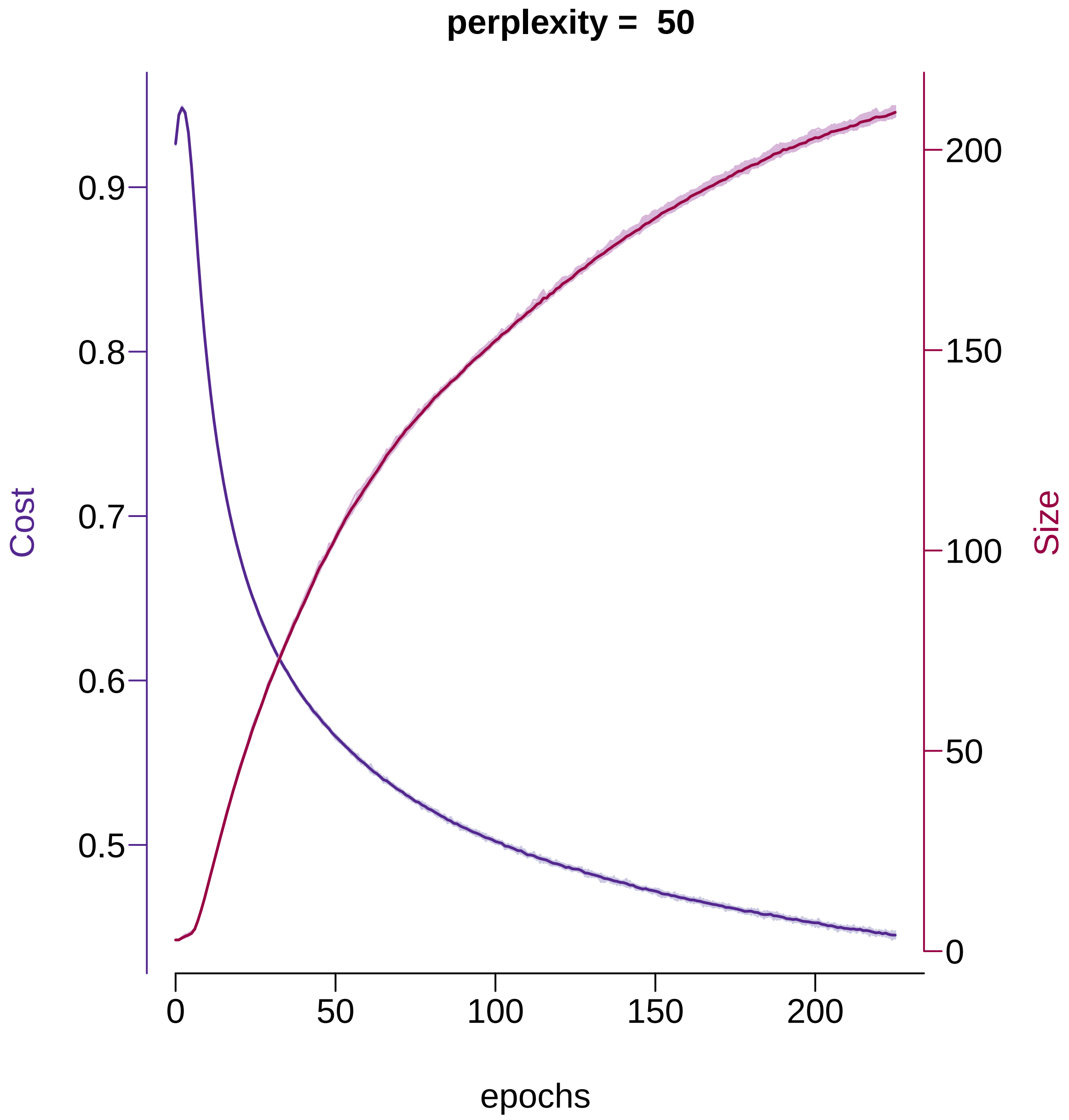
perplexity = 50
Cost
0.9
0.8
0.7
0.6
0.5
Size
200
150
100
50
0
0
50
100
150
200
epochs

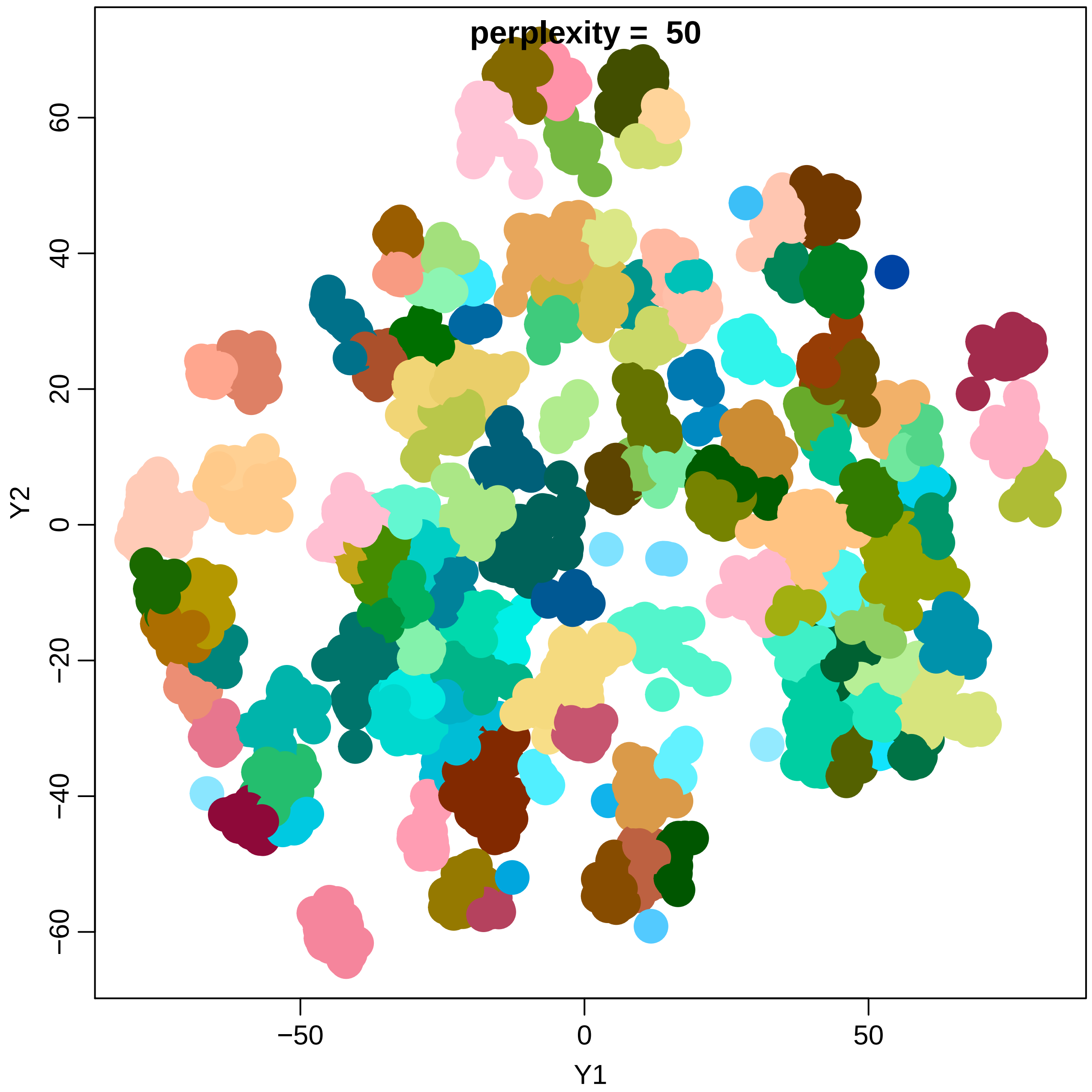

perplexity = 50
Y2
60
40
20
0
−20
−40
−60
Y1
−50
0
50

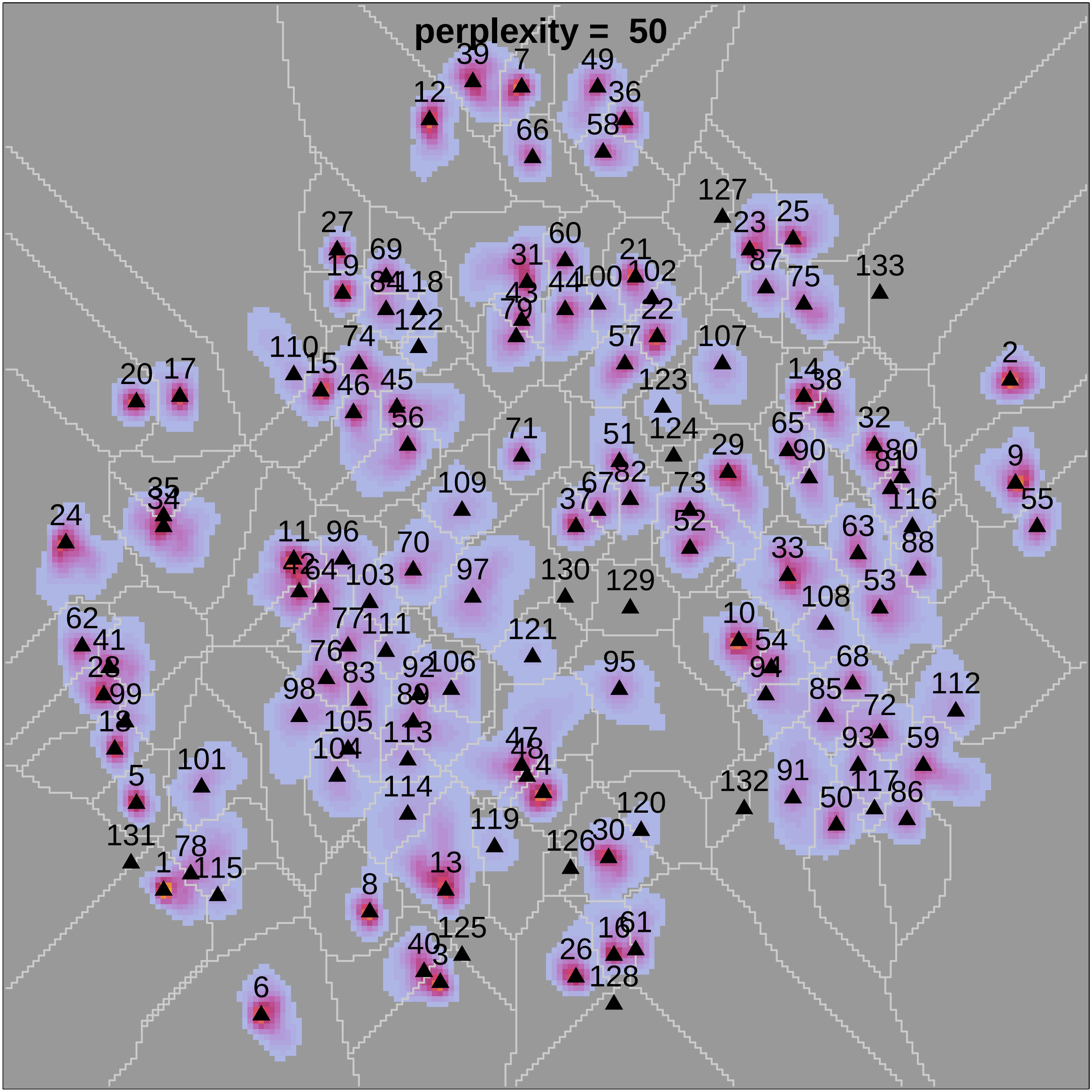

perplexity = 50

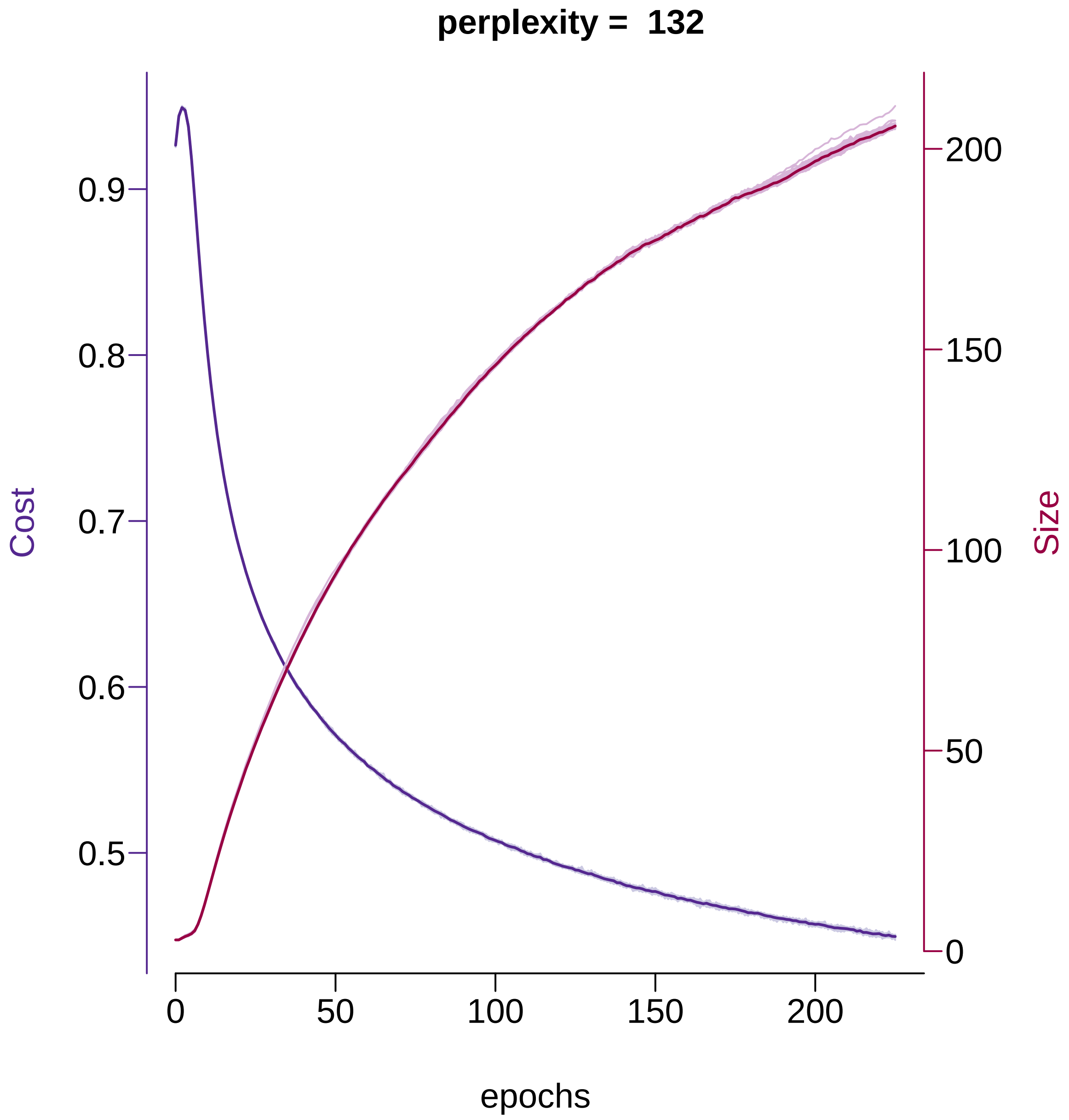
perplexity = 132
Cost
0.9
0.8
0.7
0.6
0.5
Size
200
150
100
50
0
0
50
100
150
200
epochs

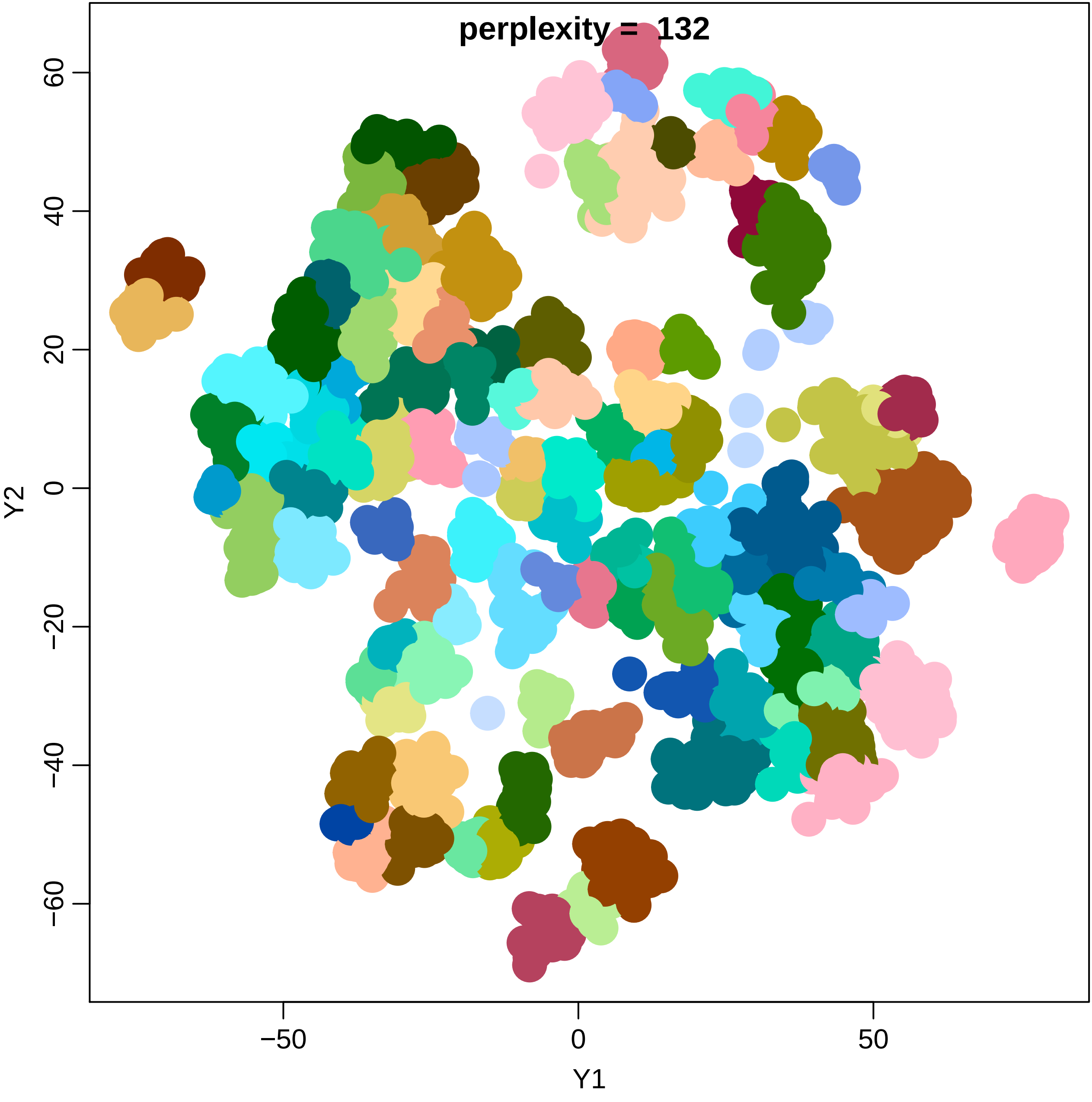

perplexity = 132
60
40
20
0
−20
−40
−60
Y2
−50
0
50
Y1

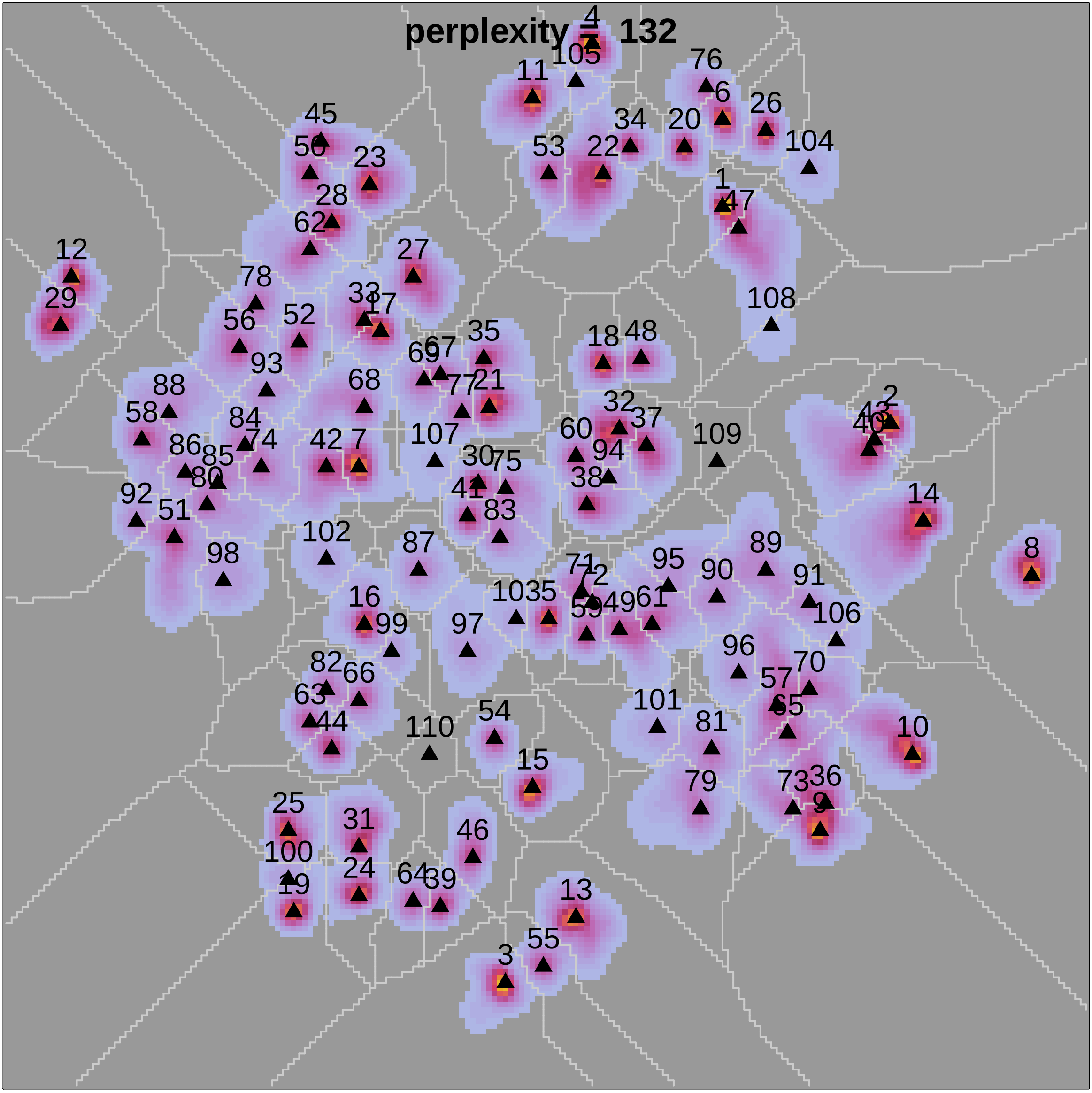
perplexity = 132

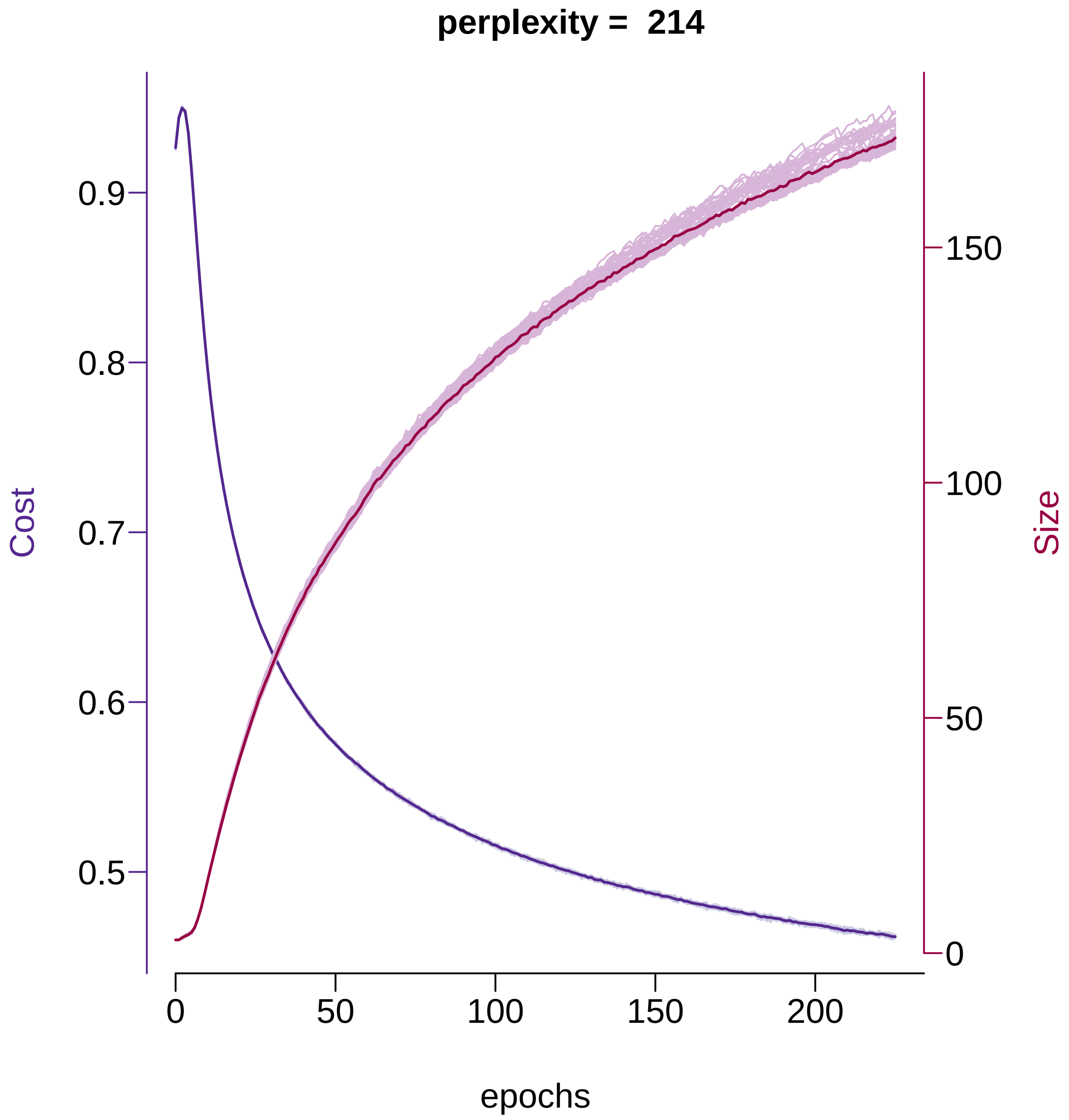

perplexity = 214
Cost
0.9
0.8
0.7
0.6
0.5
Size
150
100
50
0
0
50
100
150
200
epochs

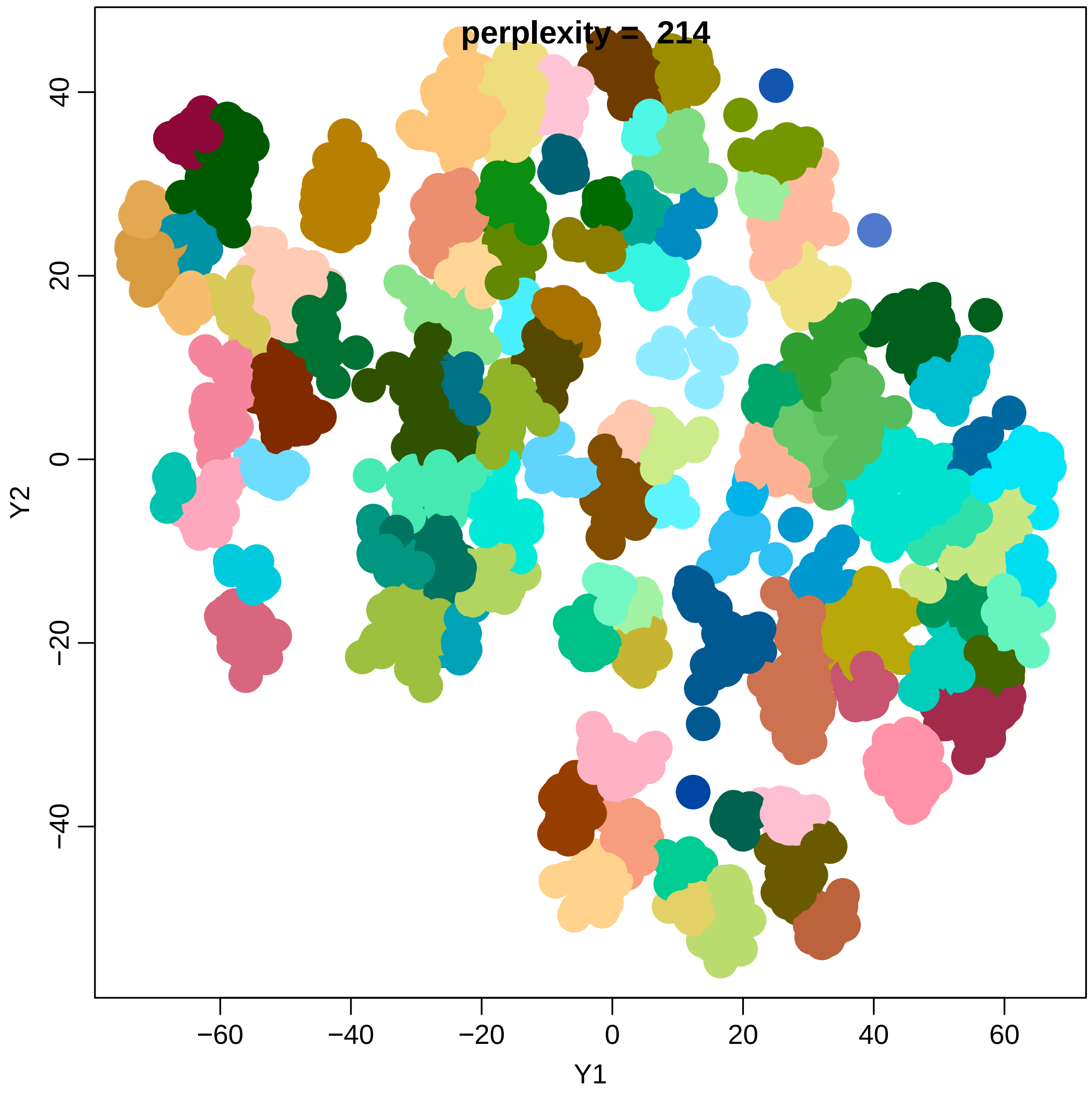
perplexity = 214
Y1
Y2
−60
−40
−20
0
20
40
60
−40
−20
0
20
40

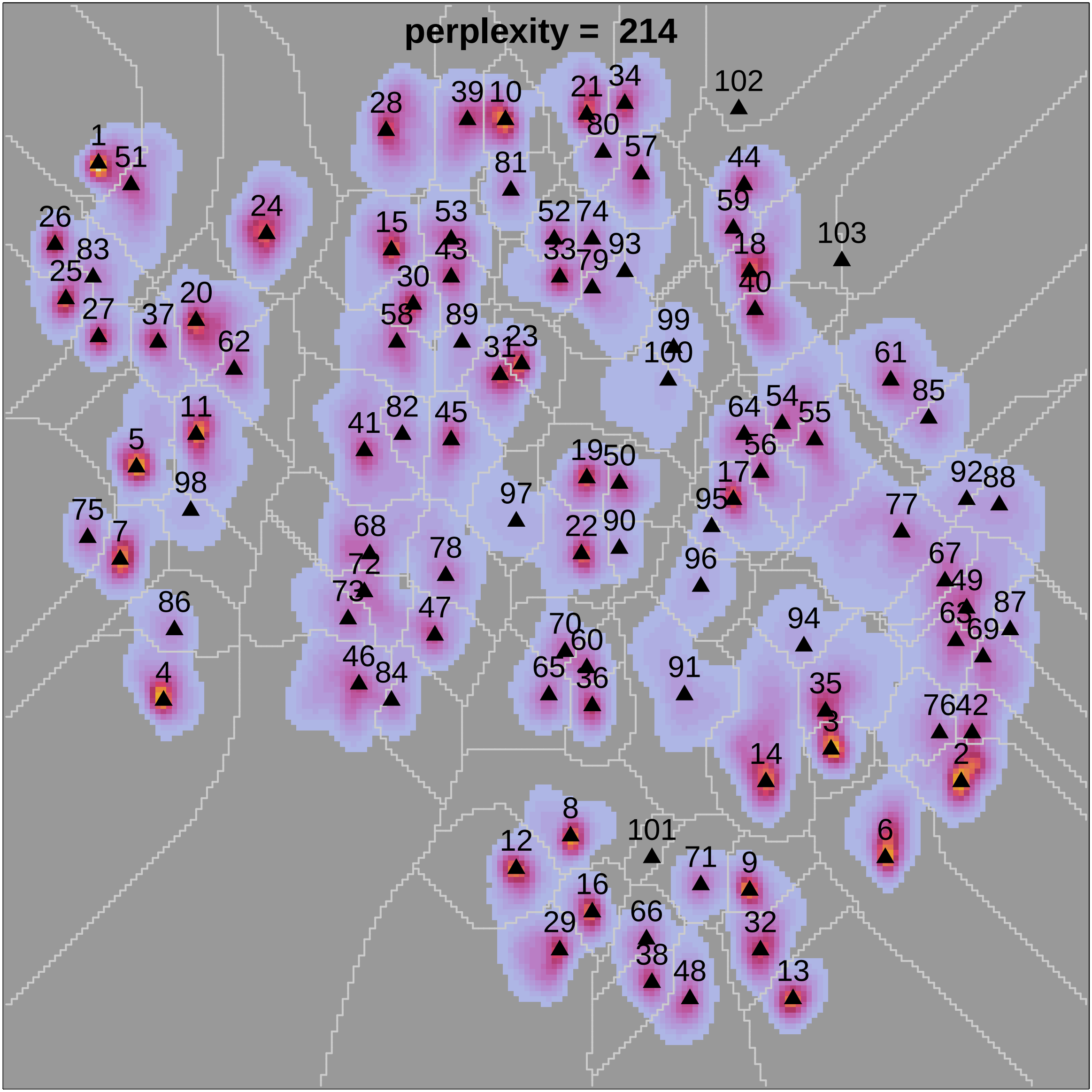
perplexity = 214
1
51
26
83
25
27
37
20
62
24
28
39
10
21
34
80
57
81
15
53
43
30
52
74
33
79
93
58
89
31
23
102
44
59
18
40
103
99
100
61
85
64
54
55
56
17
95
11
5
98
75
7
86
4
41
82
45
19
50
97
22
90
68
72
73
78
47
46
84
70
60
65
36
96
91
94
77
92
88
67
49
63
87
69
35
3
76
42
2
14
6
8
12
101
71
9
16
29
66
38
48
32
13

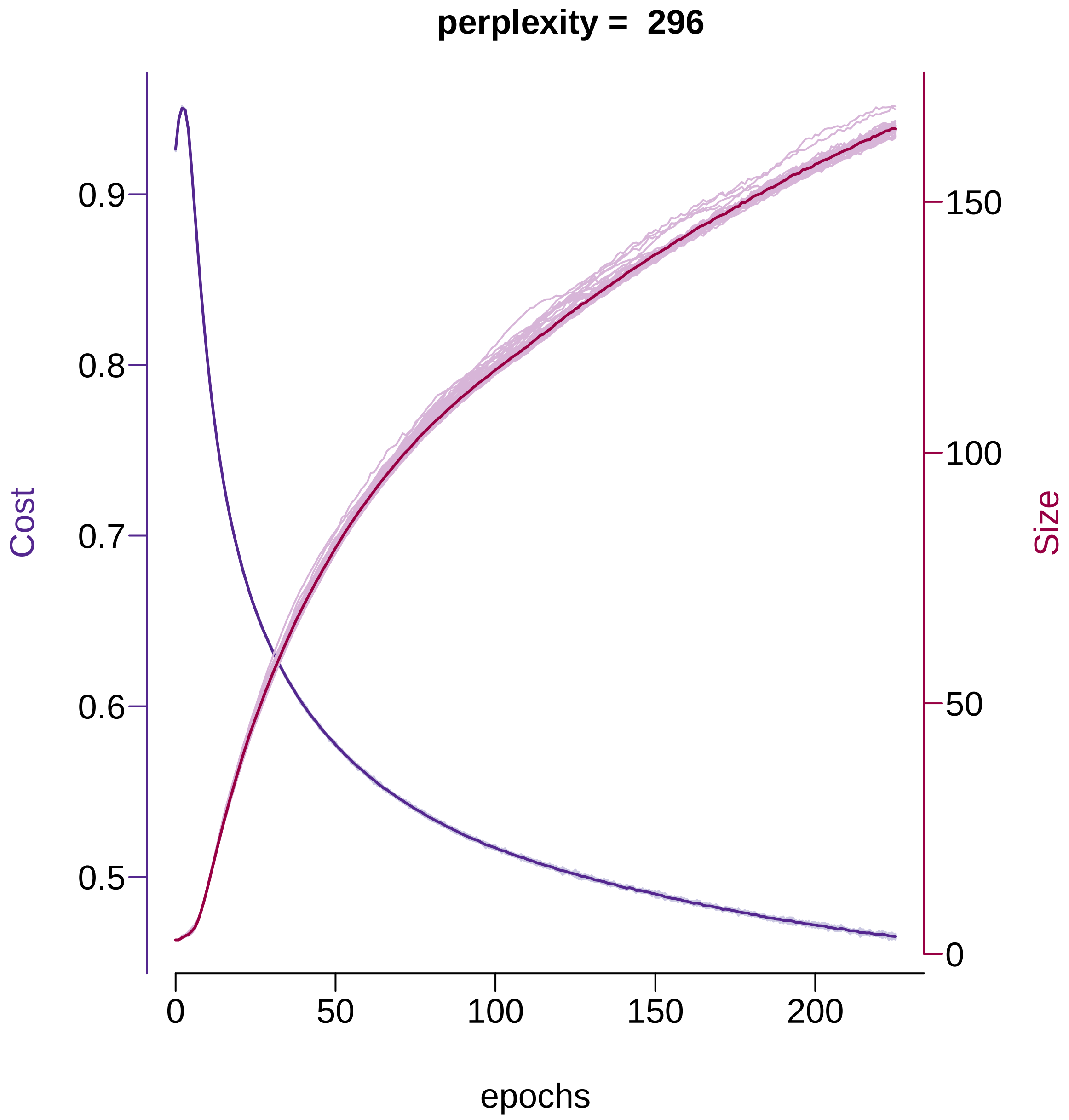
perplexity = 296
Cost
0.9
0.8
0.7
0.6
0.5
Size
150
100
50
0
0
50
100
150
200
epochs

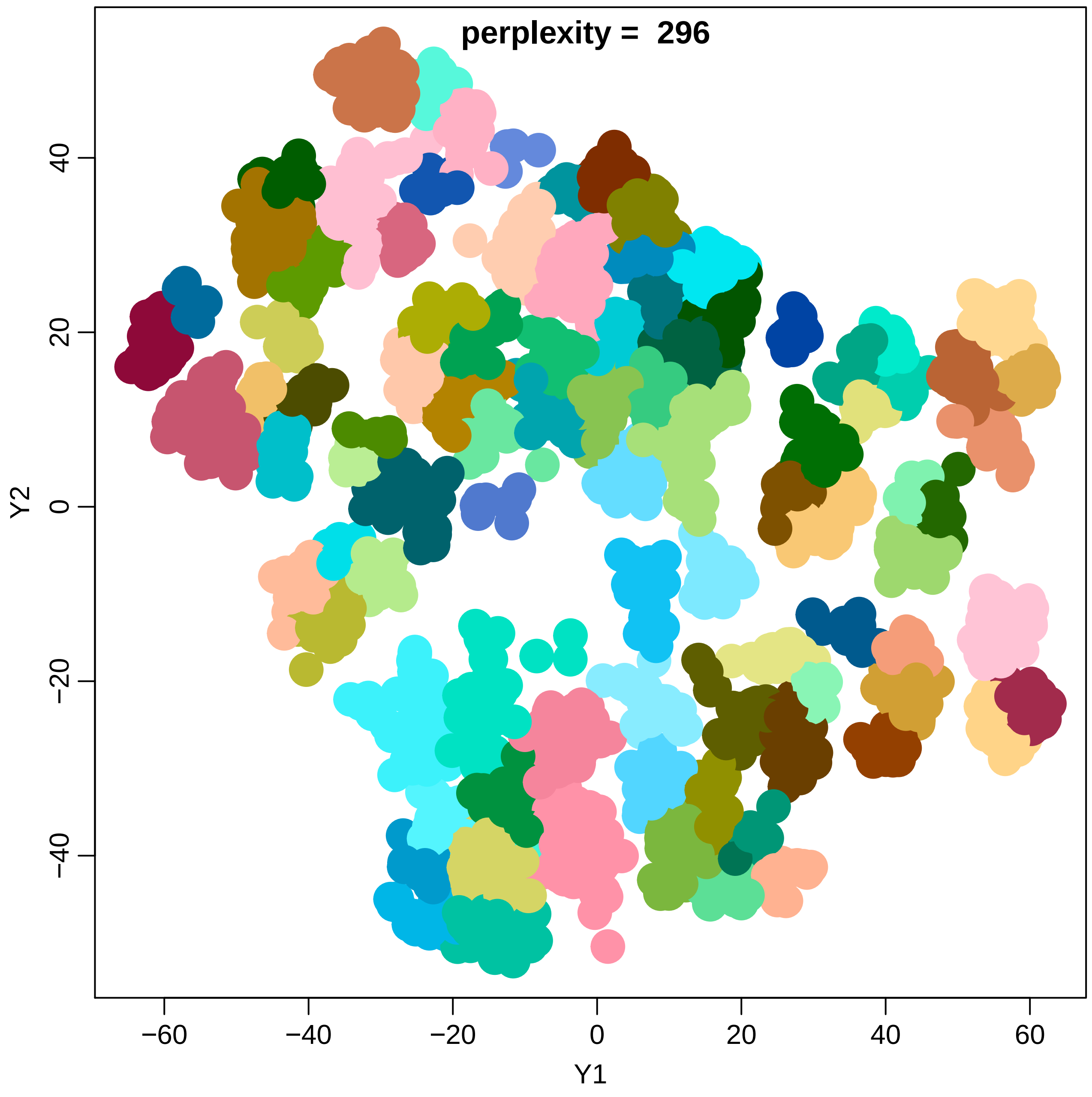

perplexity = 296
Y2
Y1
40
20
0
−20
−40
−60
−40
−20
0
20
40
60

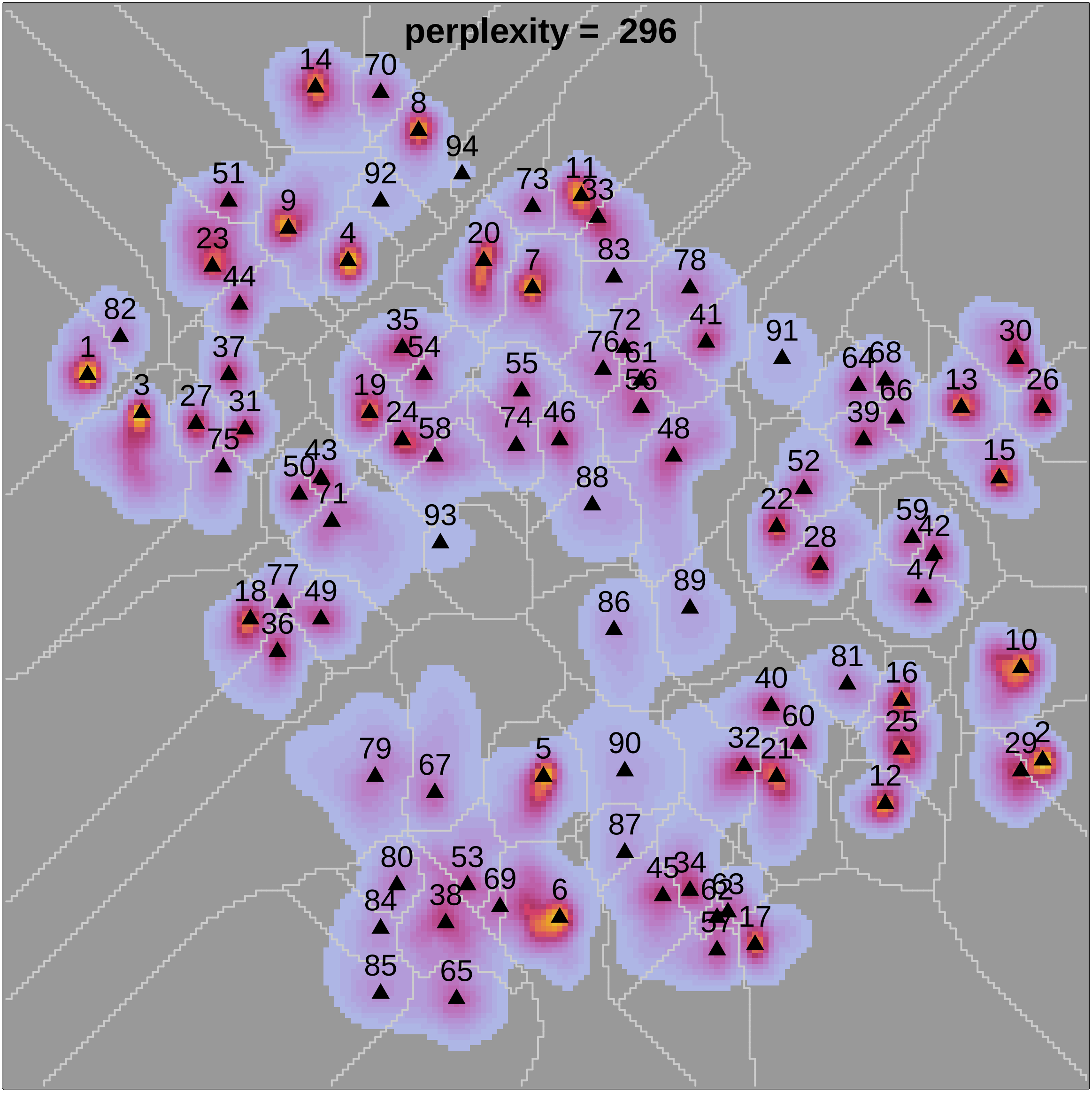
perplexity = 296

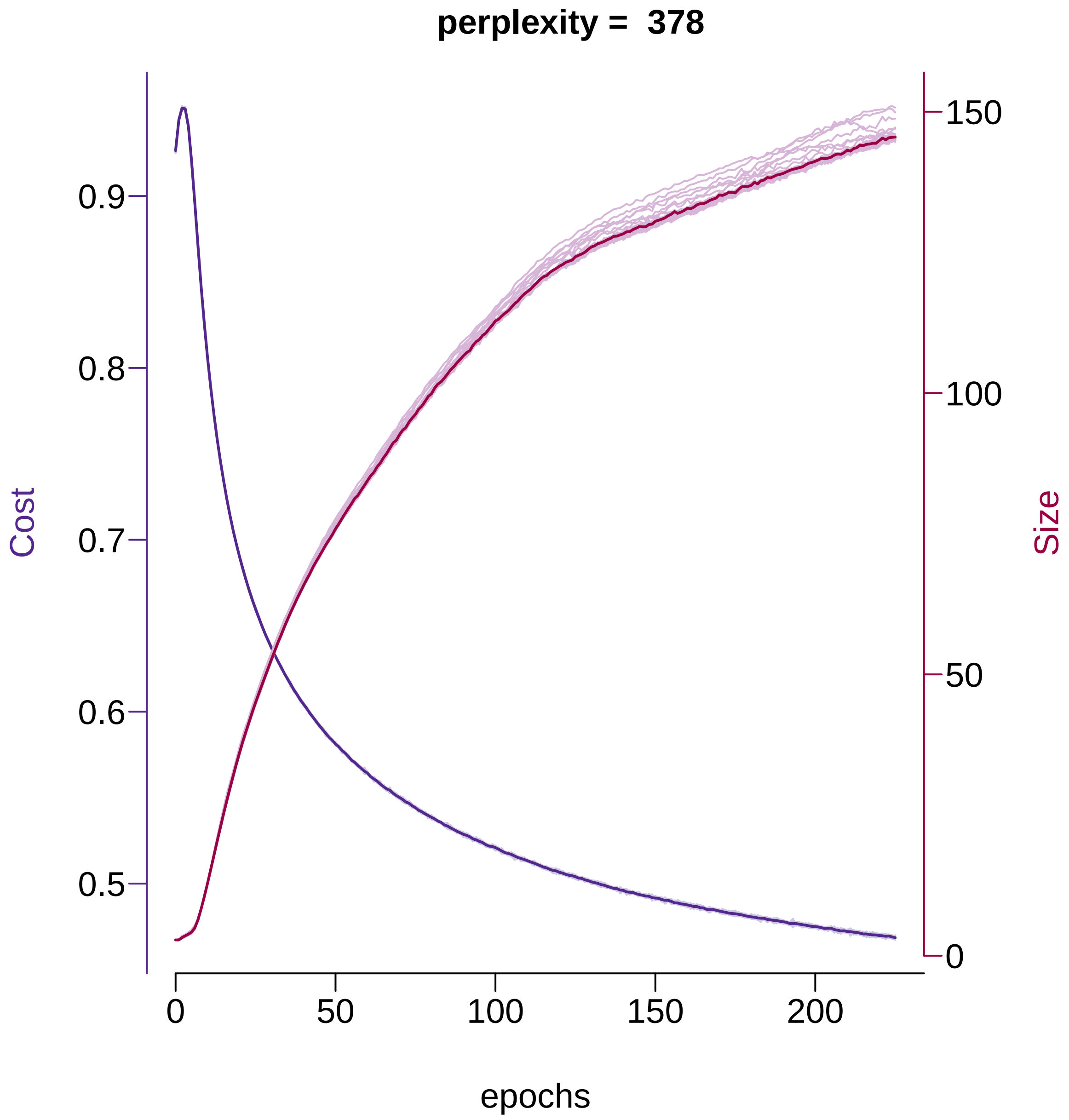
perplexity = 378
Cost
0.9
0.8
0.7
0.6
0.5
Size
150
100
50
0
0
50
100
150
200
epochs

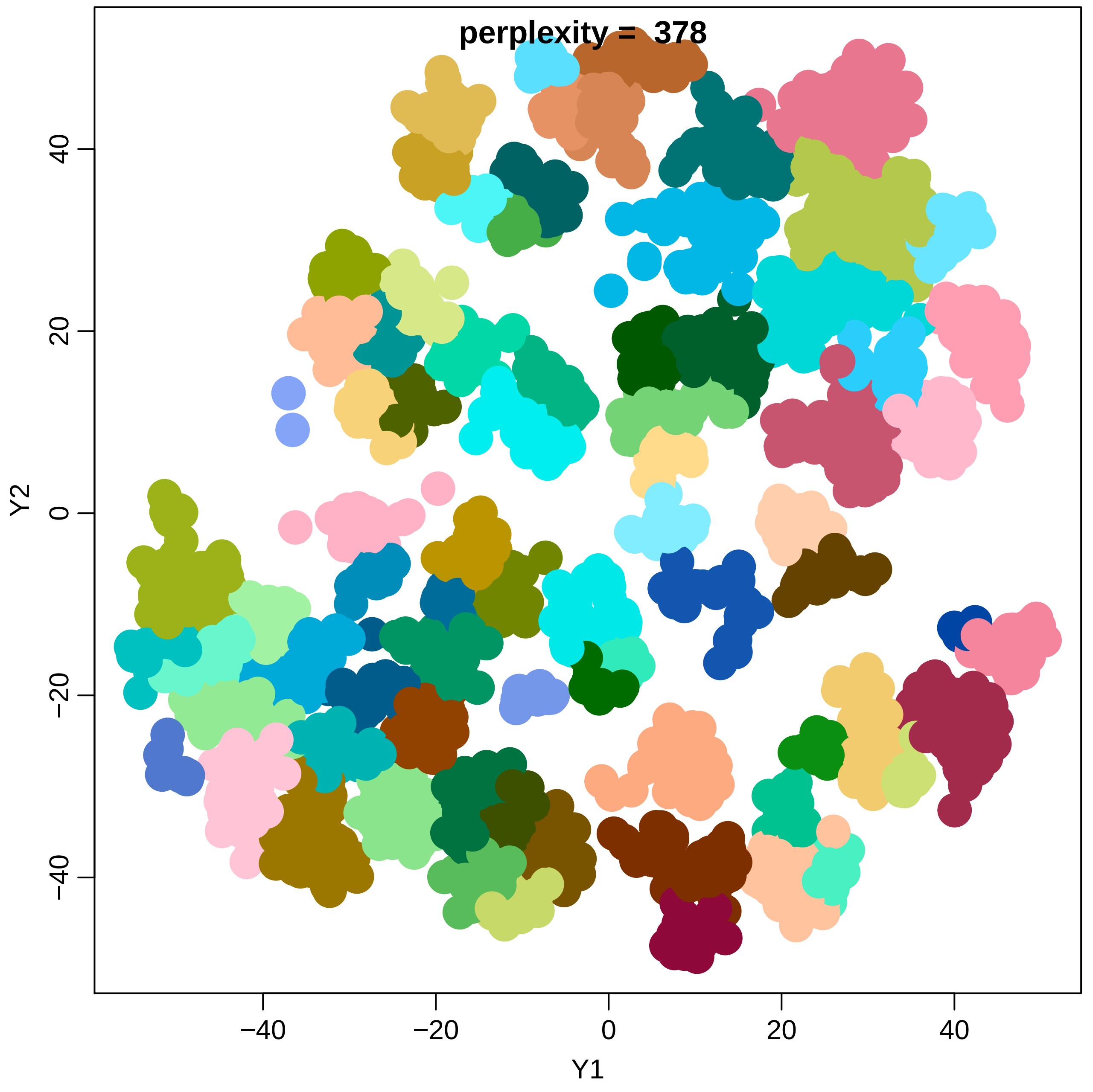
perplexity = 378
Y2
Y1
40
20
0
−20
−40
−40
−20
0
20
40

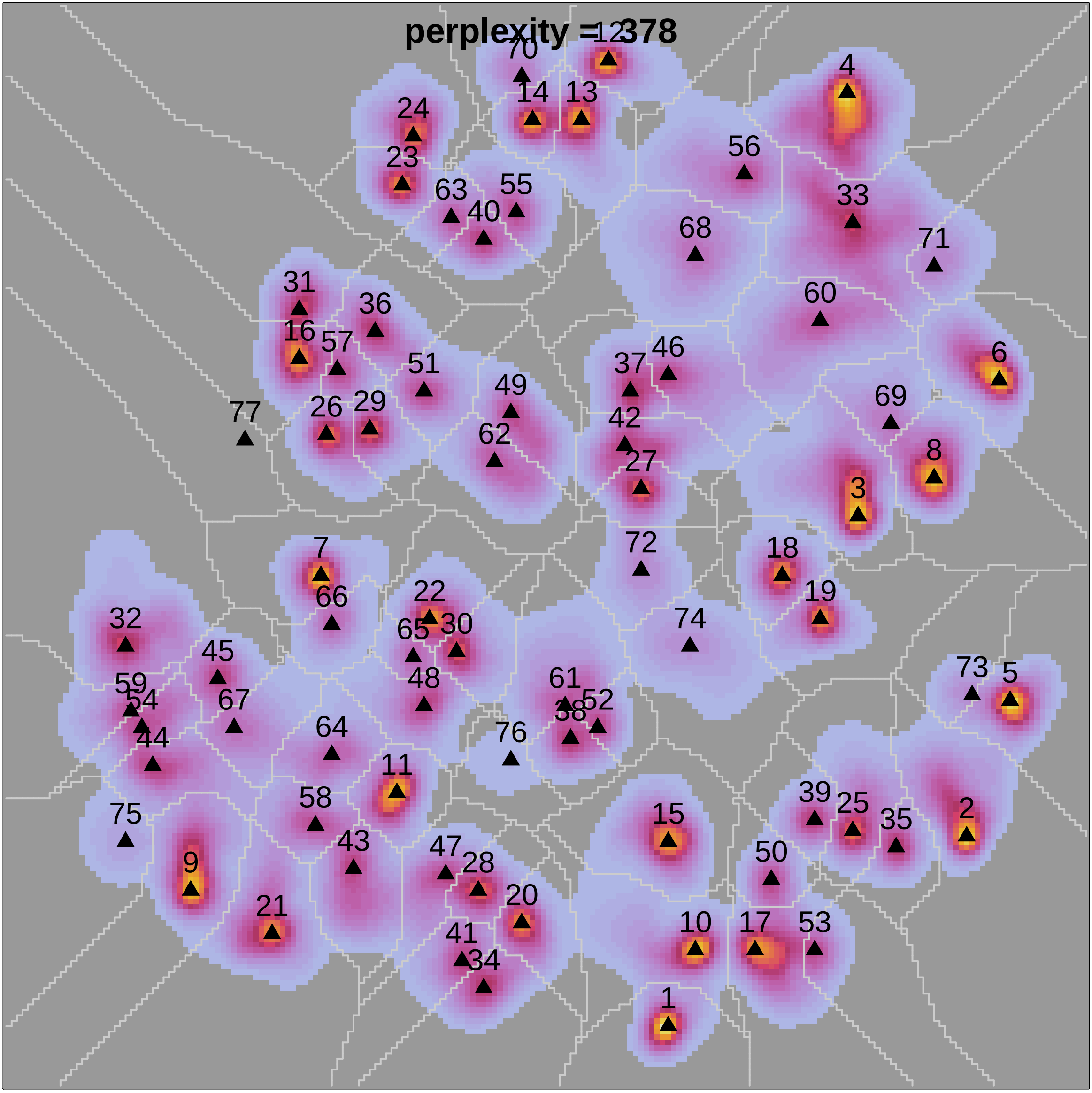

perplexity = 378

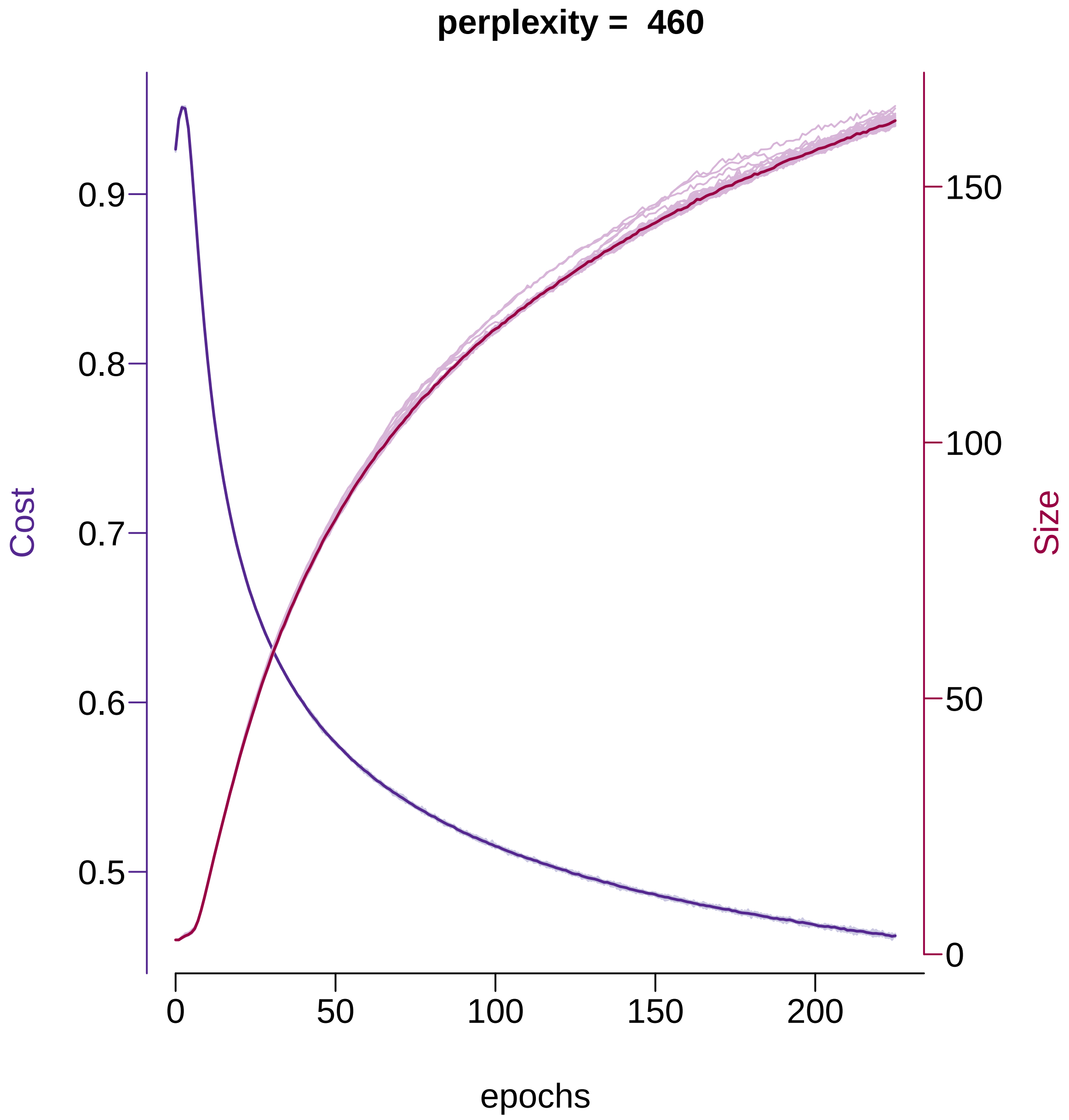

perplexity = 460
Cost
0.9
0.8
0.7
0.6
0.5
Size
150
100
50
0
0
50
100
150
200
epochs

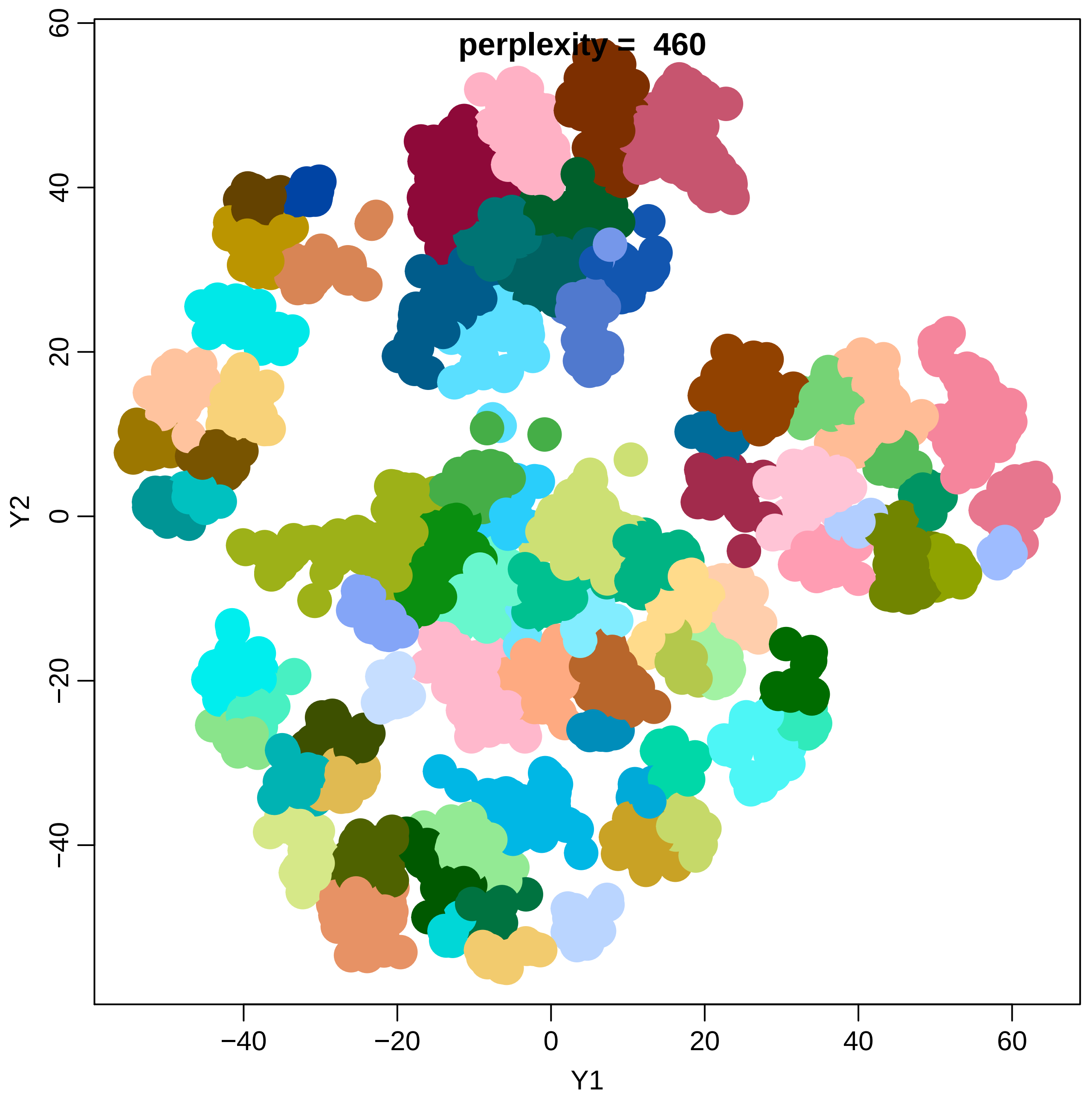

perplexity = 460
Y1
Y2
−40
−20
0
20
40
60

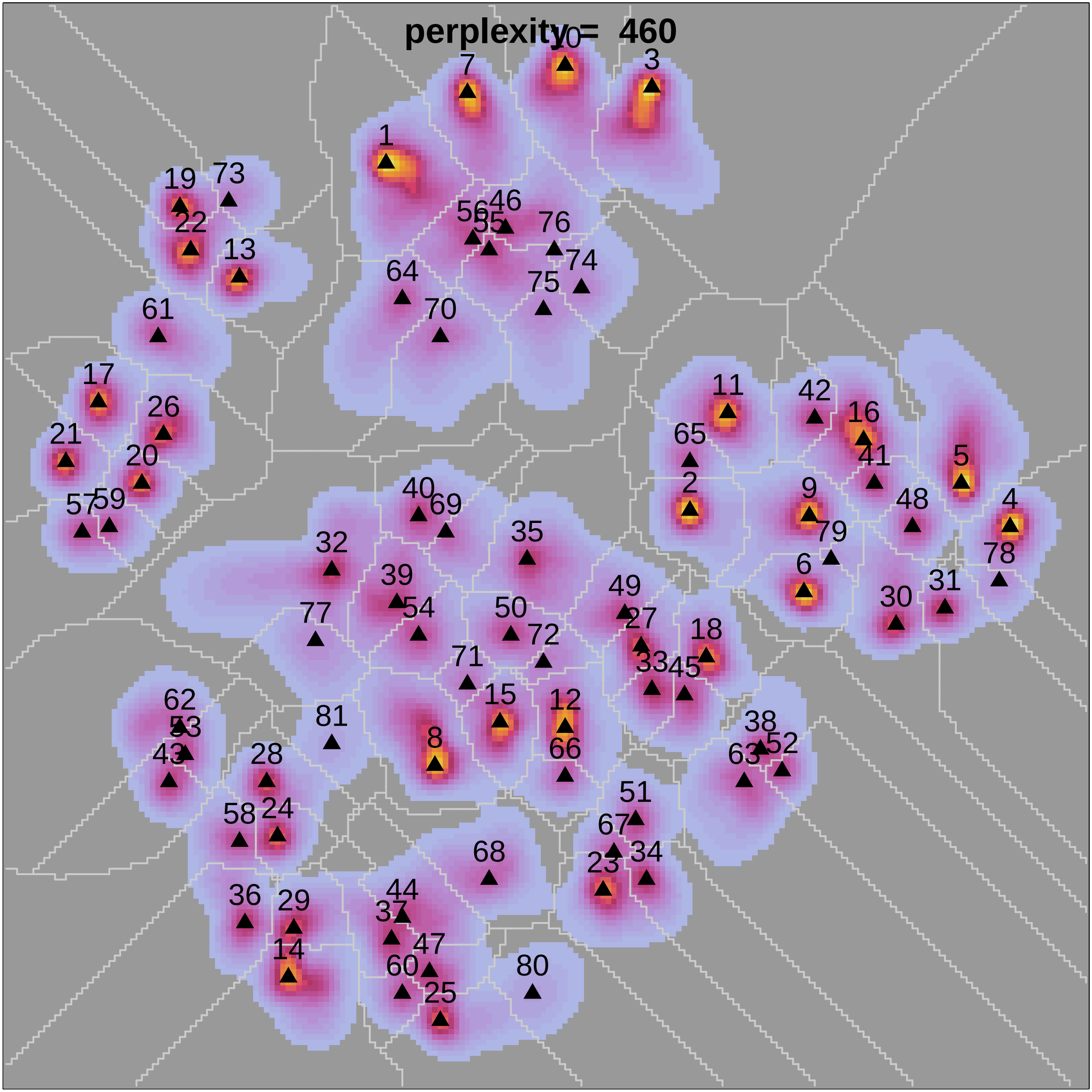
perplexity = 460
10
3
7
1
19
73
46
56
55
76
22
13
74
64
75
70
61
17
11
42
26
16
21
65
20
5
41
2
9
57
59
40
69
48
4
32
35
79
78
6
39
31
49
30
77
54
50
27
72
18
71
33
45
62
15
12
81
53
38
8
43
28
63
52
66
51
58
24
67
68
34
23
44
36
29
37
47
14
60
80
25

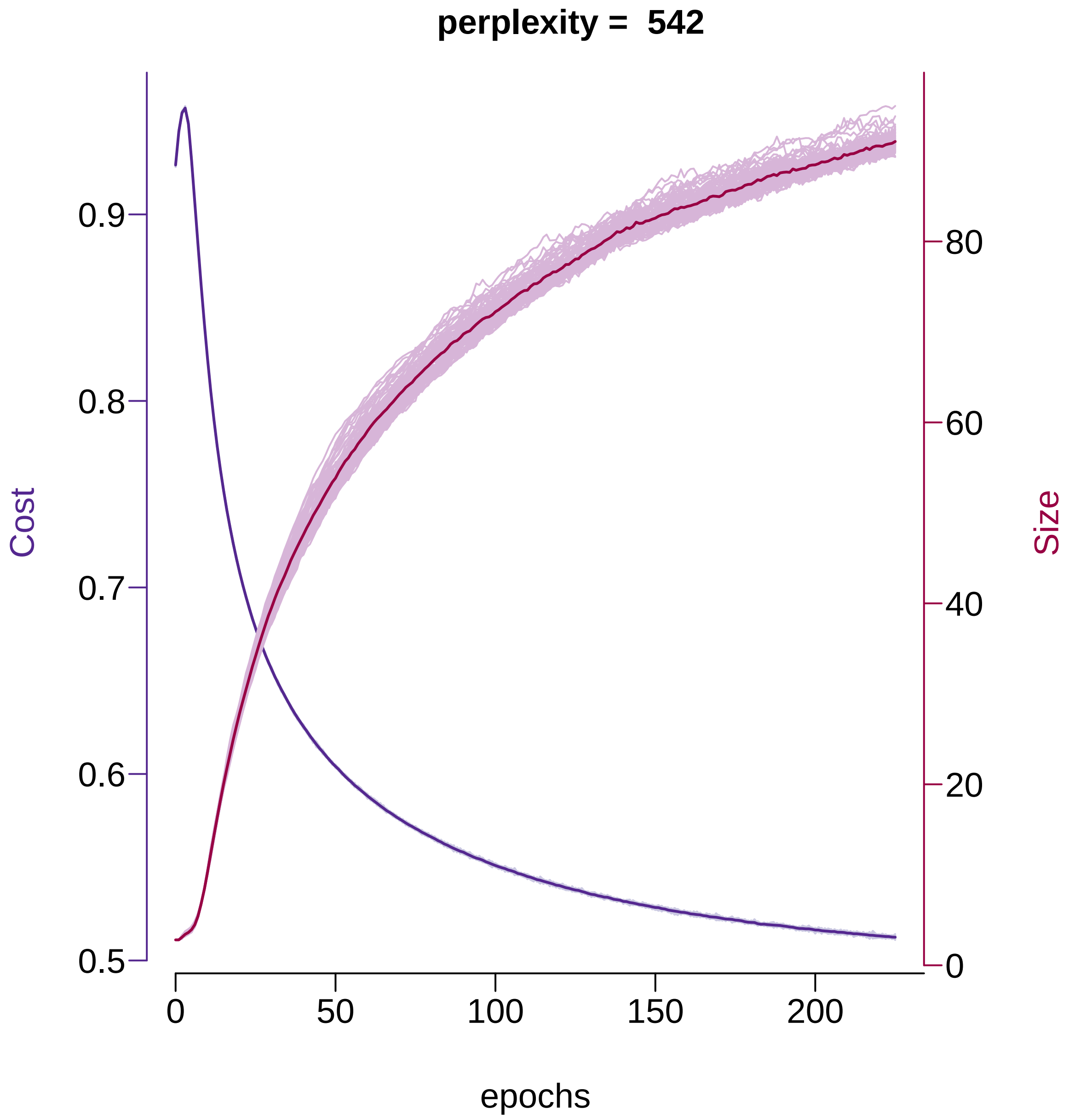

perplexity = 542
Cost
Size
epochs
0.5
0.6
0.7
0.8
0.9
0
20
40
60
80
0
50
100
150
200

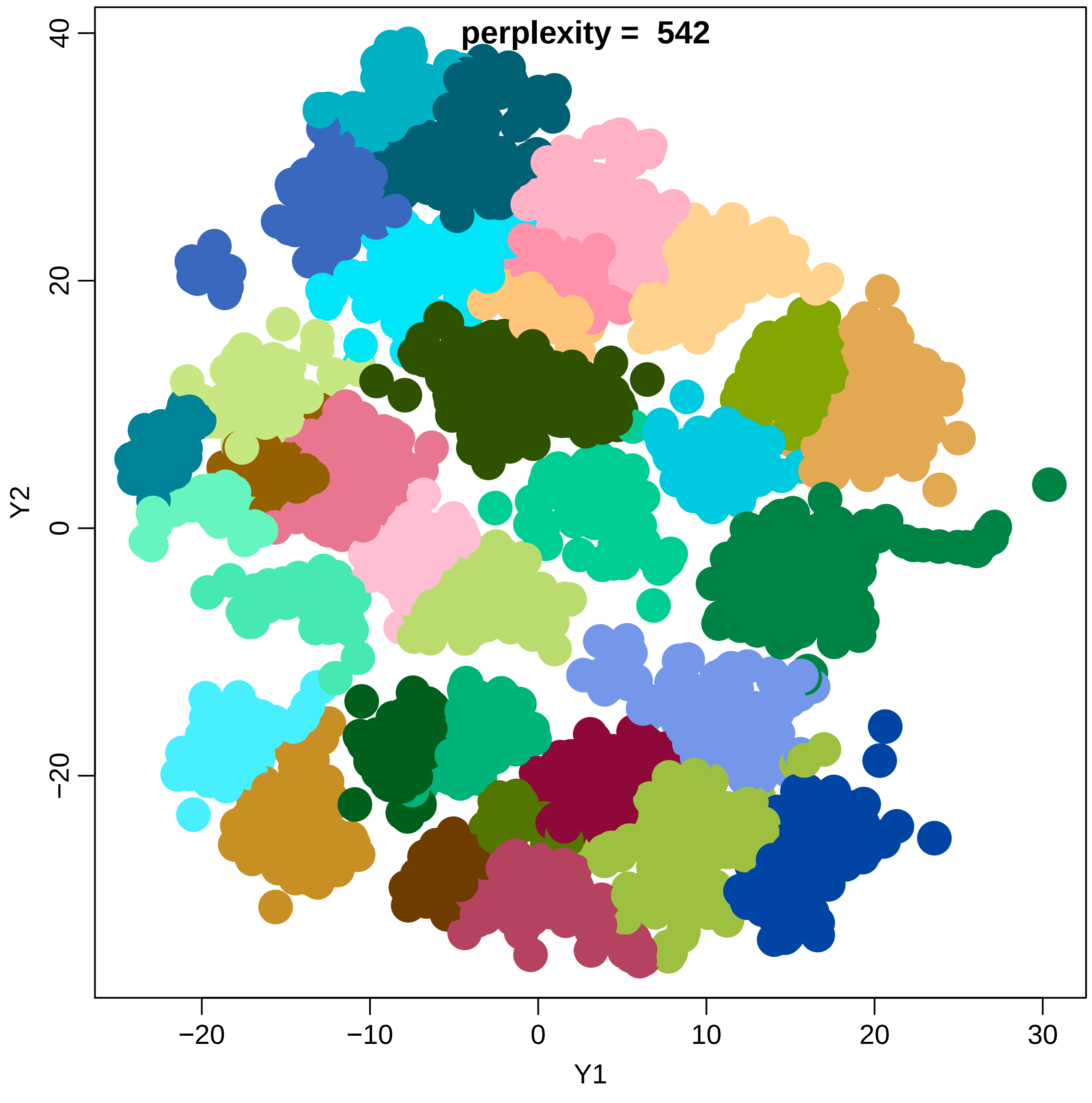
perplexity = 542
Y2
40
20
0
−20
Y1
−20
−10
0
10
20
30

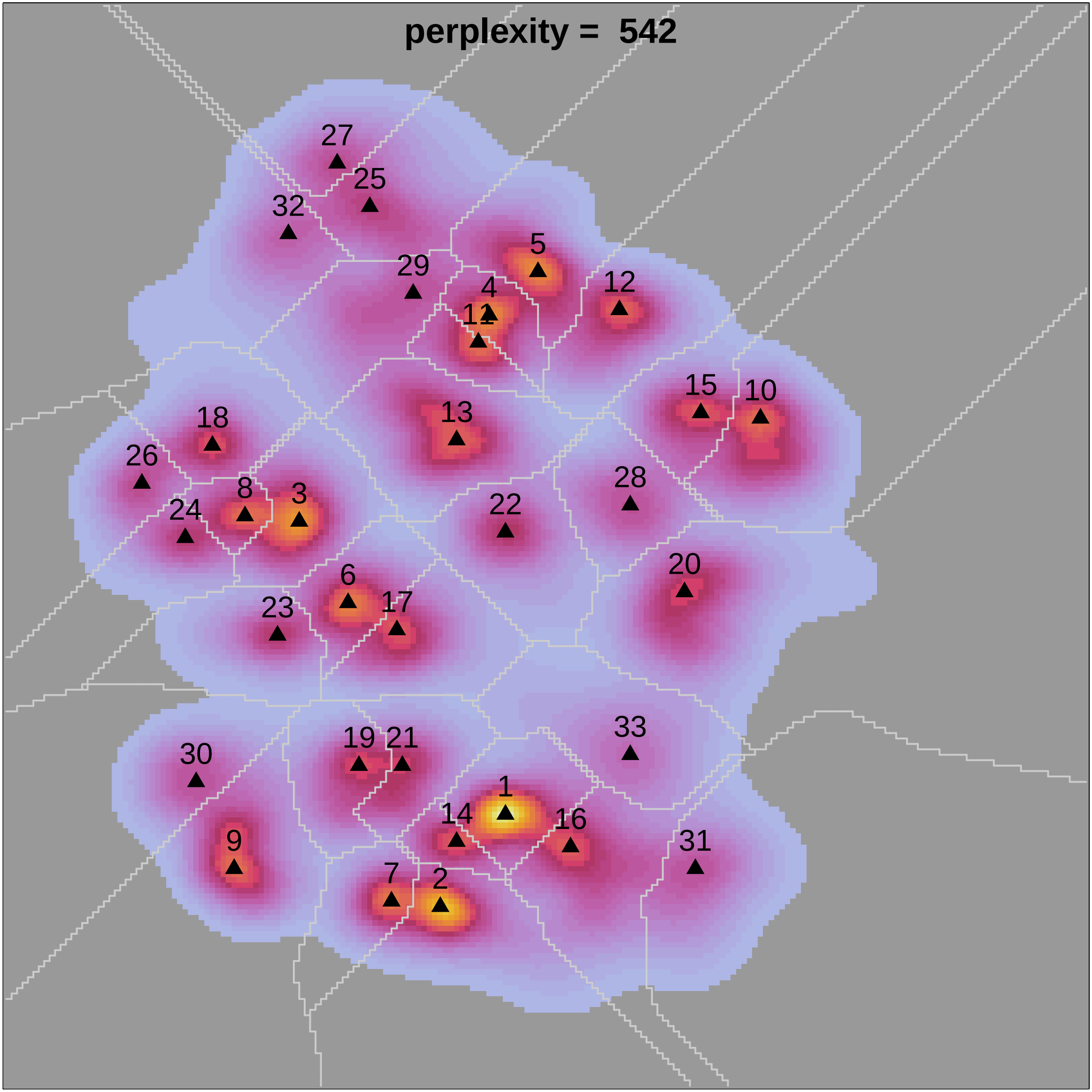
perplexity = 542
27
25
32
29
5
4
11
12
15
10
18
13
26
28
8
3
24
22
20
6
17
23
33
19
21
30
1
14
16
9
31
7
2

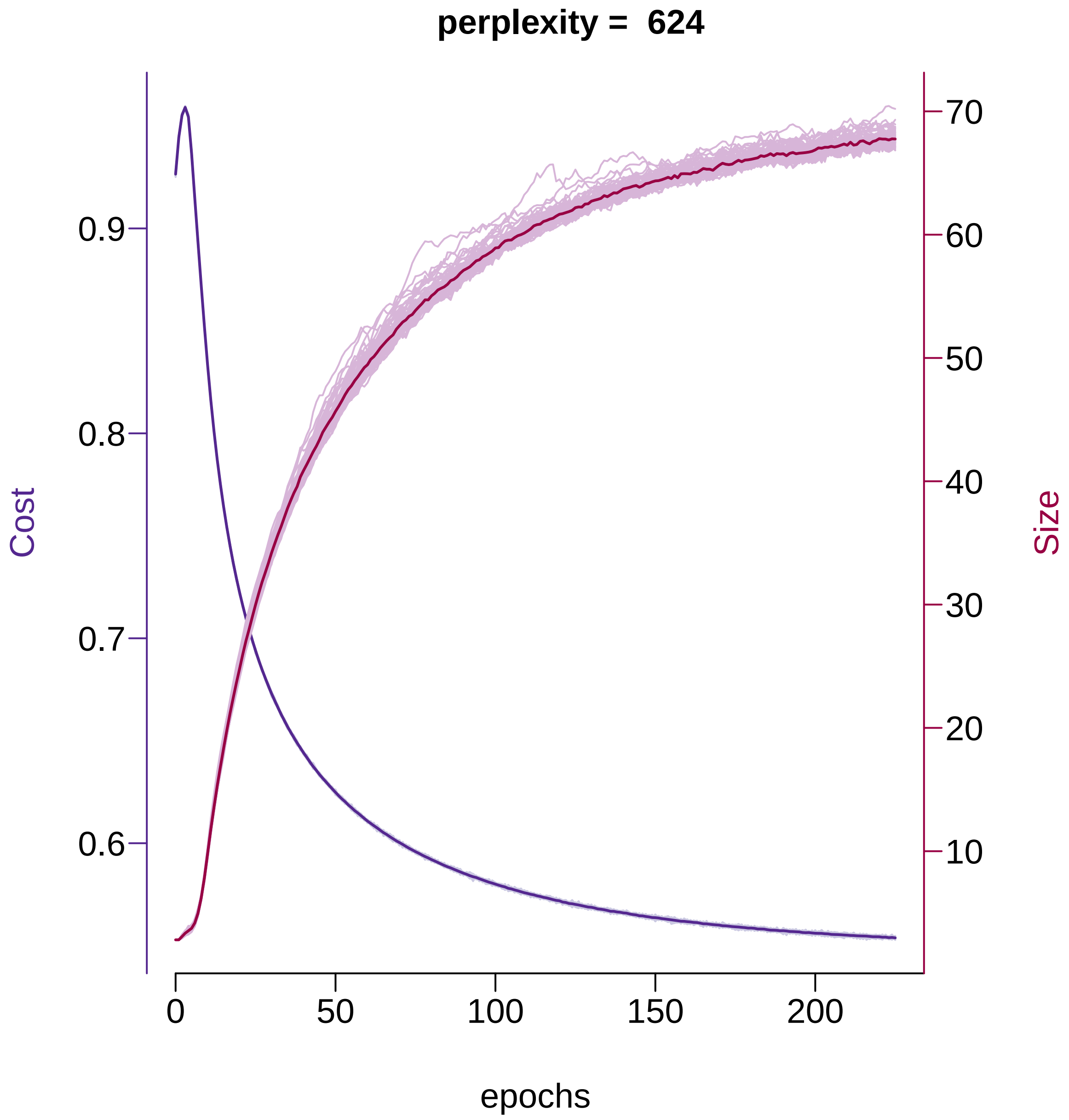

perplexity = 624
Cost
0.9
0.8
0.7
0.6
Size
70
60
50
40
30
20
10
0
50
100
150
200
epochs

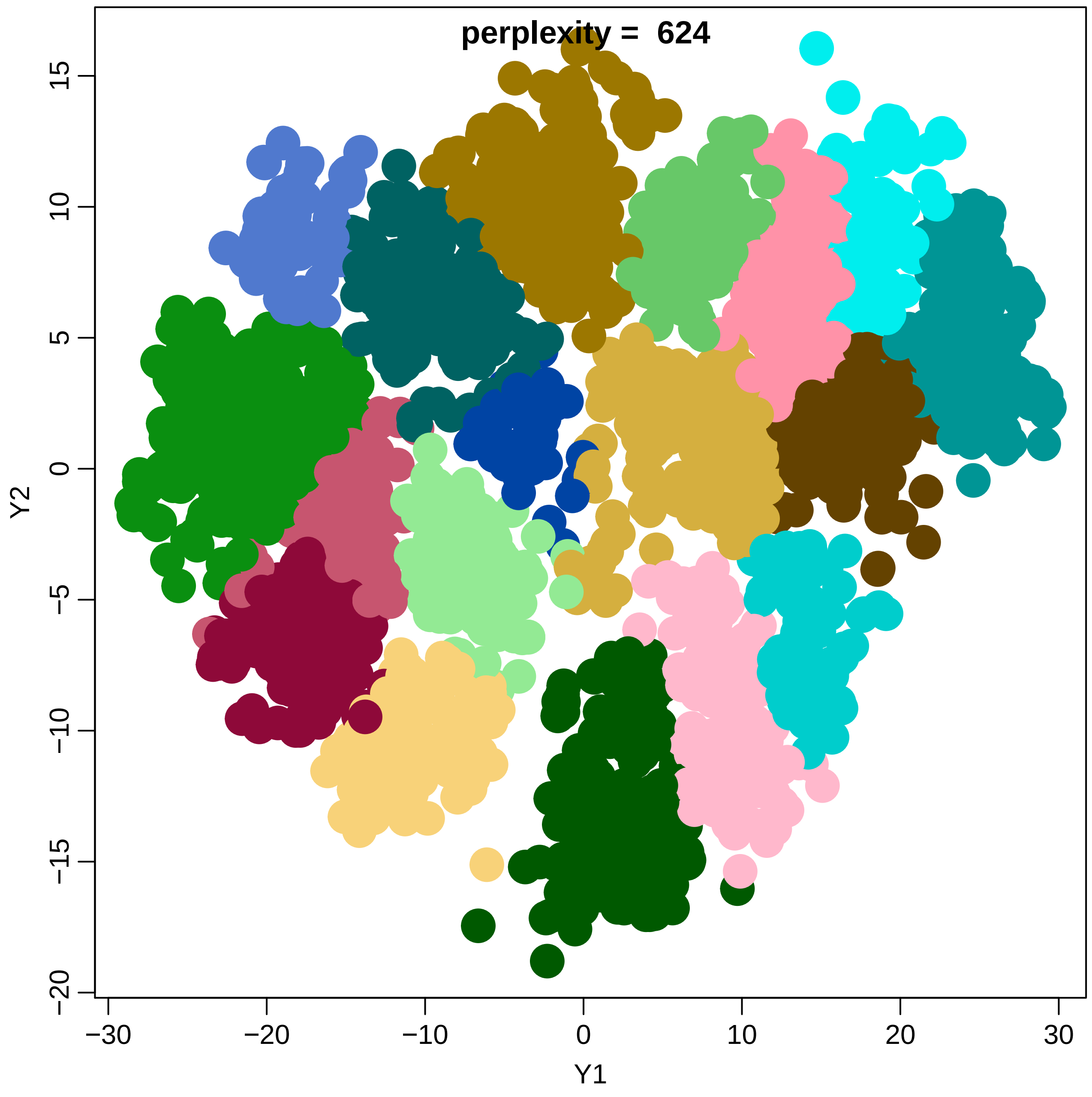

perplexity = 624
Y1
Y2
−30
−20
−10
0
10
20
30
−20
−15
−10
−5
0
5
10
15

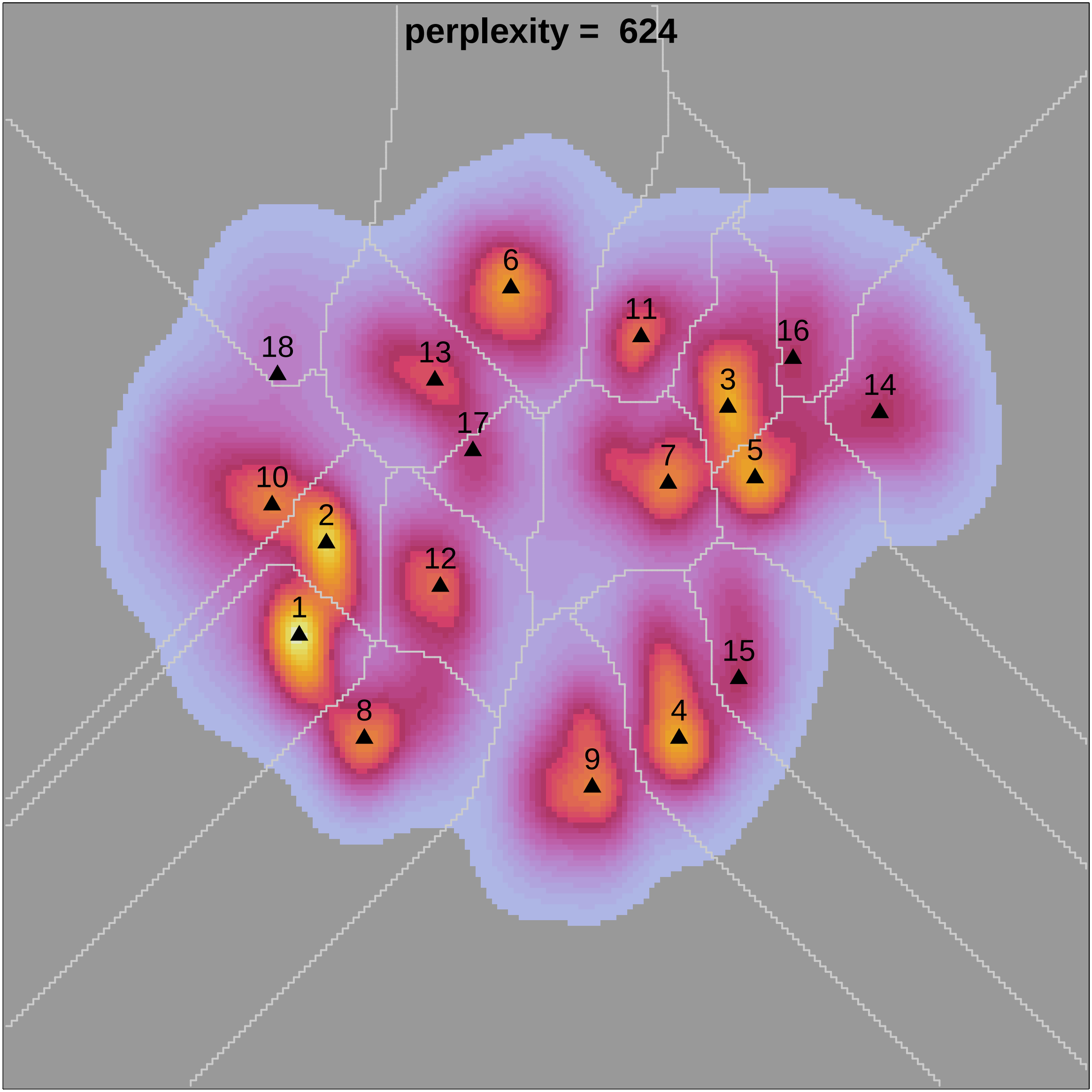
perplexity = 624
6
11
16
18
13
3
14
17
7
5
10
2
12
1
15
8
4
9

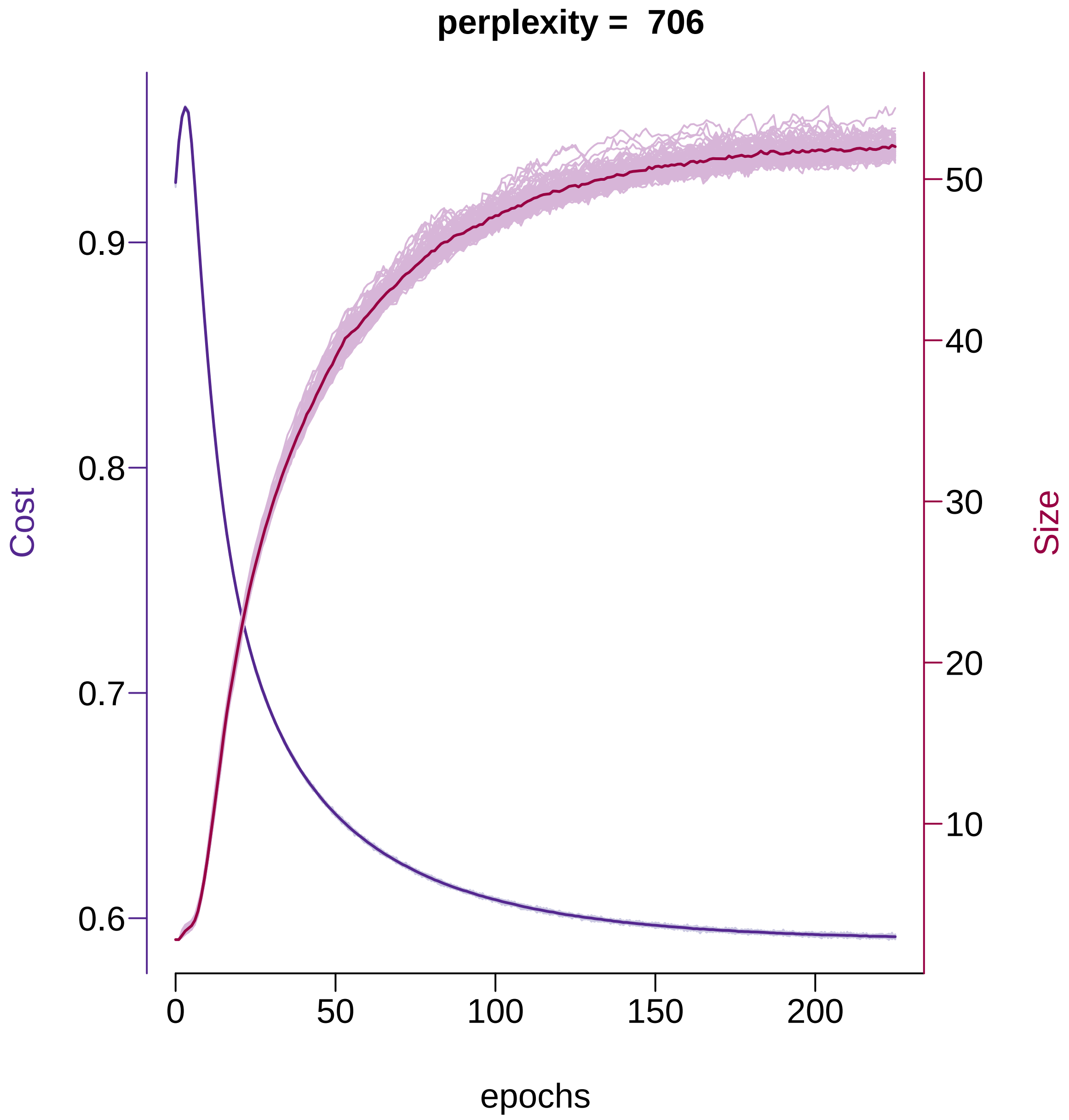
perplexity = 706
Cost
Size
epochs
0.6
0.7
0.8
0.9
10
20
30
40
50
0
50
100
150
200

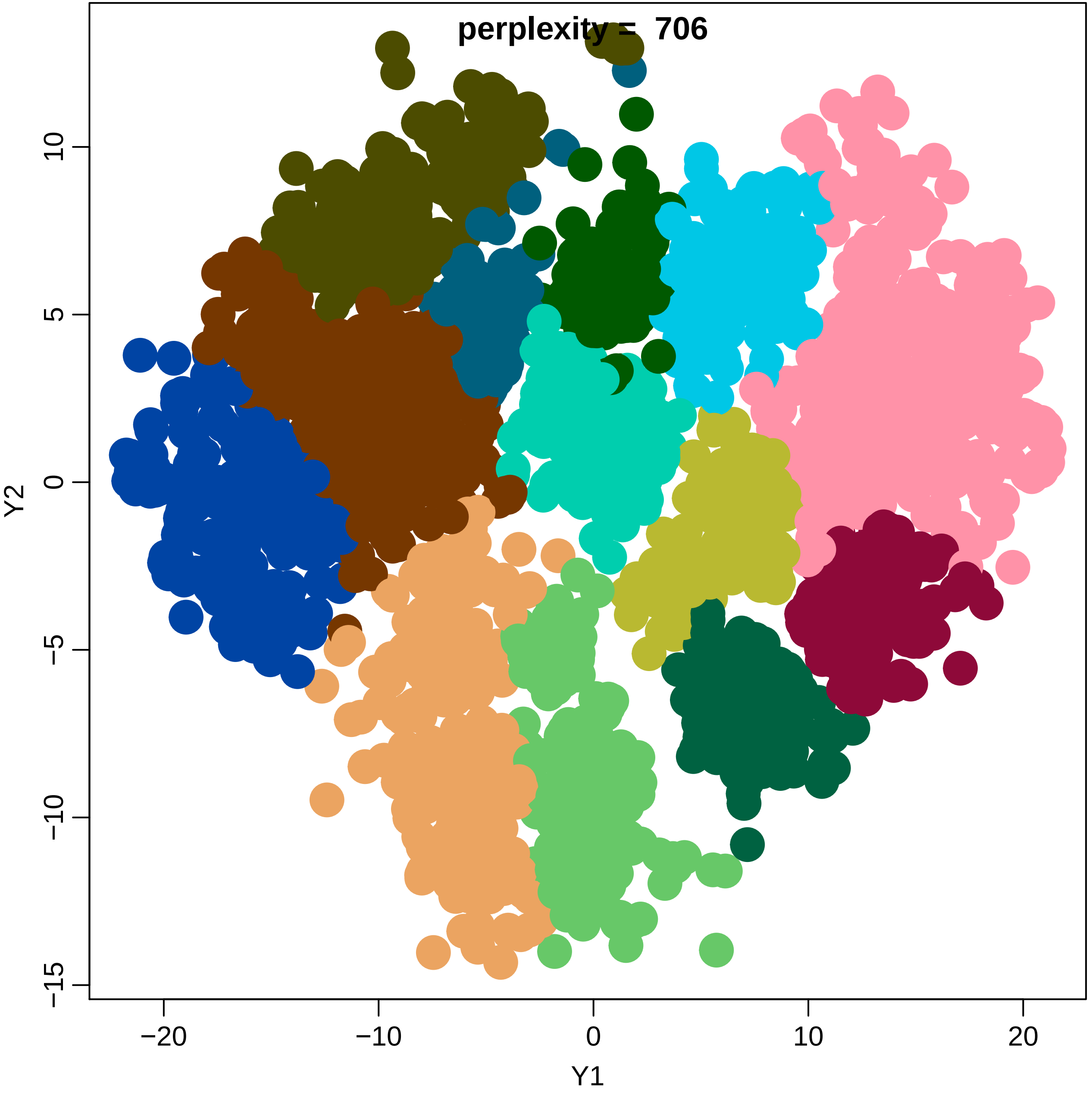

perplexity = 706
10
5
0
−5
−10
−15
Y2
−20
−10
0
10
20
Y1

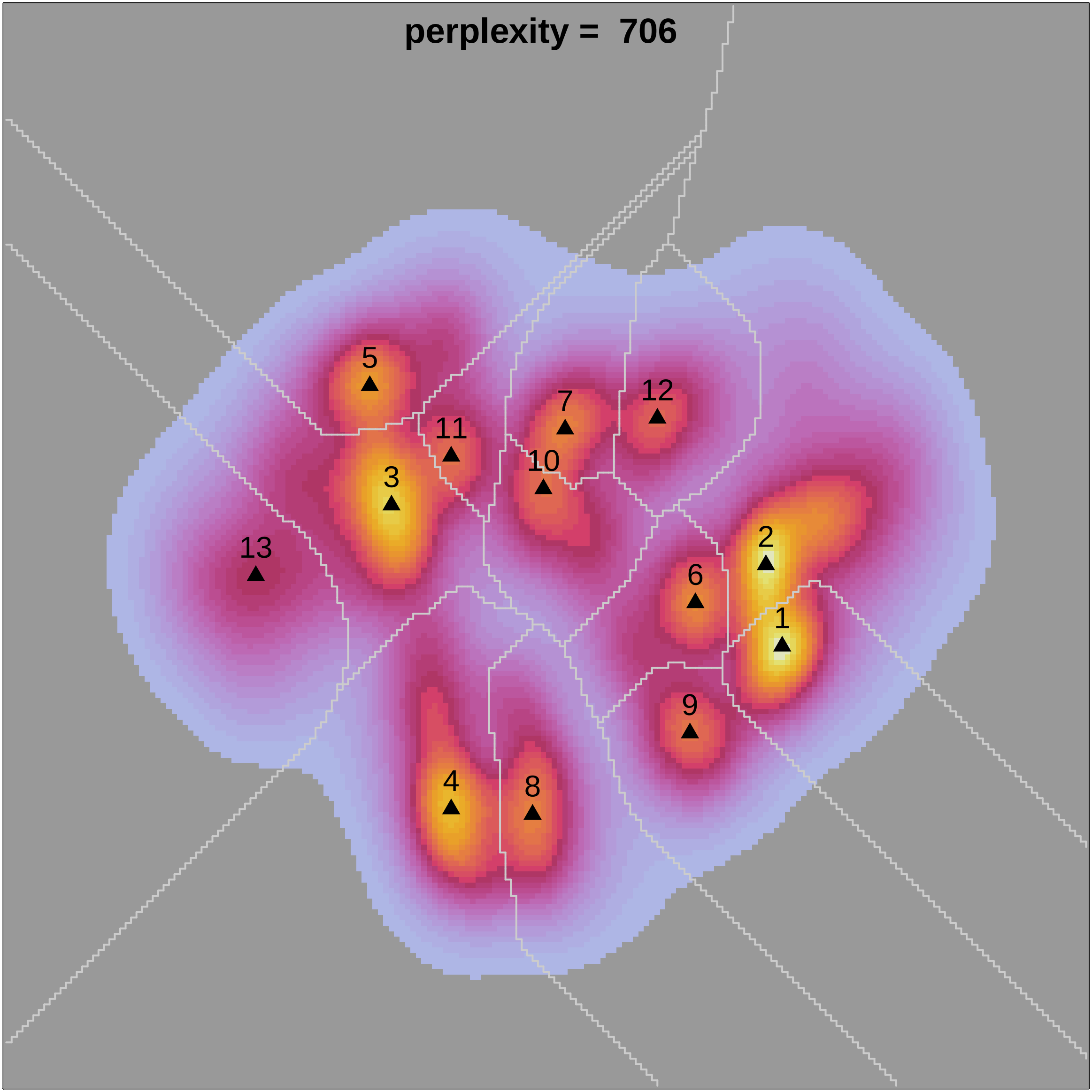
perplexity = 706
5
12
7
11
10
3
13
2
6
1
9
4
8

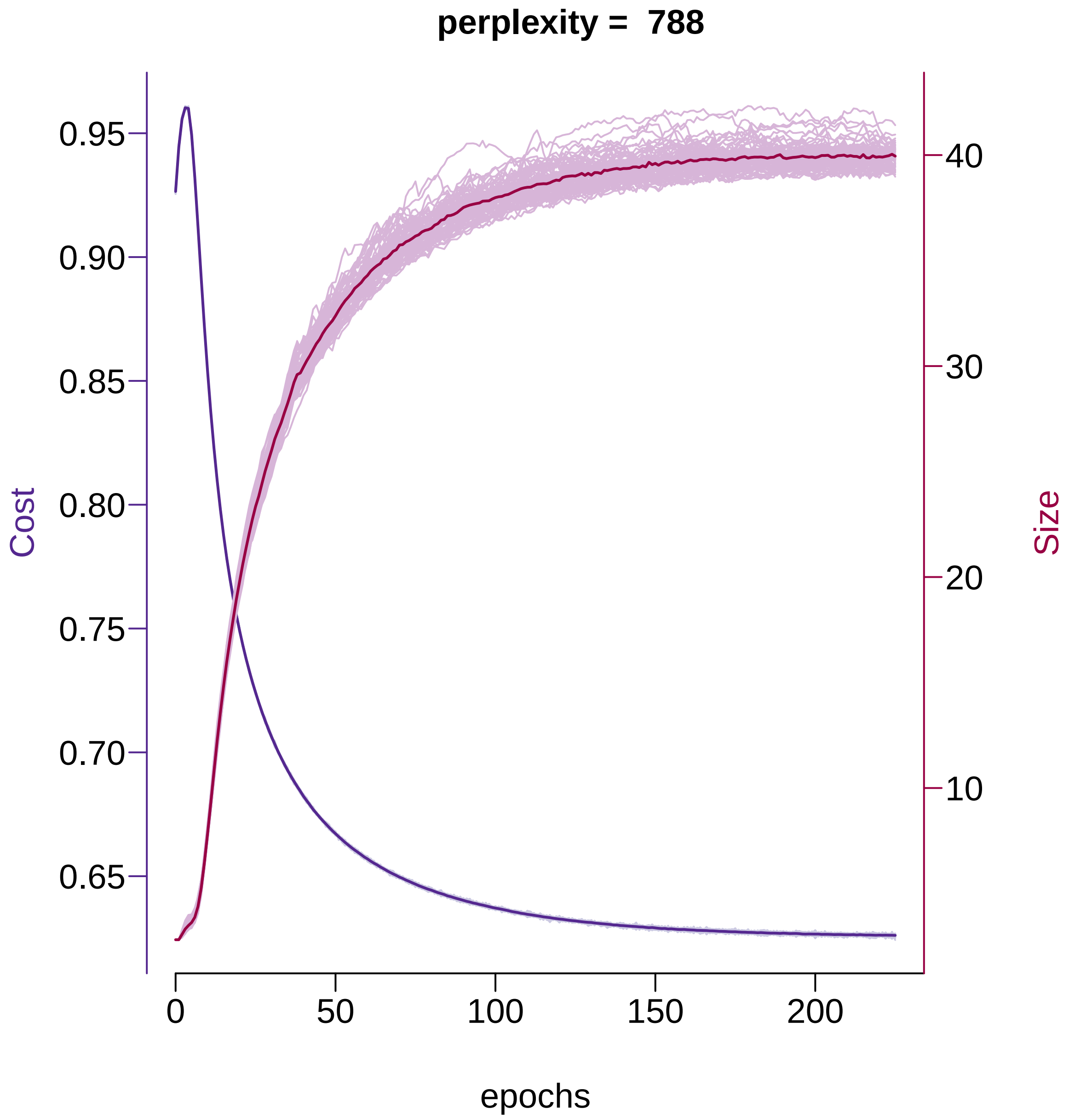
perplexity = 788
Cost
0.95
0.90
0.85
0.80
0.75
0.70
0.65
Size
40
30
20
10
0
50
100
150
200
epochs

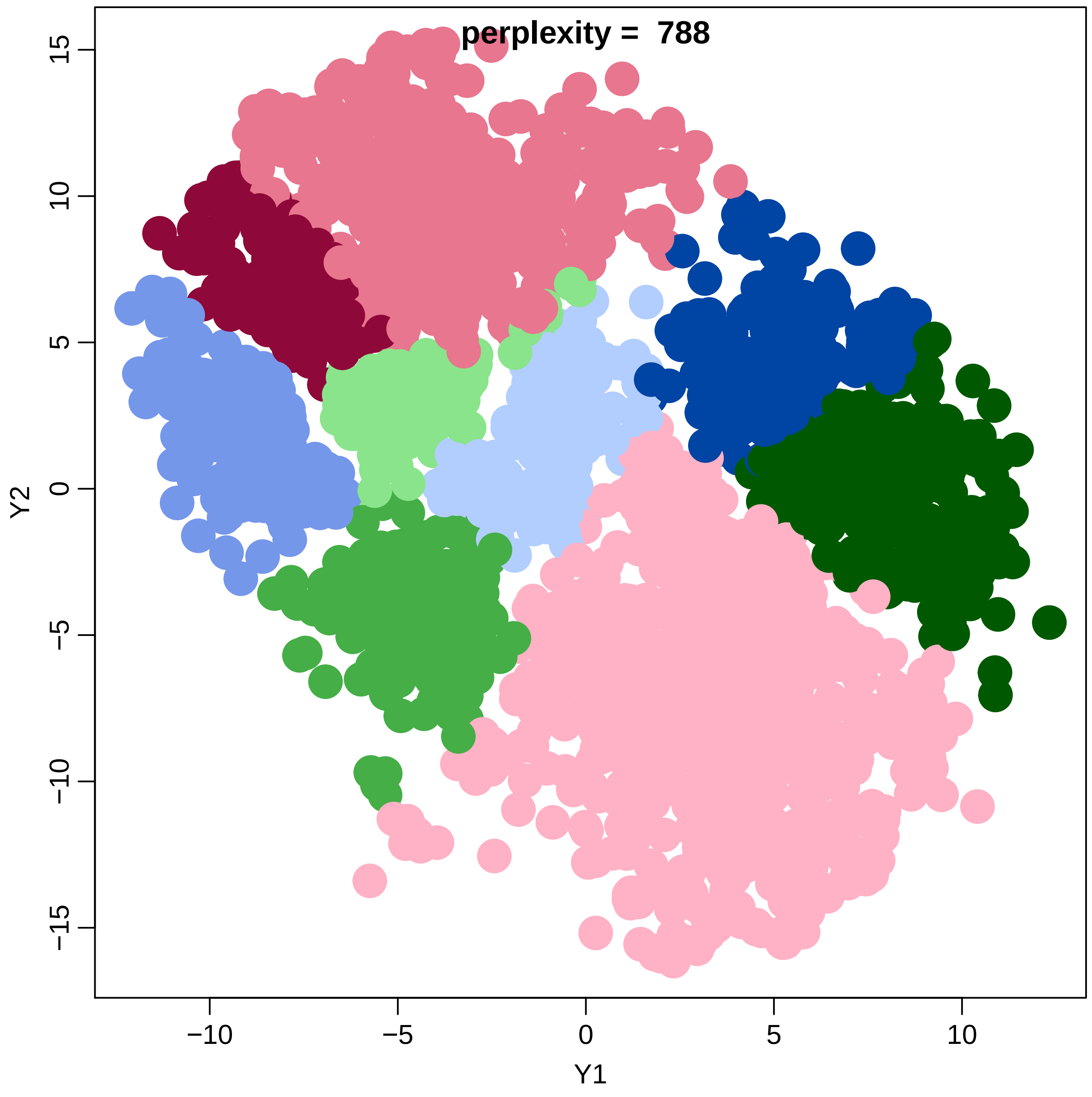

perplexity = 788
Y2
Y1
15
10
5
0
−5
−10
−15
−10
−5
0
5
10

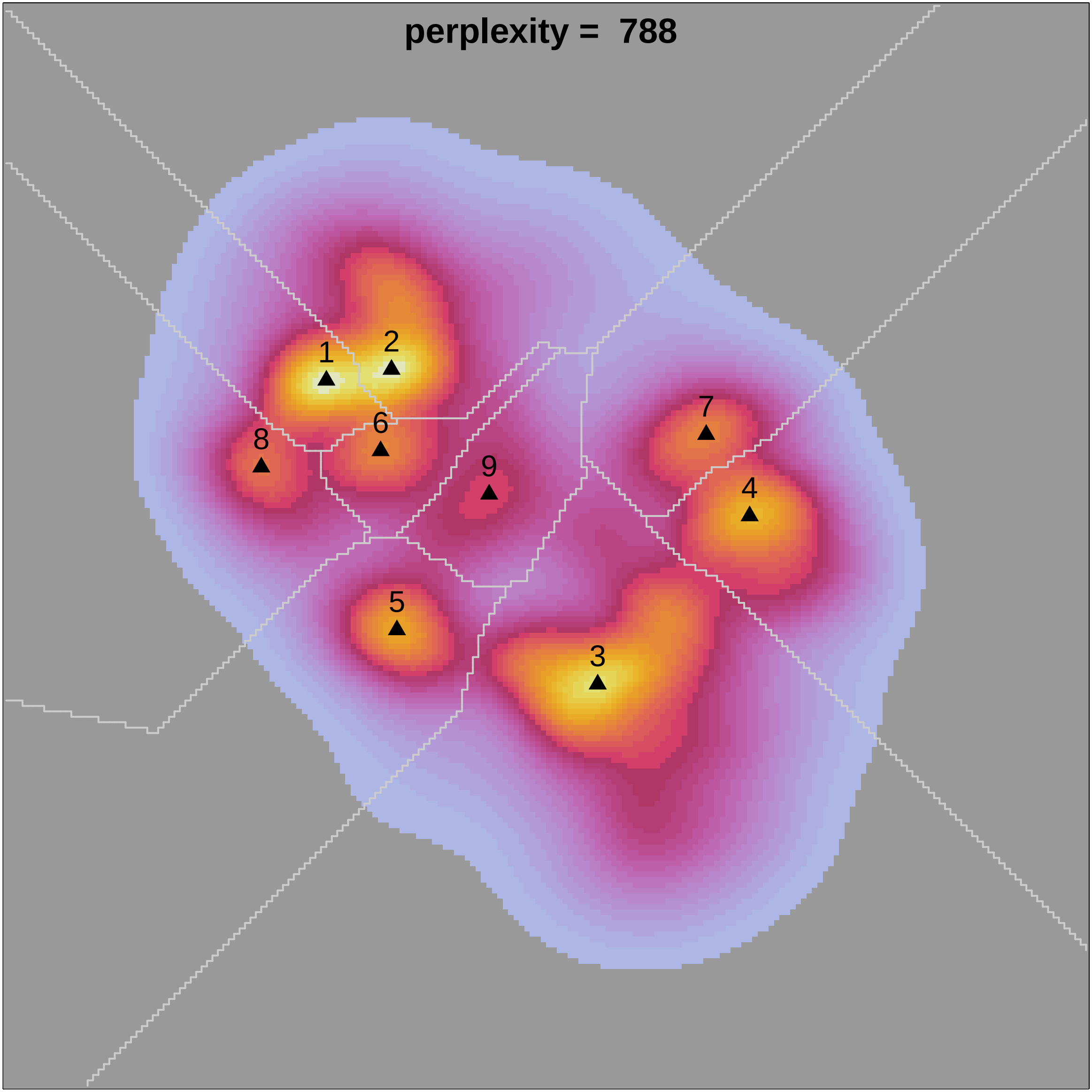
perplexity = 788
1
2
6
8
9
7
4
5
3

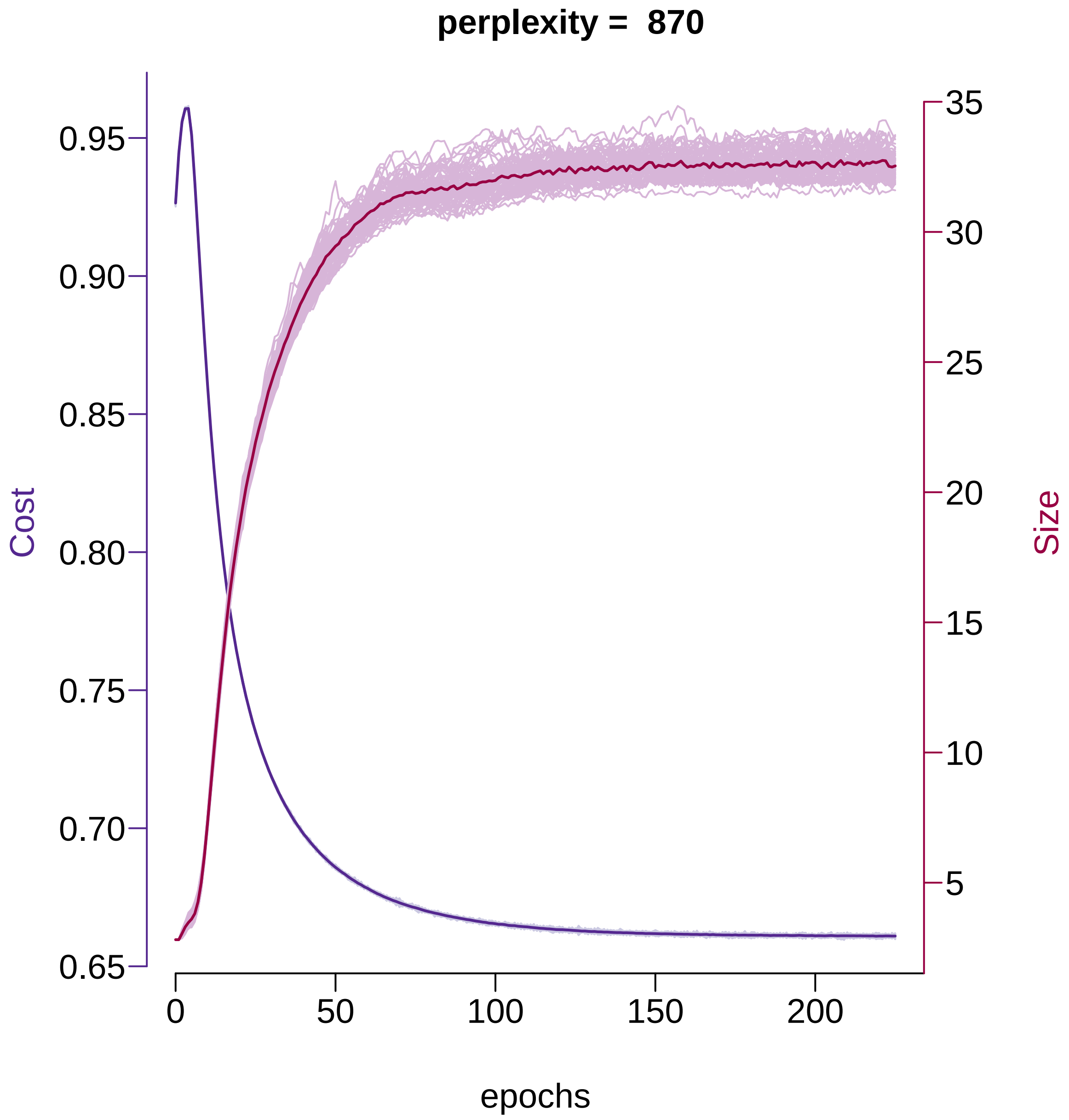
perplexity = 870
Cost
Size
epochs
0.95
0.90
0.85
0.80
0.75
0.70
0.65
35
30
25
20
15
10
5
0
50
100
150
200

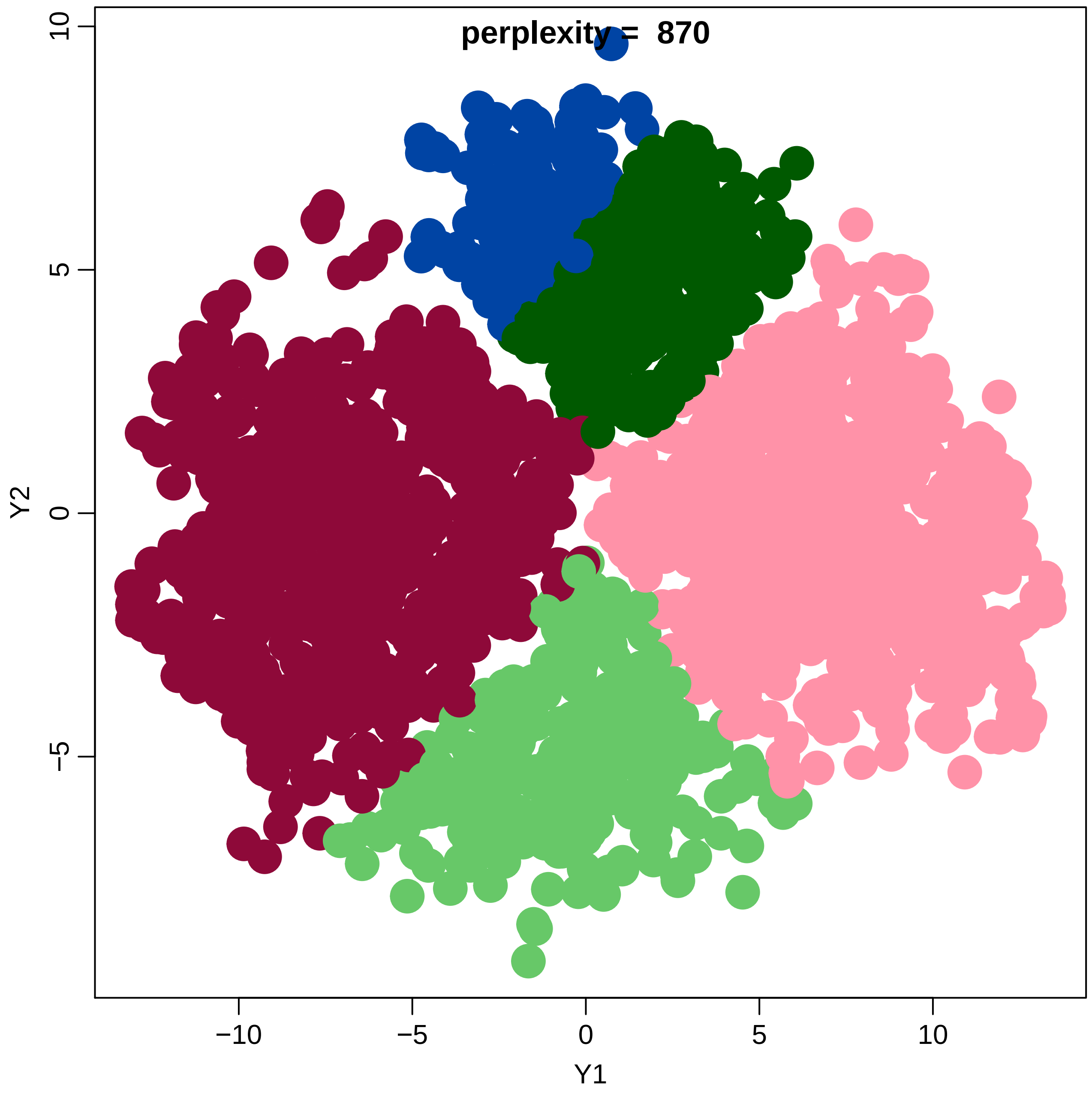
perplexity = 870
Y1
Y2
−10
−5
0
5
10
−5
0
5
10

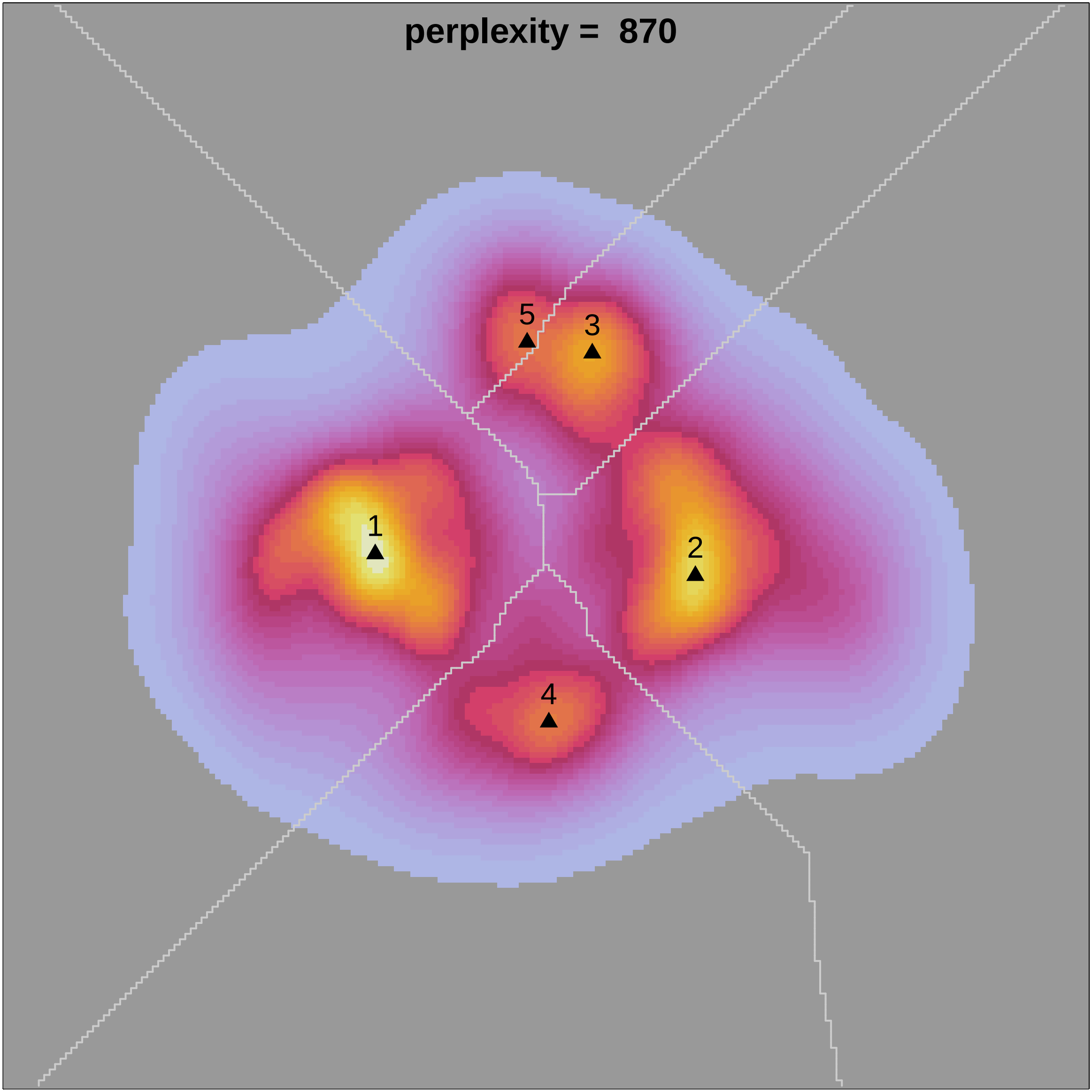
perplexity = 870
1
2
3
4
5

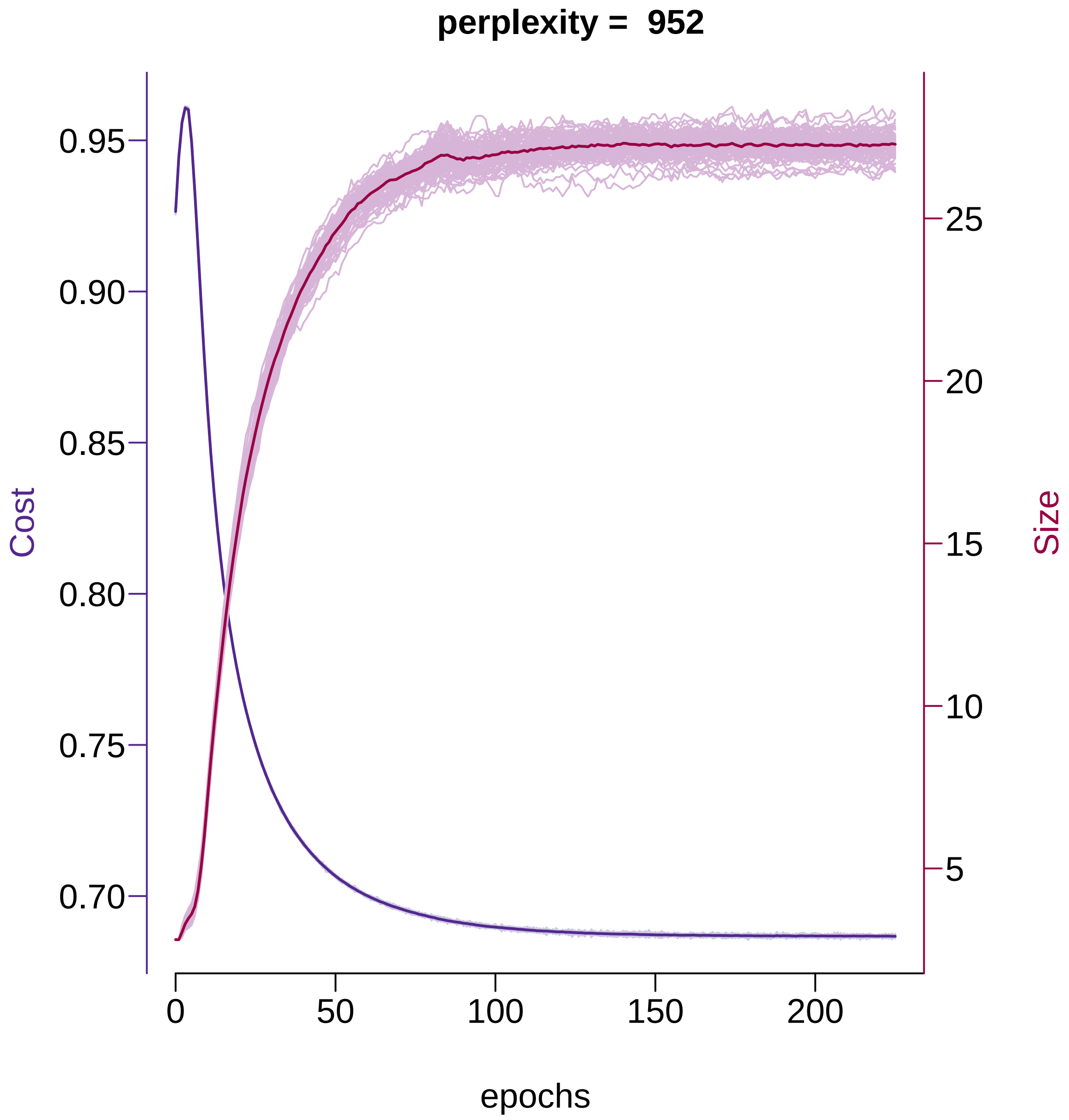
perplexity = 952
Cost
0.95
0.90
0.85
0.80
0.75
0.70
Size
25
20
15
10
5
0
50
100
150
200
epochs

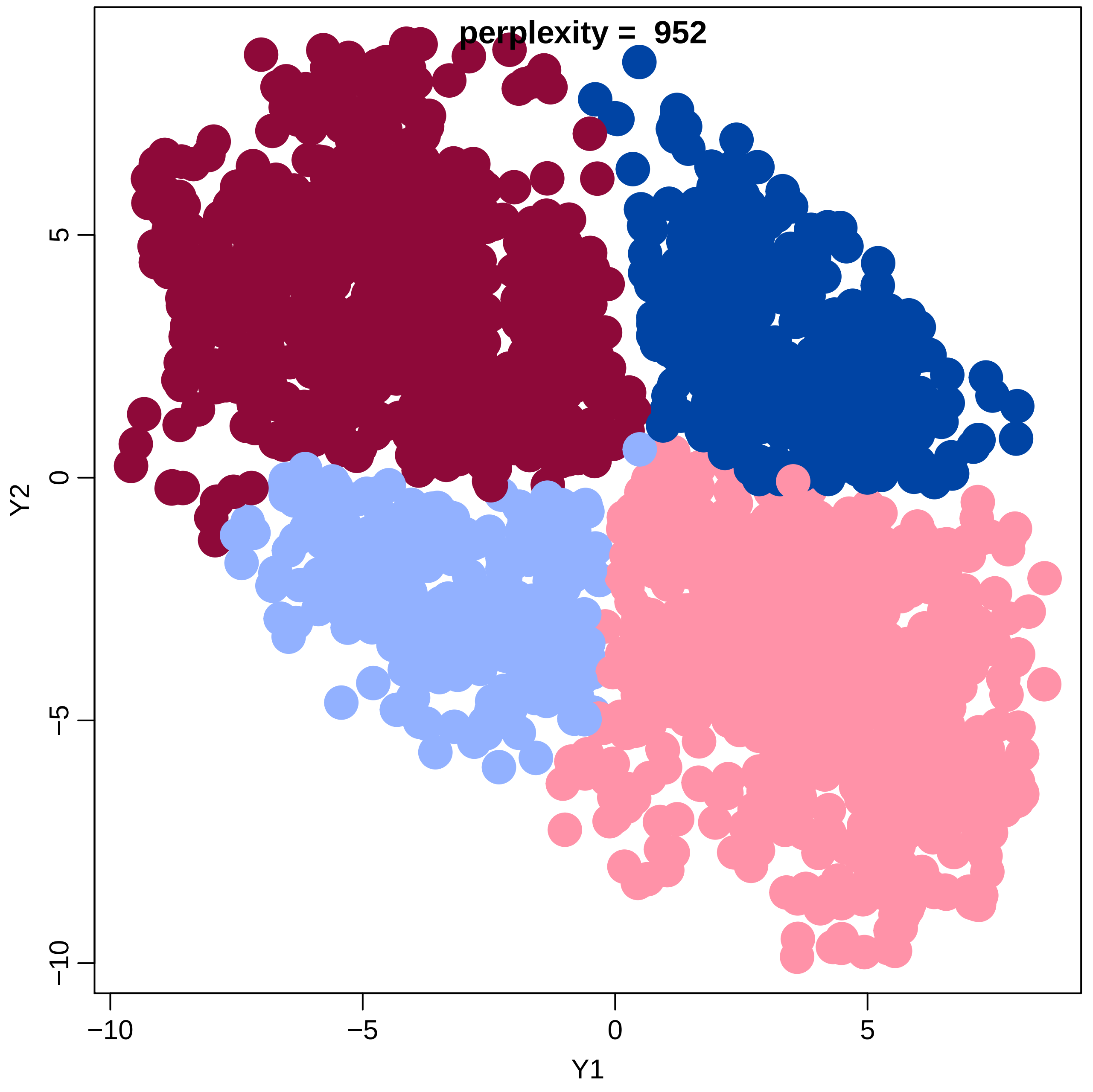
perplexity = 952
Y1
Y2
−10
−5
0
5
−10
−5
0
5

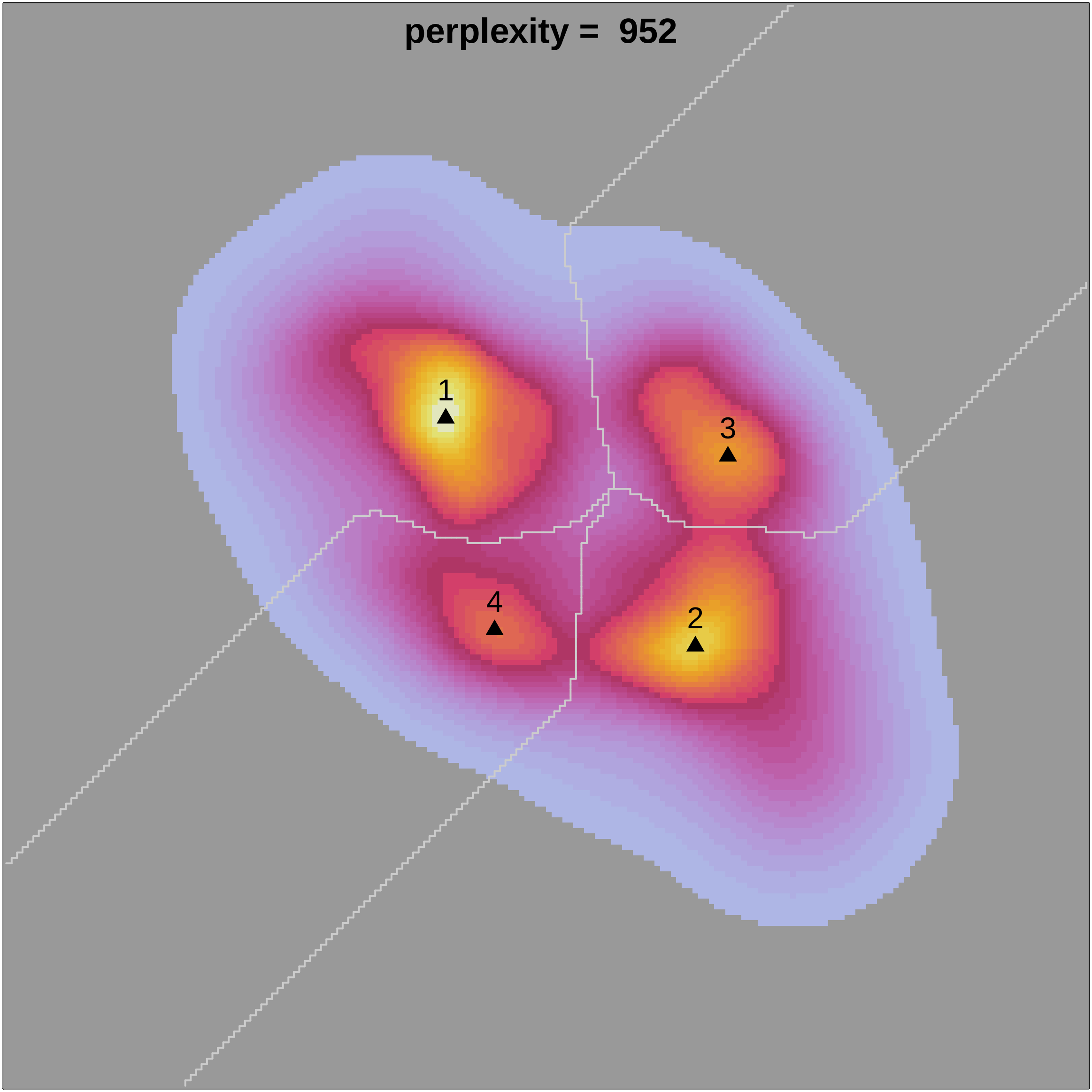

perplexity = 952
1
2
3
4

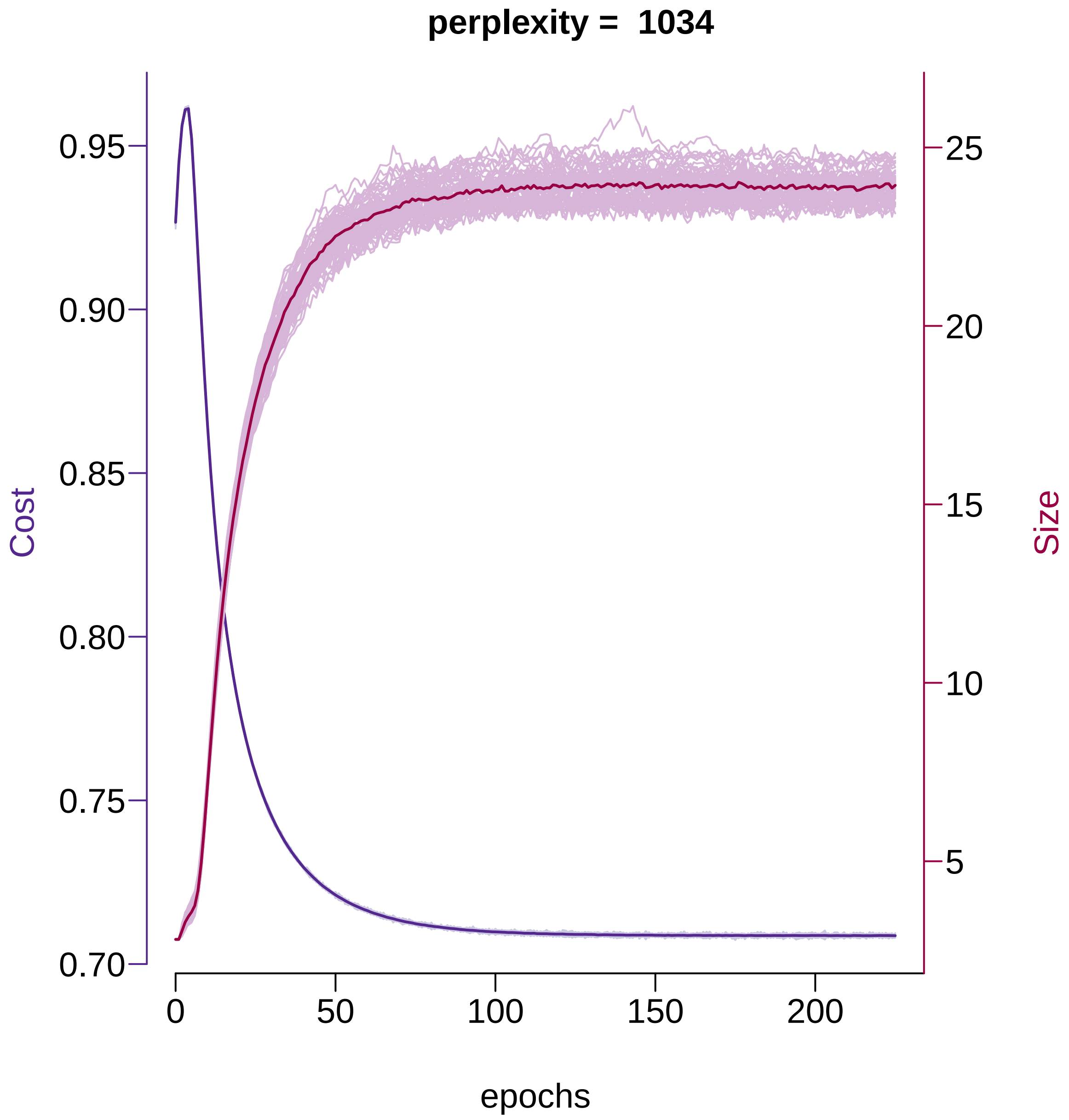
perplexity = 1034
Cost
0.95
0.90
0.85
0.80
0.75
0.70
Size
25
20
15
10
5
0
50
100
150
200
epochs

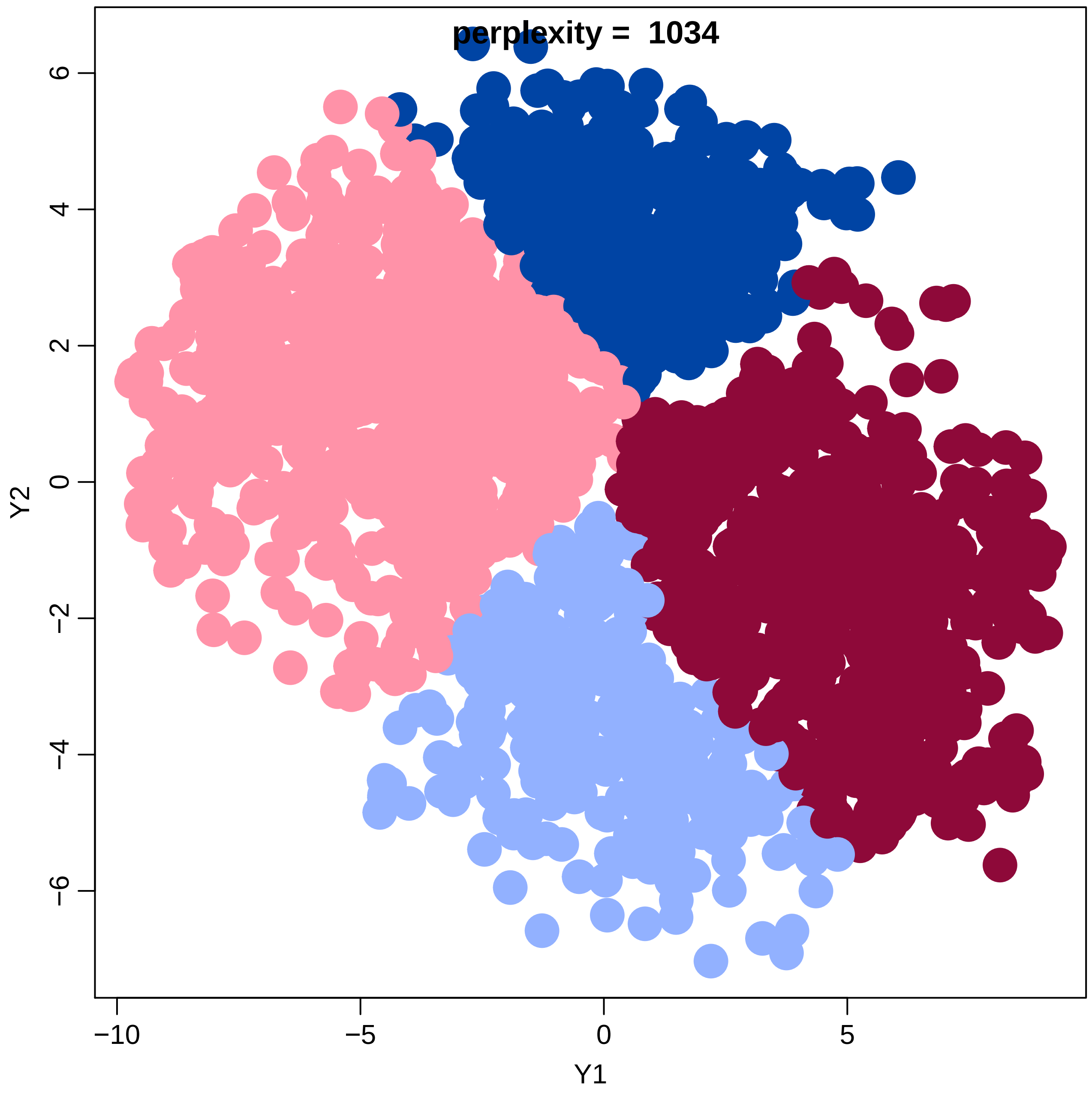
perplexity = 1034
Y2
Y1
6
4
2
0
−2
−4
−6
−10
−5
0
5

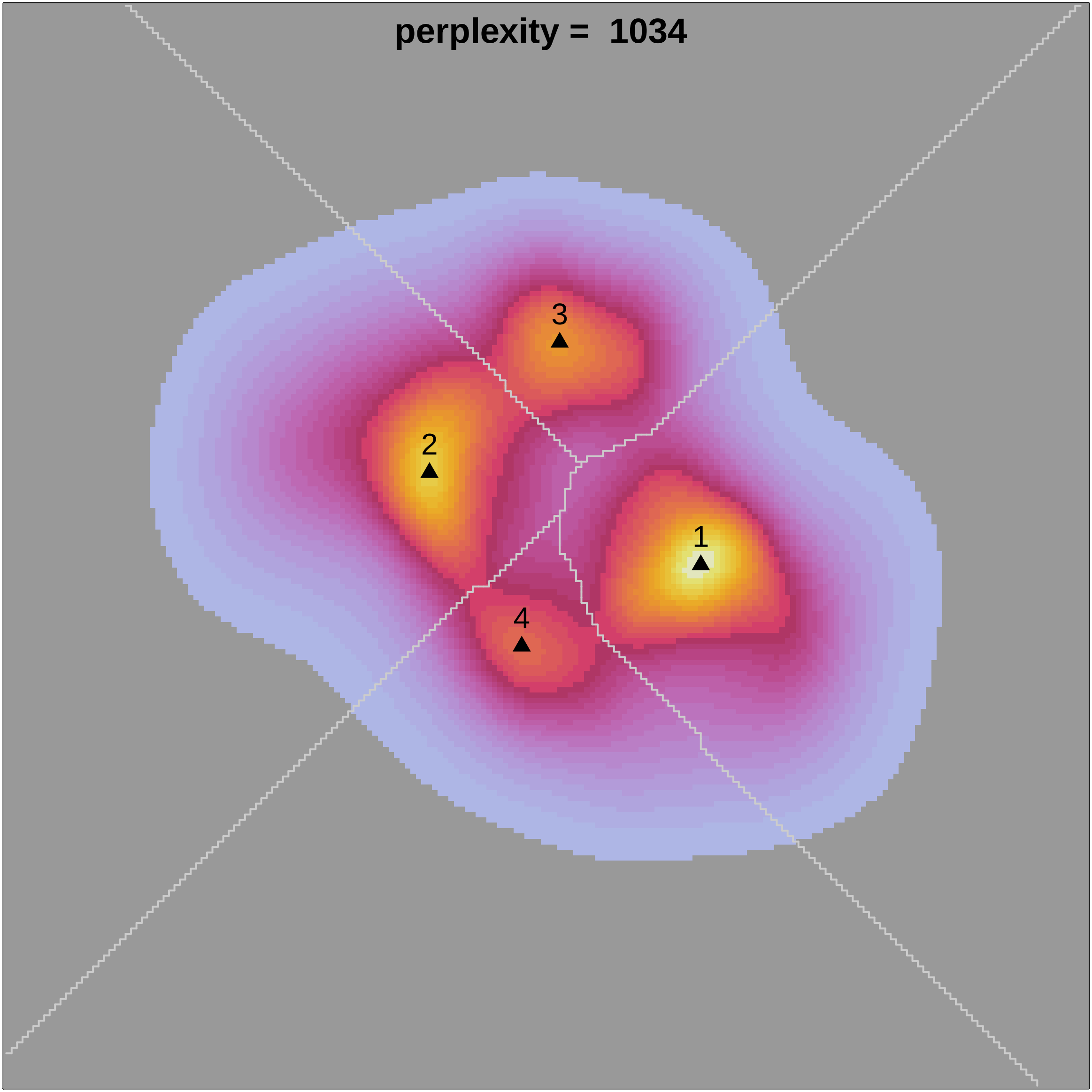
perplexity = 1034
3
2
1
4

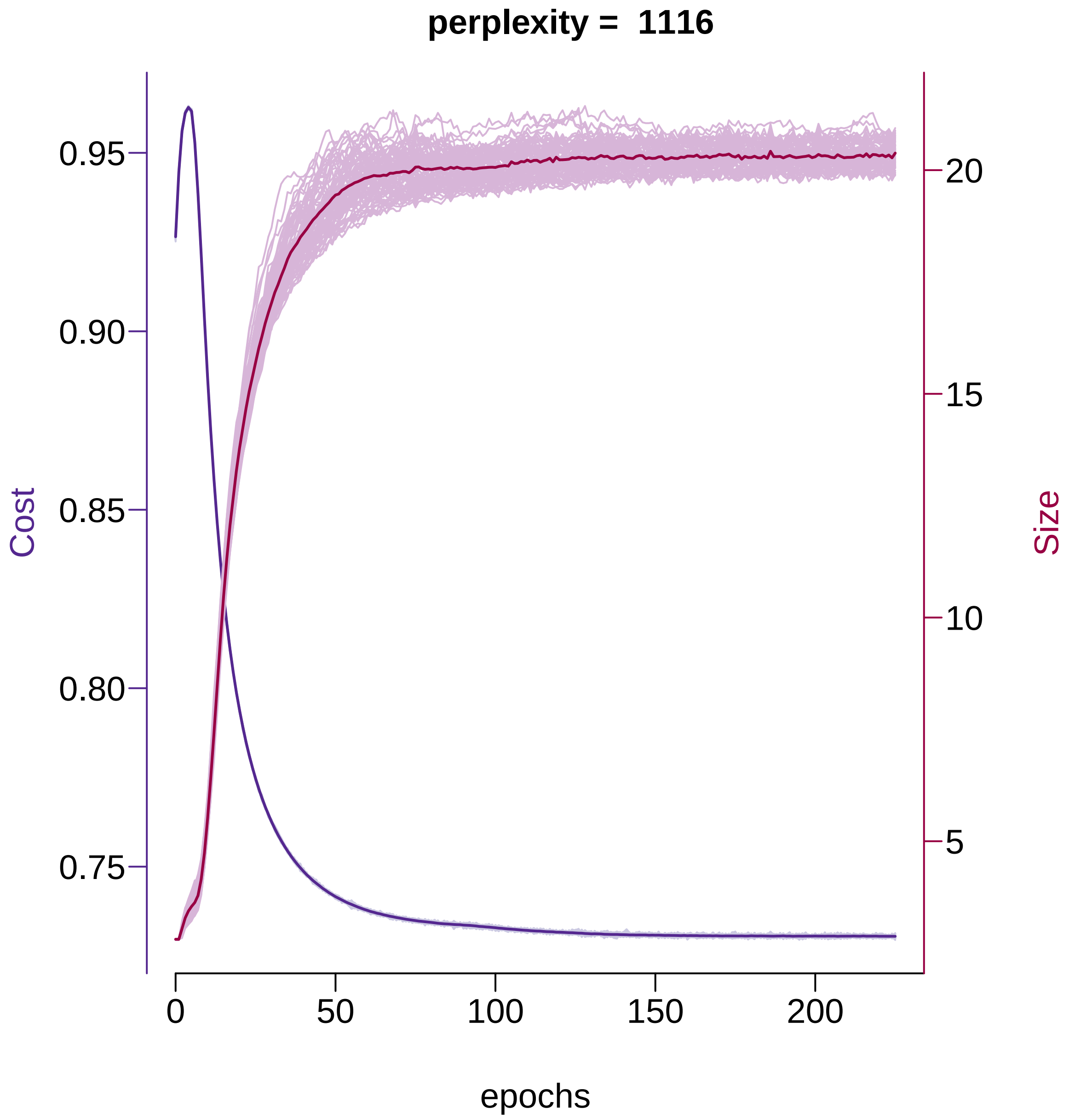

perplexity = 1116
Cost
0.95
0.90
0.85
0.80
0.75
Size
20
15
10
5
0
50
100
150
200
epochs

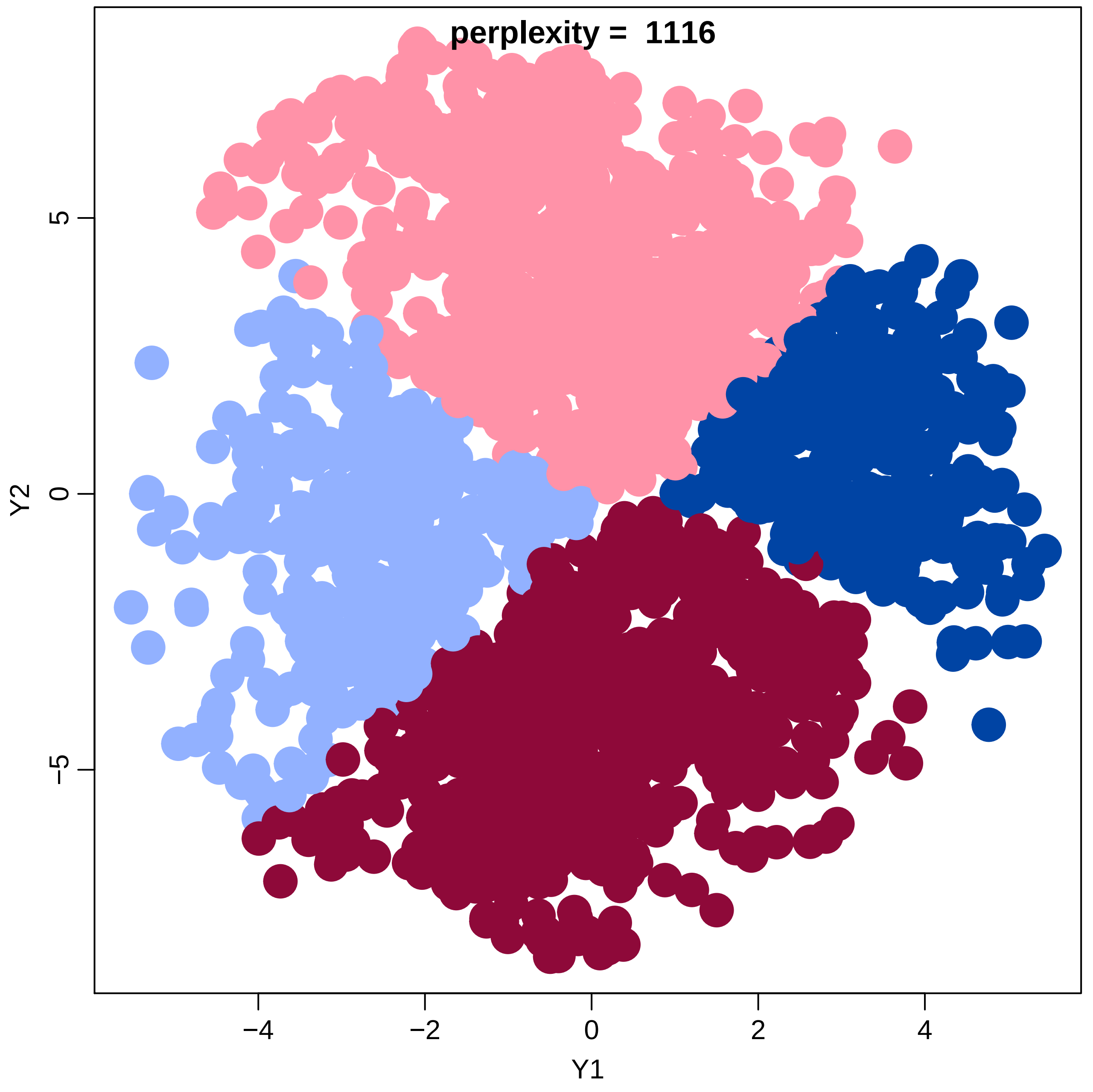
perplexity = 1116
Y2
5
0
−5
−4
−2
0
2
4
Y1

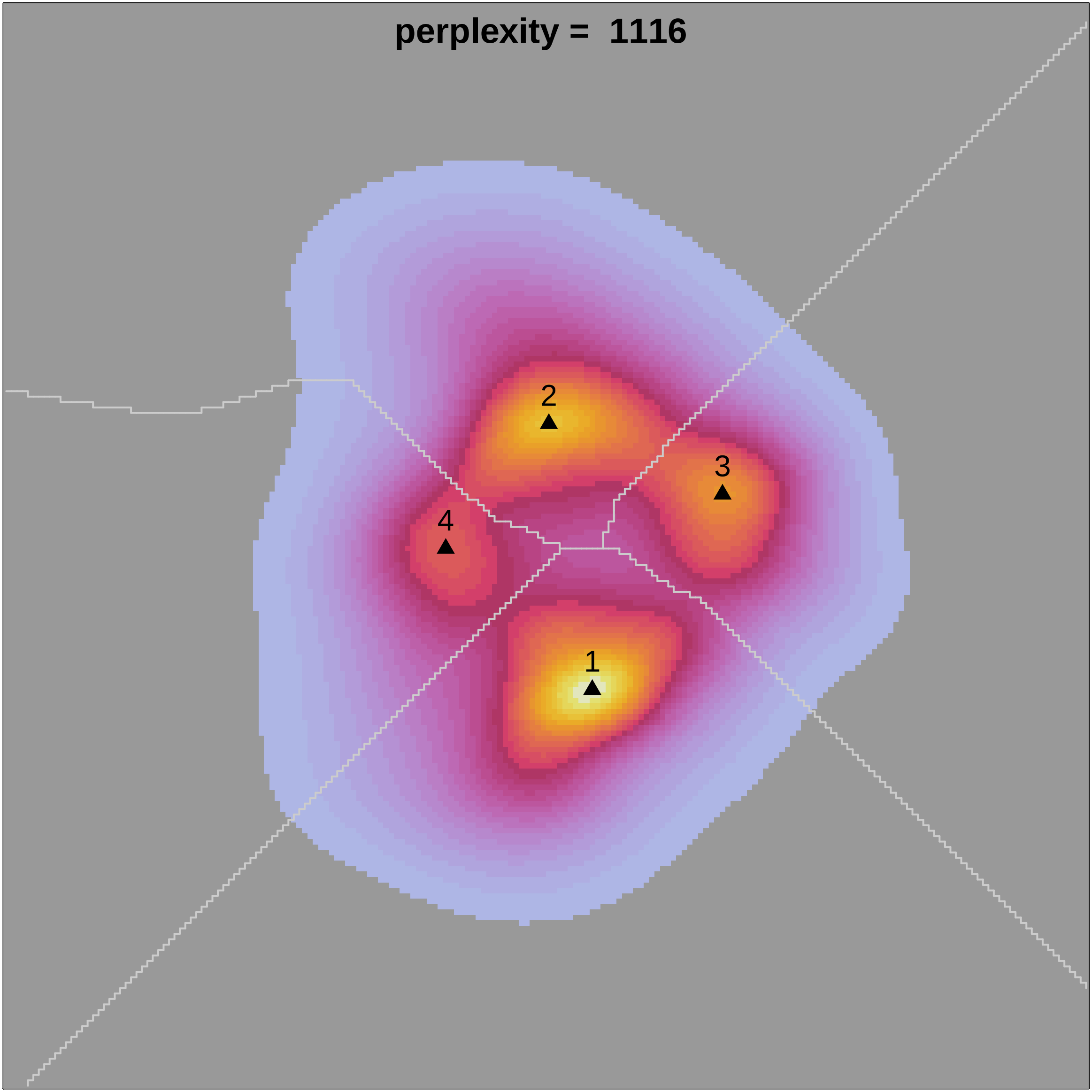
perplexity = 1116
1
2
3
4

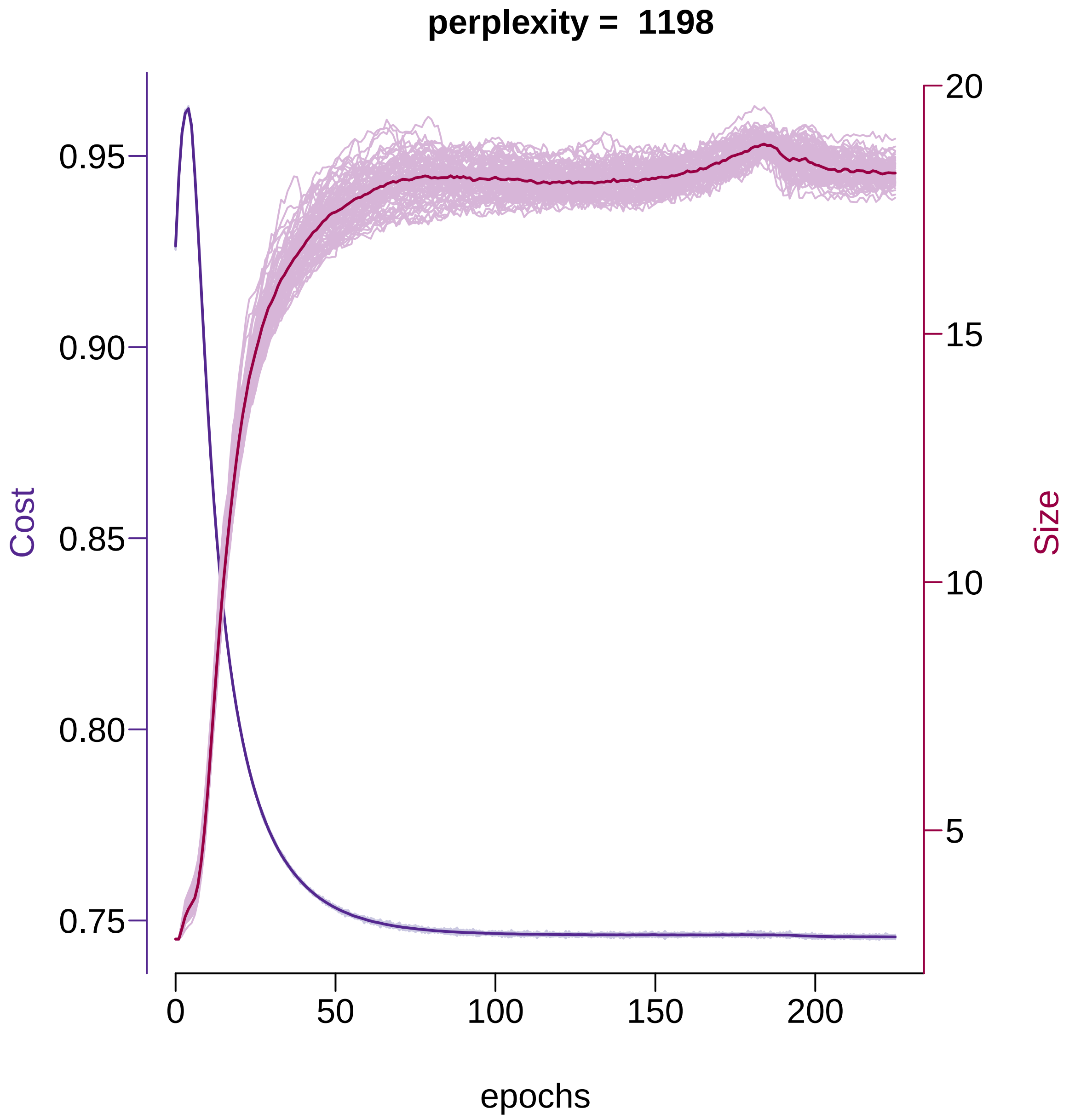
perplexity = 1198
Cost
0.95
0.90
0.85
0.80
0.75
Size
20
15
10
5
0
50
100
150
200
epochs

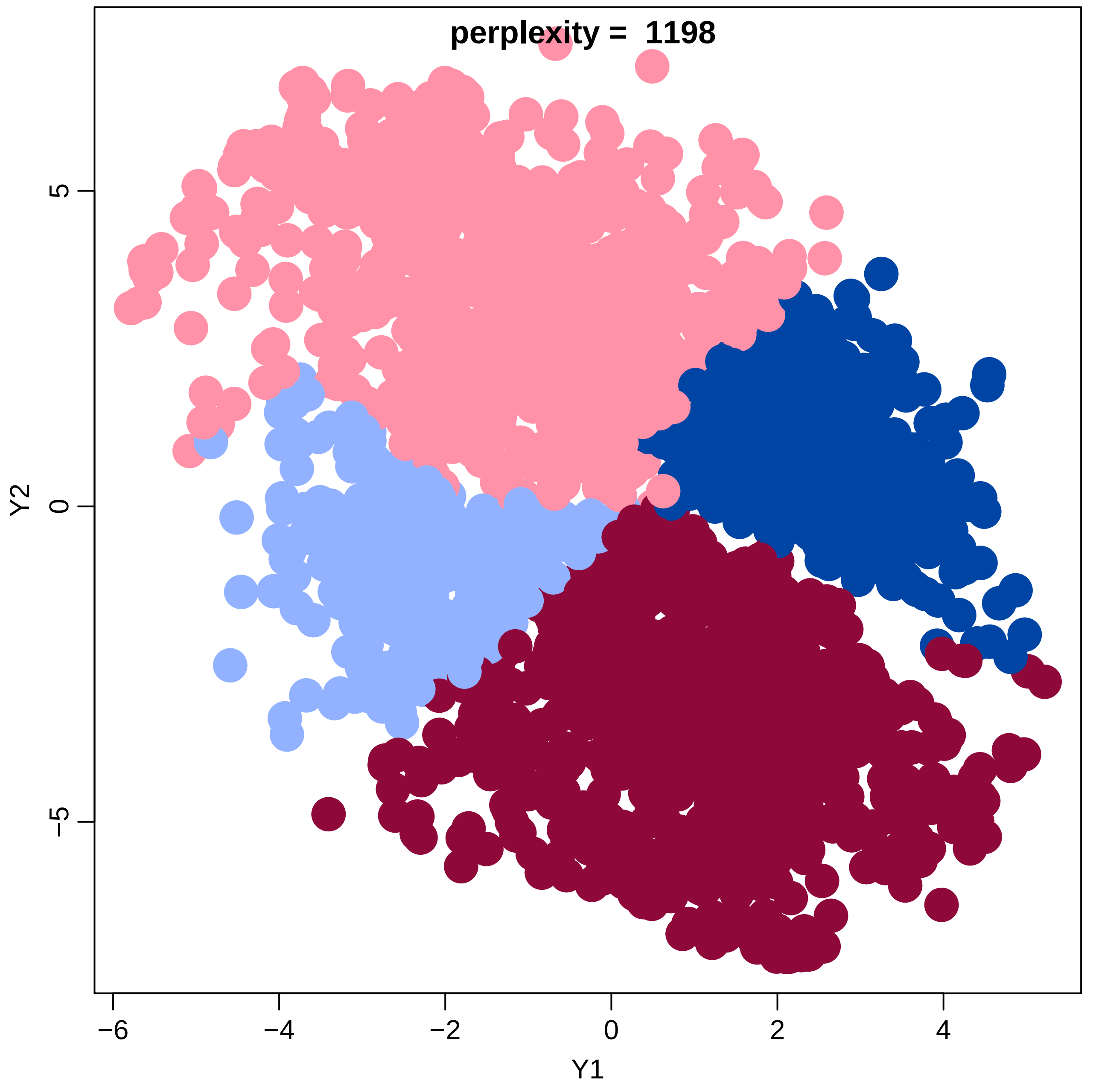

perplexity = 1198
Y2
Y1
5
0
−5
−6
−4
−2
0
2
4

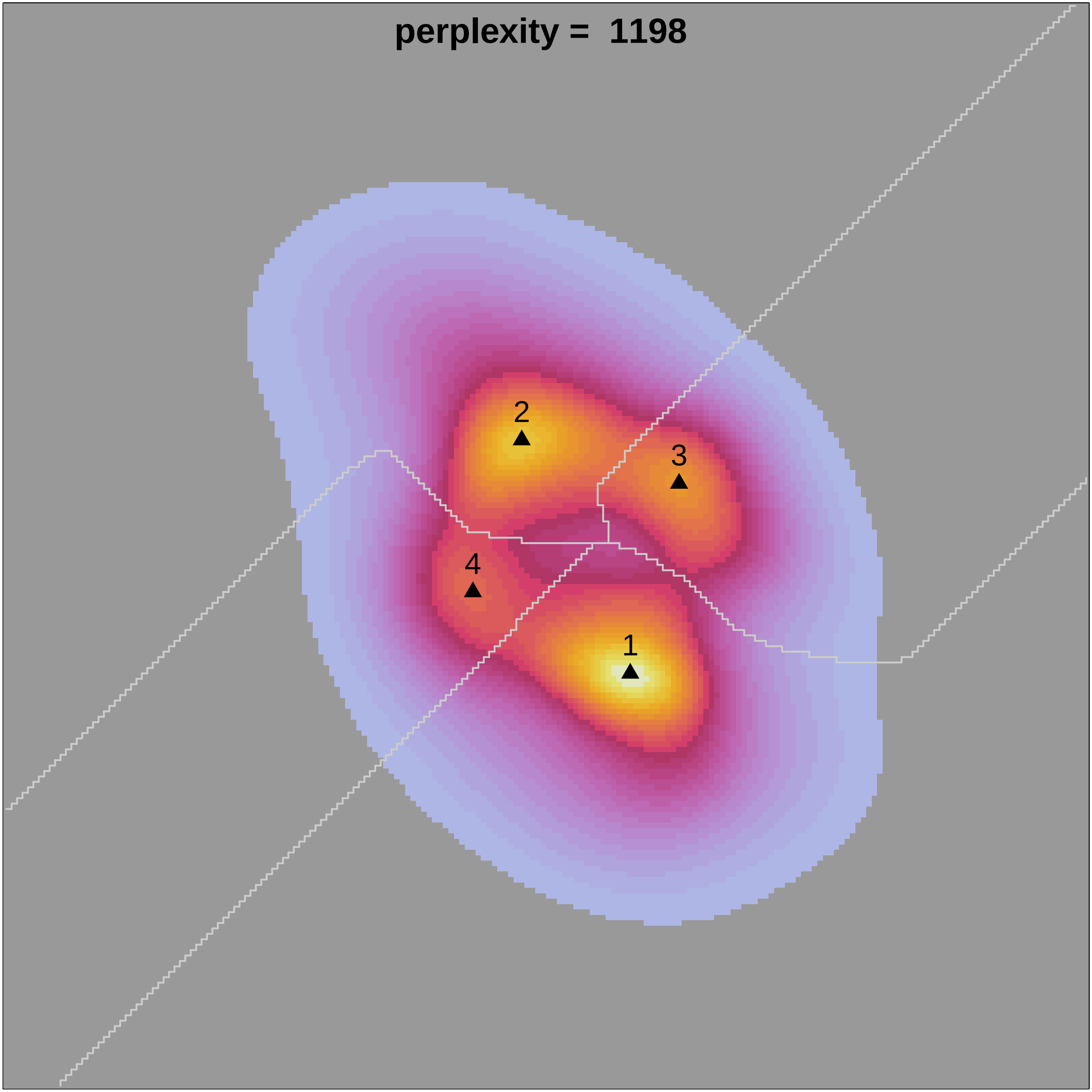
perplexity = 1198
2
3
4
1

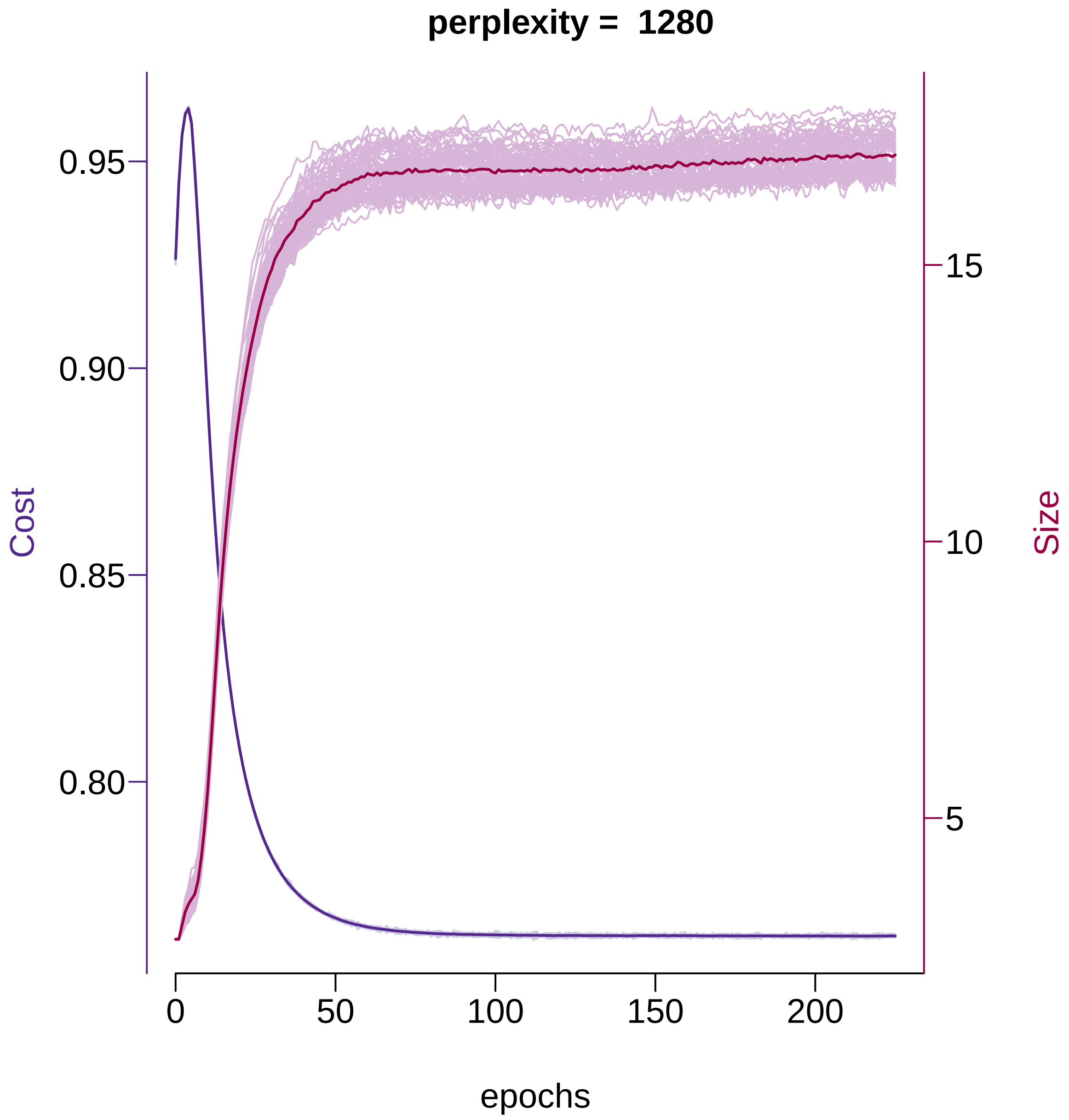
perplexity = 1280
Cost
0.95
0.90
0.85
0.80
Size
15
10
5
0
50
100
150
200
epochs

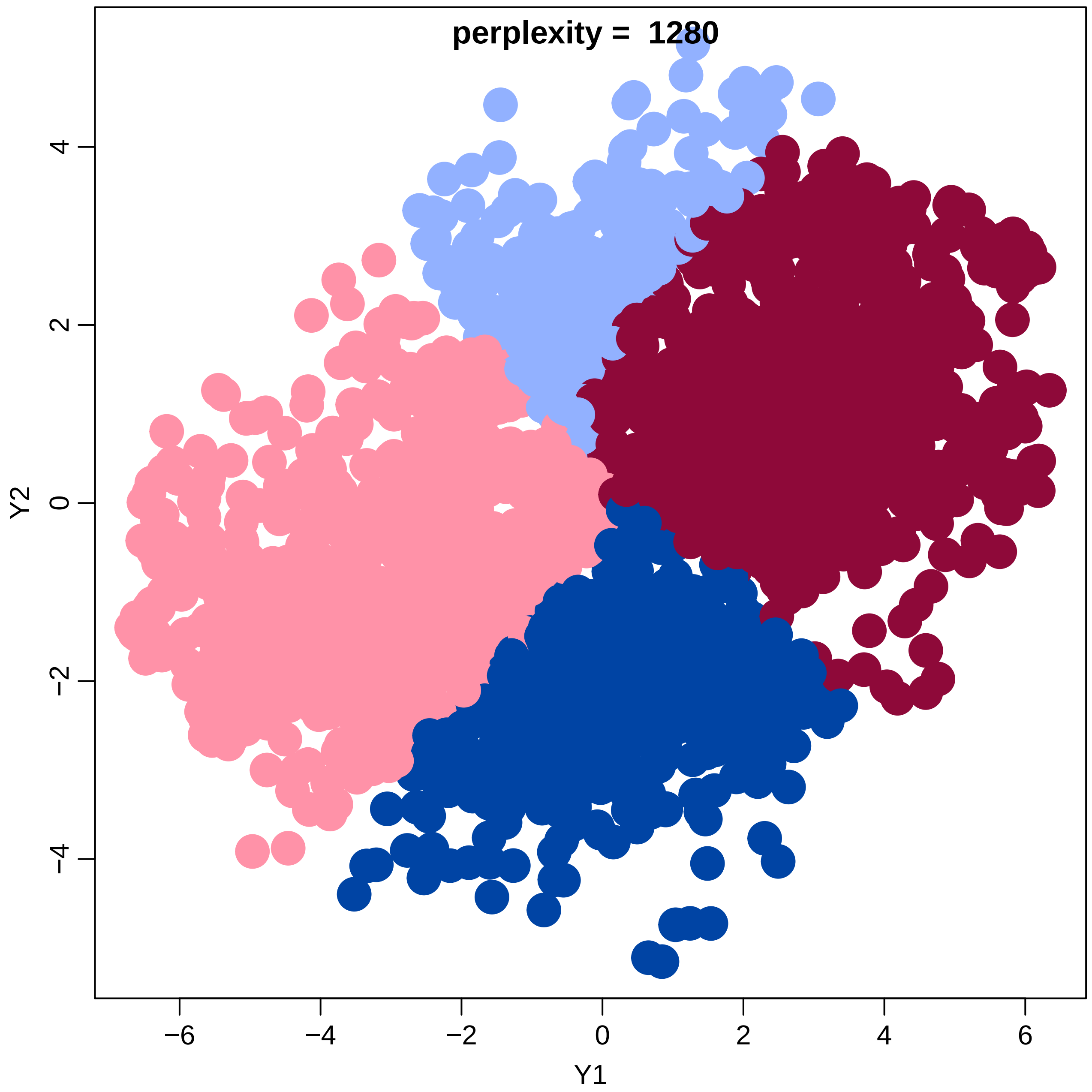
perplexity = 1280
Y2
4
2
0
−2
−4
Y1
−6
−4
−2
0
2
4
6

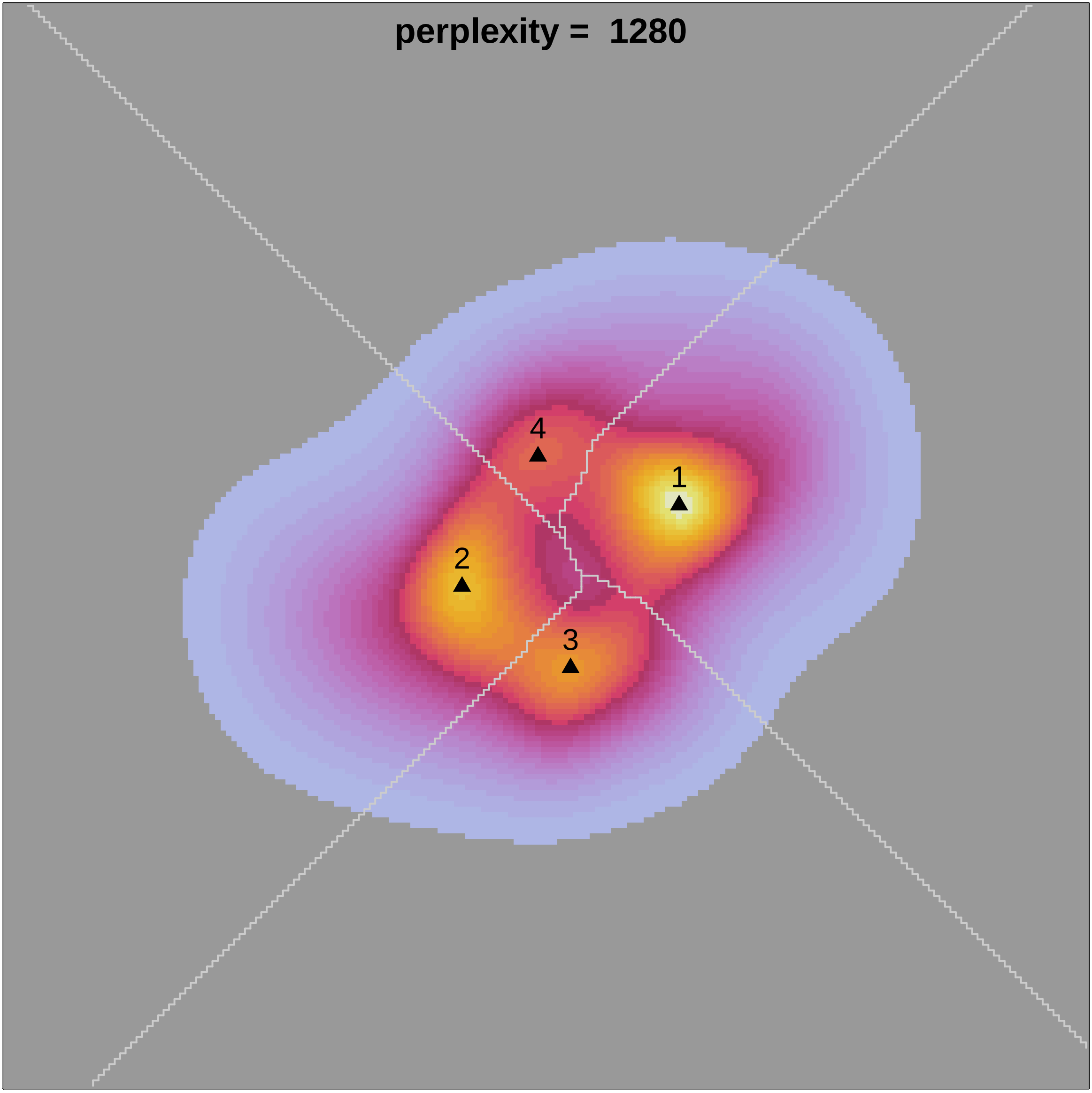
perplexity = 1280
4
1
2
3

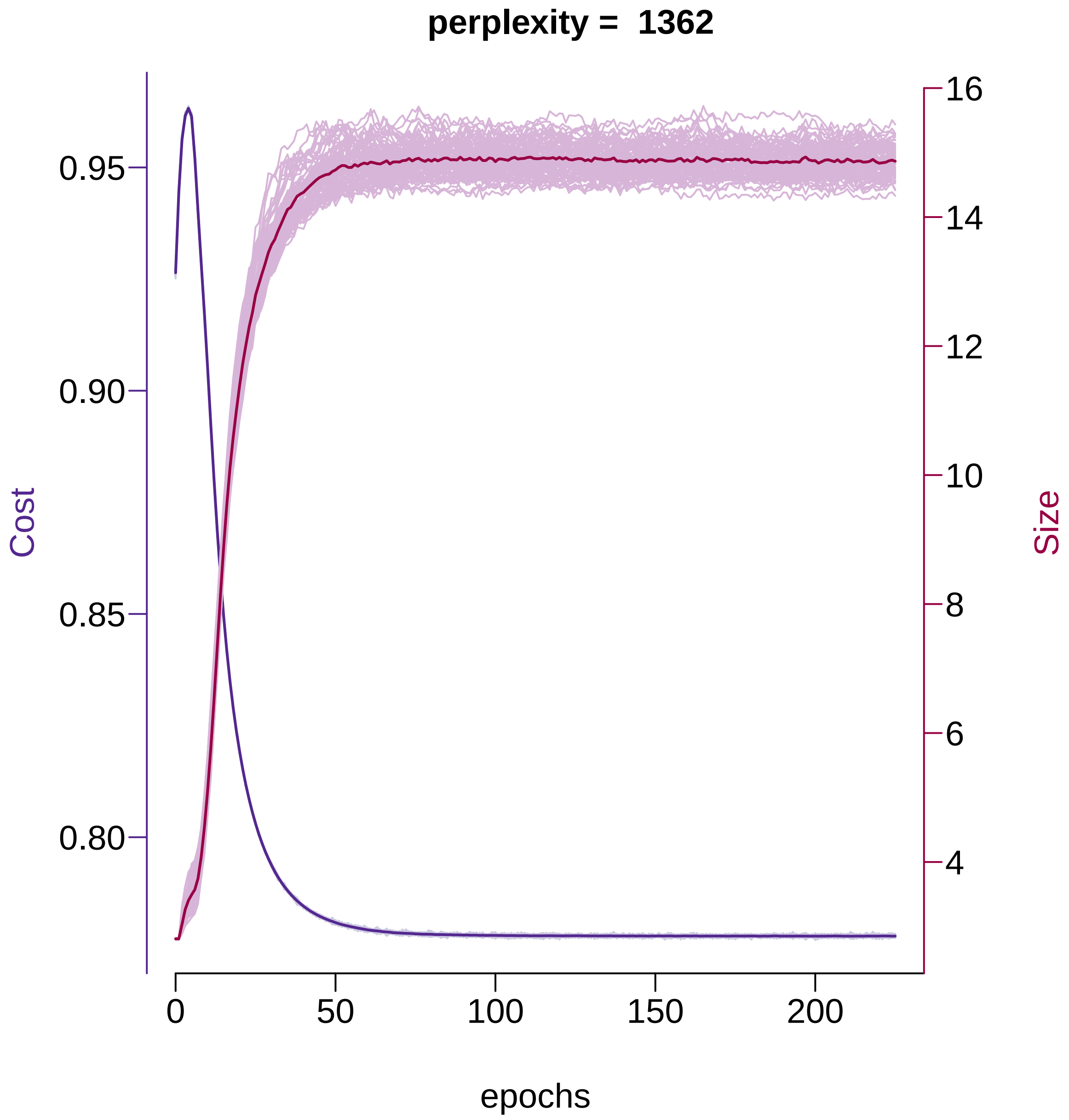
perplexity = 1362
Cost
0.95
0.90
0.85
0.80
Size
16
14
12
10
8
6
4
0
50
100
150
200
epochs

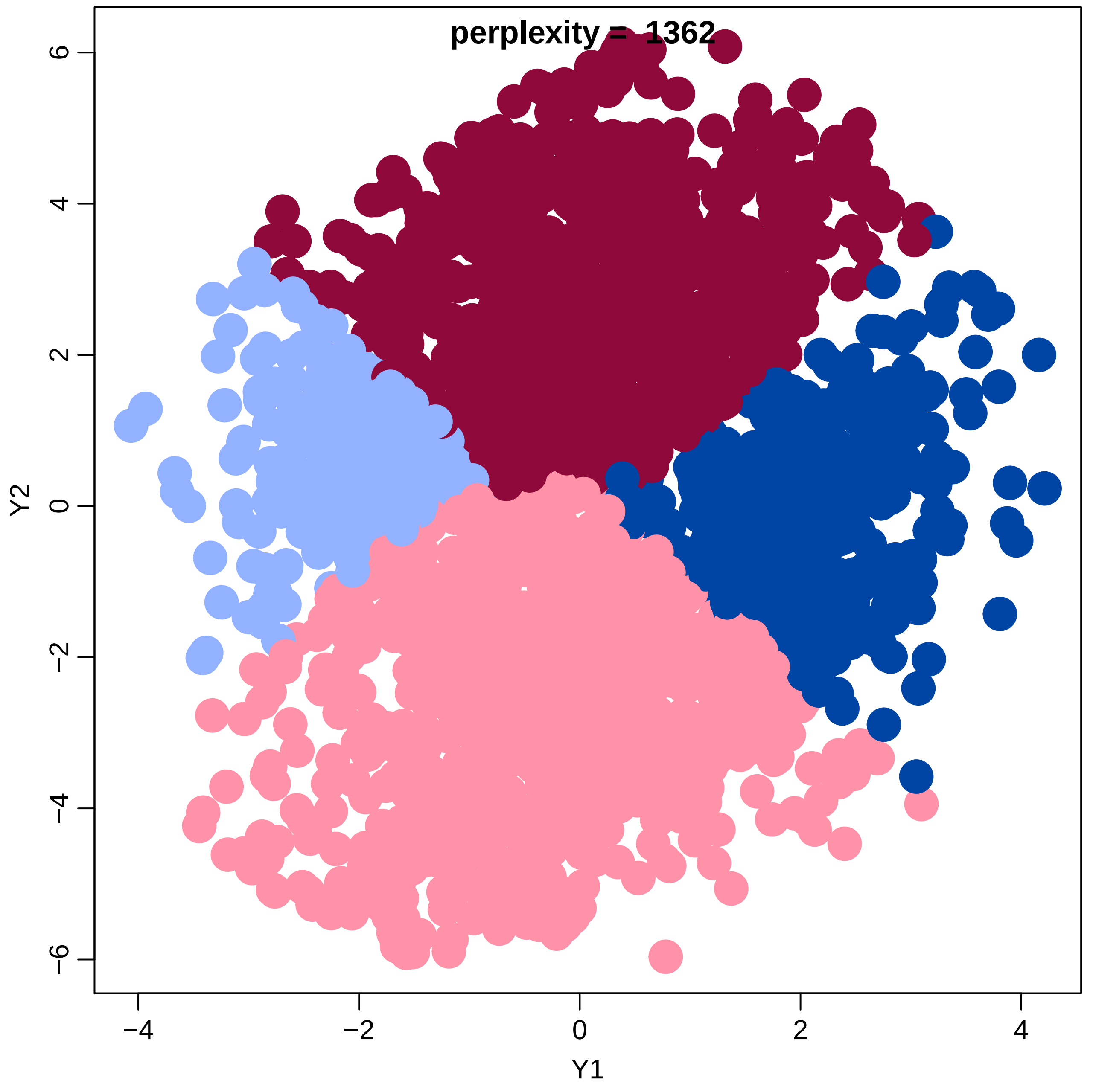
perplexity = 1362
Y2
Y1
6
4
2
0
−2
−4
−6
−4
−2
0
2
4

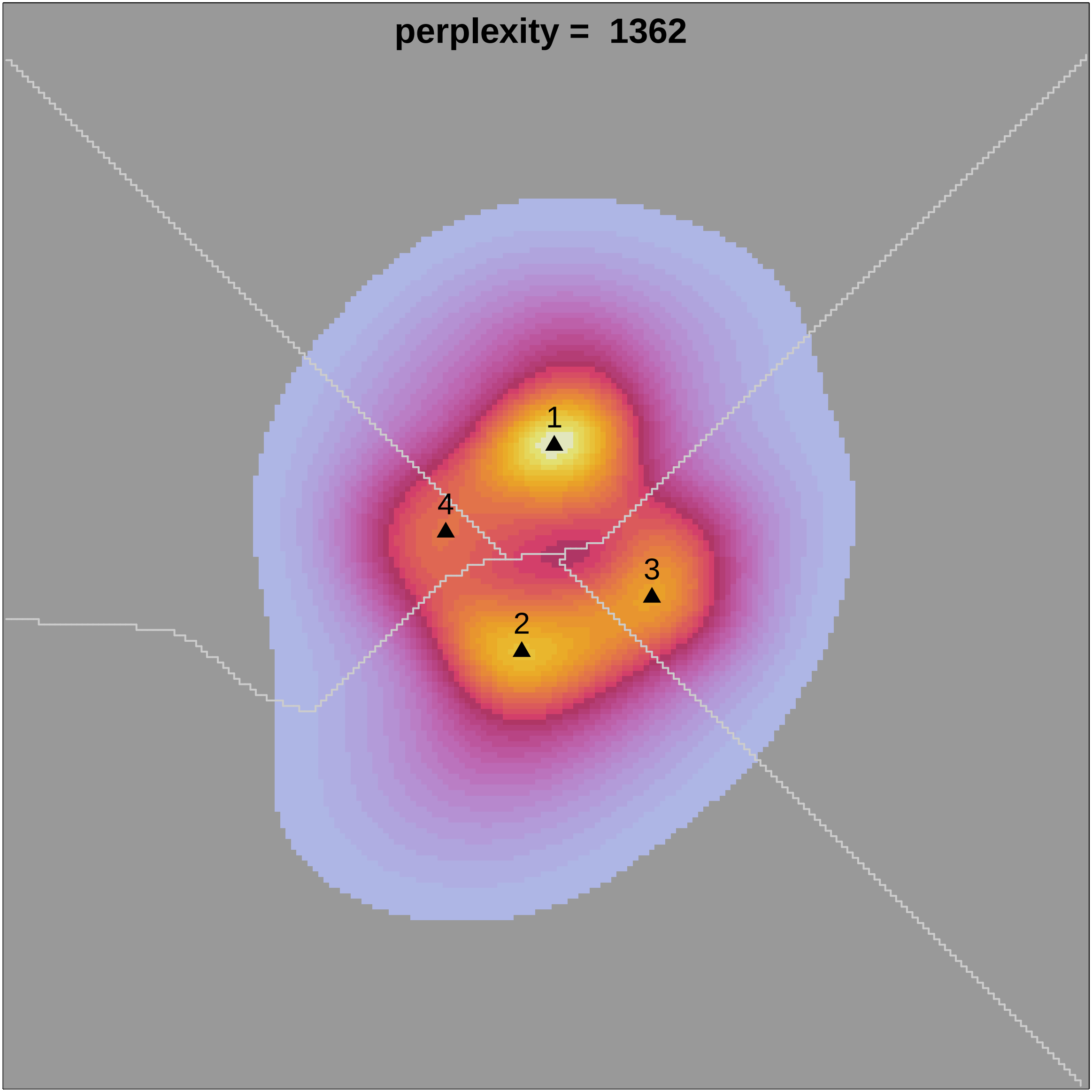

perplexity = 1362
1
4
3
2

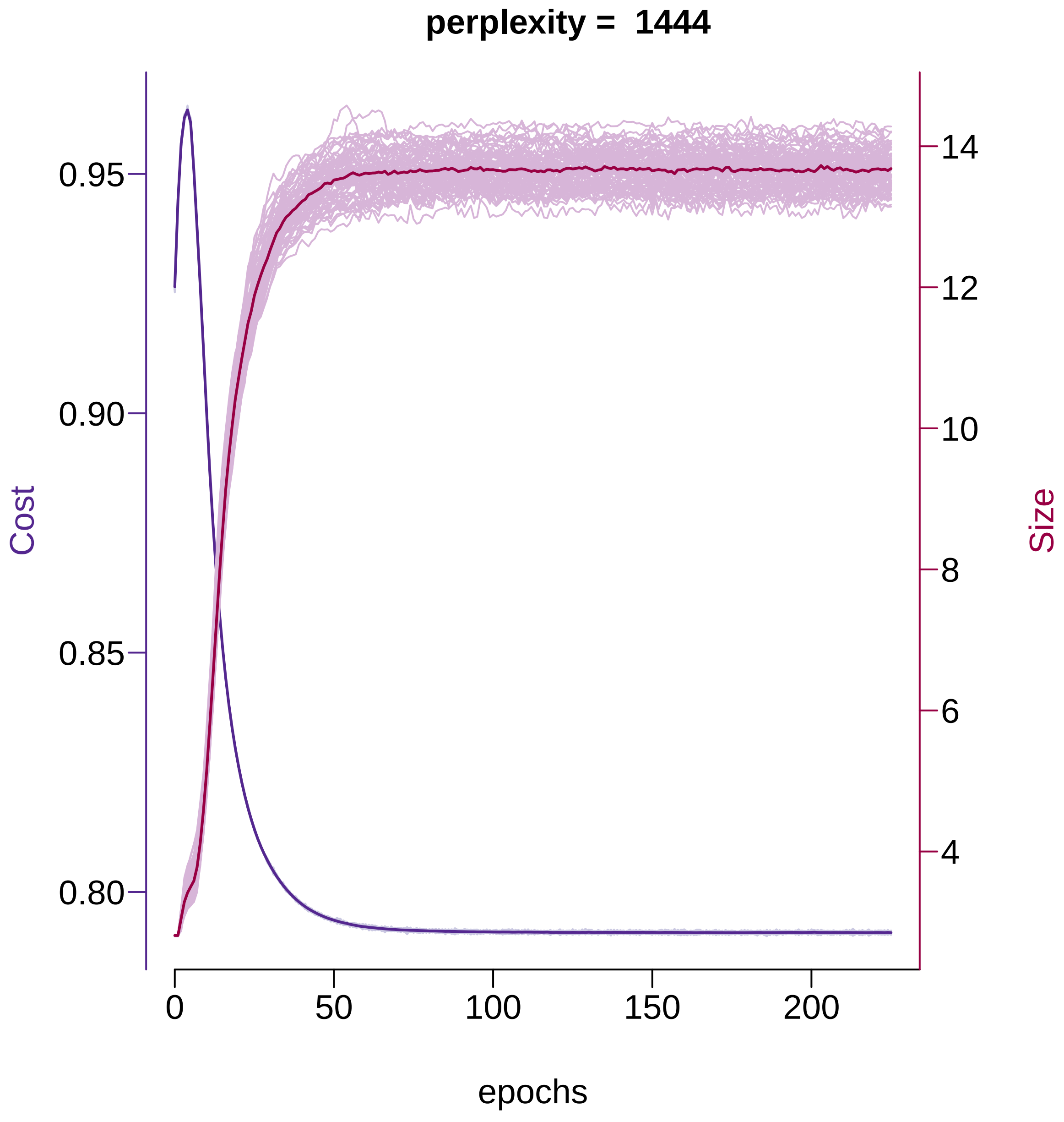
perplexity = 1444
Cost
Size
epochs
0.95
0.90
0.85
0.80
14
12
10
8
6
4
0
50
100
150
200

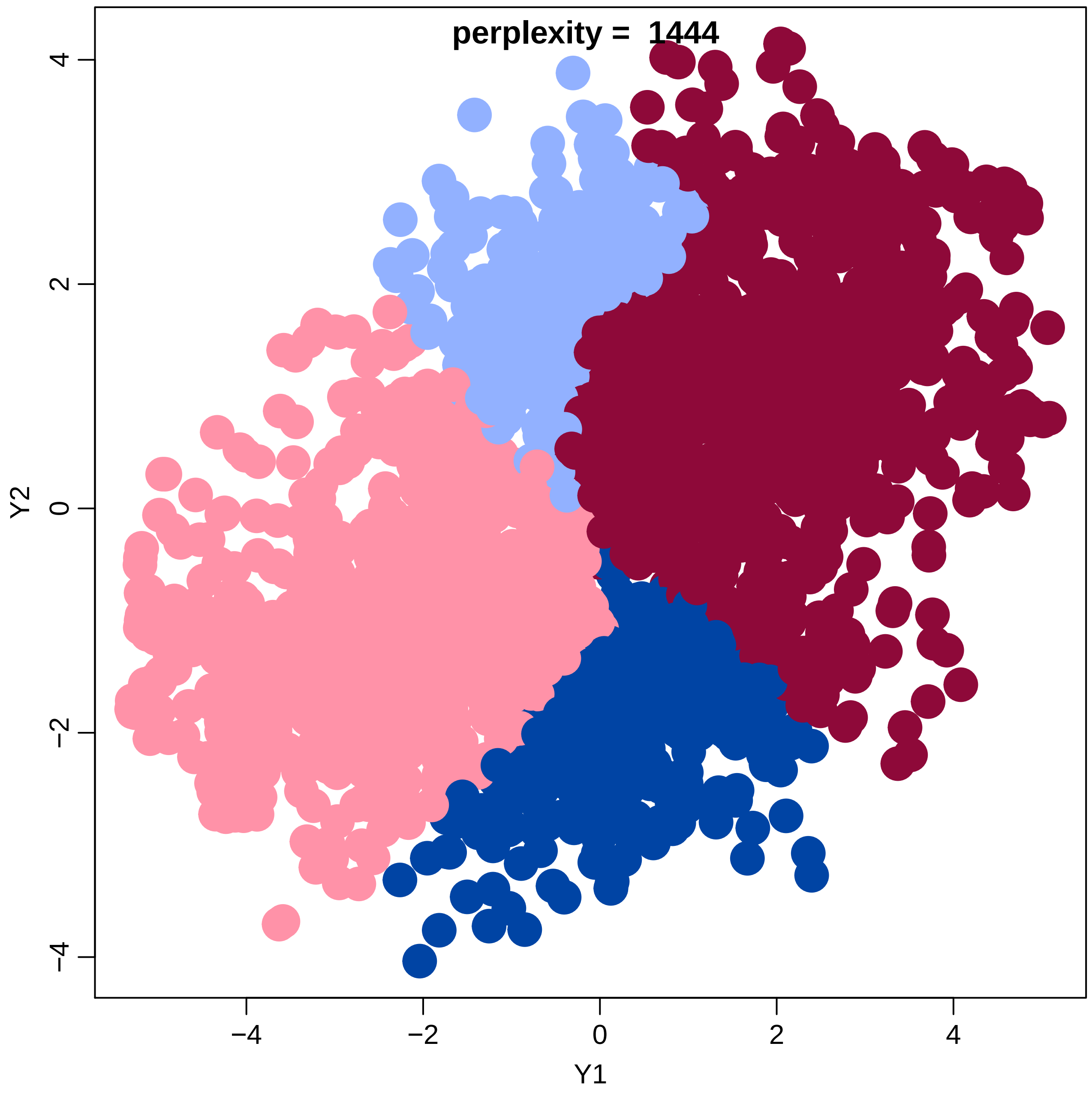
perplexity = 1444
Y2
Y1
4
2
0
−2
−4
−4
−2
0
2
4

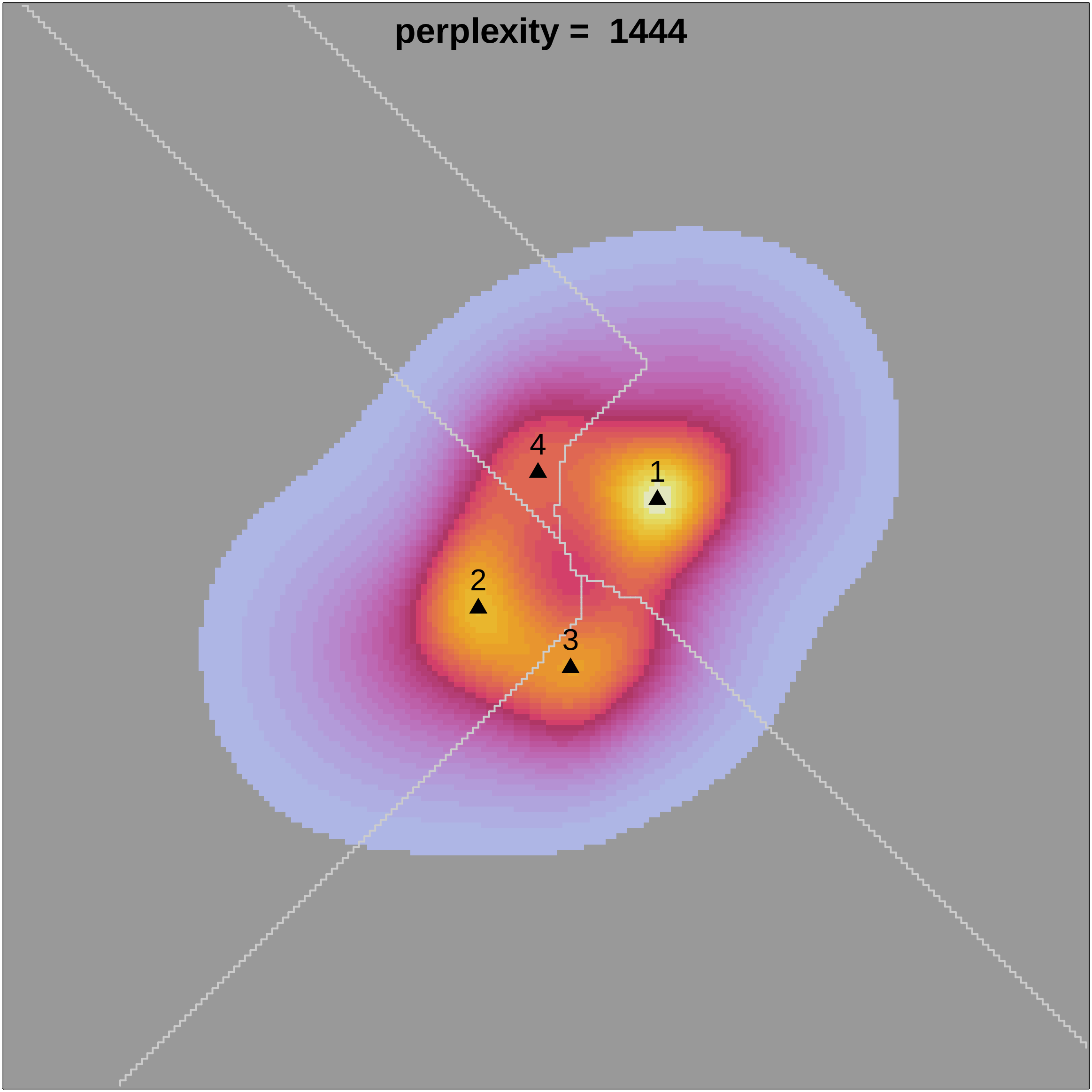

perplexity = 1444
1
2
3
4

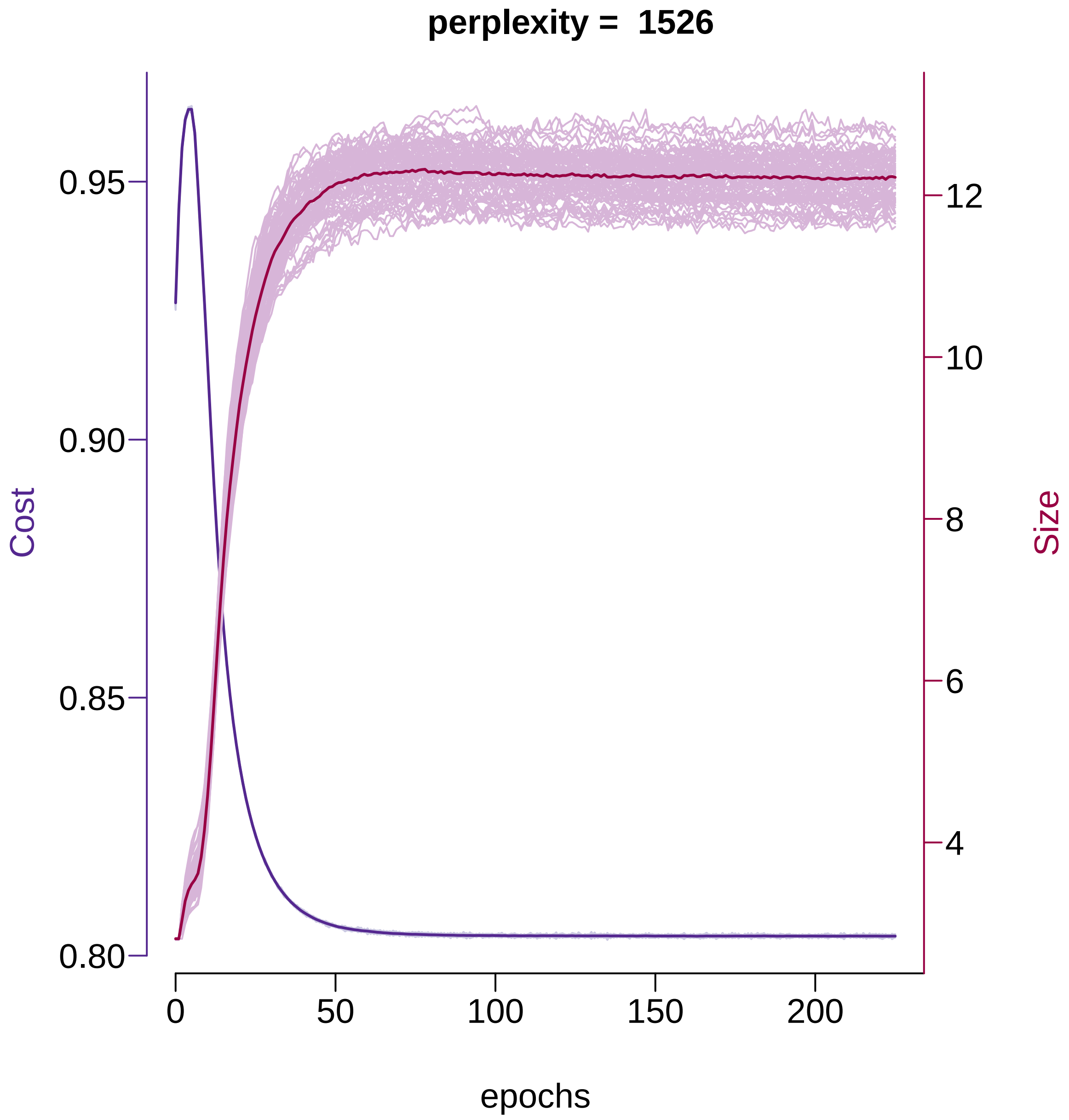
perplexity = 1526
Cost
0.95
0.90
0.85
0.80
Size
12
10
8
6
4
0
50
100
150
200
epochs

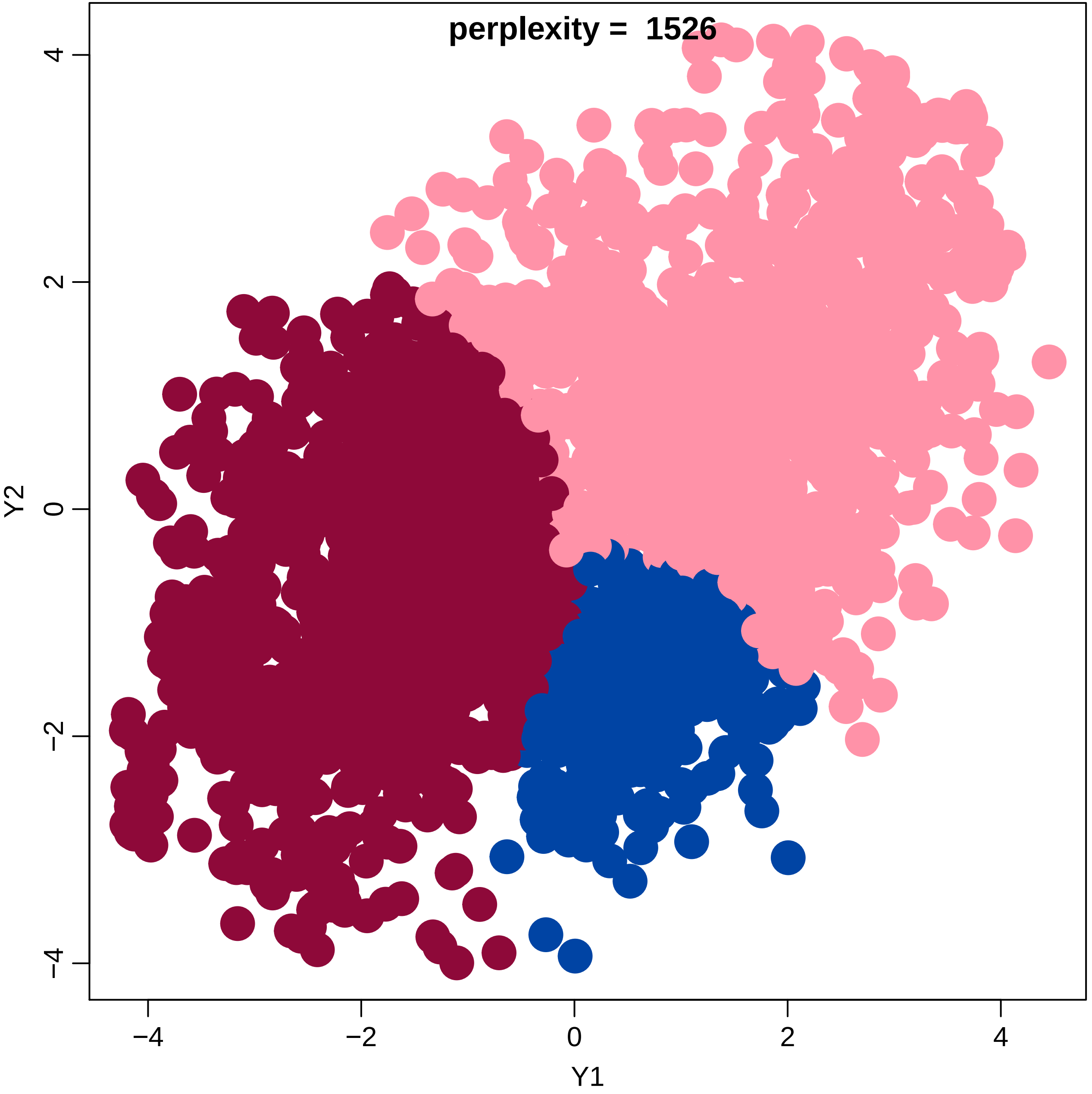

perplexity = 1526
Y2
Y1
4
2
0
−2
−4
−4
−2
0
2
4

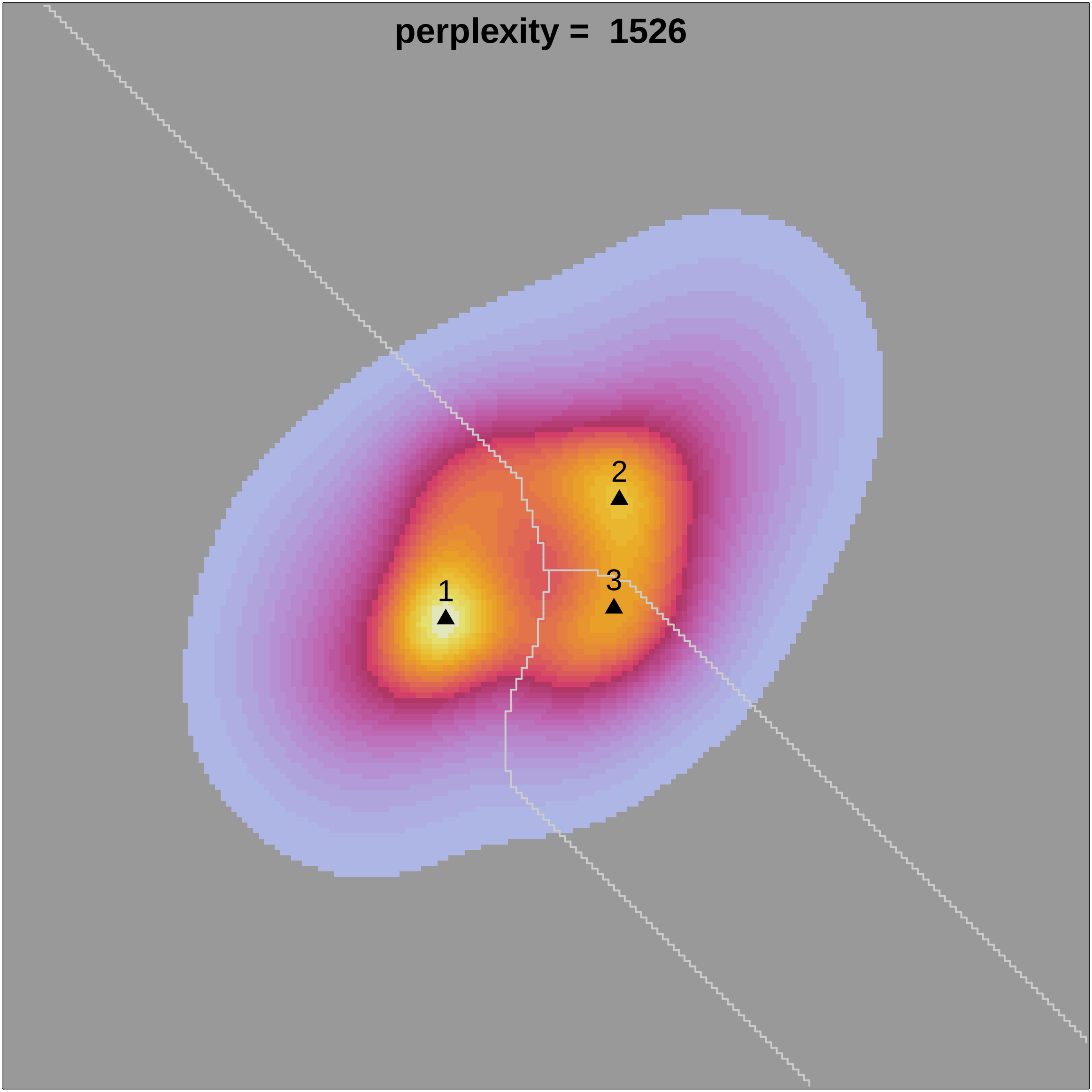
perplexity = 1526
1
2
3

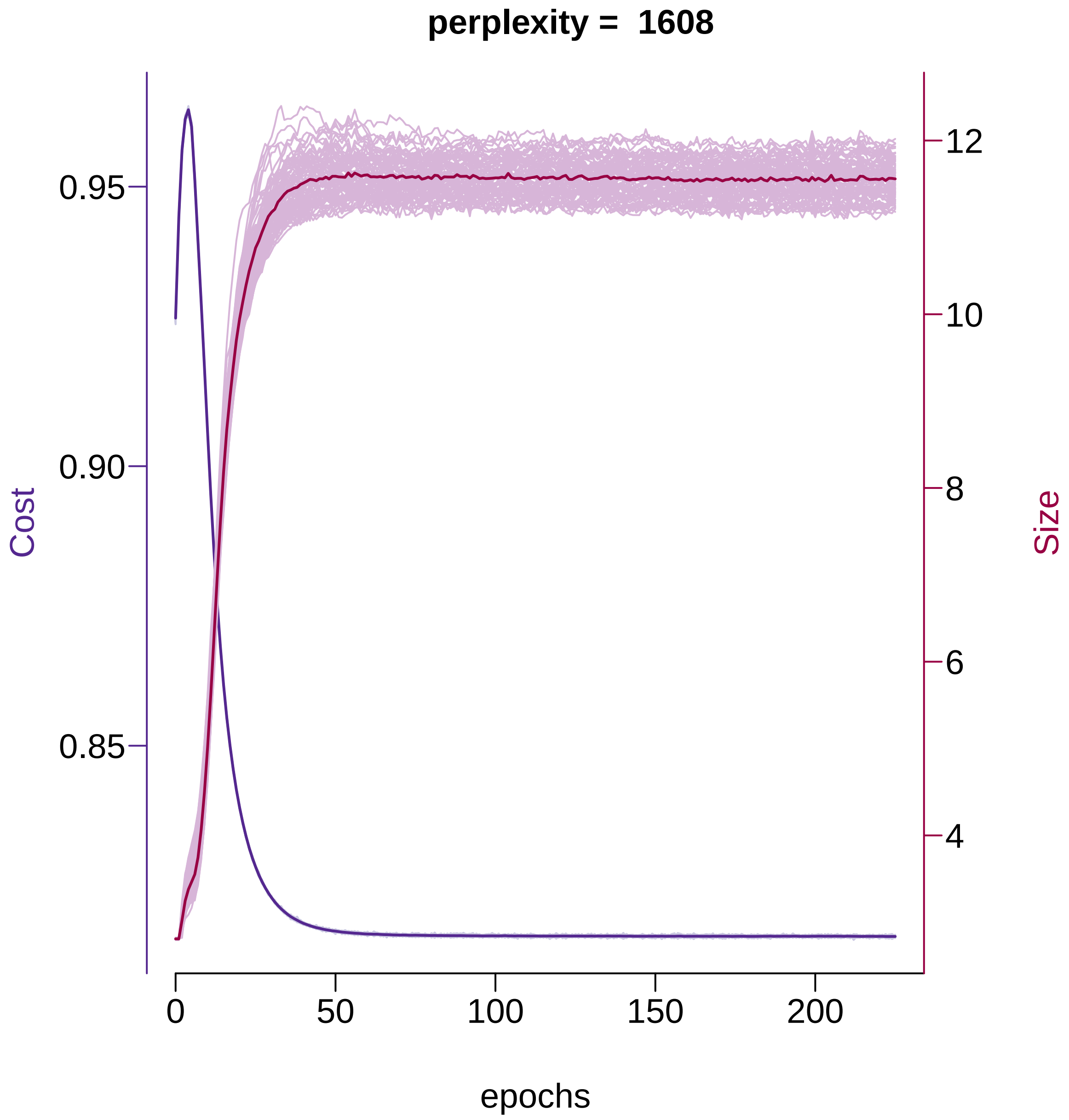
perplexity = 1608
Cost
0.95
0.90
0.85
Size
12
10
8
6
4
0
50
100
150
200
epochs

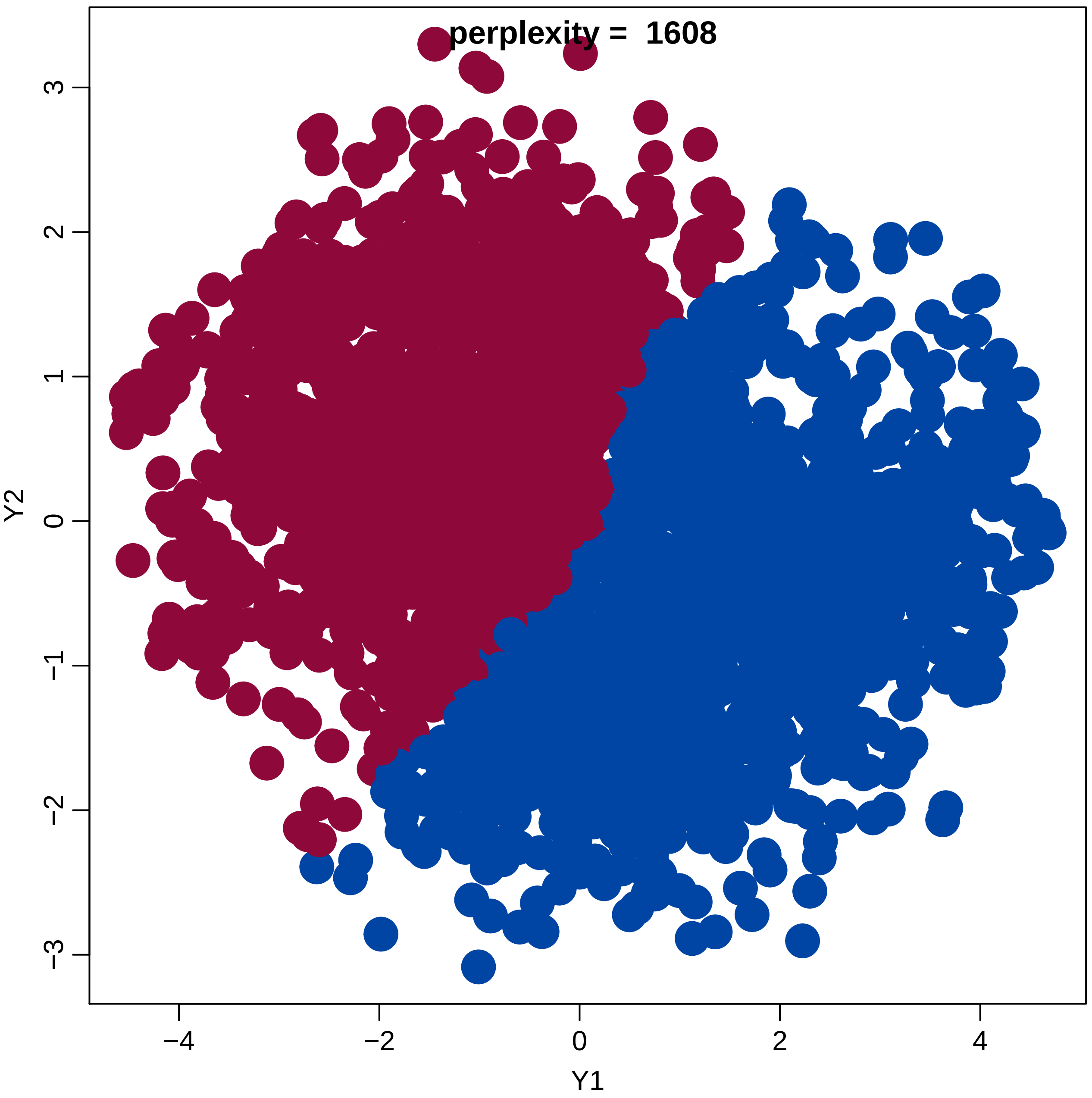
perplexity = 1608
Y2
Y1
3
2
1
0
−1
−2
−3
−4
−2
0
2
4

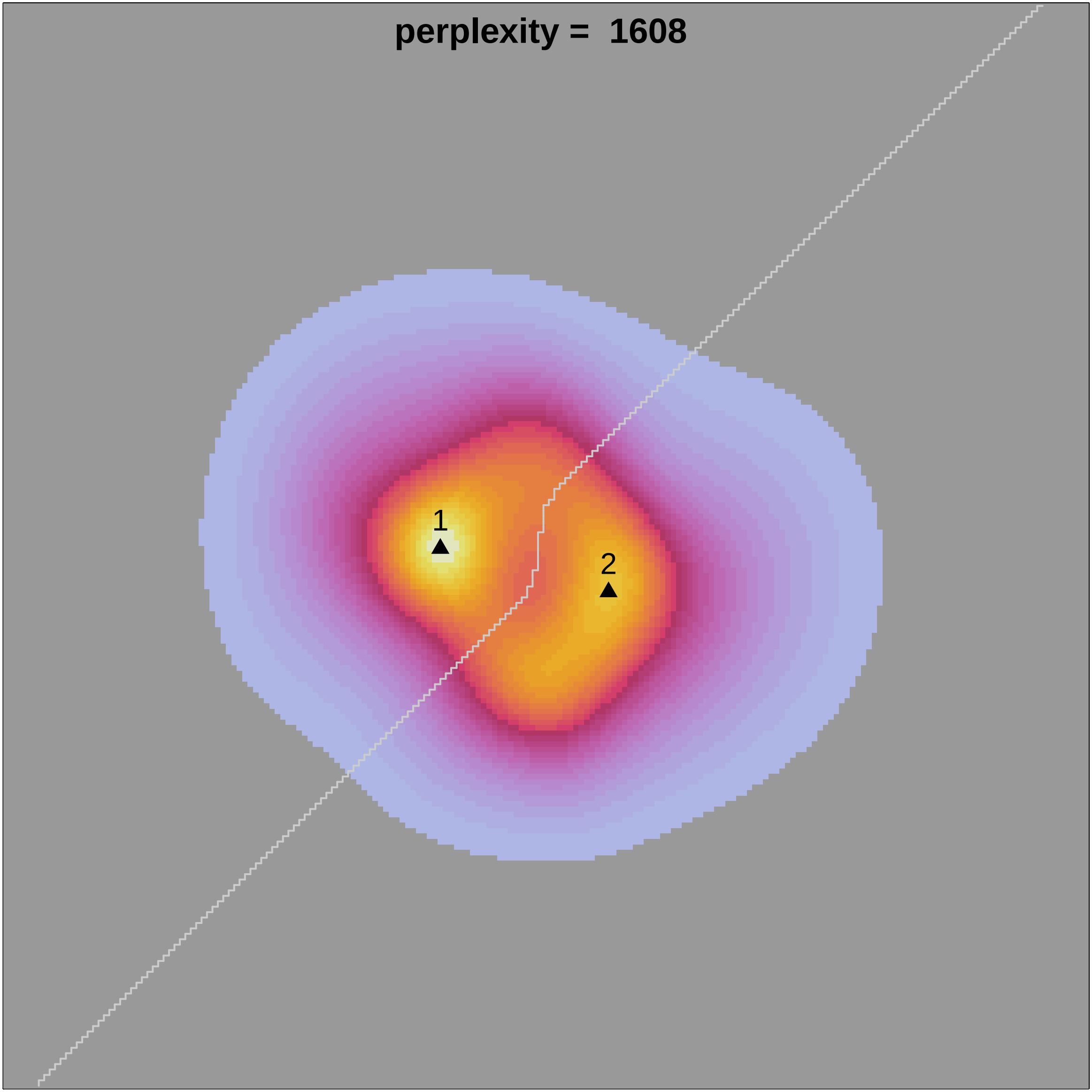

perplexity = 1608
1
2

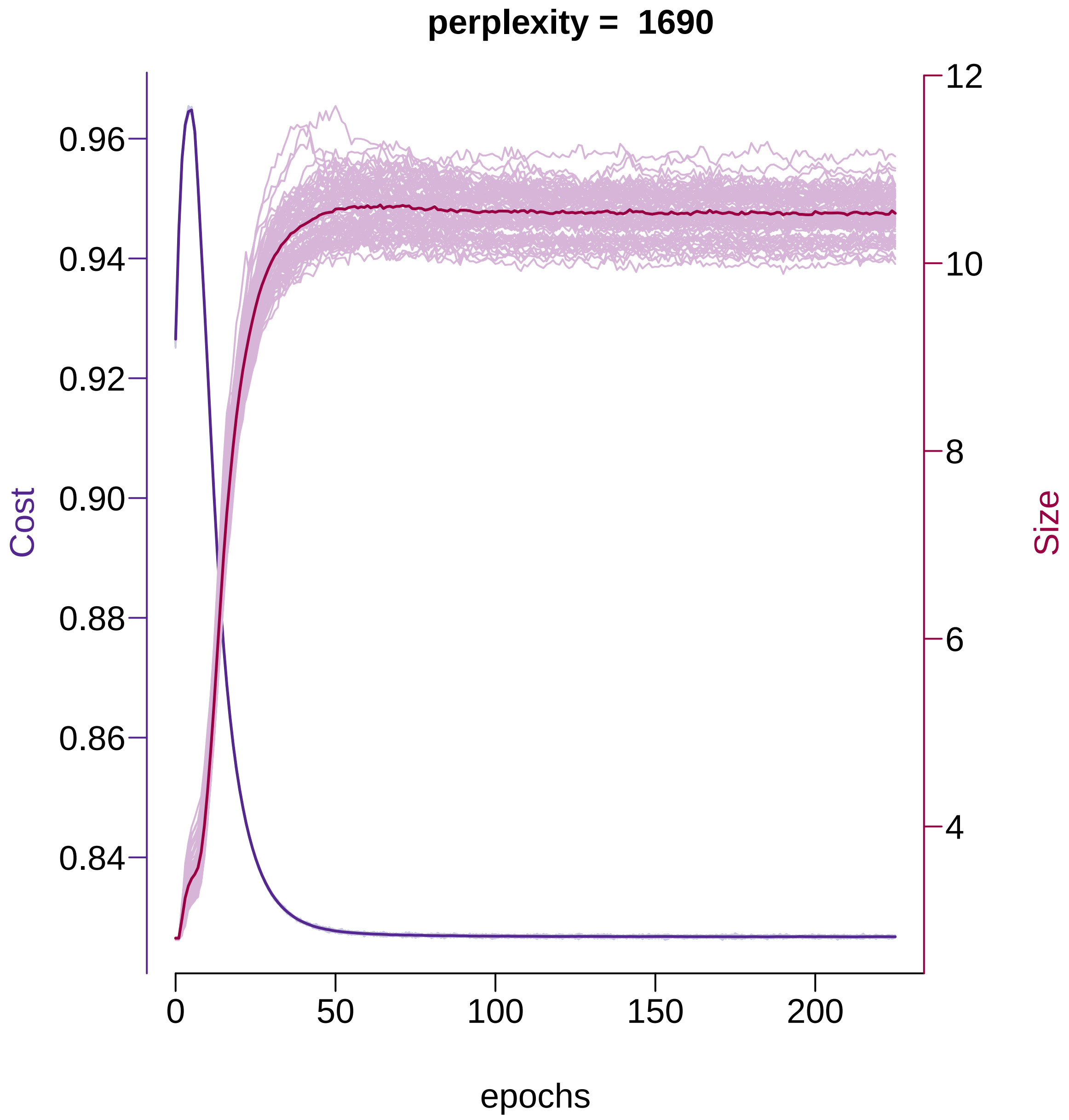
perplexity = 1690
Cost
0.96
0.94
0.92
0.90
0.88
0.86
0.84
Size
12
10
8
6
4
0
50
100
150
200
epochs

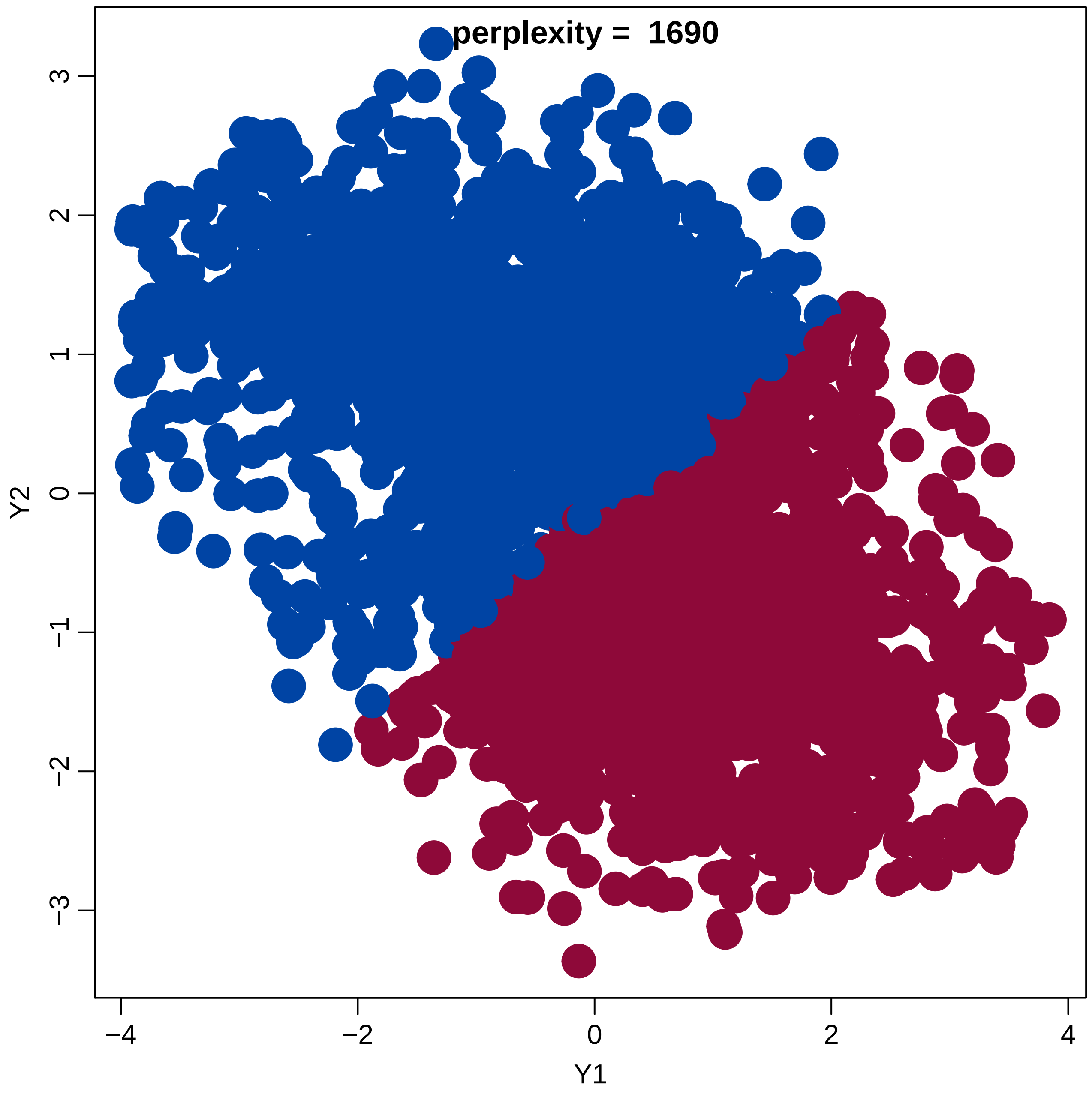
perplexity = 1690
Y1
Y2
−4
−2
0
2
4
−3
−2
−1
0
1
2
3

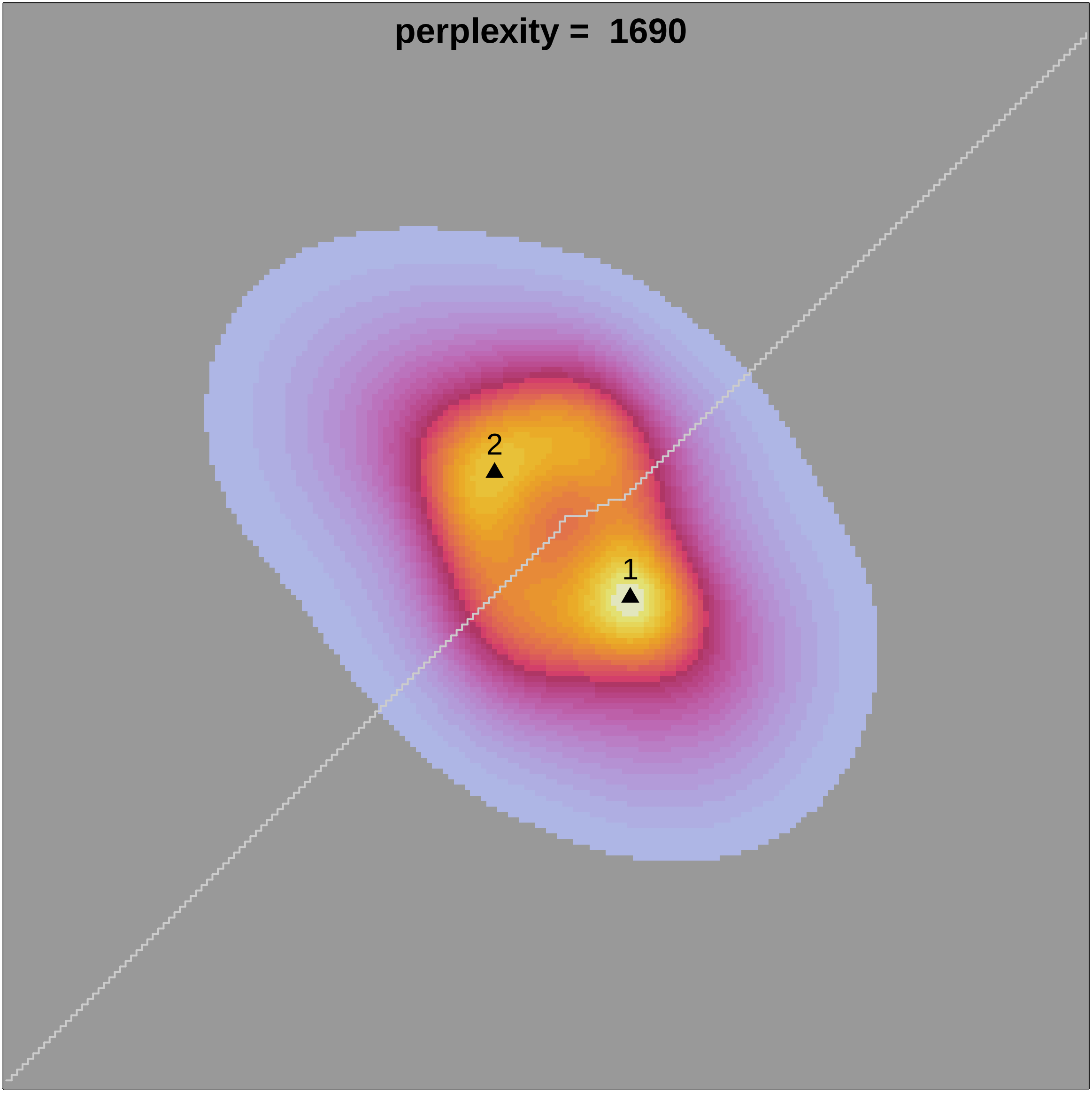
perplexity = 1690
2
1

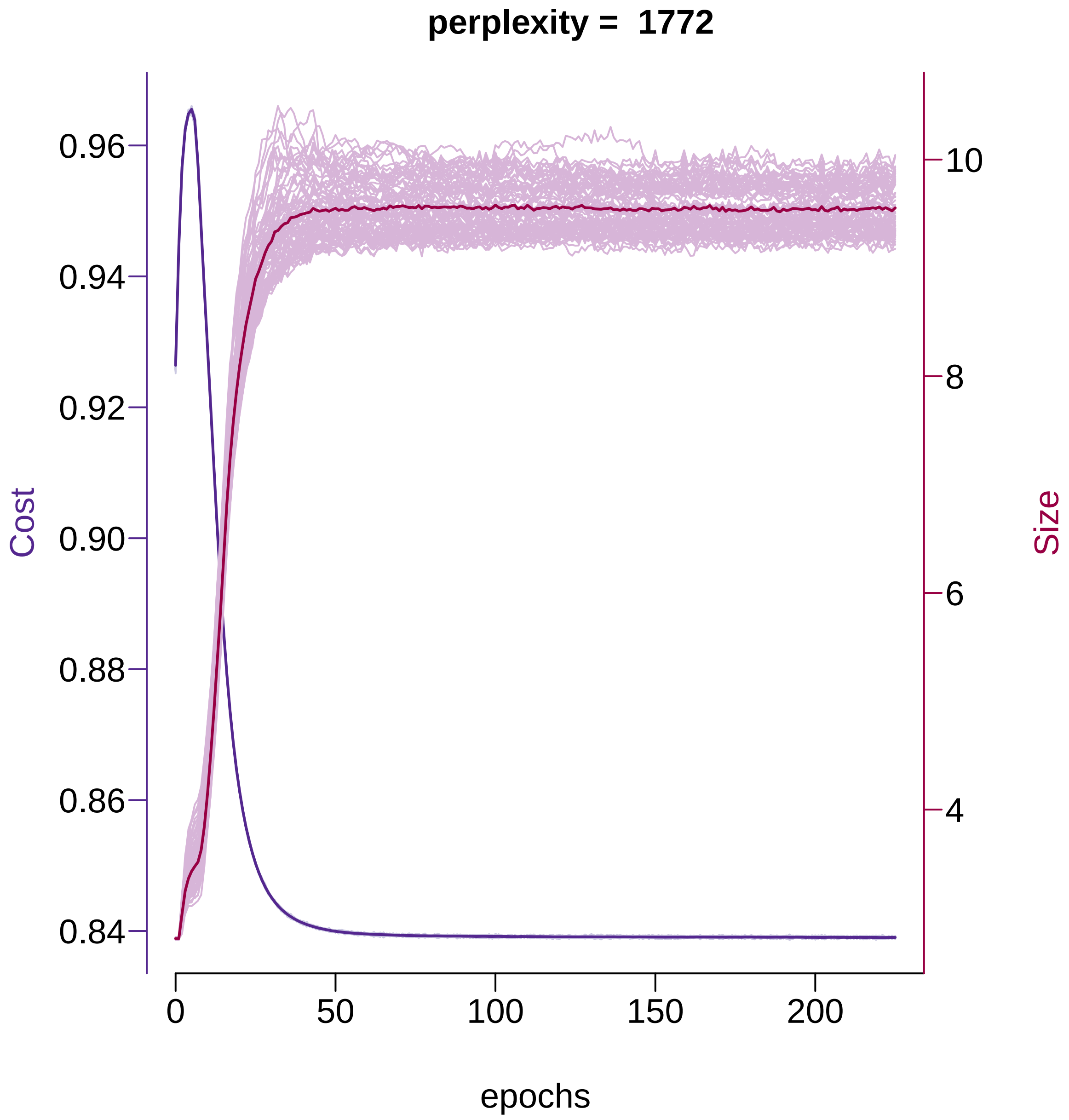
perplexity = 1772
Cost
Size
epochs
0.96
0.94
0.92
0.90
0.88
0.86
0.84
10
8
6
4
0
50
100
150
200

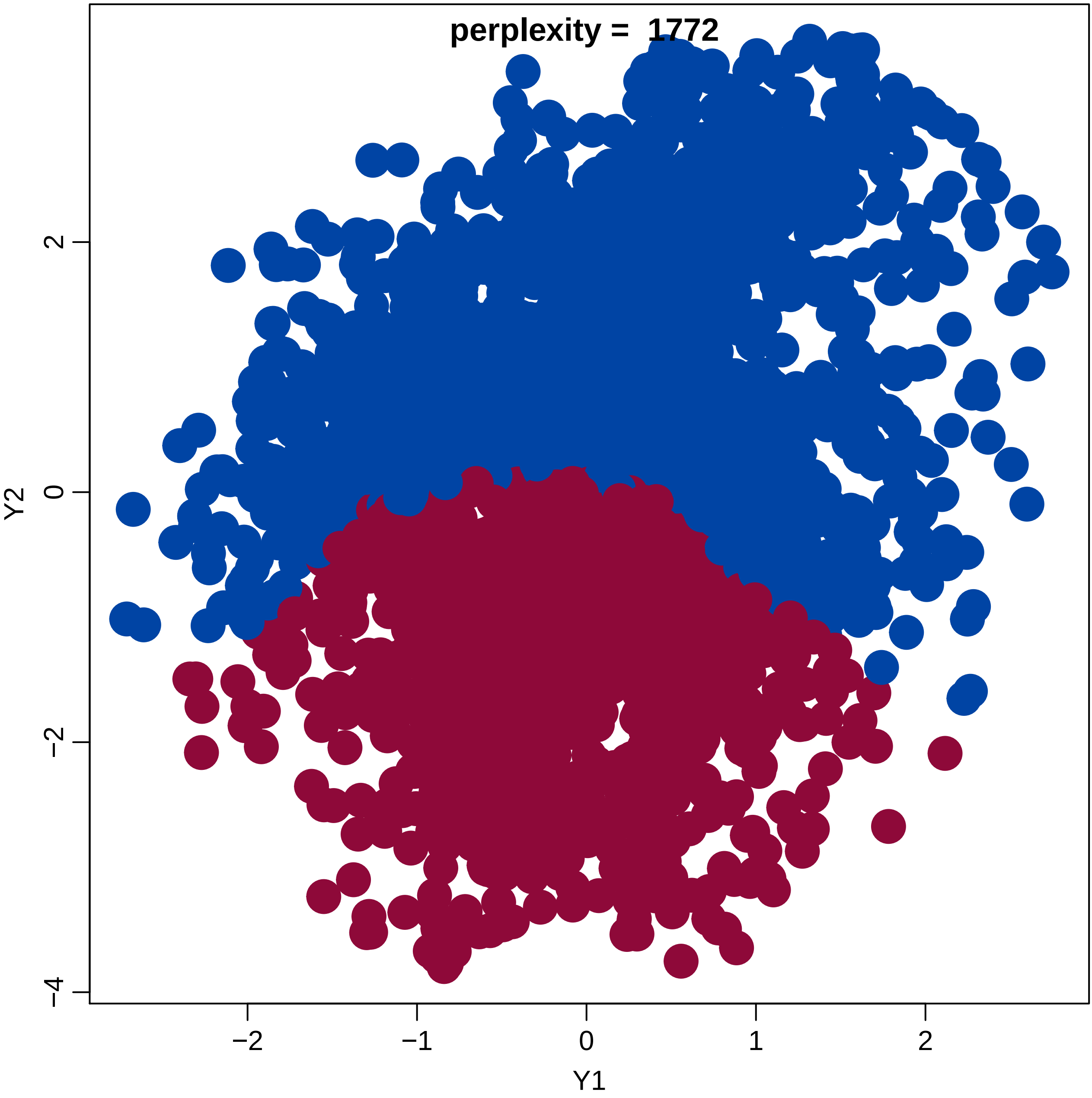
perplexity = 1772
Y2
Y1
2
0
−2
−4
−2
−1
0
1
2

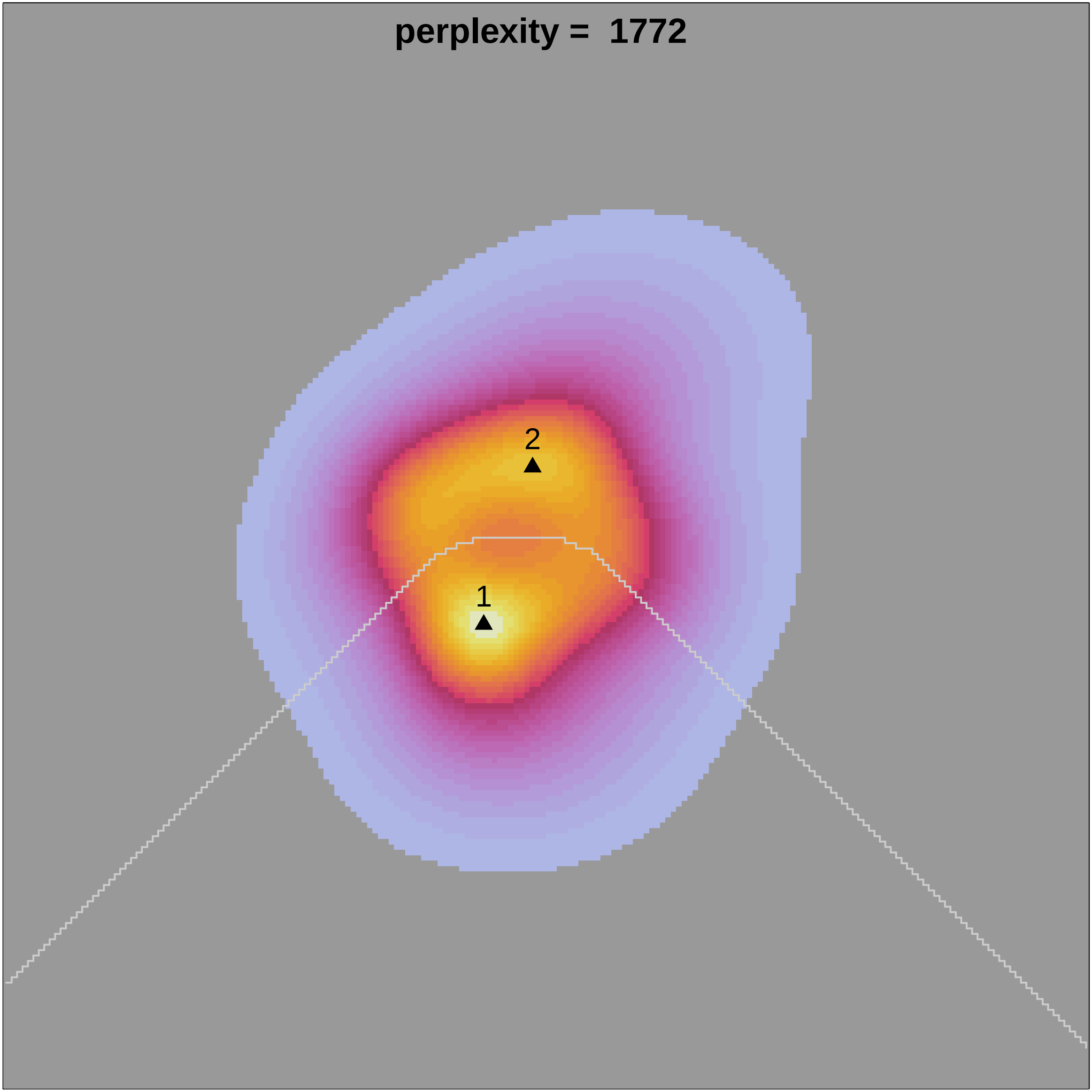
perplexity =  1772
2
1

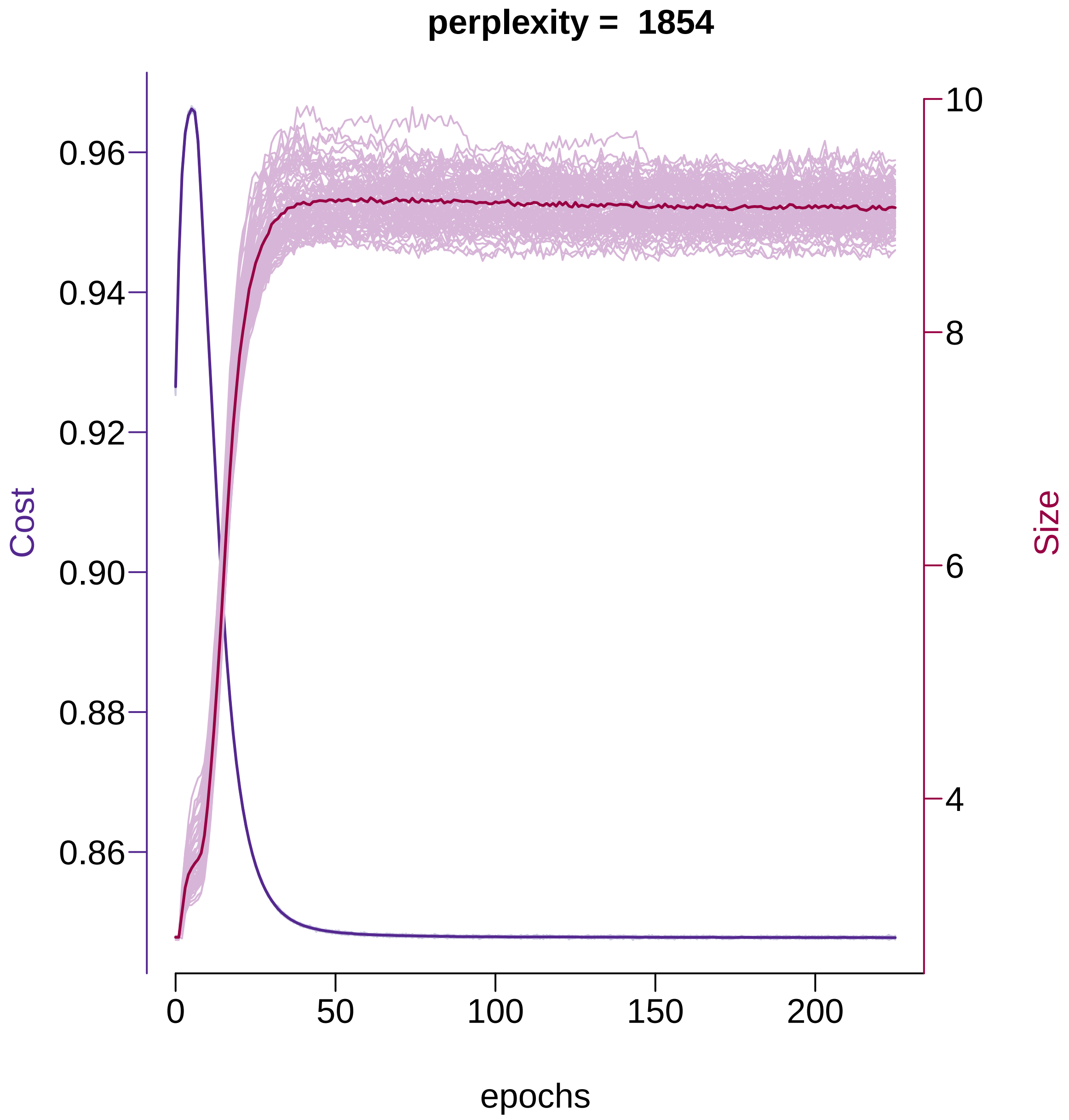
perplexity = 1854
Cost
0.96
0.94
0.92
0.90
0.88
0.86
Size
10
8
6
4
0
50
100
150
200
epochs

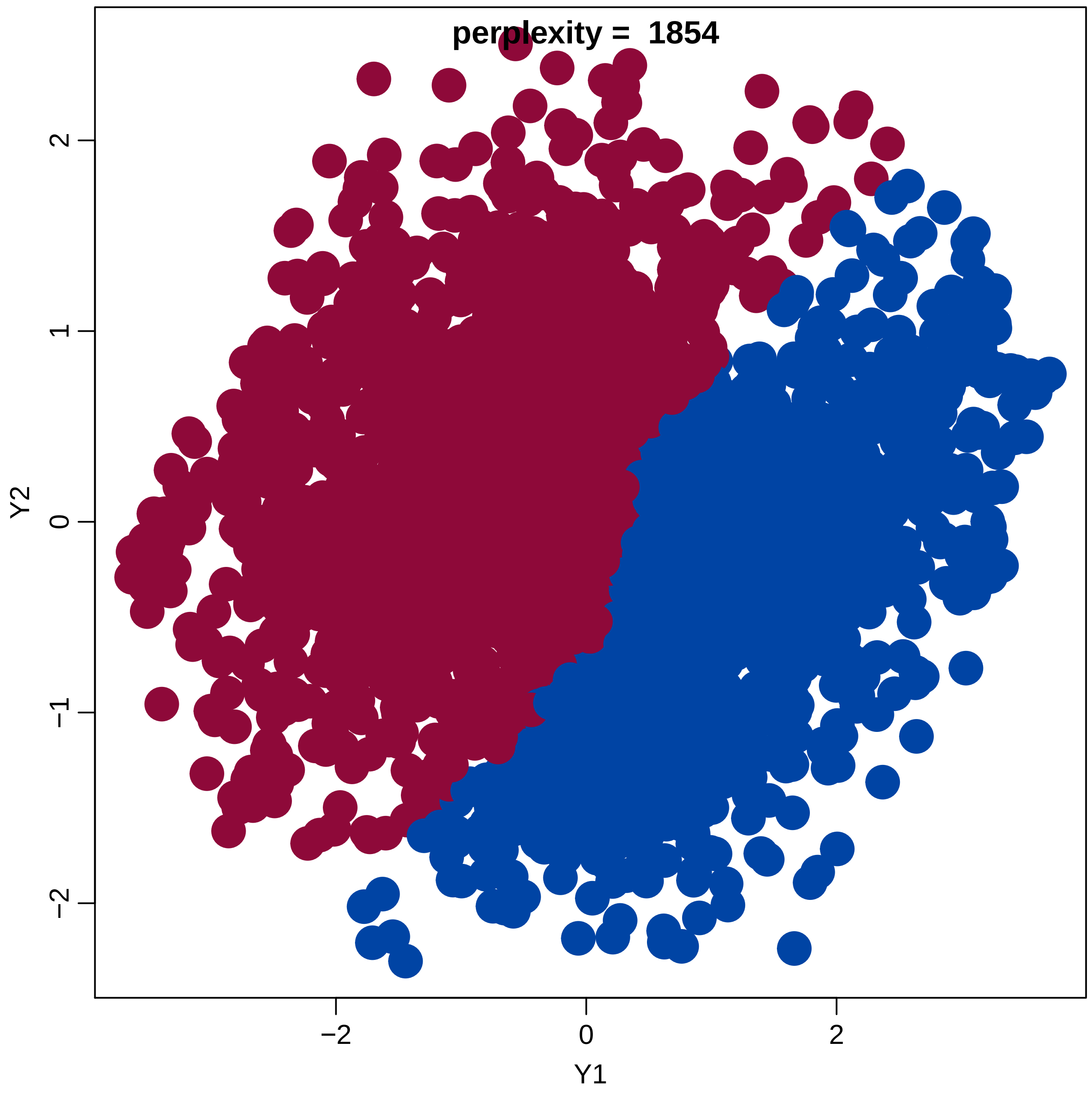

perplexity = 1854
Y2
2
1
0
−1
−2
Y1
−2
0
2

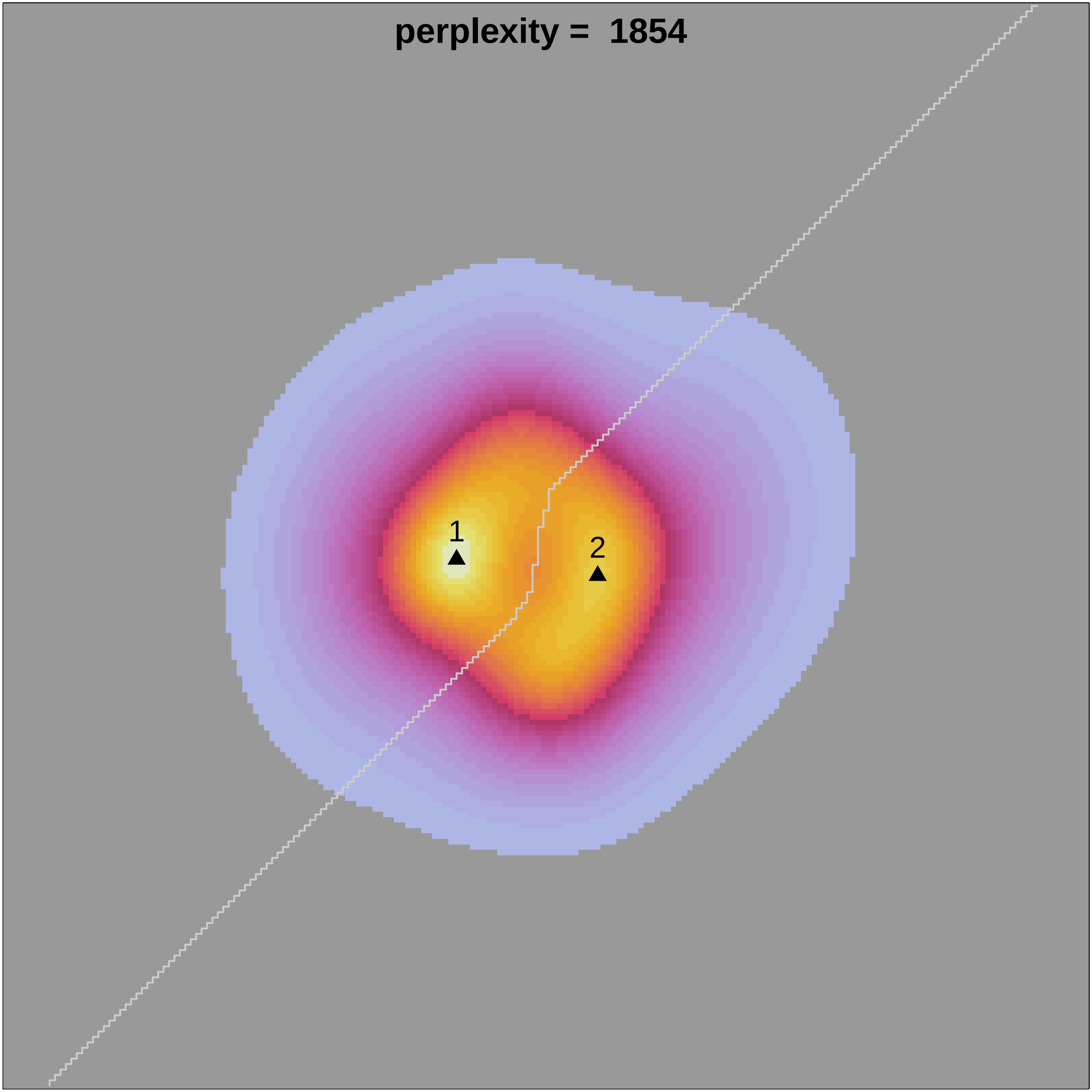
perplexity = 1854
1
2

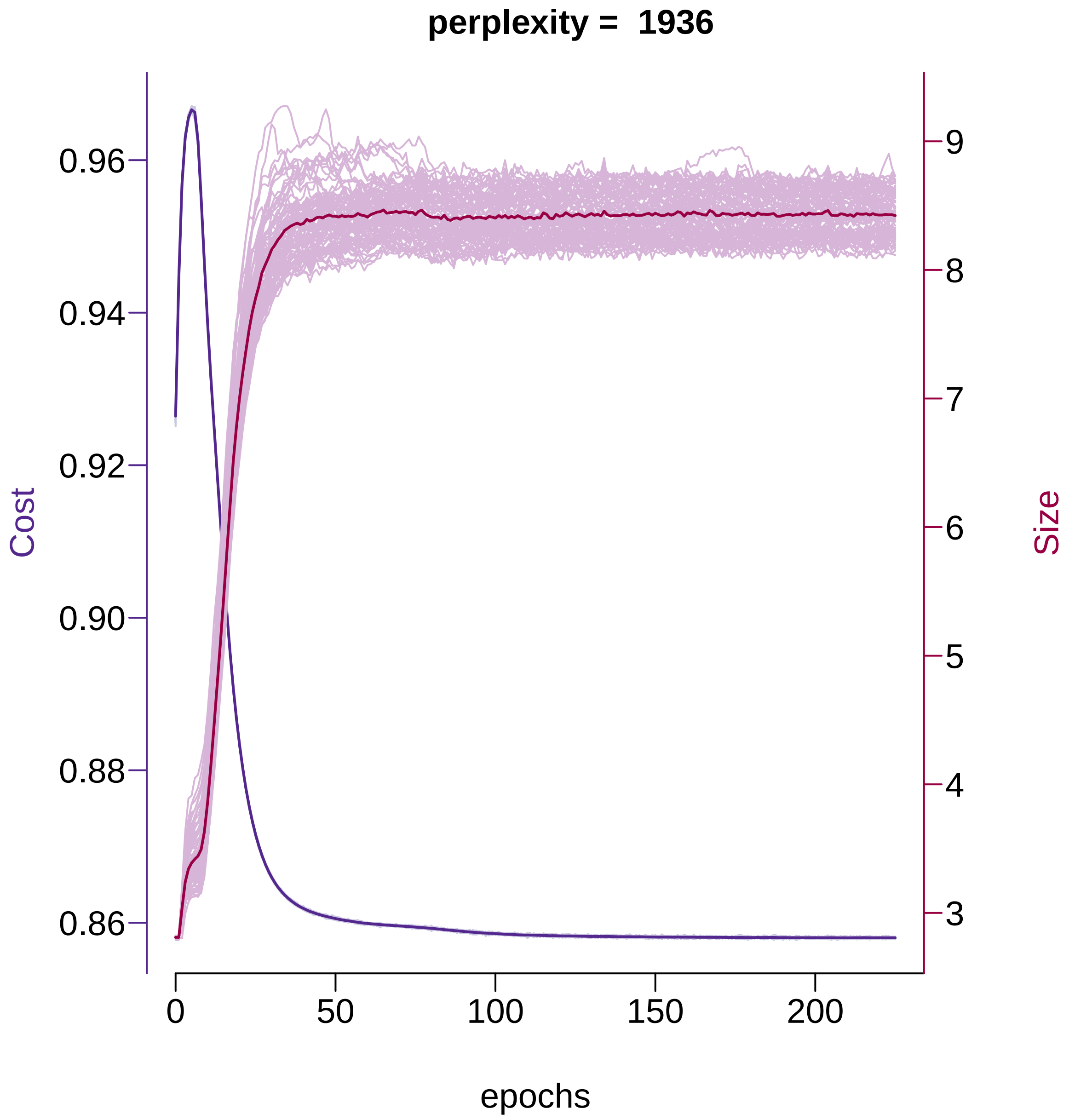

perplexity = 1936
Cost
0.96
0.94
0.92
0.90
0.88
0.86
Size
9
8
7
6
5
4
3
0
50
100
150
200
epochs

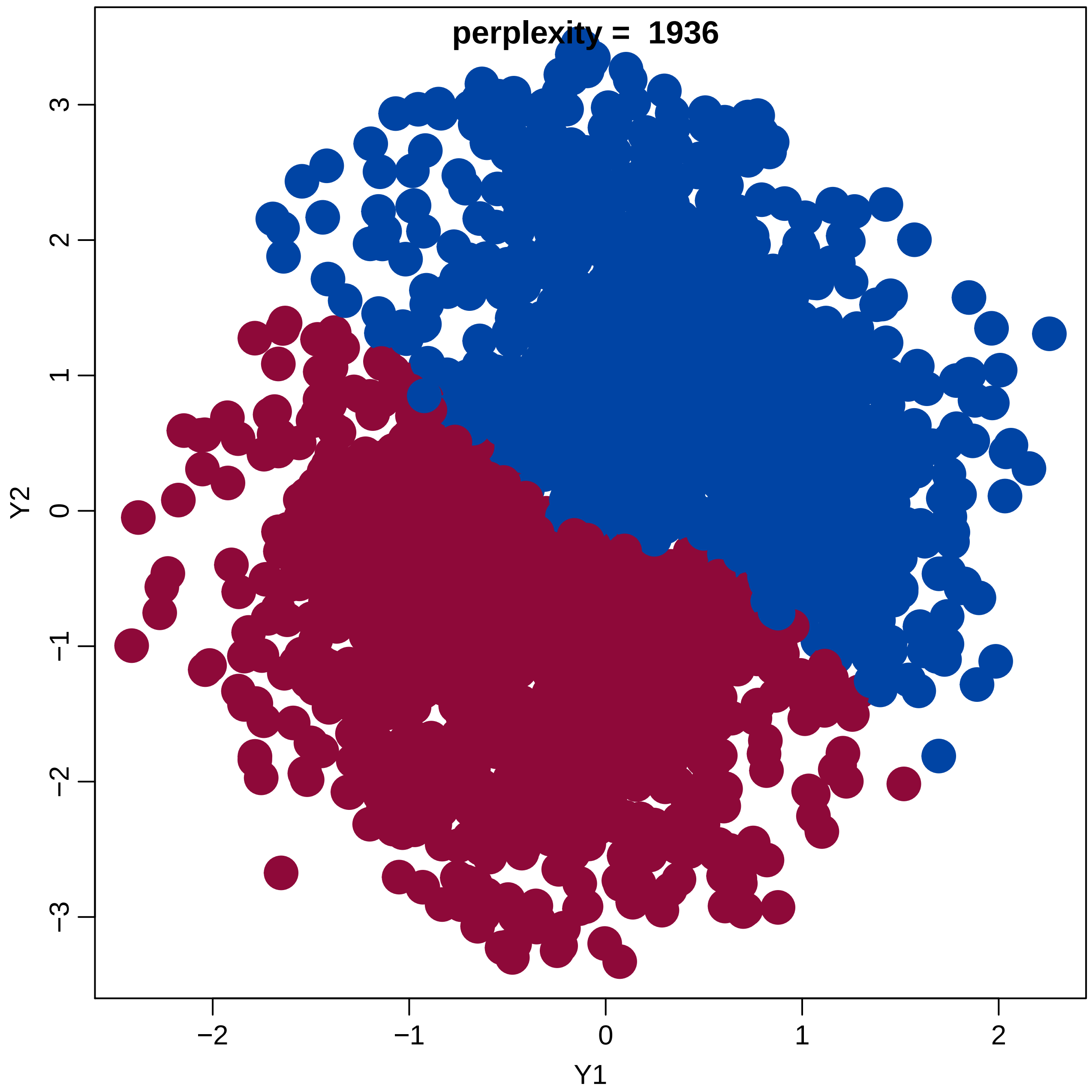
perplexity = 1936
Y2
Y1
−3
−2
−1
0
1
2
3
−2
−1
0
1
2

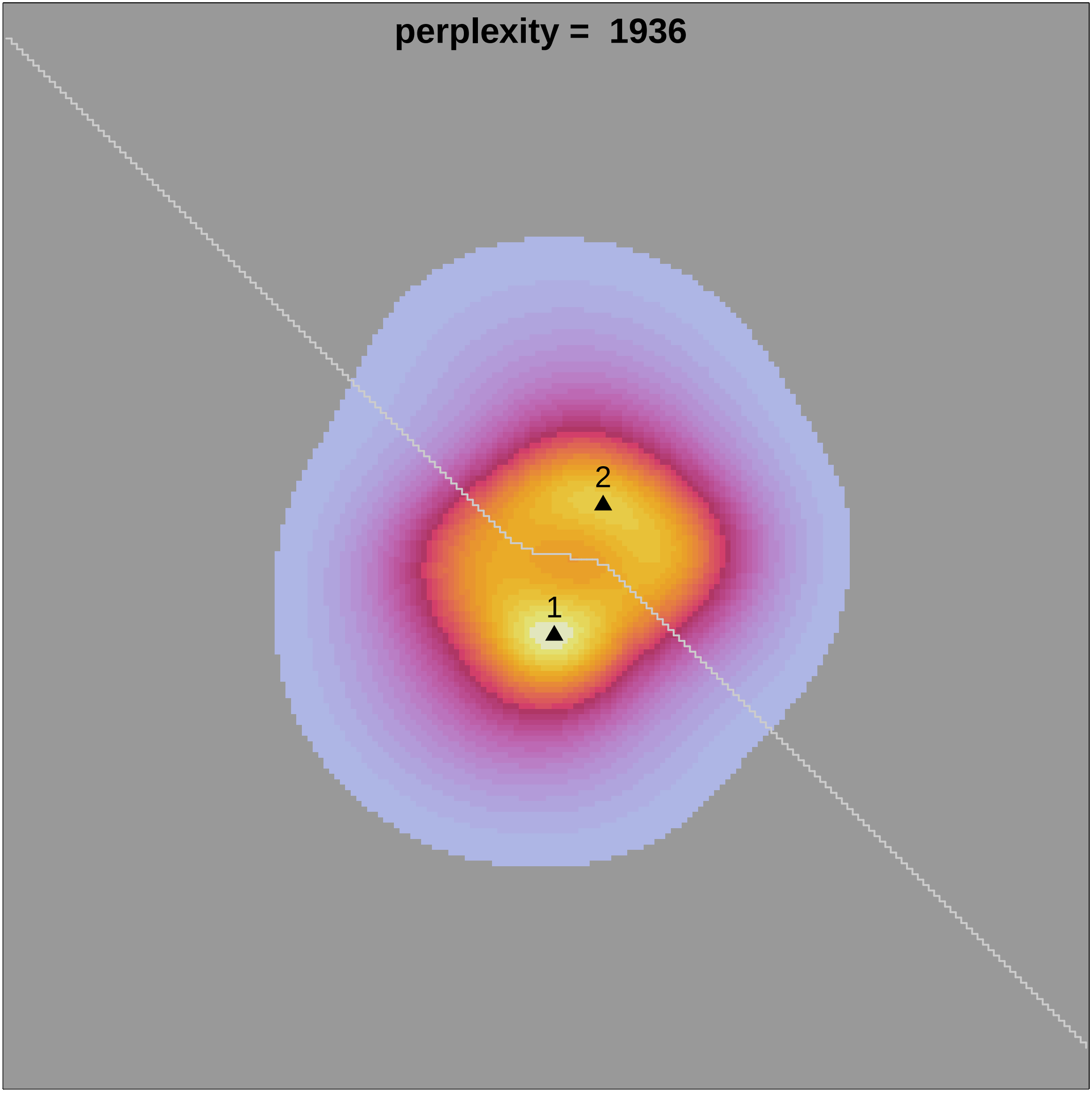
perplexity = 1936
2
1

**perplexity = 2018**

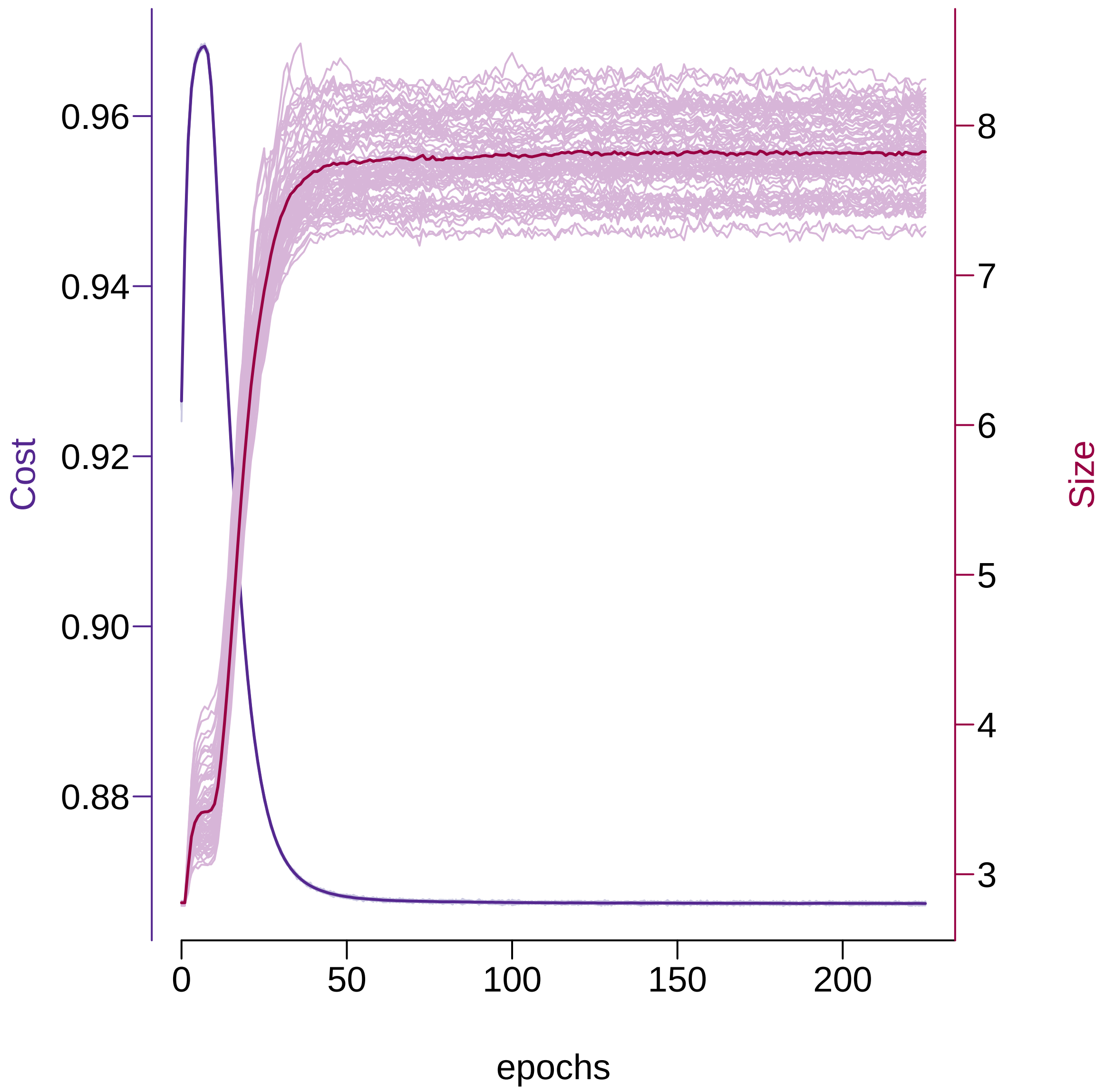

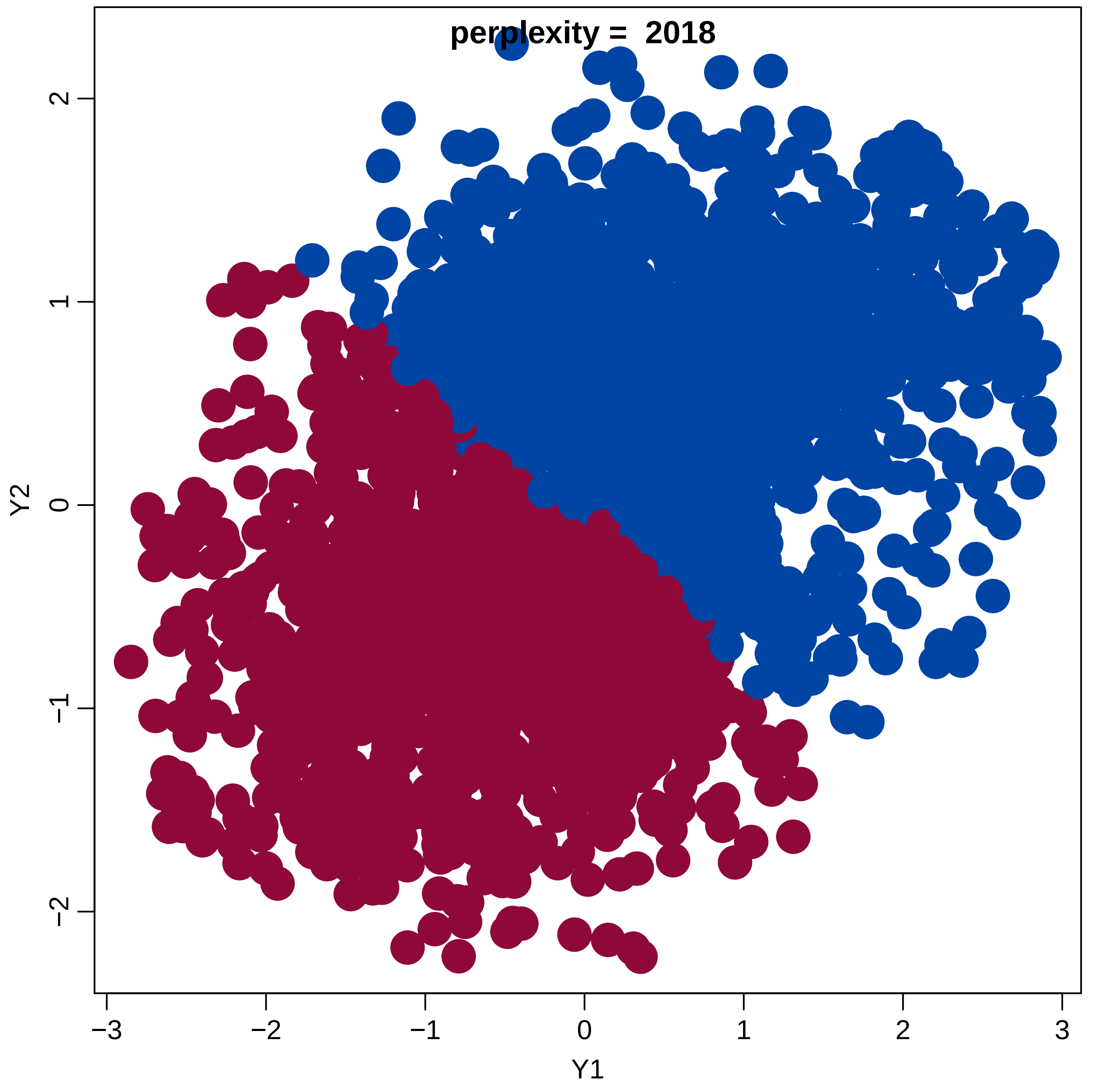

perplexity = 2018
Y2
2
1
0
−1
−2
−3
−2
−1
0
1
2
3
Y1

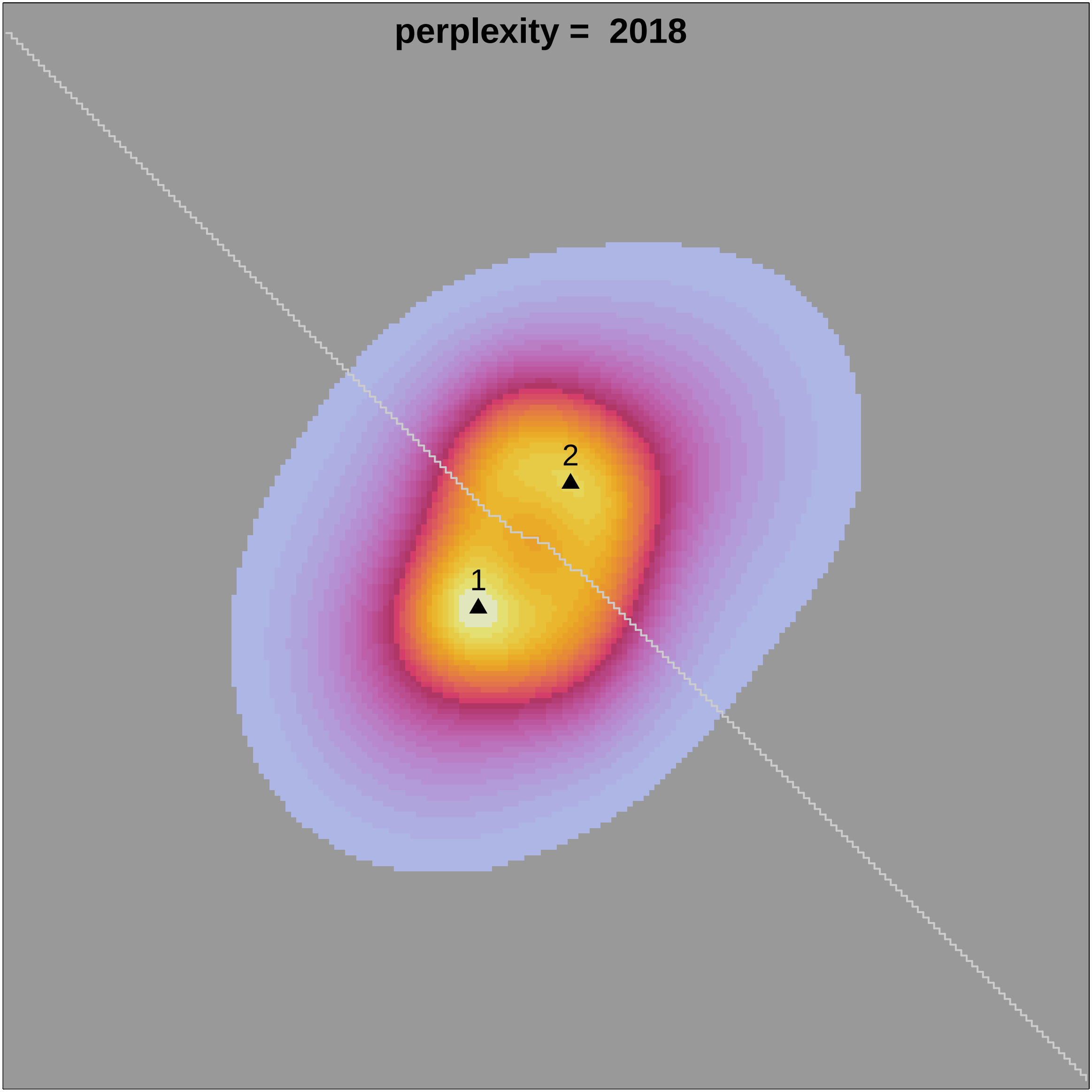
perplexity = 2018
1
2

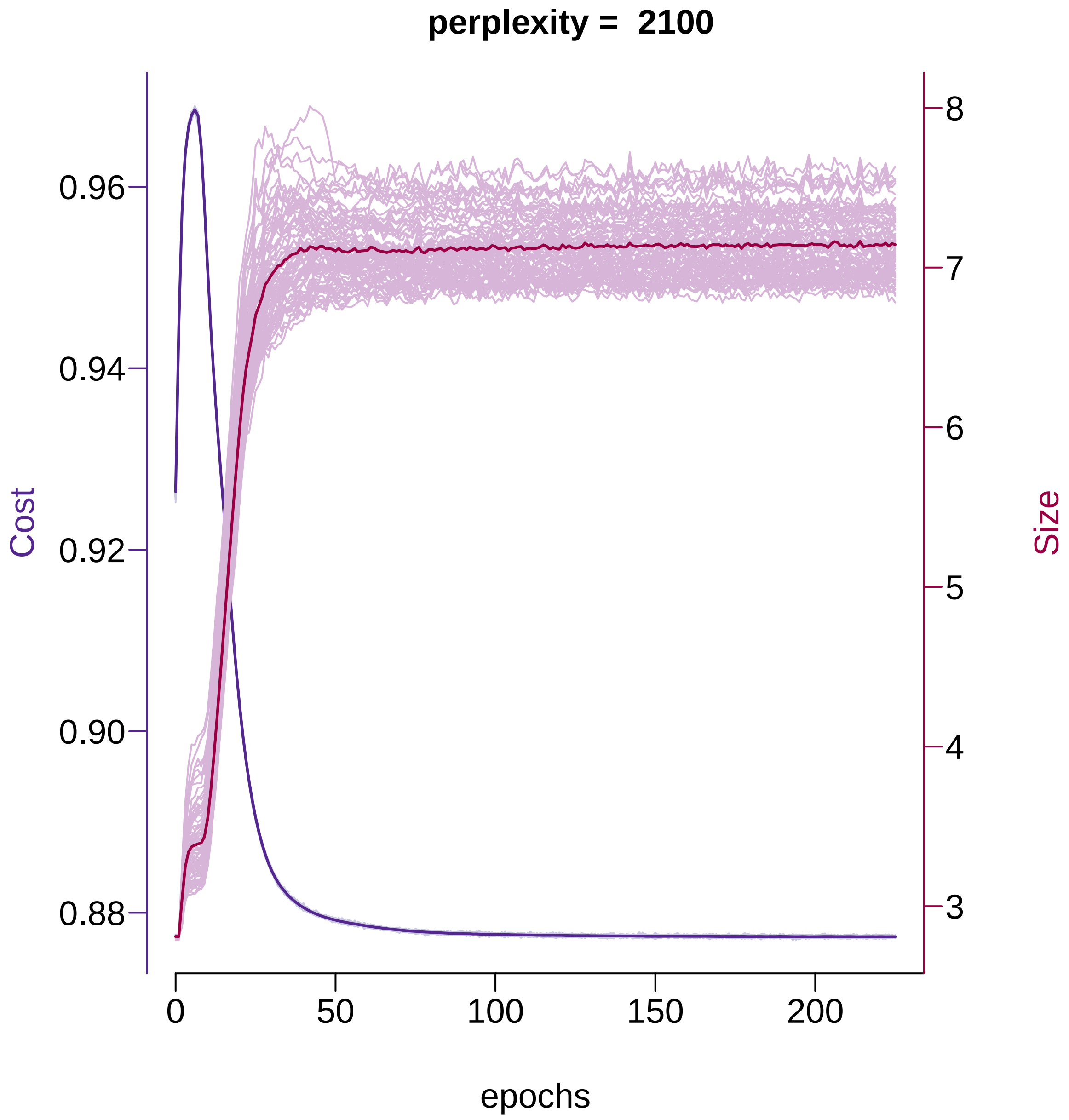

perplexity = 2100
Cost
Size
0.96
0.94
0.92
0.90
0.88
8
7
6
5
4
3
0
50
100
150
200
epochs

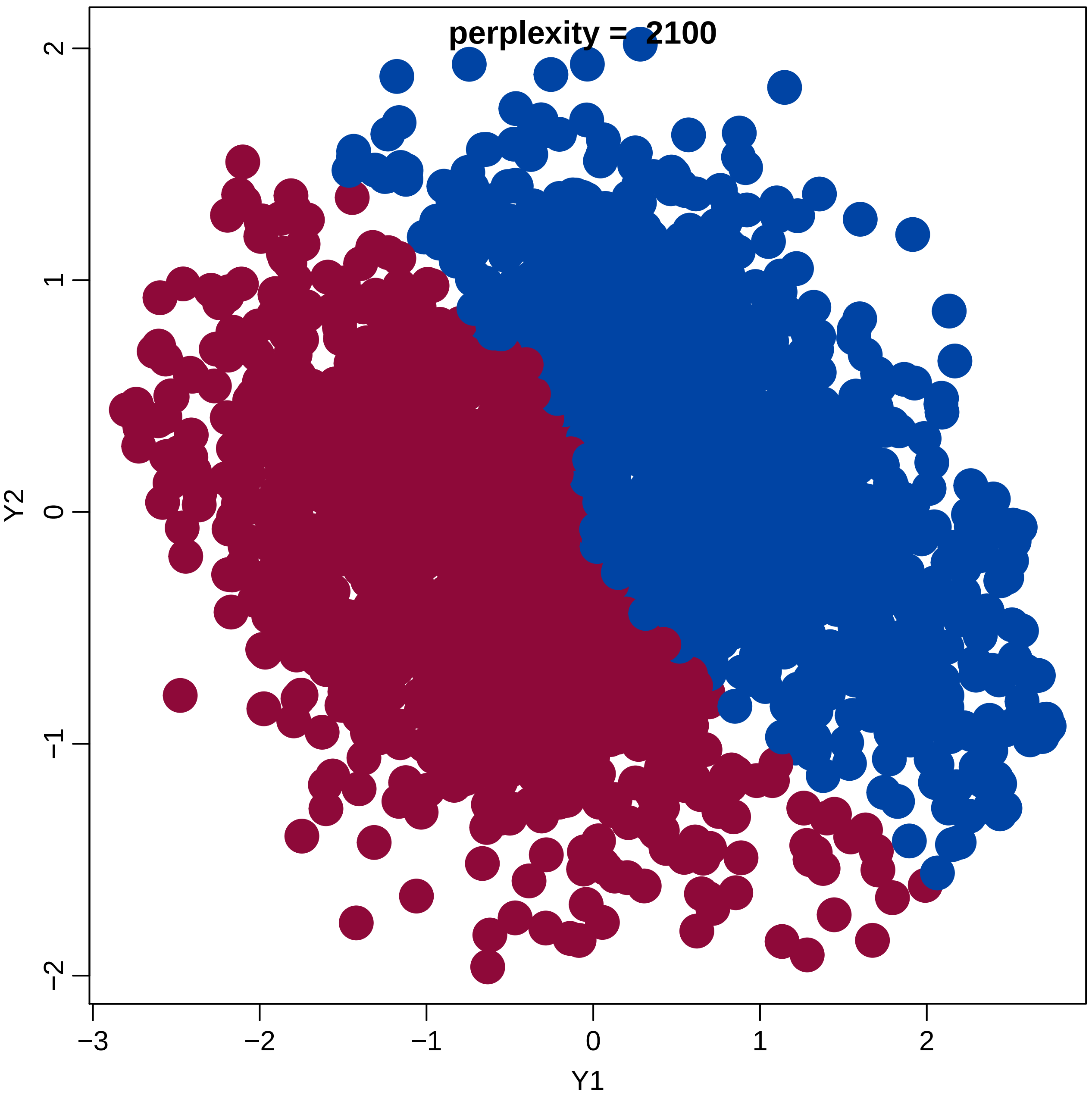
perplexity = 2100
Y1
Y2
−3
−2
−1
0
1
2
−2
−1
0
1
2

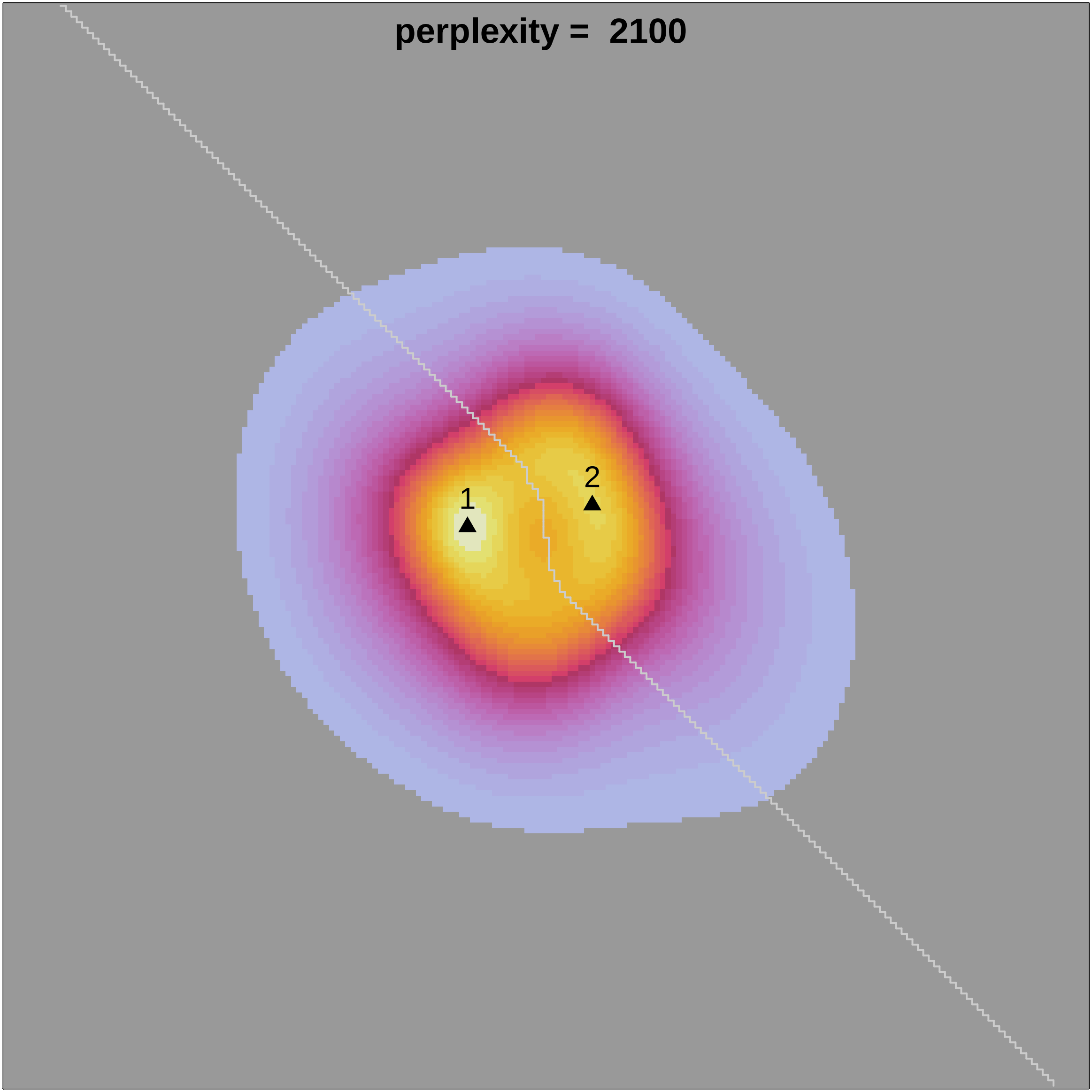
perplexity = 2100
1
2
